## Supplemental Material for "Synthesis and Biological Evaluation of New Isoxazolyl Steroids as Anti-Prostate Cancer Agents"

**Table S1**. AR transcriptional and antiproliferative activities of novel derivatives.

| Cmp. | AR transcriptional activity (%) | | | | | | Viability after 72 h ($GI_{50}$)[c] | | |
|---|---|---|---|---|---|---|---|---|---|
| | ANTAGONIST MODE[a] | | | AGONIST MODE[b] | | | | | |
| | 50 μM | 10 μM | 2 μM | 50 μM | 10 μM | 2 μM | LNCaP | LAPC-4 | DU145 |
| **24d** | 103.6 ± 42.6 | 112.8 ± 21.1 | 109.0 ± 12.1 | 18.6 ± 3.6 | 21.1 ± 1.6 | 17.8 ± 0.1 | > 50 | > 50 | > 50 |
| **24e** | 125.3 ± 27.4 | 101.1 ± 13.6 | 101.3 ± 9.5 | 12.8 ± 0.2 | 14.2 ± 1.4 | 14.7 ± 0.5 | > 50 | > 50 | > 50 |
| **24g** | 111.6 ± 19.9 | 112.5 ± 14.5 | 112.9 ± 10.9 | 19.7 ± 1.6 | 17.5 ± 0.5 | 15.6 ± 0.6 | > 50 | > 50 | > 50 |
| **24j** | 21.6 ± 7.8 | 84.8 ± 17.4 | 100.2 ± 13.5 | 6.1 ± 1.5 | 12.6 ± 0.1 | 14.4 ± 2.8 | 25.8 ± 0.6 | 18.2 ± 1.2 | > 50 |
| **27** | 63.5 ± 8.0 | 62.0 ± 14.8 | 89.7 ± 9.9 | 20.4 ± 4.2 | 20.9 ± 2.9 | 17.4 ± 0.5 | > 50 | > 50 | > 50 |
| **32** | 31.1 ± 13.3 | 87.8 ± 20.9 | 99.7 ± 17.4 | 14.6 ± 0.5 | 12.1 ± 2.7 | 14.0 ± 1.1 | 19.5 ± 0.1 | 18.9 ± 7.2 | > 50 |
| **36** | 67.4 ± 21.9 | 72.5 ± 15.7 | 101.1 ± 18.1 | 16.3 ± 1.7 | 12.0 ± 0.5 | 14.7 ± 0.2 | > 50 | 19.0 ± 7.8 | > 50 |
| **38** | 78.2 ± 24.9 | 92.6 ± 20.5 | 96.1 ± 11.2 | 7.3 ± 1.1 | 13.0 ± 0.4 | 14.6 ± 1.0 | > 50 | > 50 | > 50 |
| **41a** | 50.8 ± 18.8 | 73.7 ± 14.9 | 98.2 ± 17.5 | 12.1 ± 1.3 | 14.7 ± 0.1 | 14.5 ± 1.5 | > 50 | > 50 | > 50 |
| **Gal** | 3.0 ± 1.8 | 35.1 ± 3.3 | 65.2 ± 6.1 | 2.4 ± 1.1 | 10.8 ± 3.1 | 15.4 ± 0.4 | 46.8 ± 0.1 | 28.6 ± 0.6 | 47.6 ± 0.2 |

[a] measured in the presence of compound and 1 nM R1881 and normalized to signal of 1 nM R1881 = 100%, determined in duplicate and repeated twice, mean ± SD is presented.
[b] measured in the presence of compound only, normalized to signal of 1 nM R1881 = 100%, measured in duplicate and repeated twice, mean ± SD is presented.
[c] measured at least in duplicate, mean ± SD is presented.

**Table S2**. Raw data for **Figure 2. (A)** Transcriptional activity of AR measured in reporter cell line in both antagonist (competition with 1 nM R1881) and agonist (presence of compound alone) mode upon treatment with different concentration of **24j**.

| Concentration (μM) | AR transcriptional activity (%) after treatment with **24 j** (antagonist mode) | | | | AR transcriptional activity (%) after treatment with **24 j** (agonist mode) | | | |
|---|---|---|---|---|---|---|---|---|
| 60.00 | 20.55 | 20.45 | 15.39 | 15.98 | 6.48 | 8.33 | 7.15 | 5.97 |
| 20.00 | 55.89 | 57.74 | 58.97 | 51.32 | 14.35 | 16.25 | 14.56 | 15.38 |
| 6.67 | 78.82 | 77.07 | 74.29 | 83.26 | 16.97 | 15.48 | 17.44 | 20.21 |
| 2.22 | 88.25 | 93.78 | 91.50 | 88.12 | 15.89 | 17.90 | 21.24 | 19.34 |
| 0.74 | 95.70 | 97.95 | 95.17 | 95.70 | 17.18 | 19.44 | 20.73 | 22.58 |
| 0.25 | 98.38 | 91.23 | 93.41 | 98.15 | 18.16 | 19.80 | 20.78 | 20.68 |

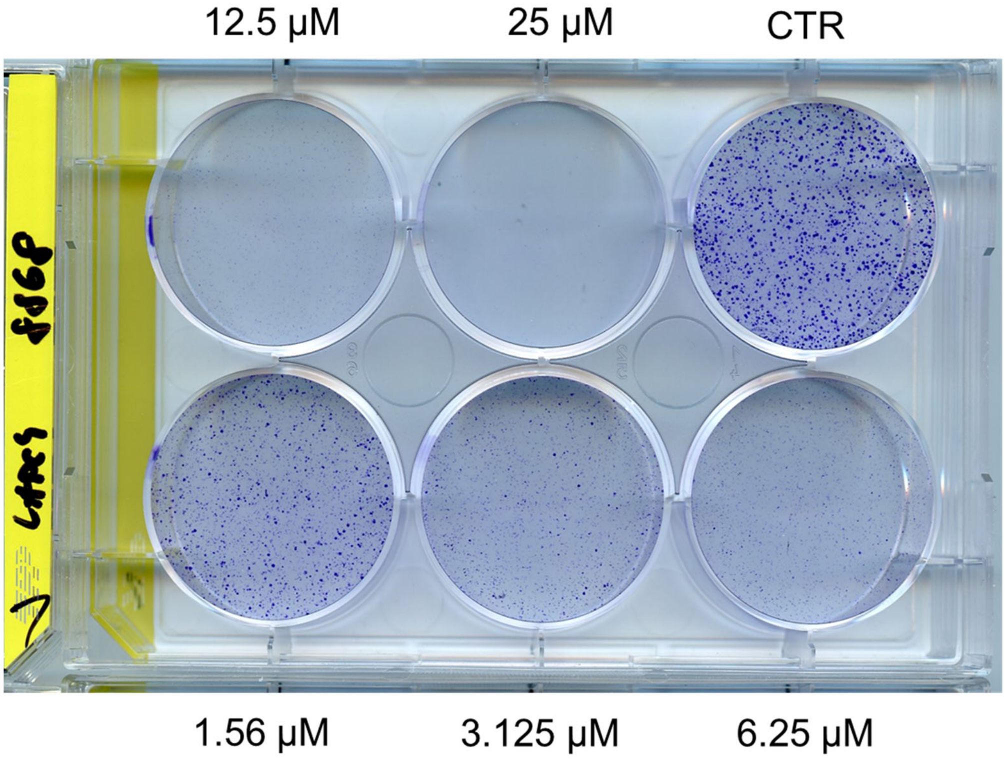


**Figure S1**. Raw picture for **Figure 2. (B)** Colony formation assay of PCa cells LAPC-4 after treatment with **24j** for 10 days.

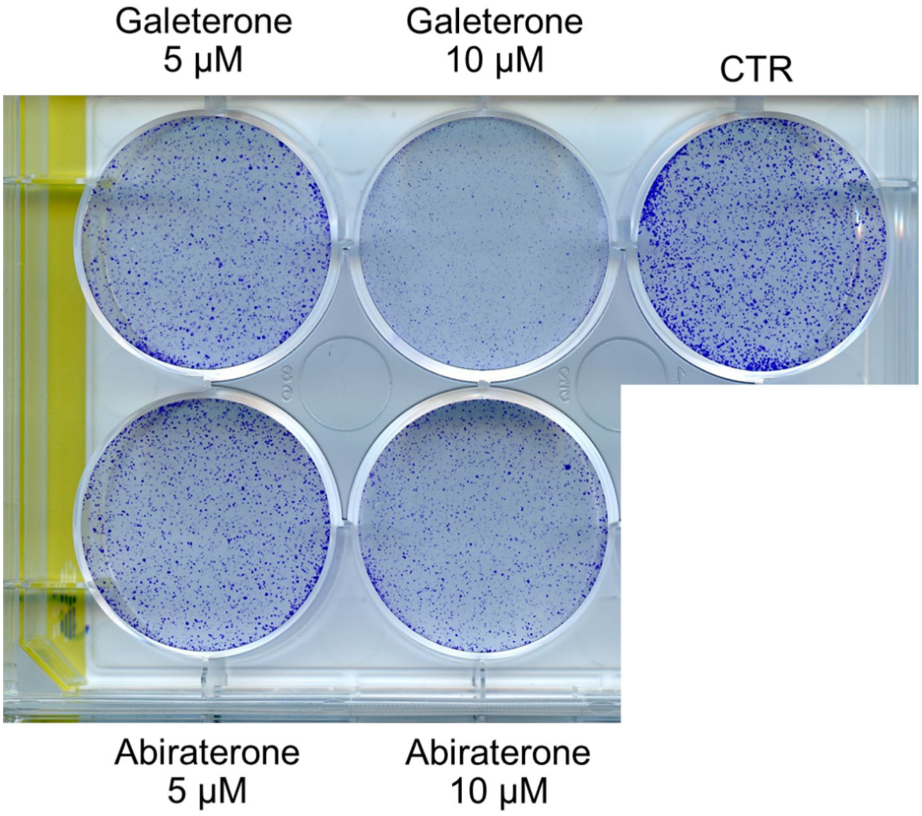


**Figure S2**. Colony formation assay of PCa cells LAPC-4 after treatment with standards for 10 days.

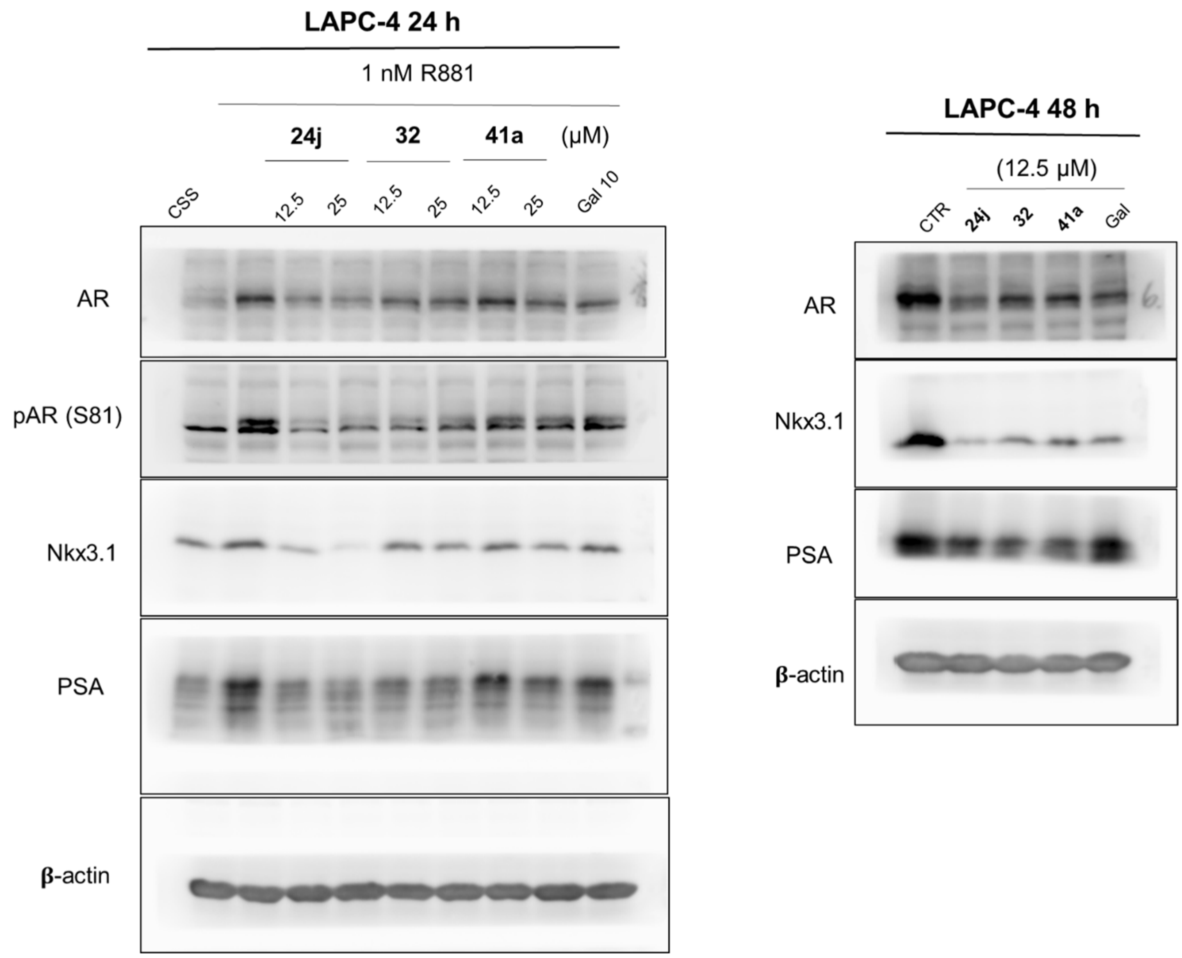


**Figure S3**. Raw picture for **Figure 3**. Western blotting analysis of AR and AR-regulated proteins in treated LAPC-4 cells.

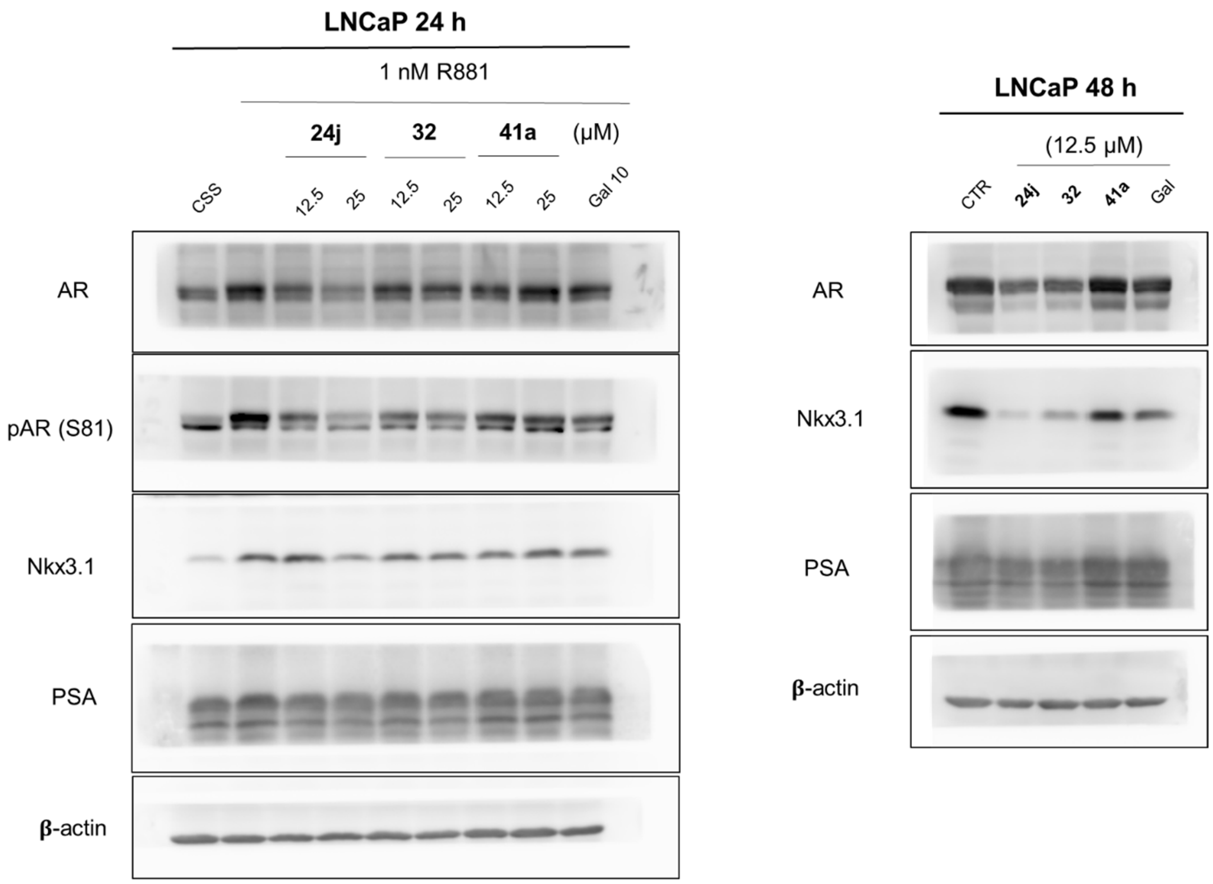


**Figure S4**. Raw picture for **Figure 3**. Western blotting analysis of AR and AR-regulated proteins in treated LAPC-4 cells.

**1H and 13C NMR spectra**

Methyl 2-(3β-((*tert*-butyldimethylsilyl)oxy)-androst-5-en-17-yl)acetate (**12**)

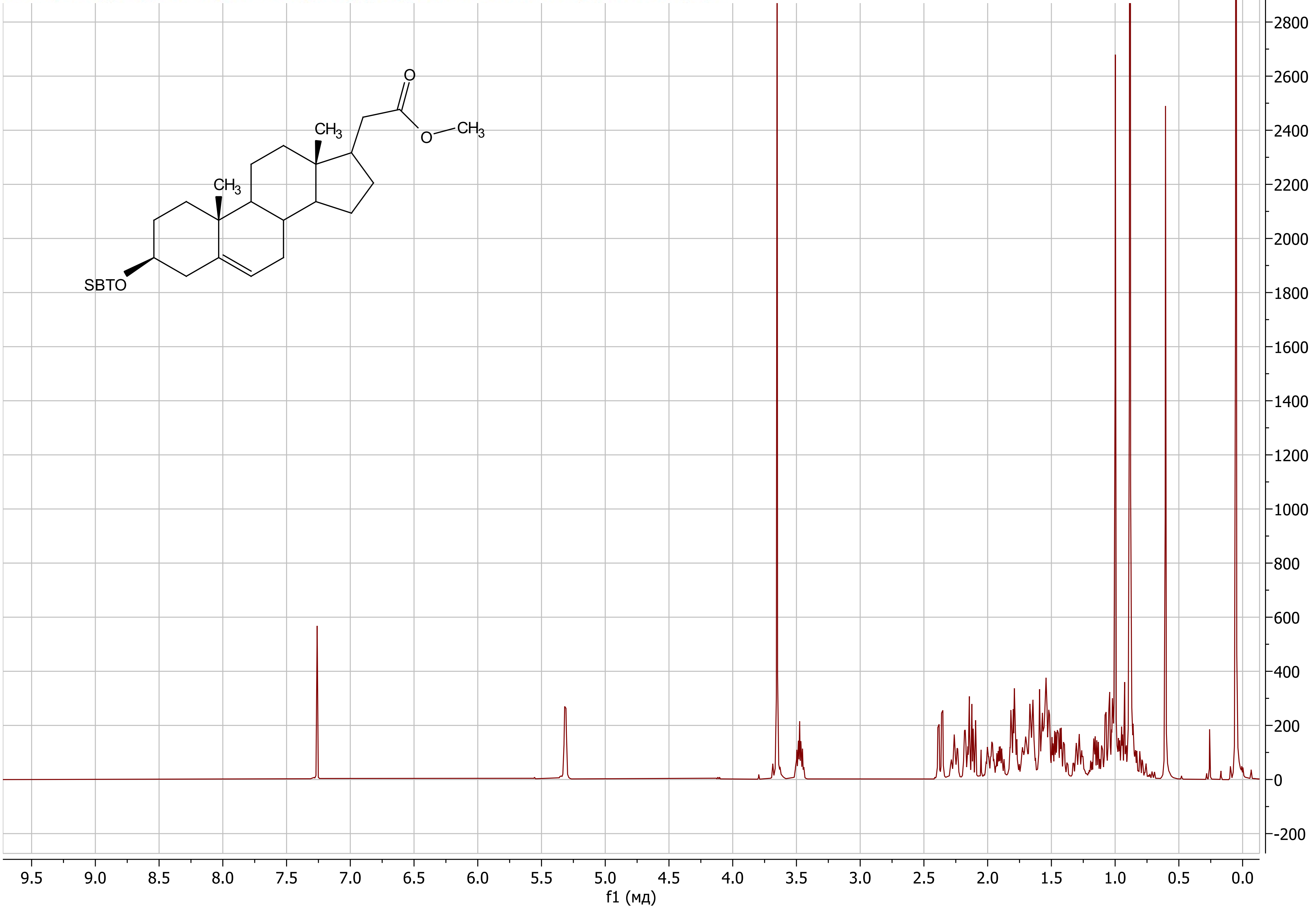

Methyl 2-(3β-((*tert*-butyldimethylsilyl)oxy)-androst-5-en-17-yl)acetate (**12**)

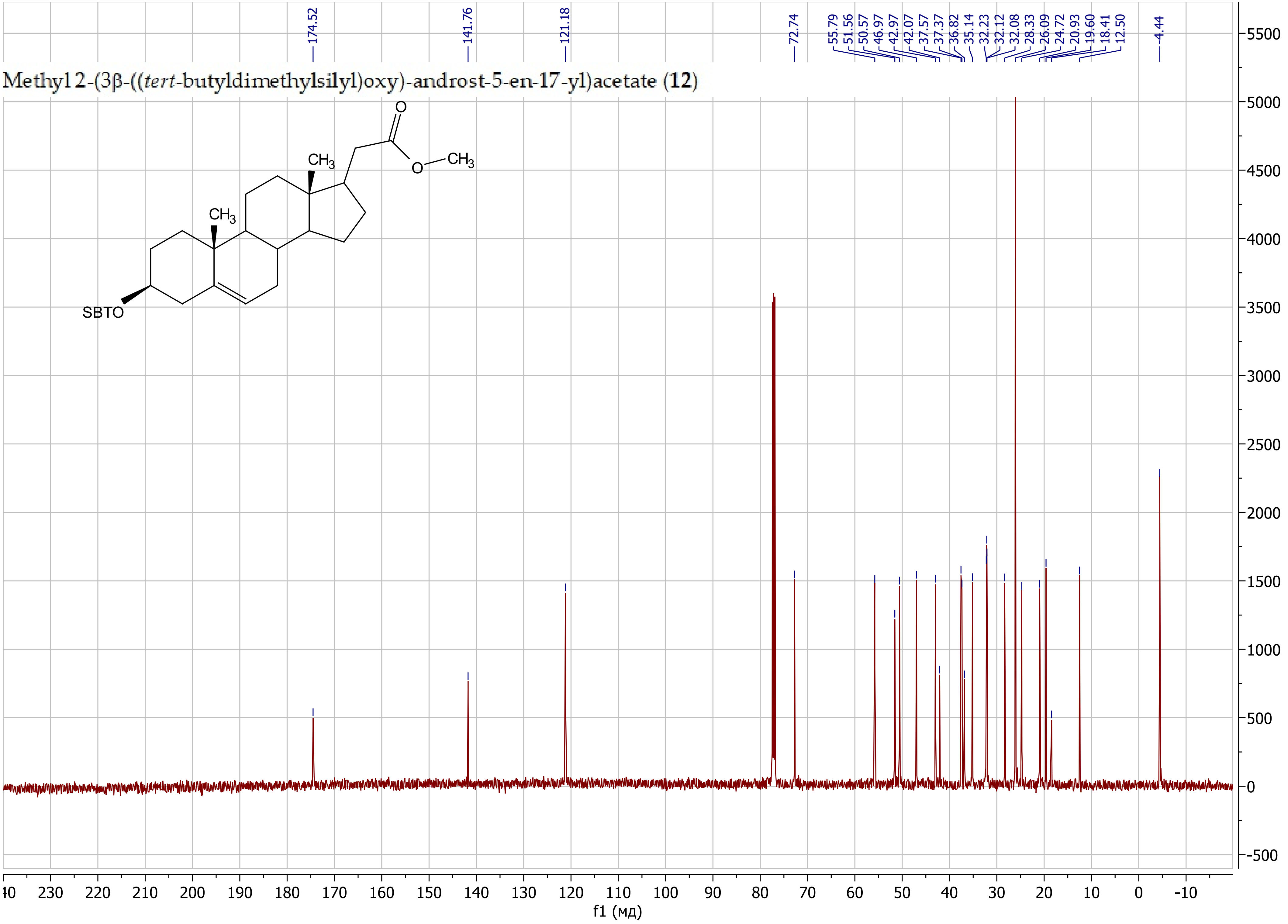

2-(3β-((*tert*-Butyldimethylsilyl)oxy)-androst-5-en-17-yl)-*N*-methoxy-*N*-methylacetamide (**14**)

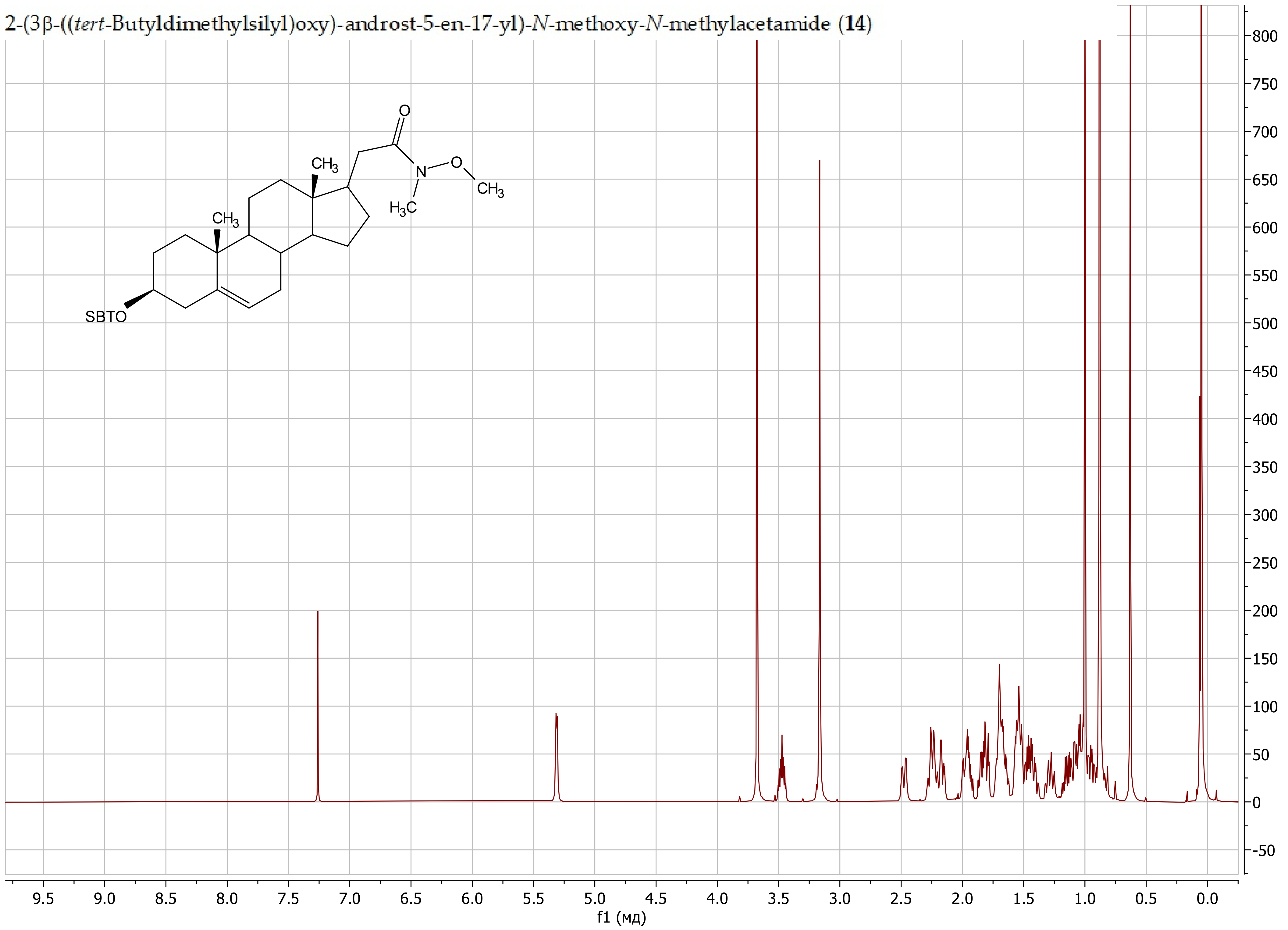

2-(3β-((*tert*-Butyldimethylsilyl)oxy)-androst-5-en-17-yl)-*N*-methoxy-*N*-methylacetamide (**14**)

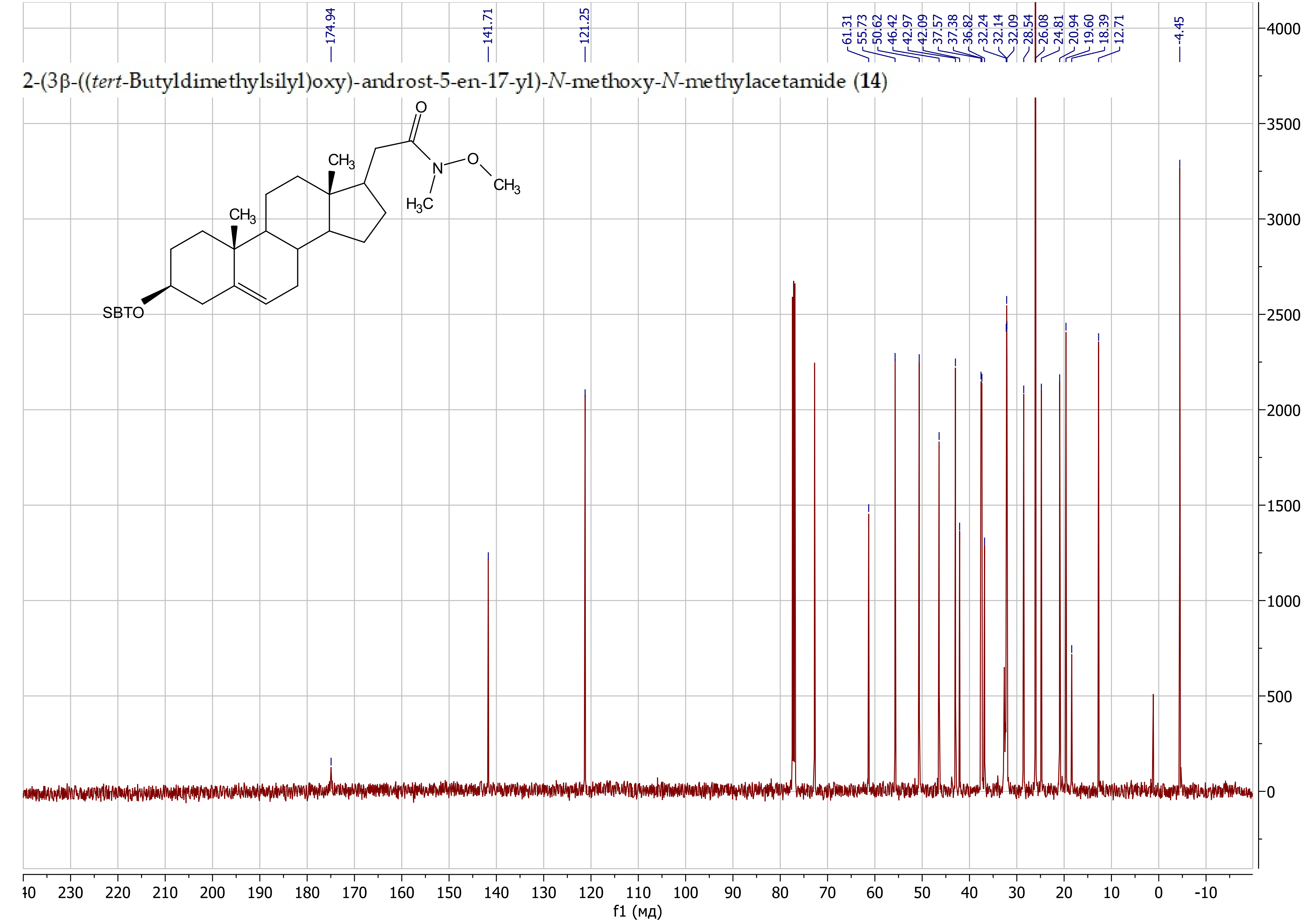

3β-((*tert*-Butyldimethylsilyl)oxy)-pregn-5-en-21-ol (**15**)

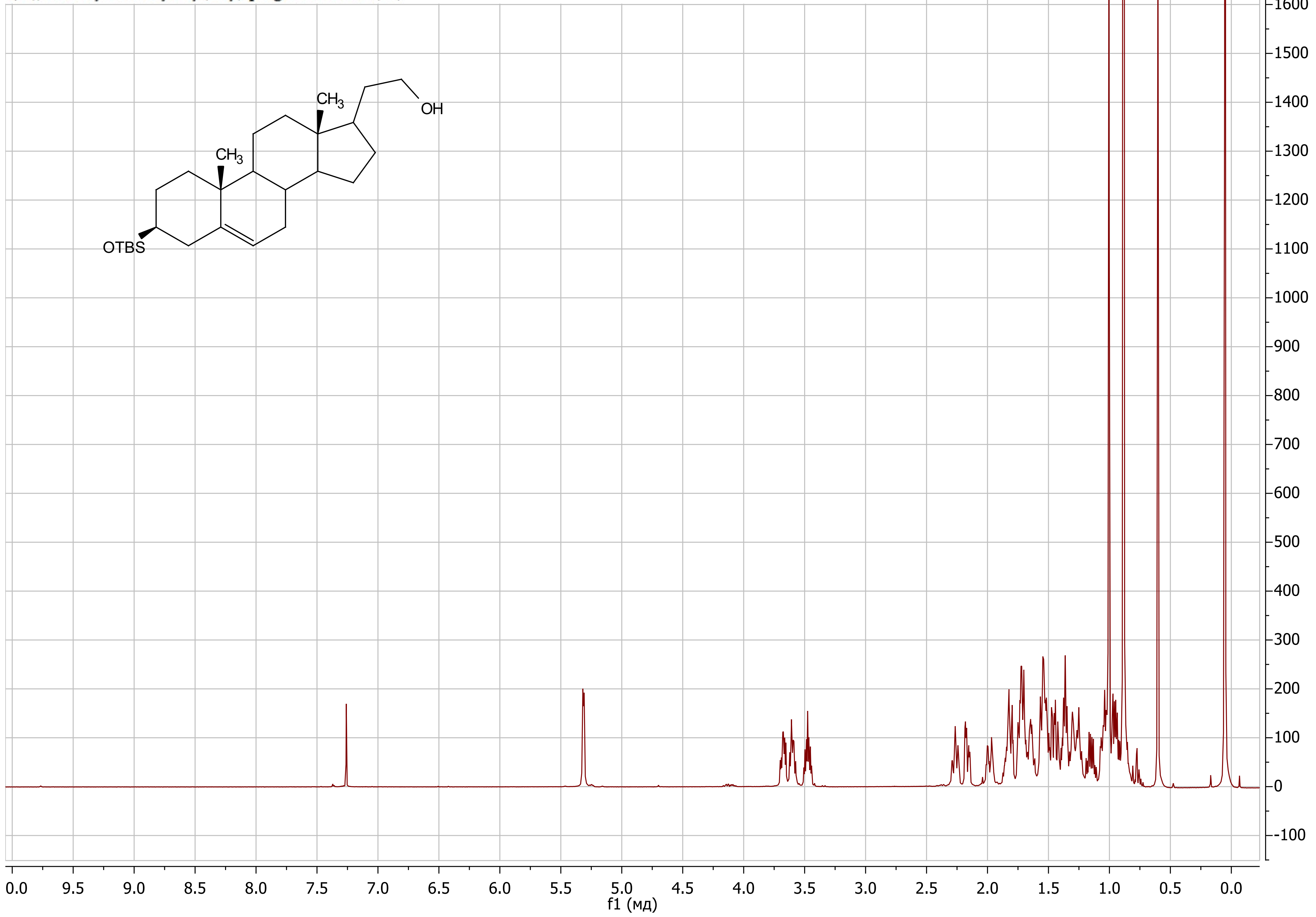

3β-((*tert*-Butyldimethylsilyl)oxy)-pregn-5-en-21-ol (**15**)

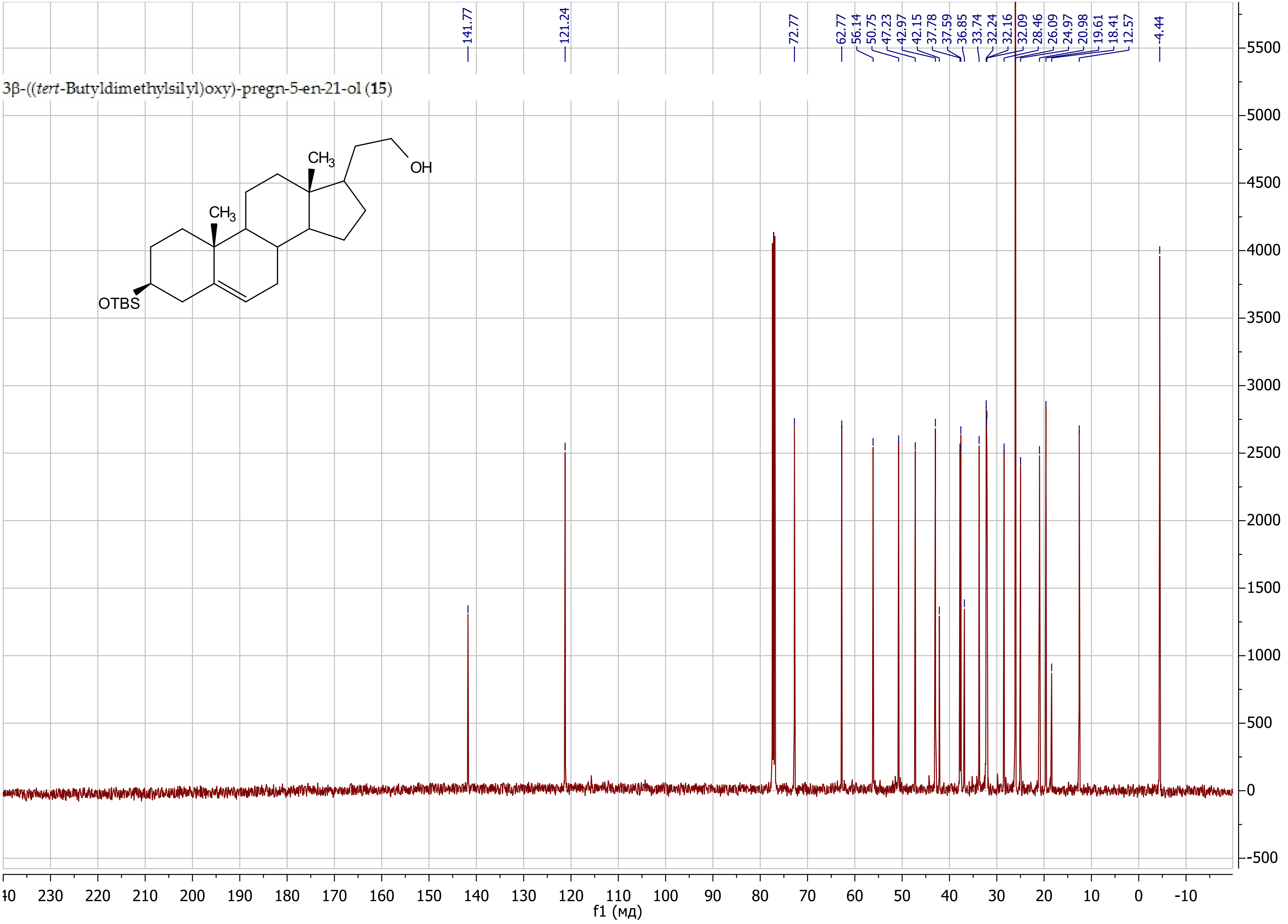

5-((*tert*-Butyldimethylsilyl)oxy)-1-((17*R*)-3β-((*tert*-butyldimethylsilyl)oxy)-androst-5-en-17-yl)pent-3-yn-2-one (**18**)

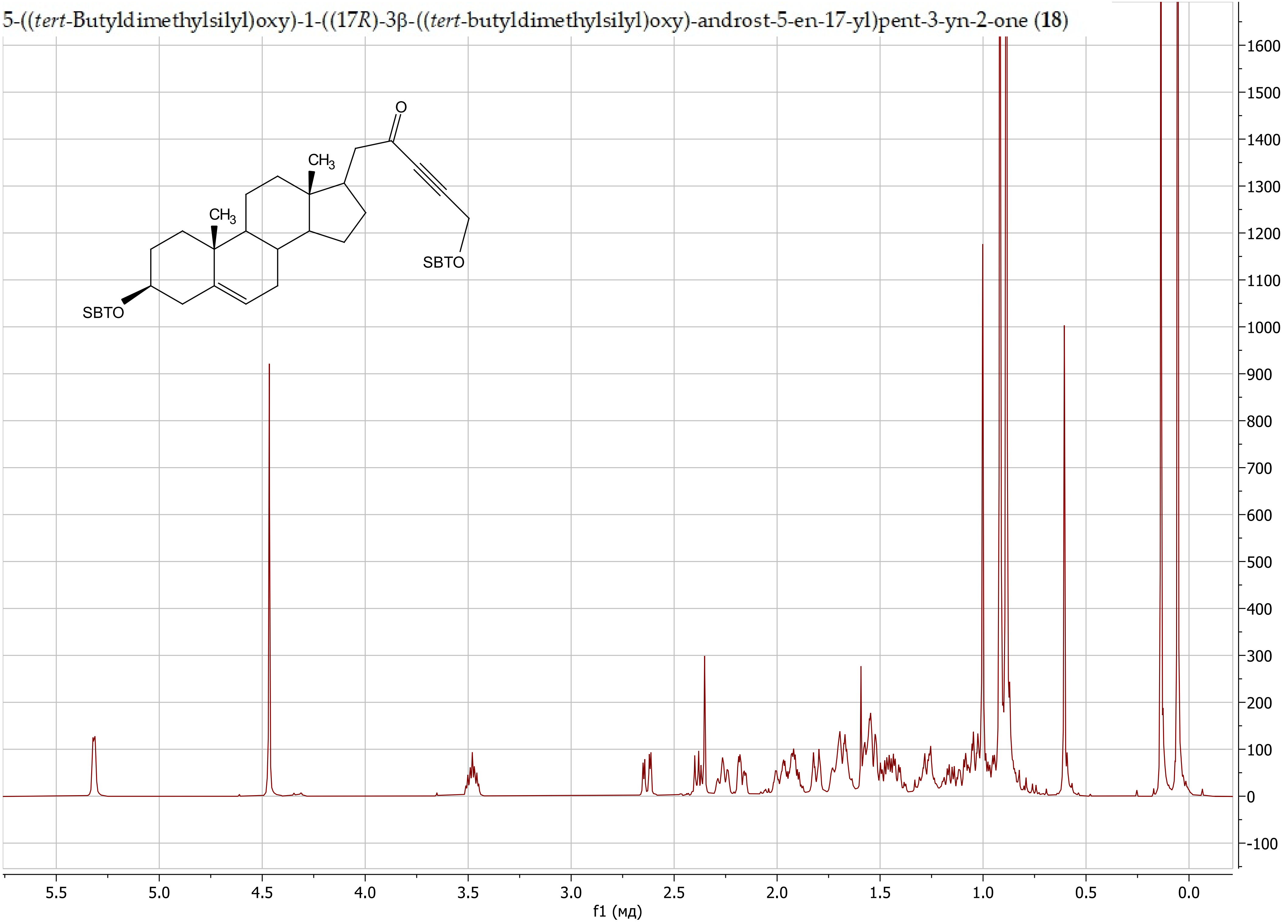

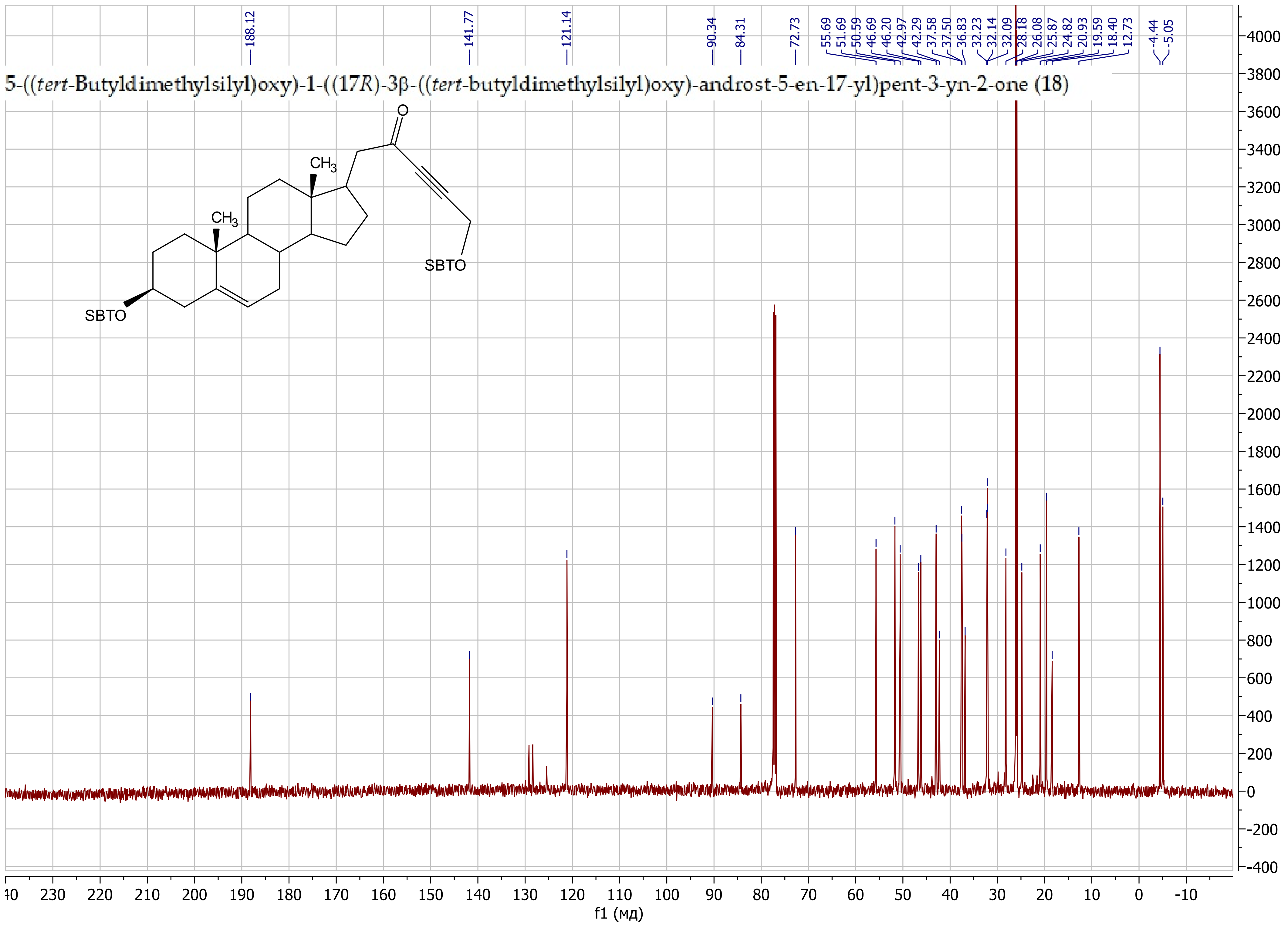
5-((tert-Butyldimethylsilyl)oxy)-1-((17R)-3β-((tert-butyldimethylsilyl)oxy)-androst-5-en-17-yl)pent-3-yn-2-one (18)
188.12
141.77
121.14
90.34
84.31
72.73
55.69
51.69
50.59
46.69
46.20
42.97
42.29
37.58
37.50
36.83
32.23
32.14
32.09
28.18
26.08
25.87
24.82
20.93
19.59
18.40
12.73
-4.44
-5.05
O
CH3
CH3
SBTO
SBTO
f1 (мд)

1-(((17*R*)-3β-((*tert*-Butyldimethylsilyl)oxy)-androst-5-en-17-yl)but-3-yn-2-one (**20a**)

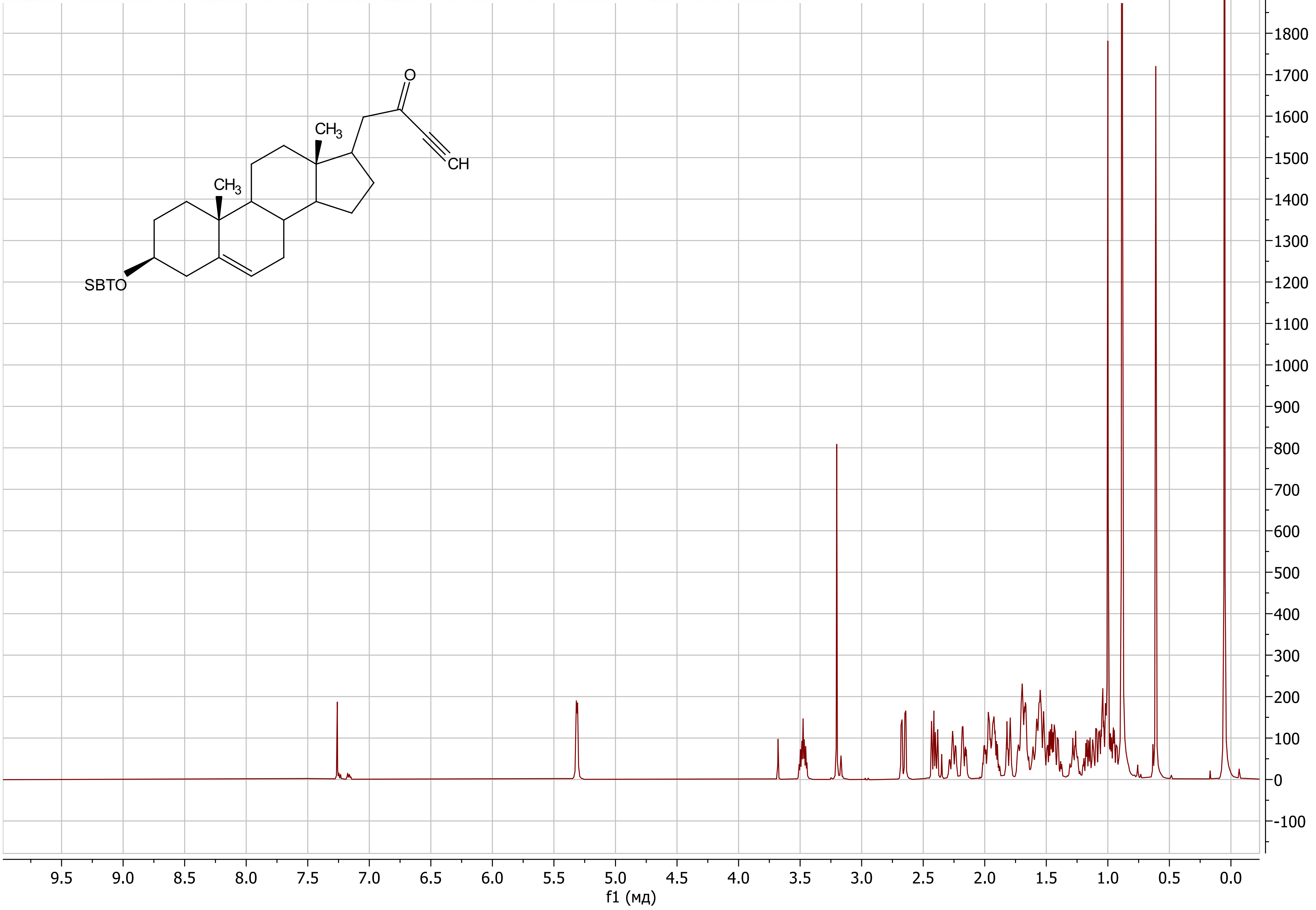

1-(((17*R*)-3β-((*tert*-Butyldimethylsilyl)oxy)-androst-5-en-17-yl)but-3-yn-2-one (**20a**)

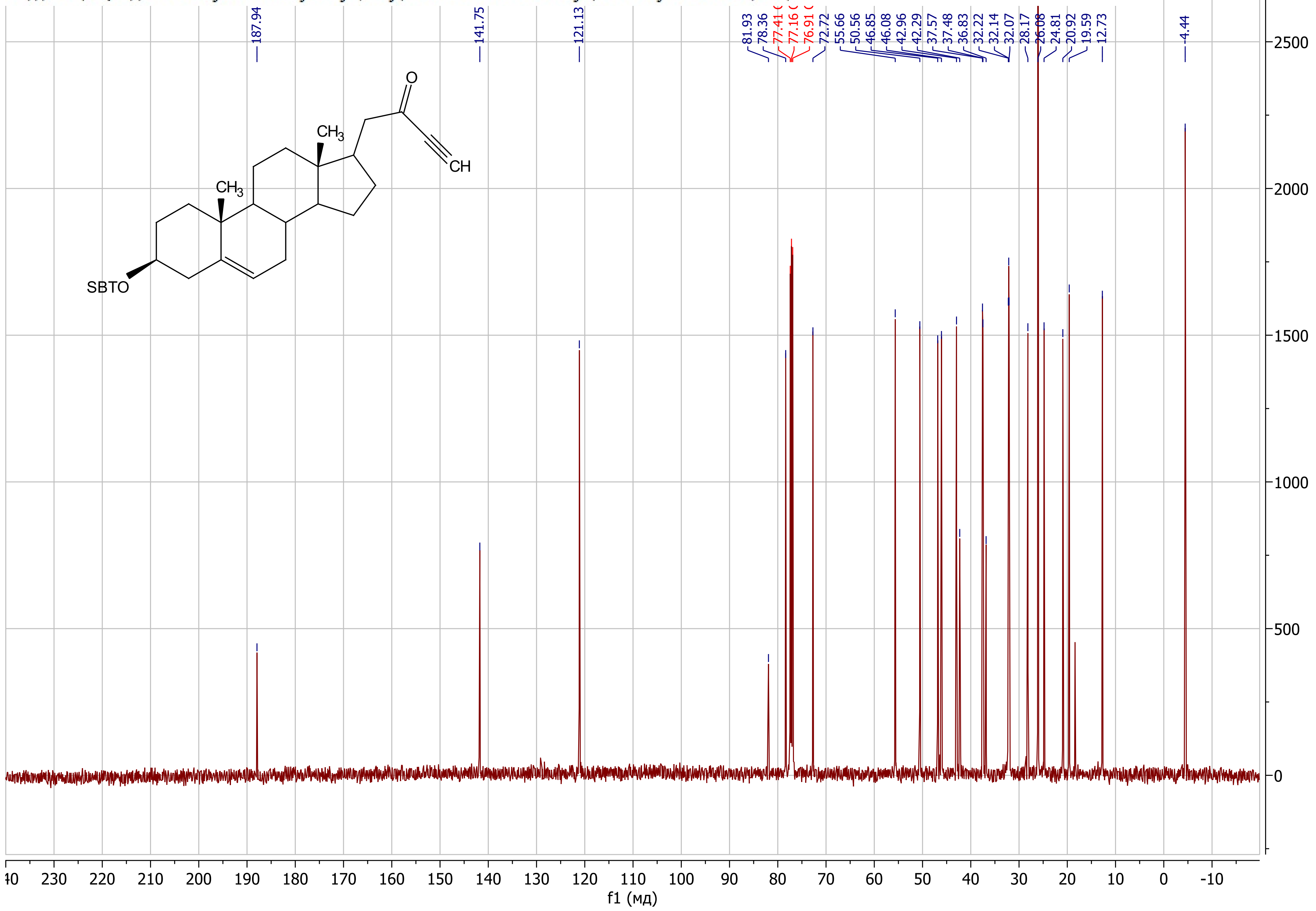

1-((17*R*)-3β-((*tert*-Butyldimethylsilyl)oxy)-androst-5-en-17-yl)-5-methylhex-3-yn-2-one (**20b**)

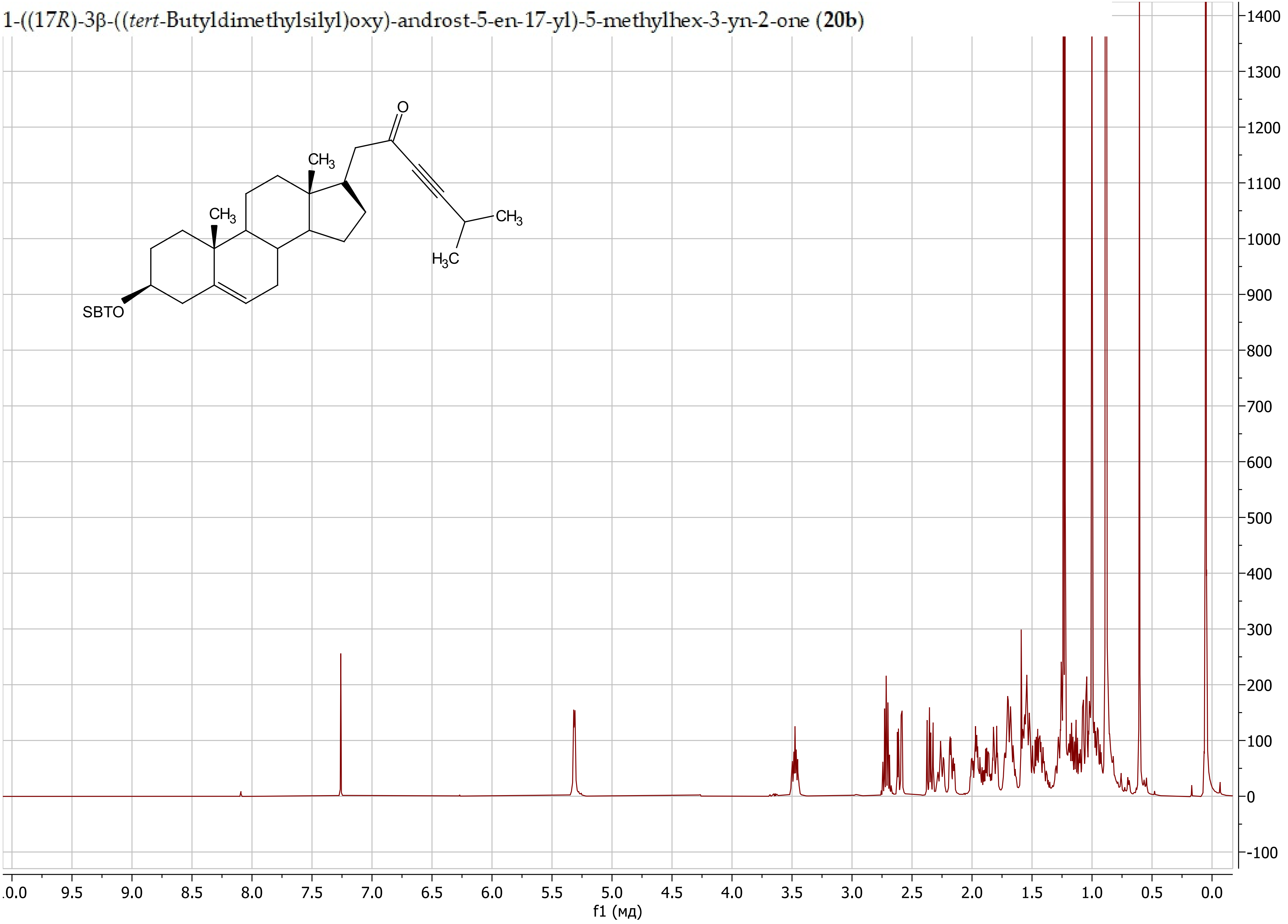

1-((17*R*)-3β-((*tert*-Butyldimethylsilyl)oxy)-androst-5-en-17-yl)-5-methylhex-3-yn-2-one (**20b**)

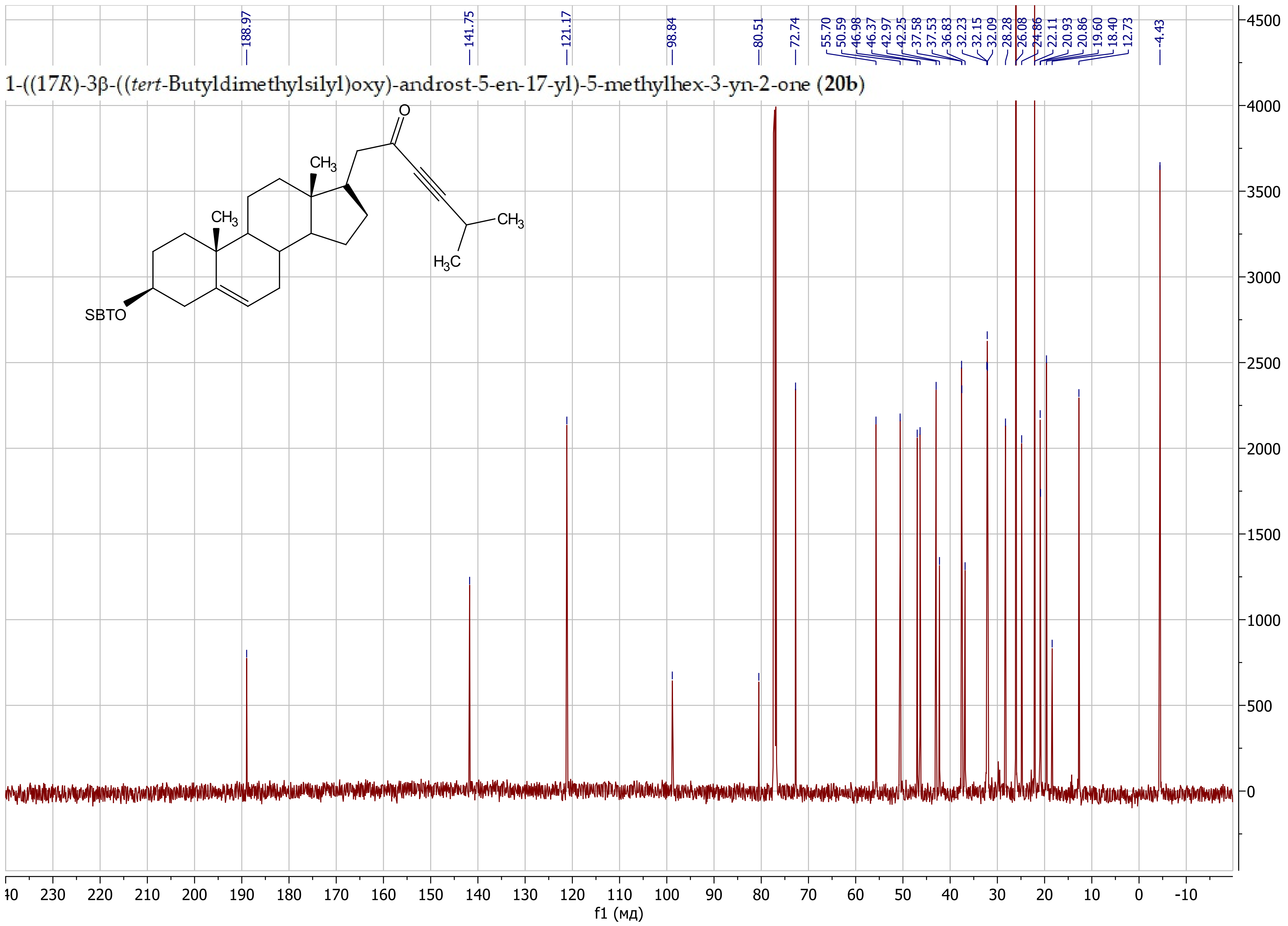

1-((17*R*)-3β-((*tert*-Butyldimethylsilyl)oxy)-androst-5-en-17-yl)-4-cyclopropylbut-3-yn-2-one (**20c**)

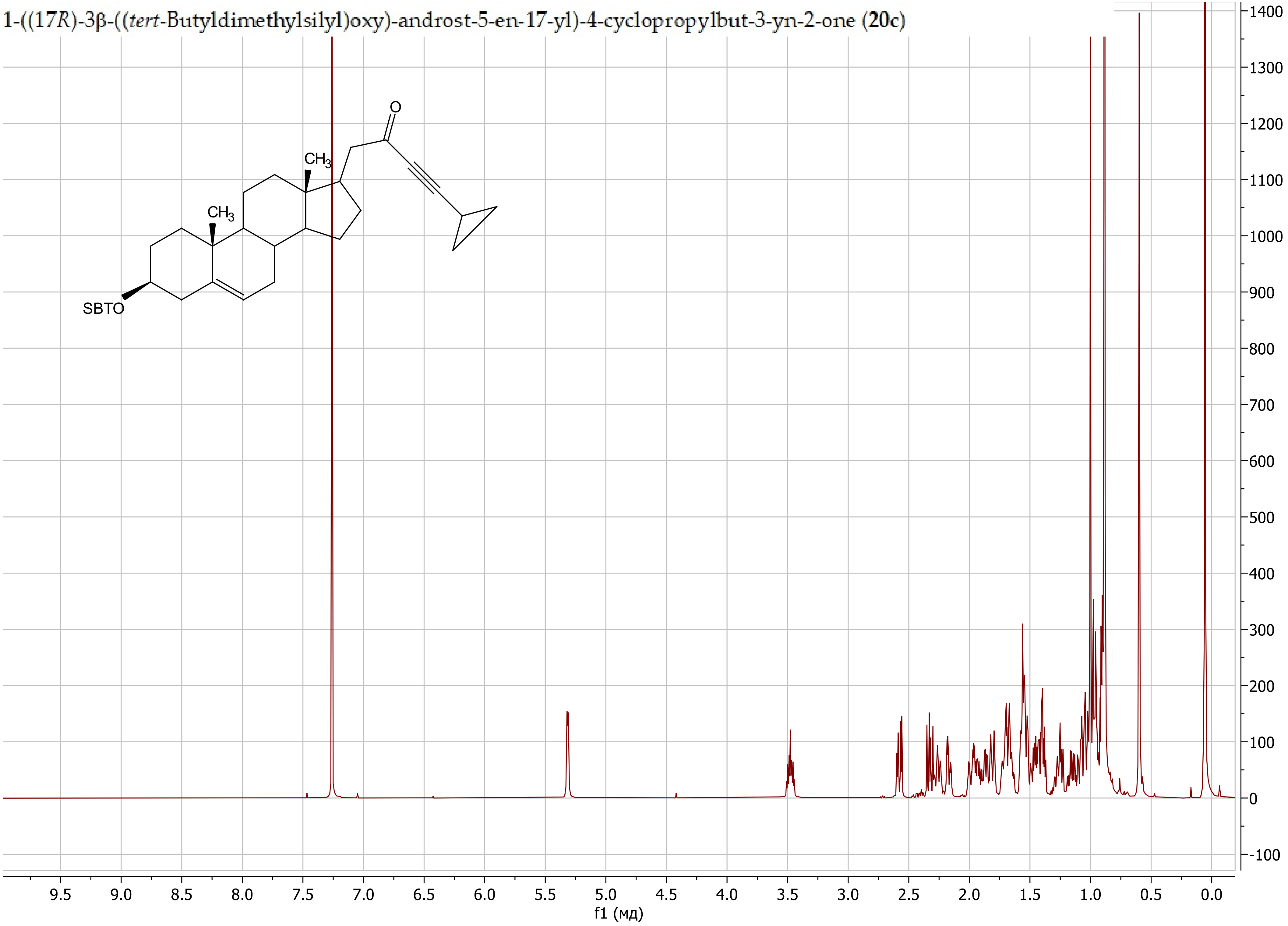

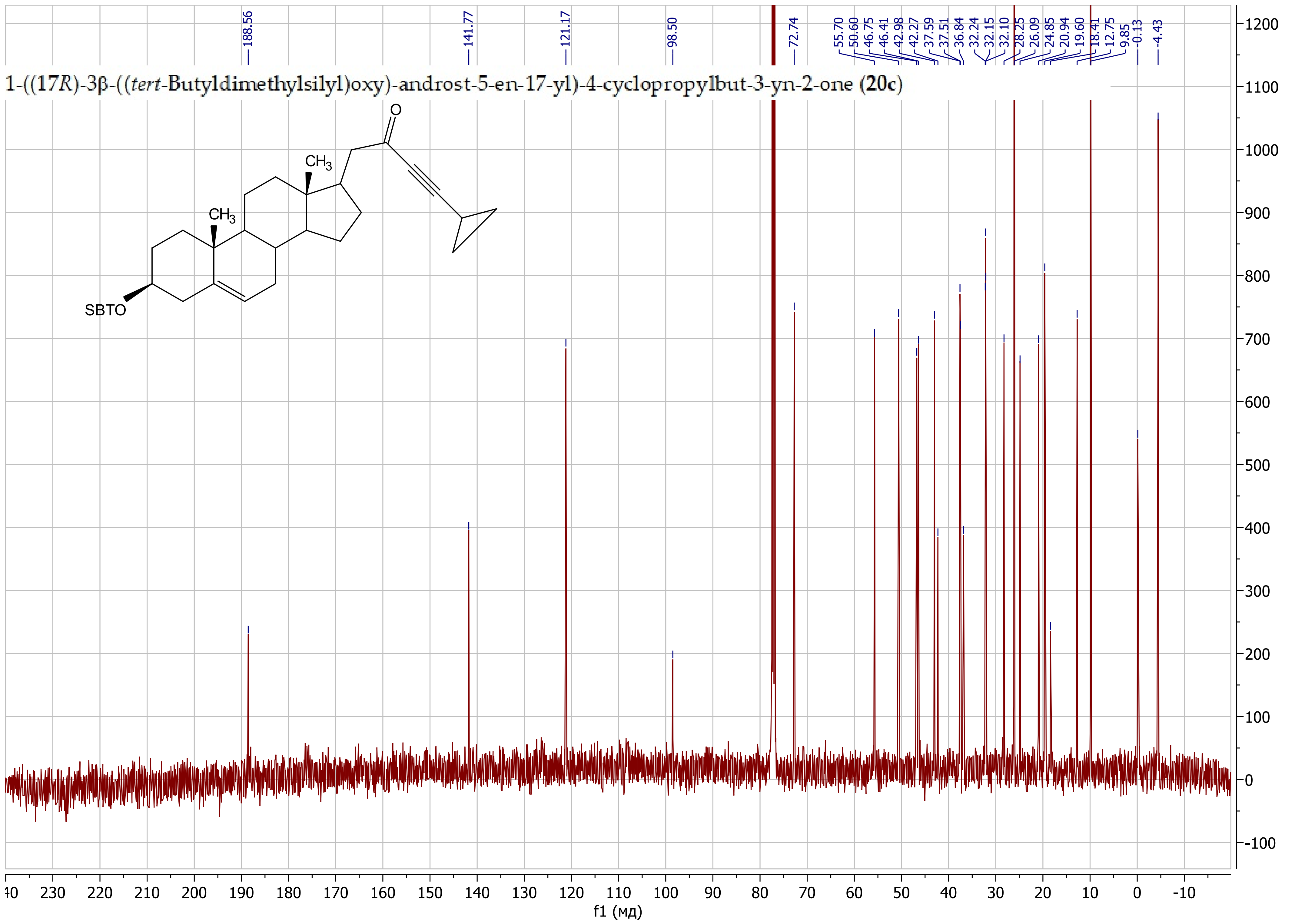
1-((17R)-3β-((tert-Butyldimethylsilyl)oxy)-androst-5-en-17-yl)-4-cyclopropylbut-3-yn-2-one (20c)
188.56
141.77
121.17
98.50
72.74
55.70
50.60
46.75
46.41
42.98
42.27
37.59
37.51
36.84
32.24
32.15
32.10
28.25
26.09
24.85
20.94
19.60
18.41
12.75
9.85
-0.13
-4.43
O
CH3
CH3
SBTO
f1 (мд)

1-((17*R*)-3β-((*tert*-Butyldimethylsilyl)oxy)-androst-5-en-17-yl)oct-3-yn-2-one (**20d**)

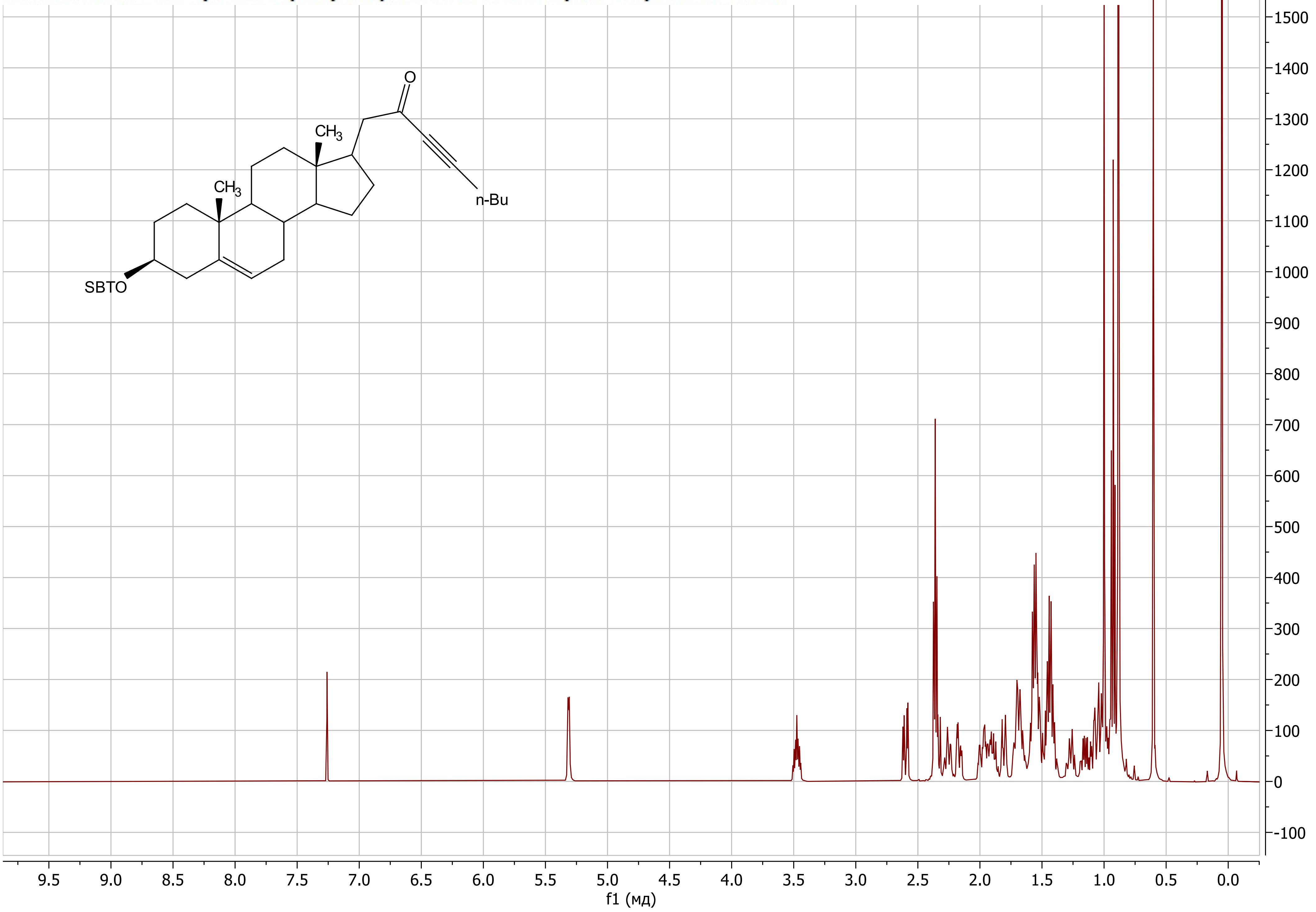

1-((17*R*)-3β-((*tert*-Butyldimethylsilyl)oxy)-androst-5-en-17-yl)oct-3-yn-2-one (**20d**)

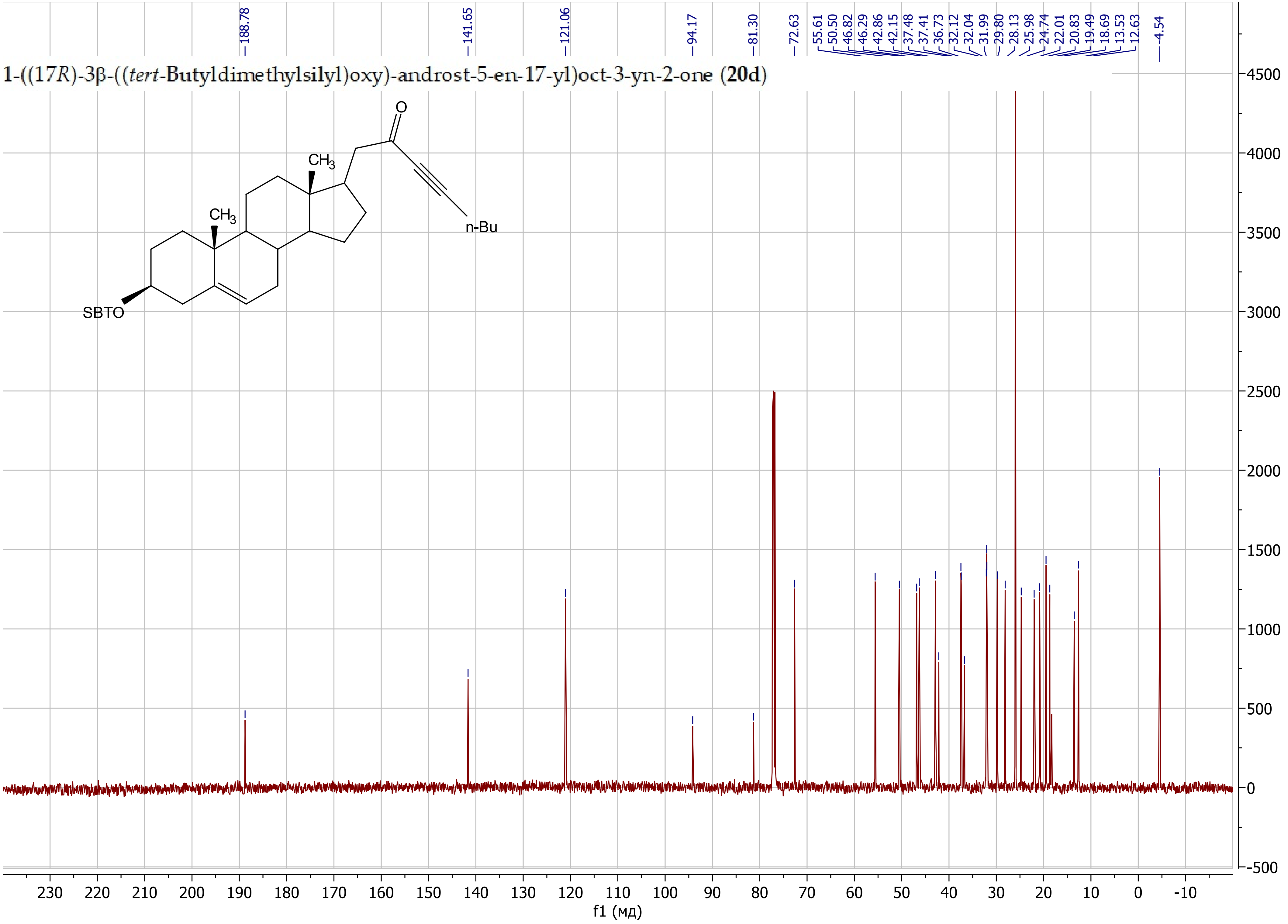

1-((17*R*)-3β-((*tert*-Butyldimethylsilyl)oxy)-androst-5-en-17-yl)-4-phenylbut-3-yn-2-one (**20e**)

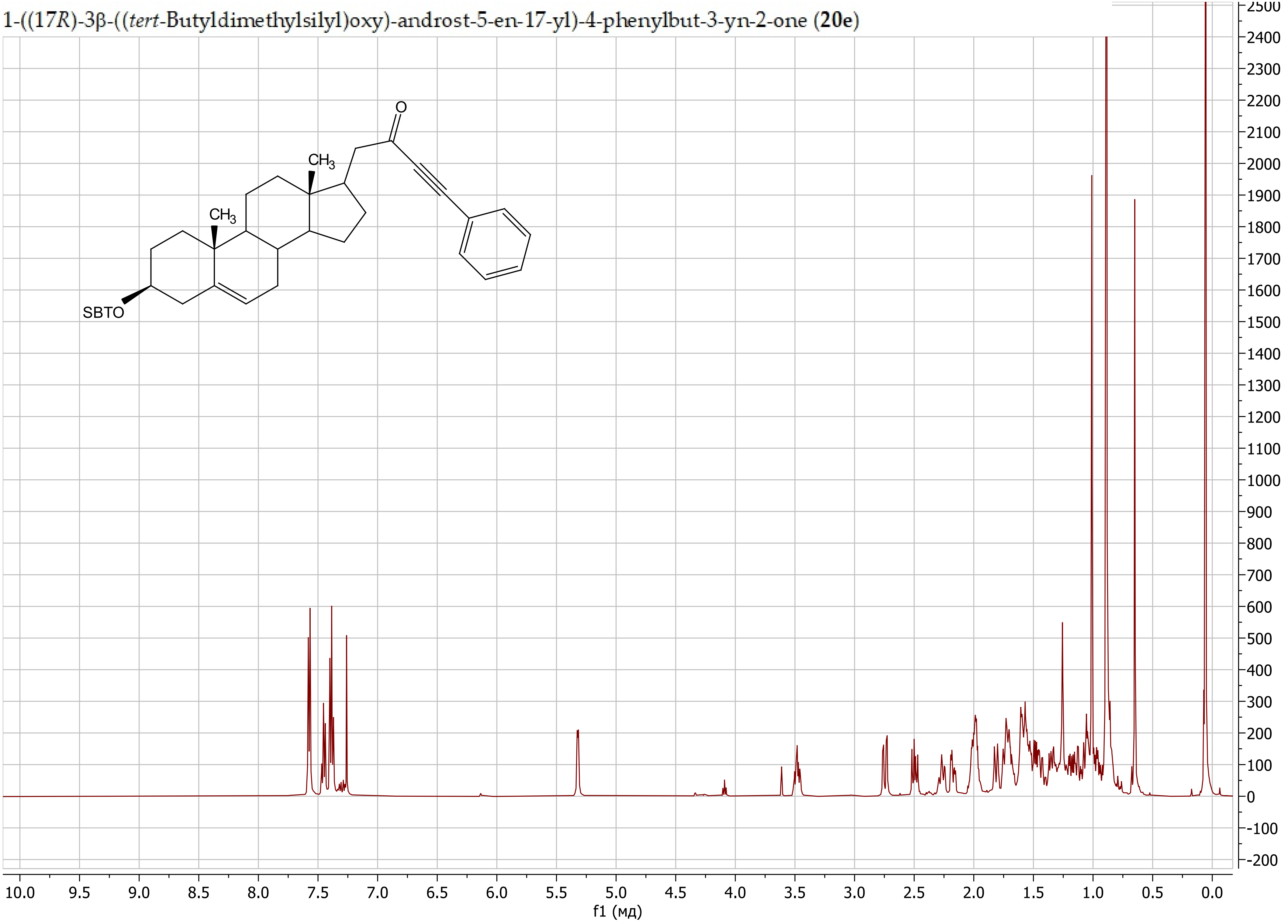

1-((17*R*)-3β-((*tert*-Butyldimethylsilyl)oxy)-androst-5-en-17-yl)-4-phenylbut-3-yn-2-one (**20e**)

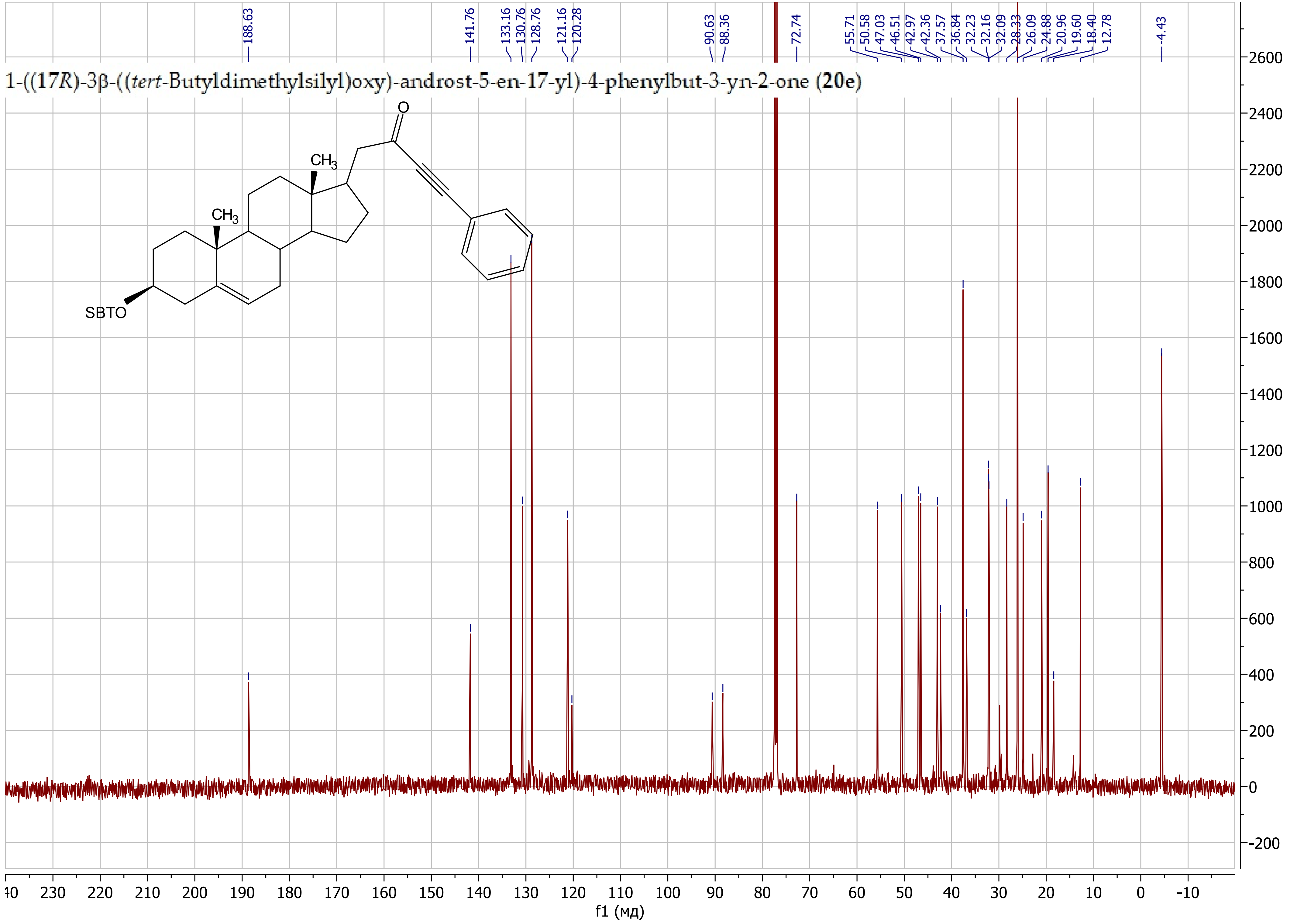

1-((17*R*)-3β-((*tert*-Butyldimethylsilyl)oxy)-androst-5-en-17-yl)-4-(pyridin-3-yl)but-3-yn-2-one (**20f**)

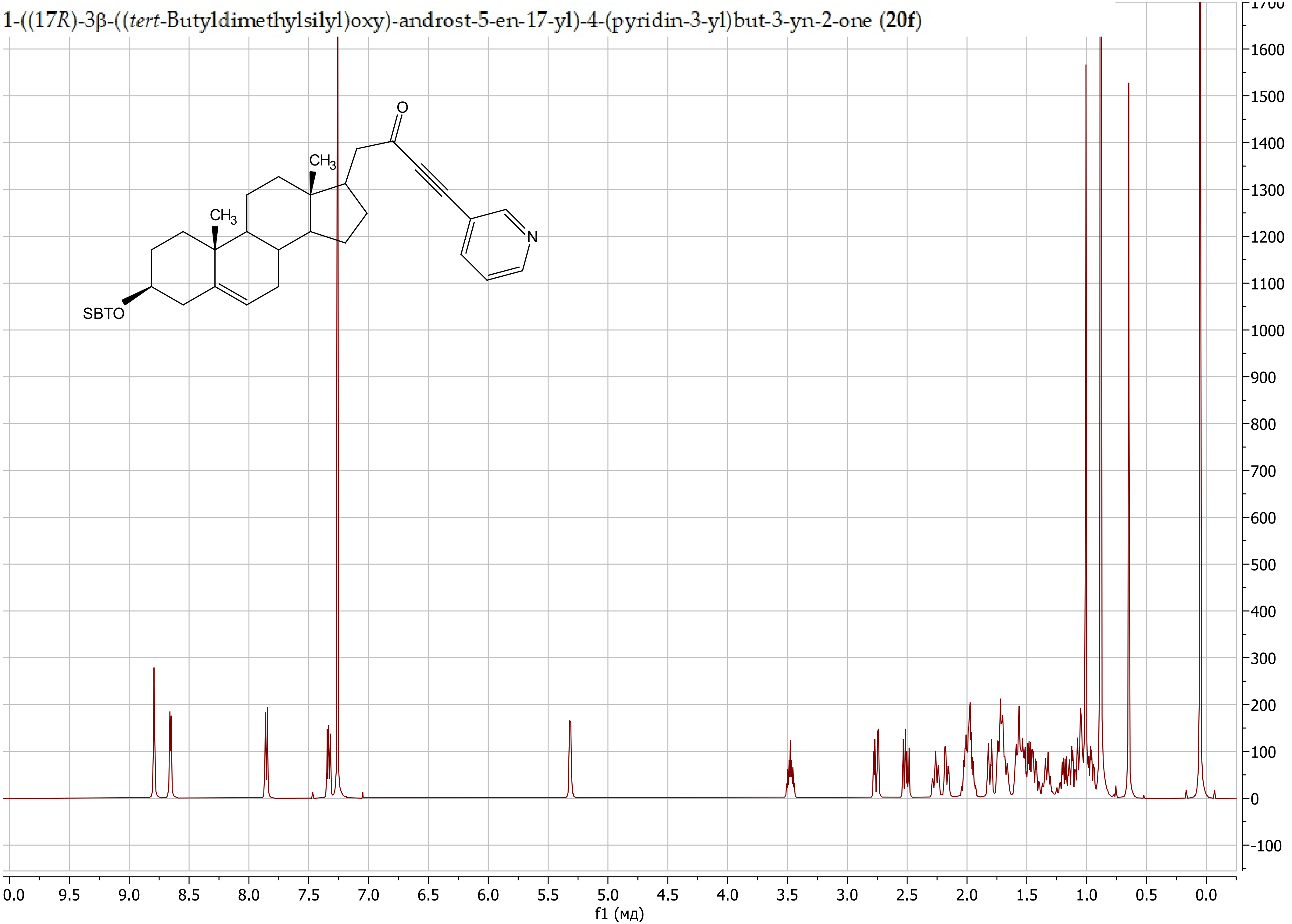

1-((17*R*)-3β-((*tert*-Butyldimethylsilyl)oxy)-androst-5-en-17-yl)-4-(pyridin-3-yl)but-3-yn-2-one (**20f**)

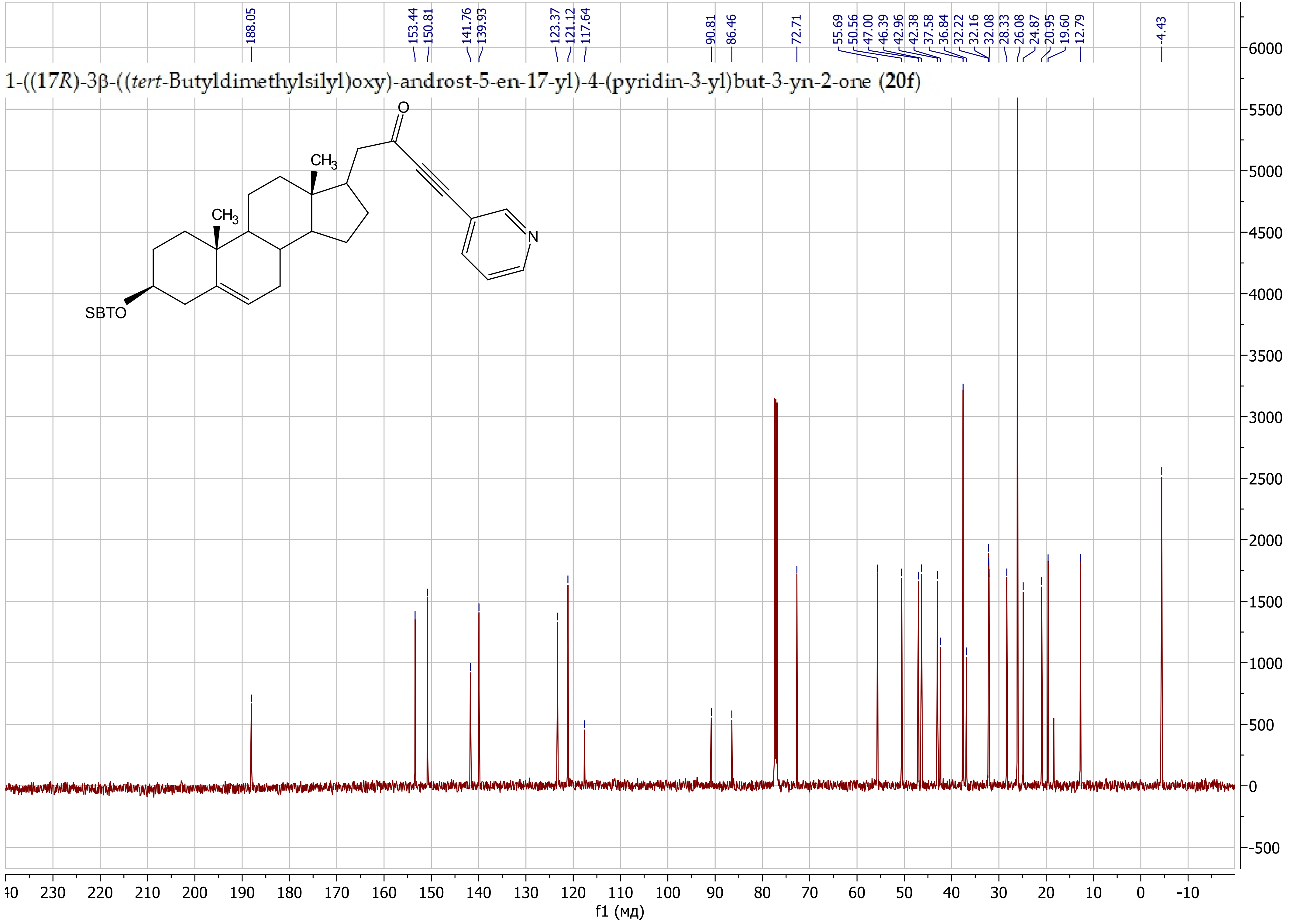

1-((17*R*)-3β-((*tert*-Butyldimethylsilyl)oxy)-androst-5-en-17-yl)-4-(2-fluorophenyl)but-3-yn-2-one (**20g**)

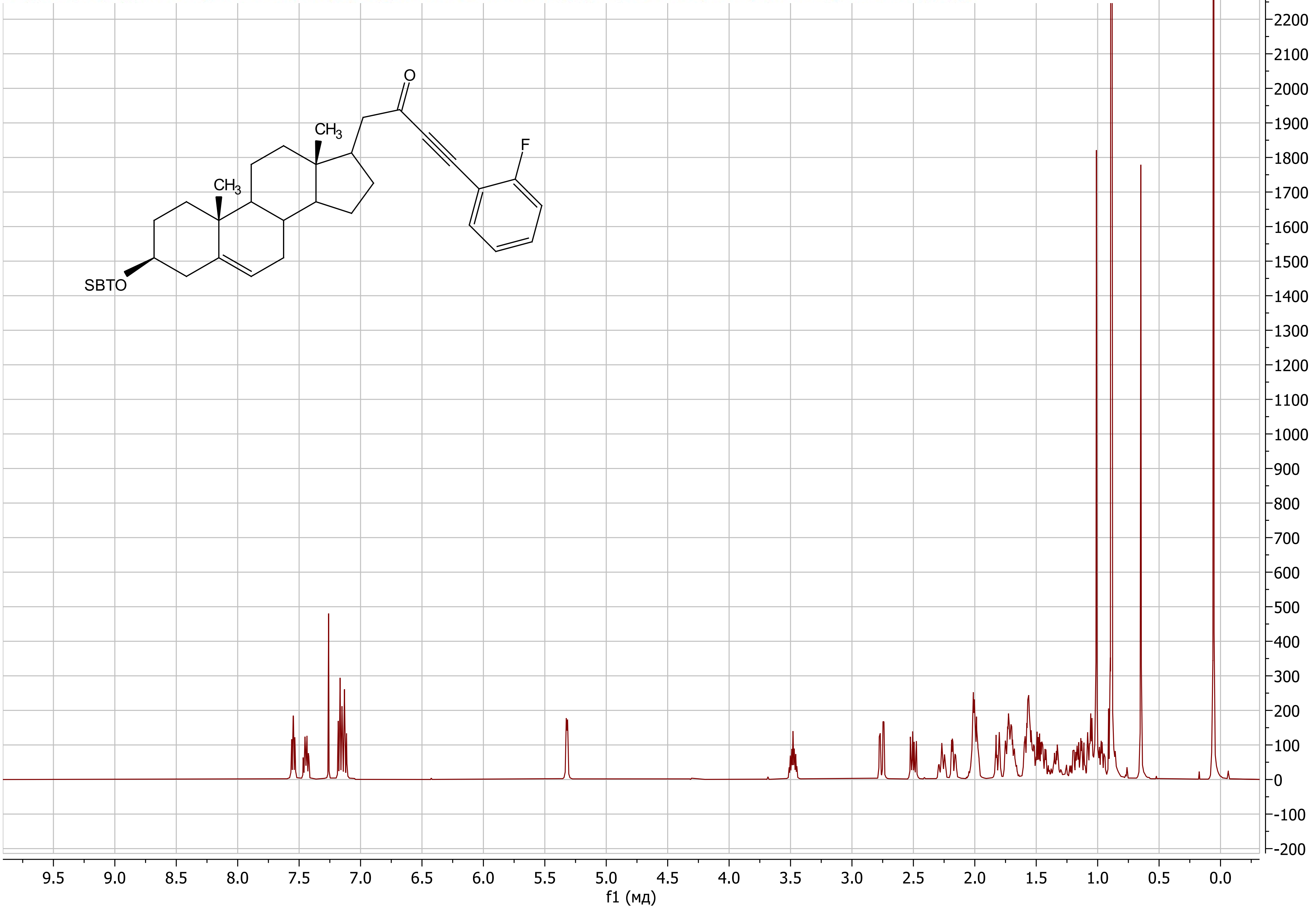

1-((17*R*)-3β-((*tert*-Butyldimethylsilyl)oxy)-androst-5-en-17-yl)-4-(2-fluorophenyl)but-3-yn-2-one (**20g**)

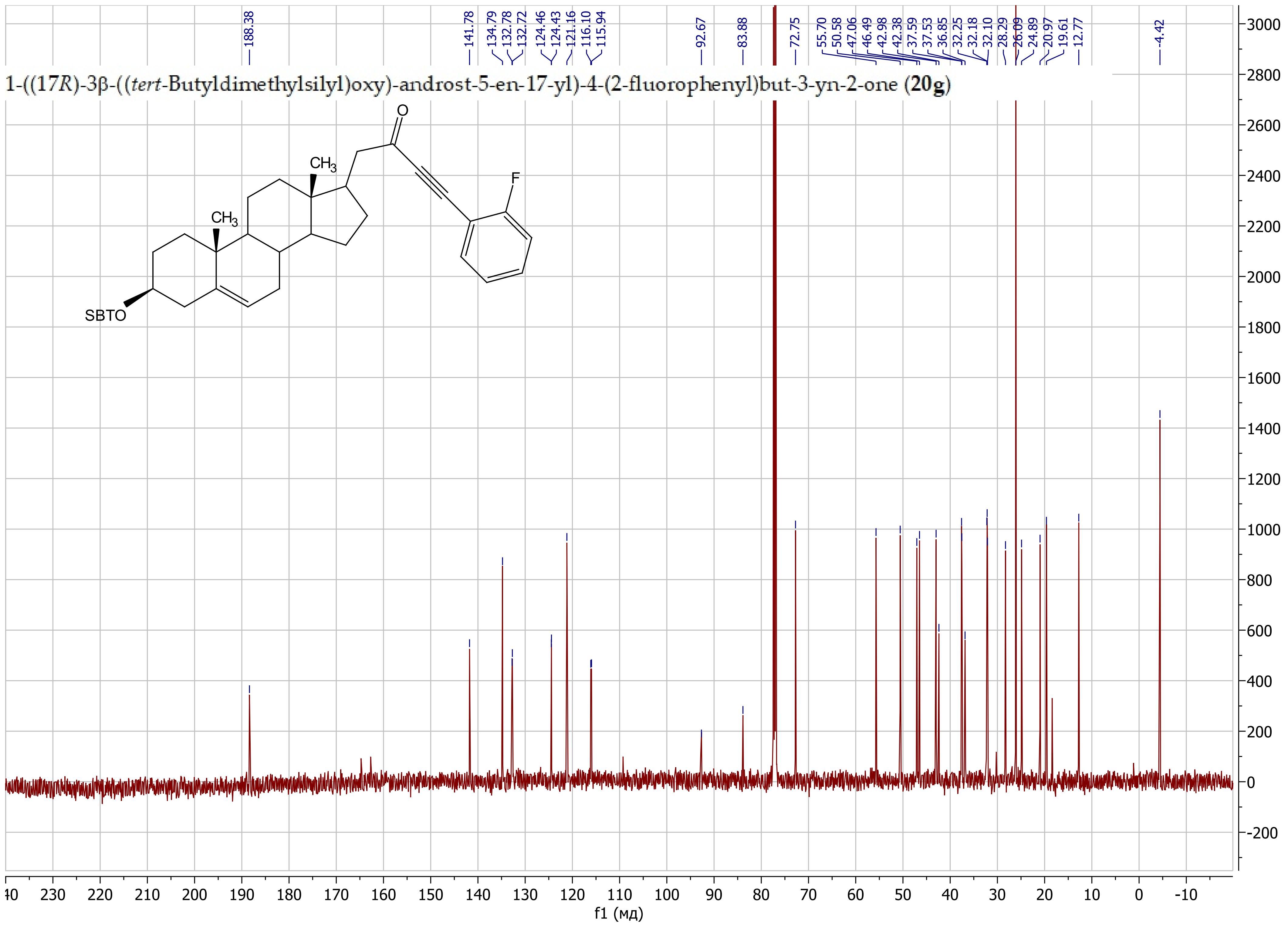

1-((17*R*)-3β-((*tert*-Butyldimethylsilyl)oxy)-androst-5-en-17-yl)-5-methyl-5-((tetrahydro-2*H*-pyran-2-yl)oxy)hex-3-yn-2-one (**20h**)

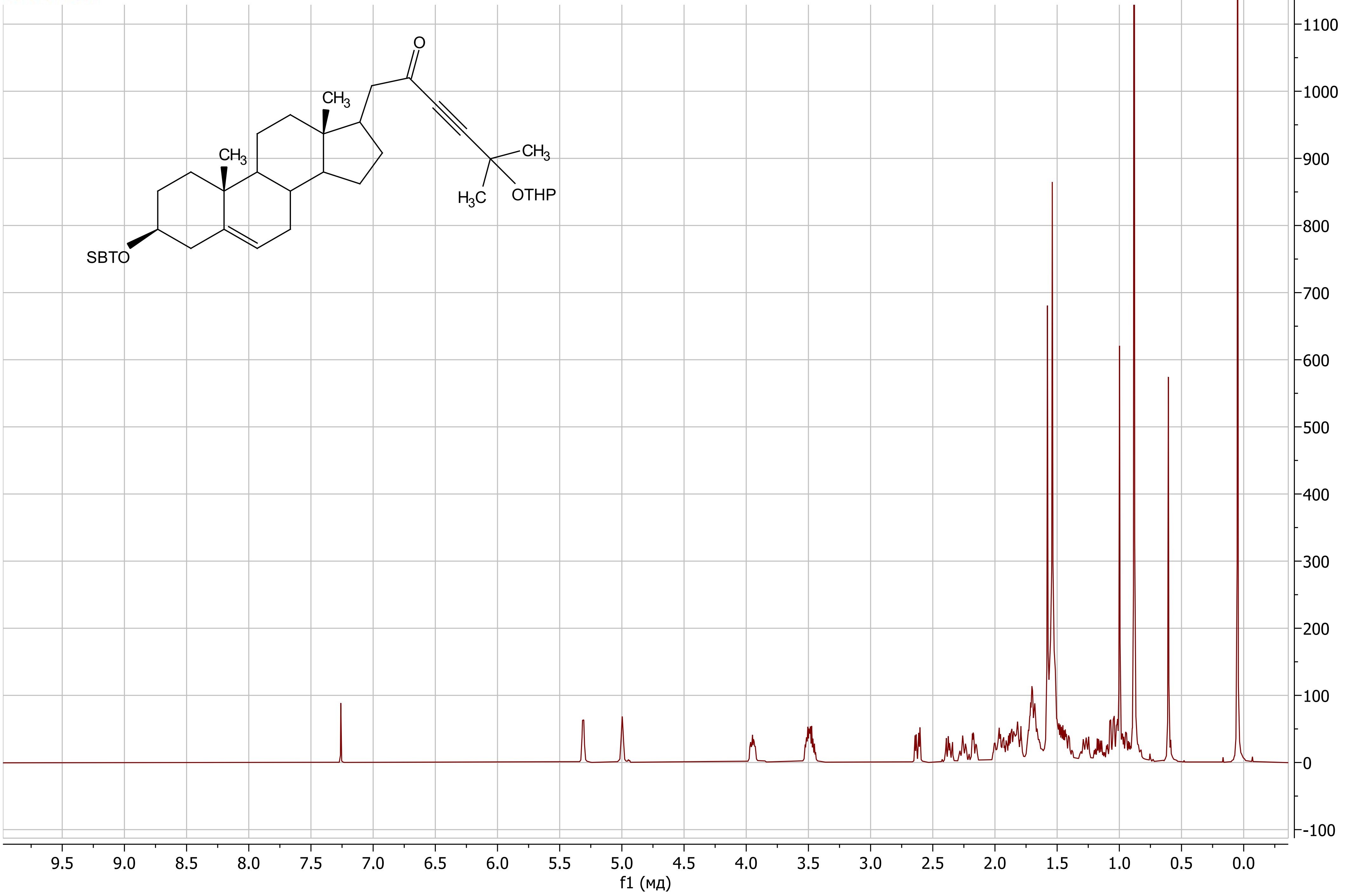

1-((17*R*)-3β-((*tert*-Butyldimethylsilyl)oxy)-androst-5-en-17-yl)-5-methyl-5-((tetrahydro-2*H*-pyran-2-yl)oxy)hex-3-yn-2-one (**20h**)

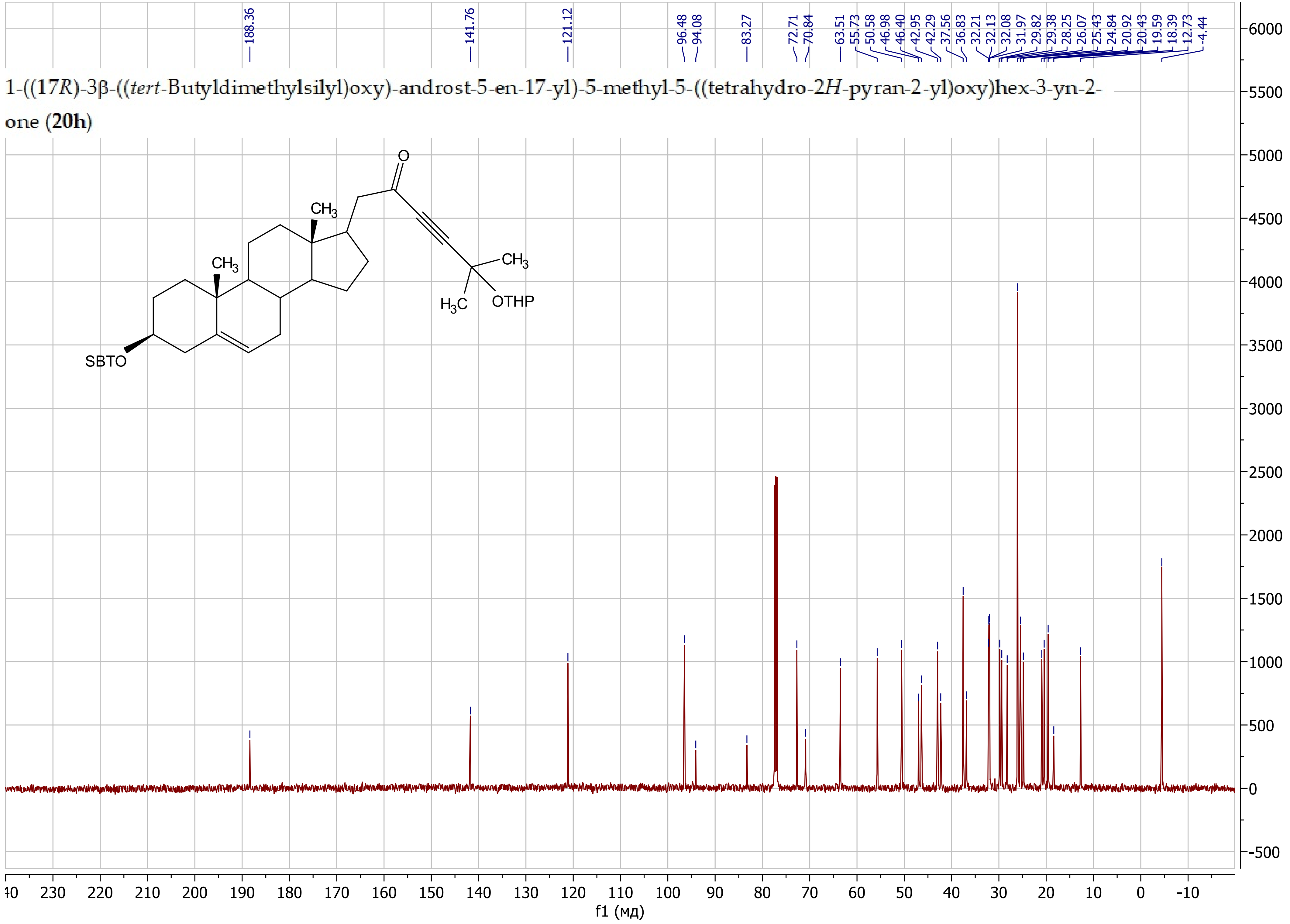

1-((17*R*)-3β-((*tert*-Butyldimethylsilyl)oxy)-androst-5-en-17-yl)-5-((tetrahydro-2*H*-pyran-2-yl)oxy)pent-3-yn-2-one (**20i**)

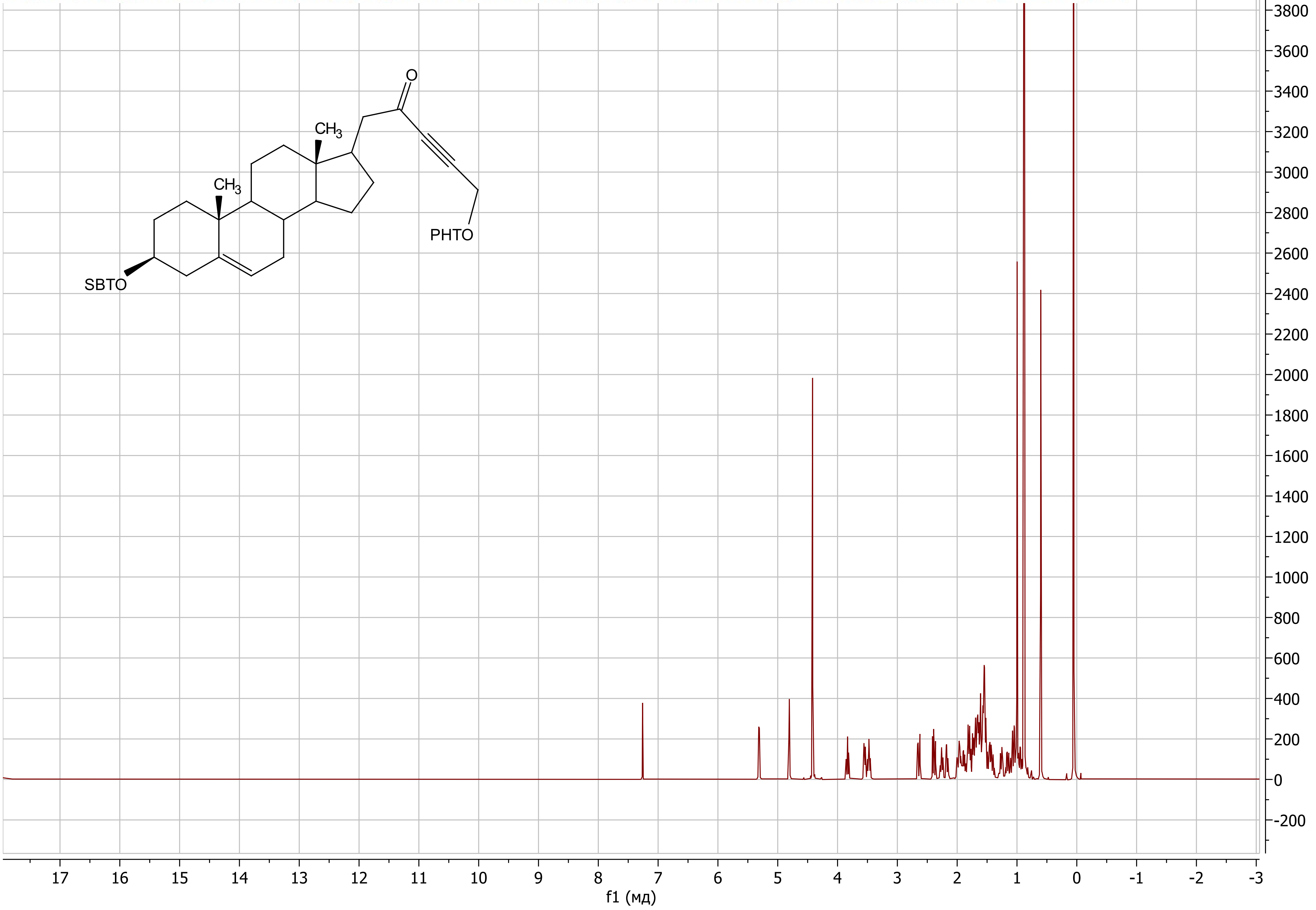

1-((17*R*)-3β-((*tert*-Butyldimethylsilyl)oxy)-androst-5-en-17-yl)-5-((tetrahydro-2*H*-pyran-2-yl)oxy)pent-3-yn-2-one (**20i**)

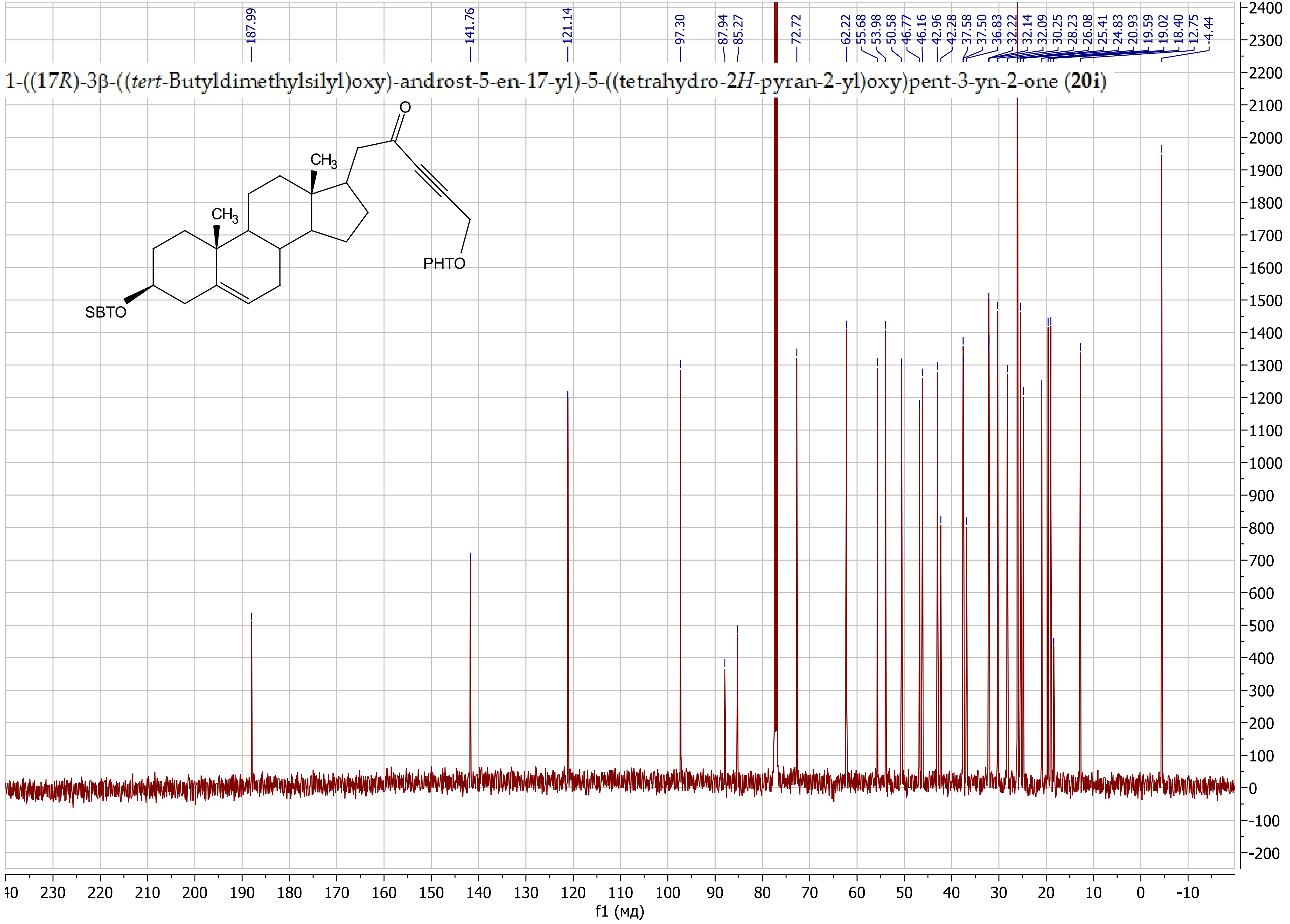

5-(((17*R*)-3β-((*tert*-Butyldimethylsilyl)oxy)-androst-5-en-17-yl)-4,5-dihydroisoxazol-5-ol (**22a**)

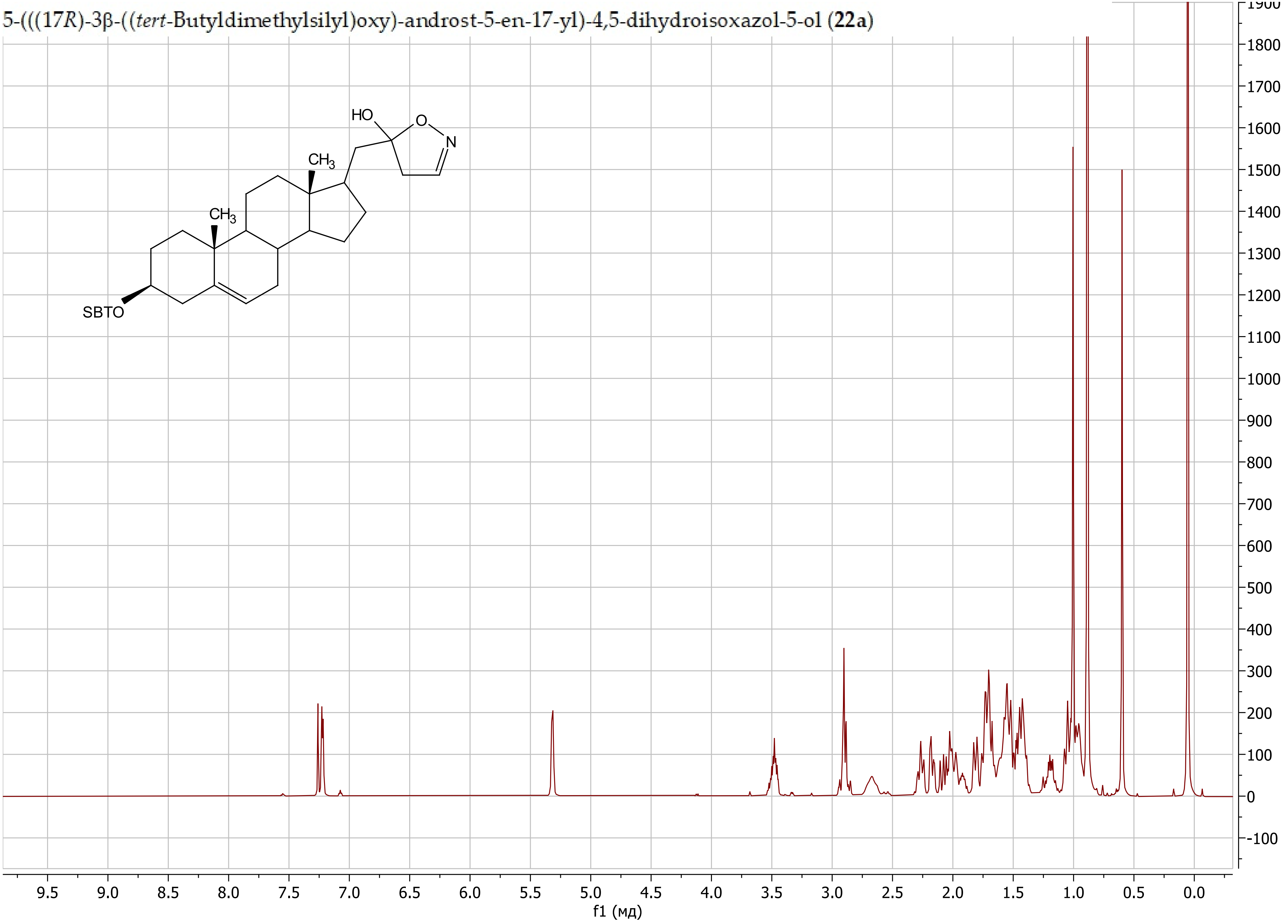

5-(((17*R*)-3β-((*tert*-Butyldimethylsilyl)oxy)-androst-5-en-17-yl)-4,5-dihydroisoxazol-5-ol (**22a**)

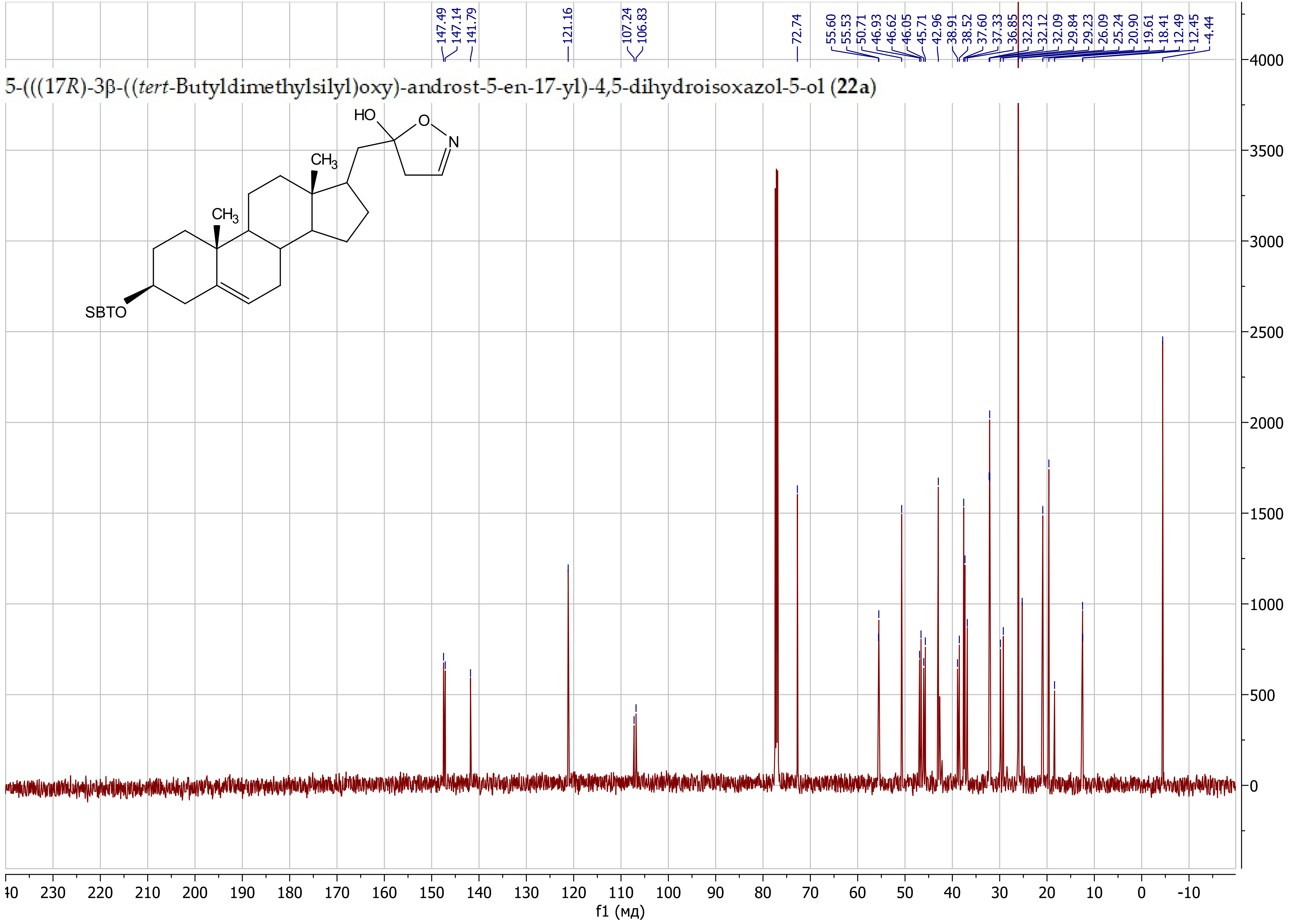

5-(((17*R*)-3β-((*tert*-Butyldimethylsilyl)oxy)-androst-5-en-17-yl)-3-isopropyl-4,5-dihydroisoxazol-5-ol (**22b**)

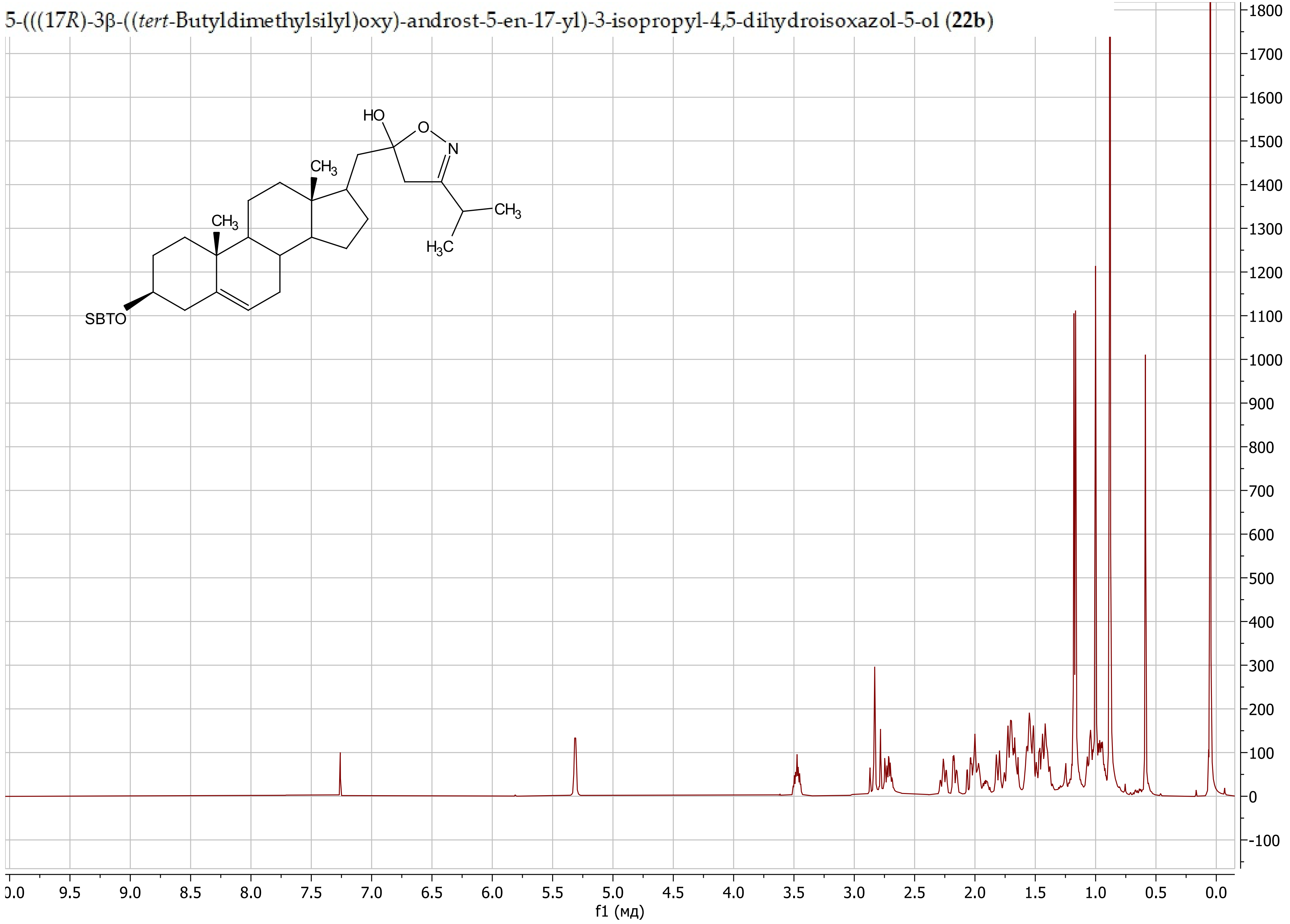

5-(((17*R*)-3β-((*tert*-Butyldimethylsilyl)oxy)-androst-5-en-17-yl)-3-isopropyl-4,5-dihydroisoxazol-5-ol (**22b**)

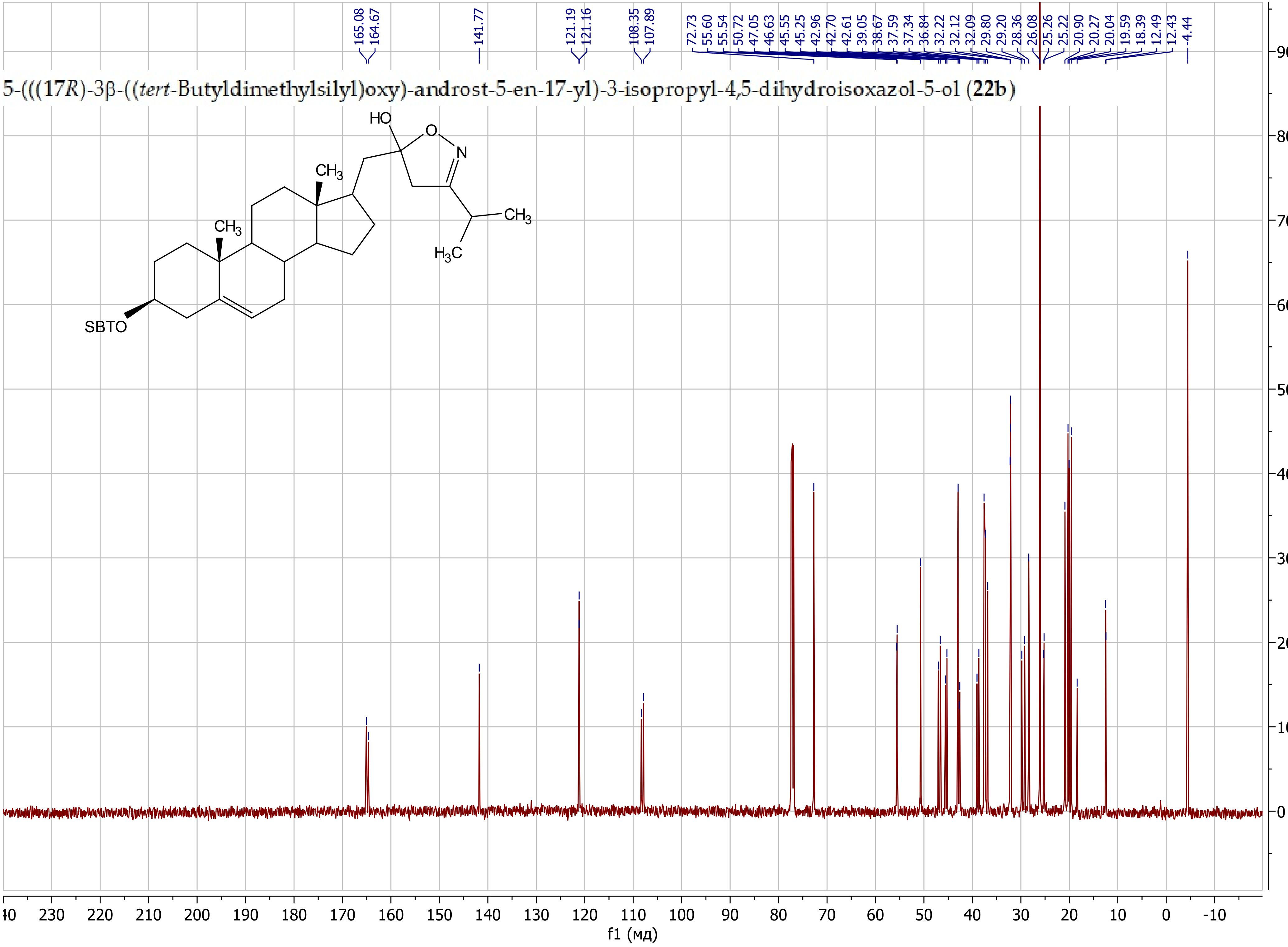

5-(((17*R*)-3β-((*tert*-Butyldimethylsilyl)oxy)-androst-5-en-17-yl)-3-cyclopropyl-4,5-dihydroisoxazol-5-ol (**22c**)

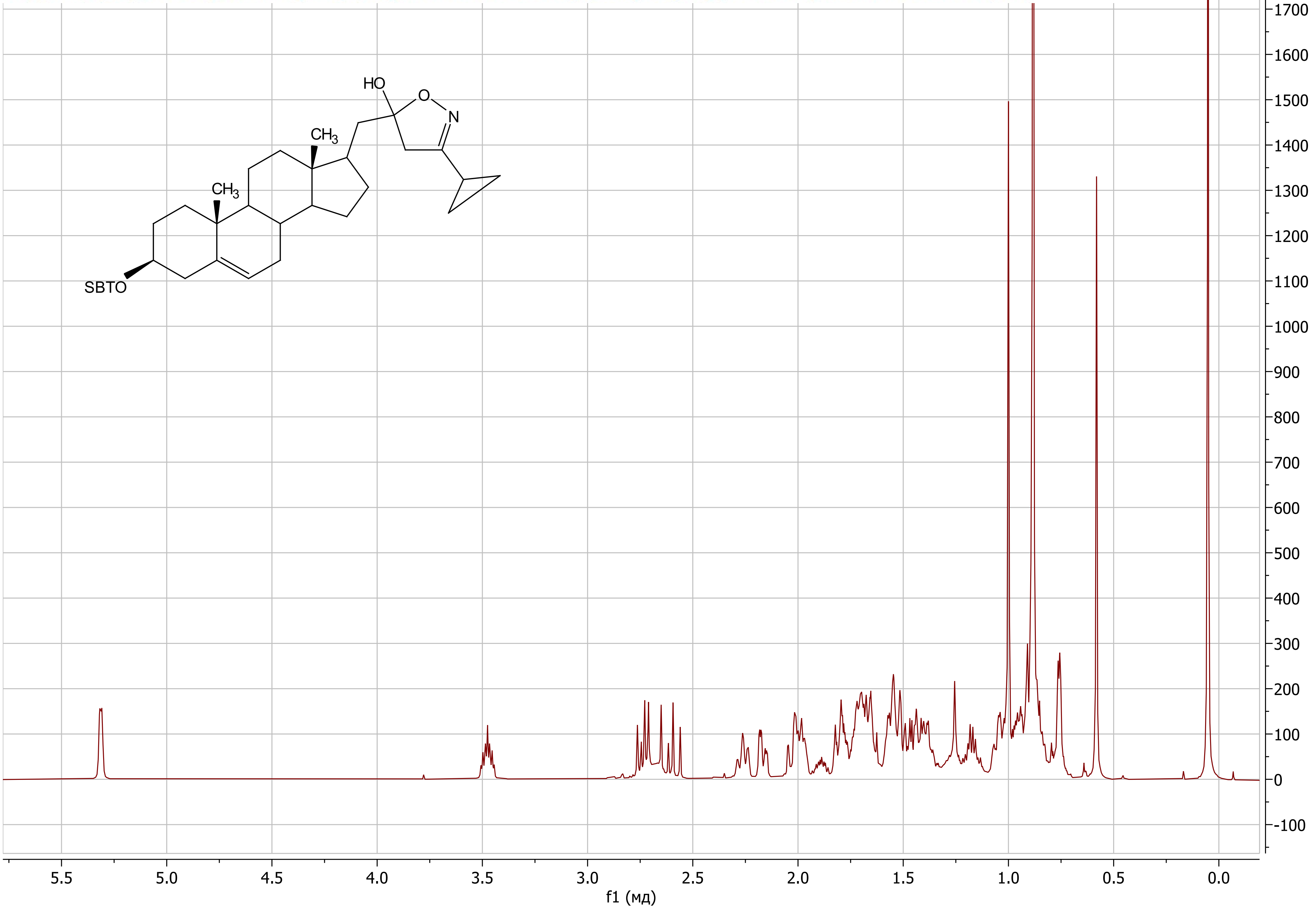

5-(((17*R*)-3β-((*tert*-Butyldimethylsilyl)oxy)-androst-5-en-17-yl)-3-cyclopropyl-4,5-dihydroisoxazol-5-ol (**22c**)

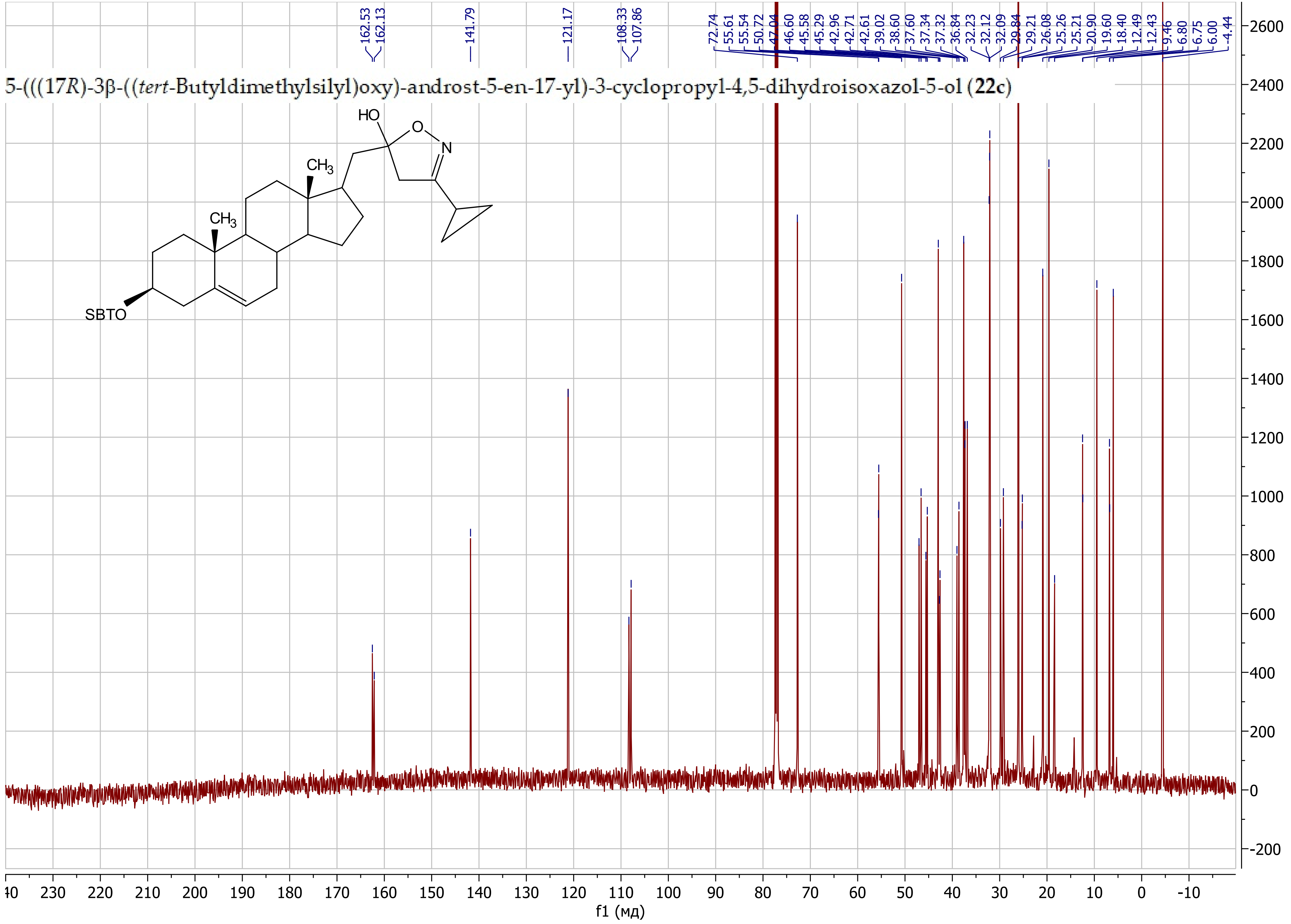

3-Butyl-5-(((17*R*)-3β-((*tert*-butyldimethylsilyl)oxy)-androst-5-en-17-yl)methyl)-4,5-dihydroisoxazol-5-ol (**22d**)

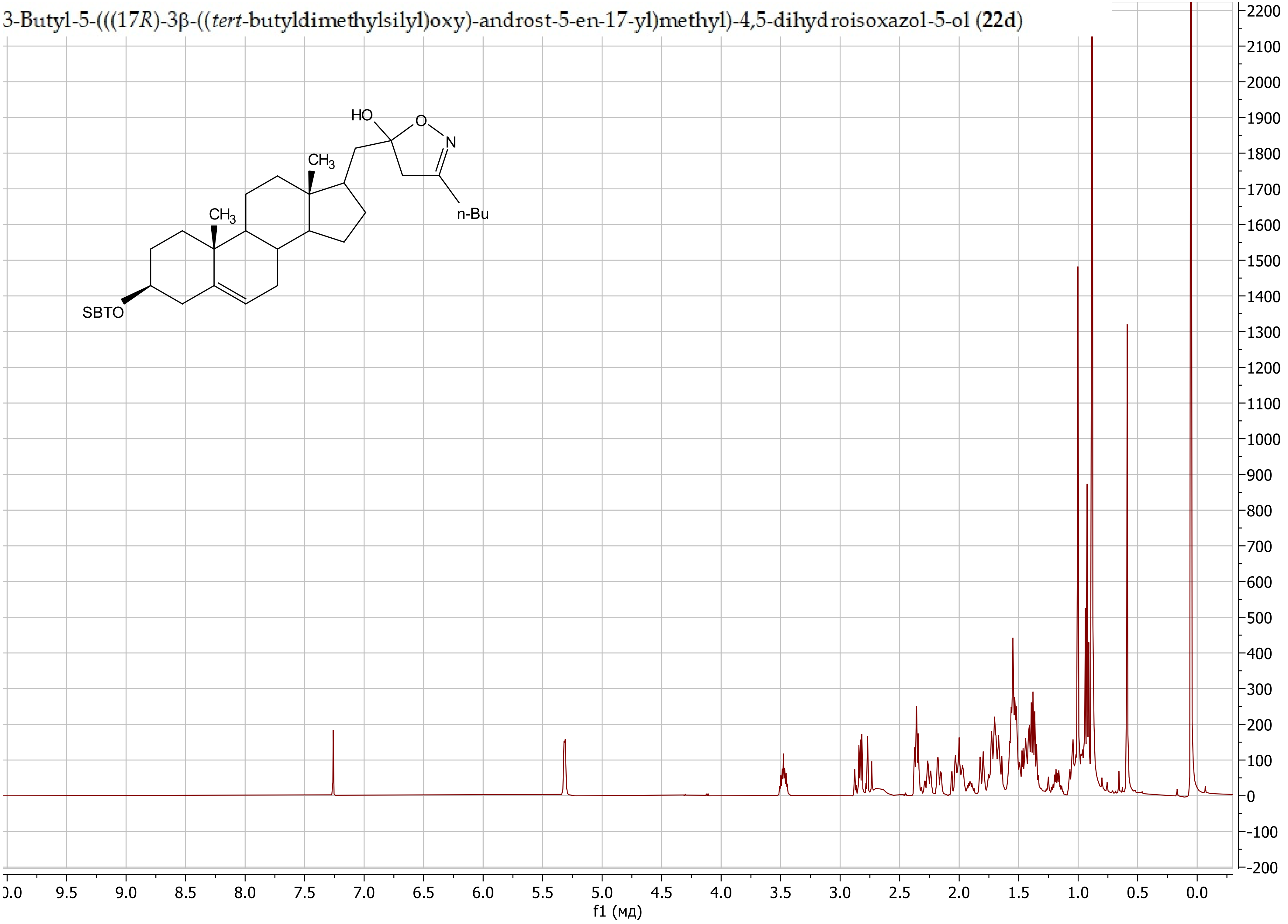

3-Butyl-5-(((17*R*)-3β-((*tert*-butyldimethylsilyl)oxy)-androst-5-en-17-yl)methyl)-4,5-dihydroisoxazol-5-ol (**22d**)

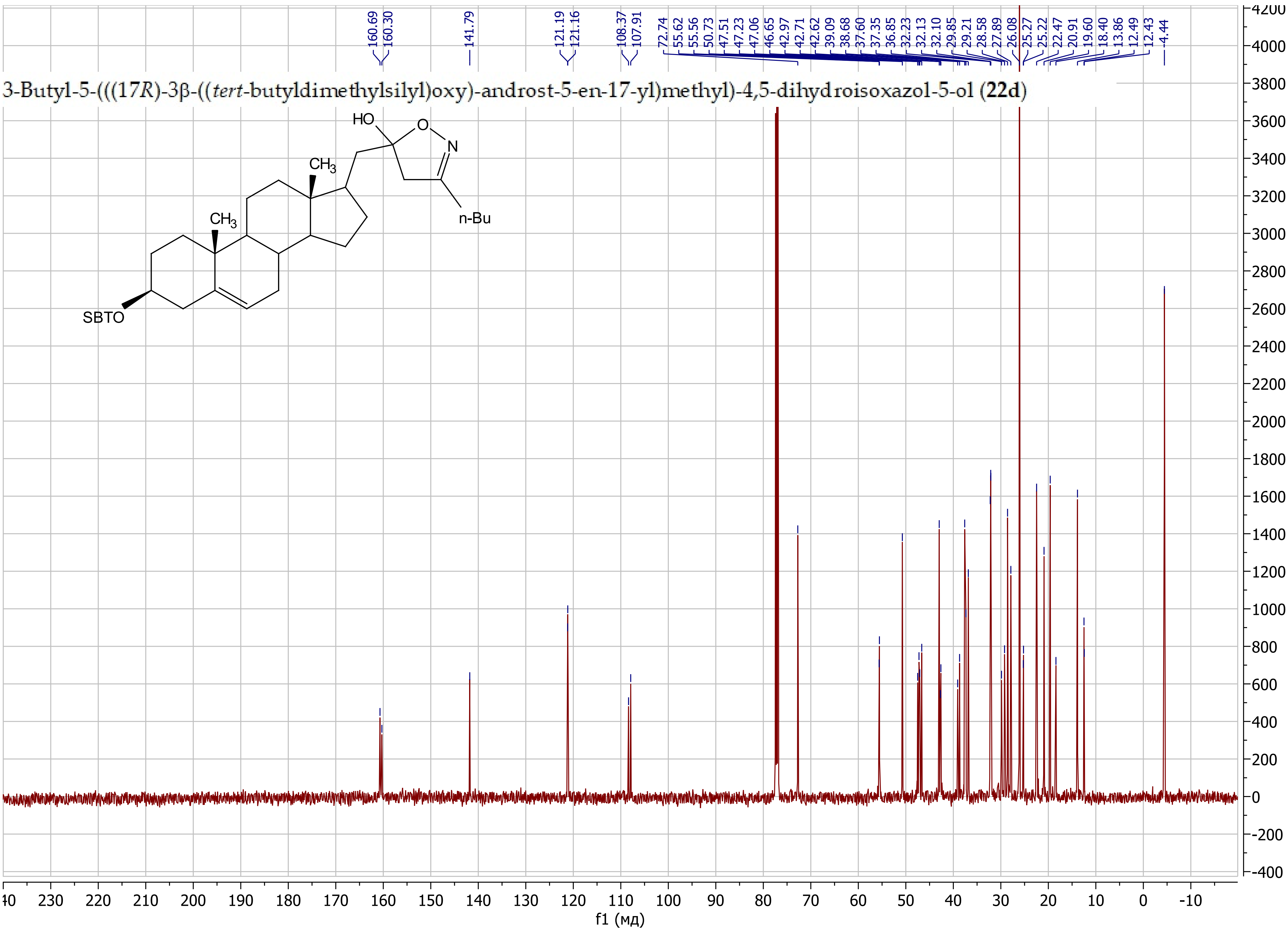

5-(((17R)-3β-((*tert*-Butyldimethylsilyl)oxy)-androst-5-en-17-yl)methyl)-3-phenyl-4,5-dihydroisoxazol-5-ol (**22e**)

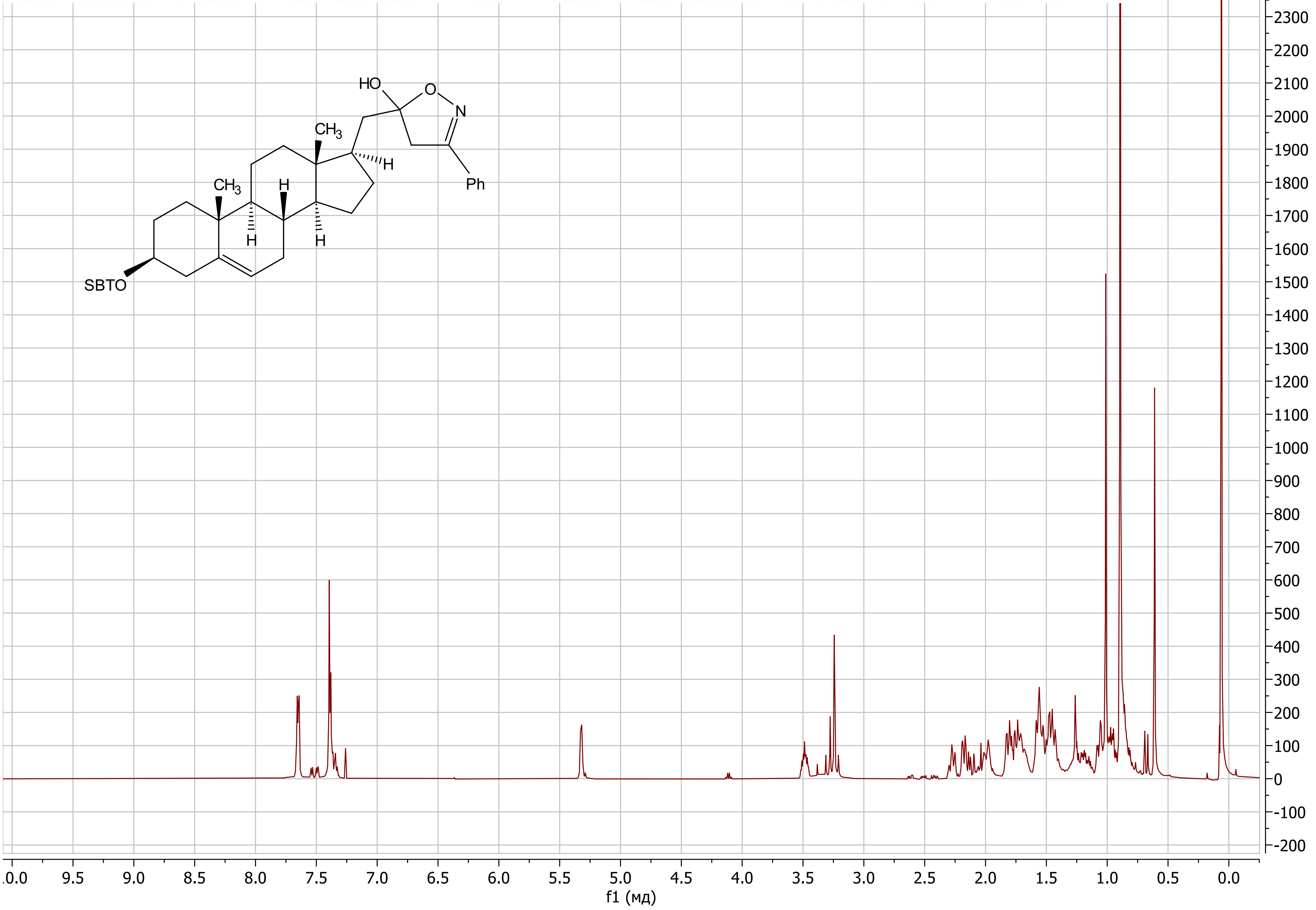

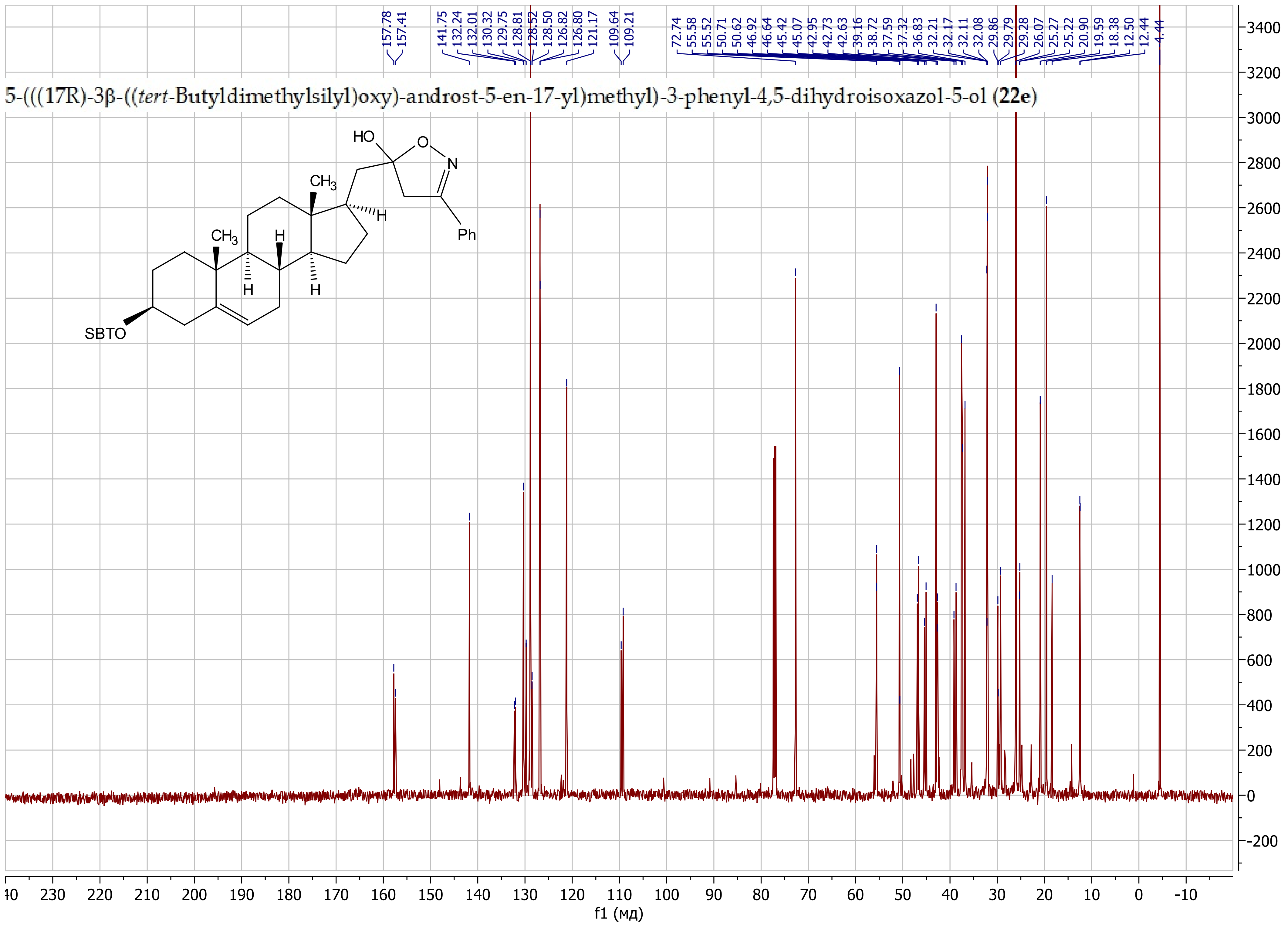
5-(((17R)-3β-((*tert*-Butyldimethylsilyl)oxy)-androst-5-en-17-yl)methyl)-3-phenyl-4,5-dihydroisoxazol-5-ol (**22e**)
157.78
157.41
141.75
132.24
132.01
130.32
129.75
128.81
128.52
128.50
126.82
126.80
121.17
109.64
109.21
72.74
55.58
55.52
50.71
50.62
46.92
46.64
45.42
45.07
42.95
42.73
42.63
39.16
38.72
37.59
37.32
36.83
32.21
32.17
32.11
32.08
29.86
29.79
29.28
26.07
25.27
25.22
20.90
19.59
18.38
12.50
12.44
4.44
HO
O
N
$CH_3$
H
Ph
SBTO
f1 (мд)

5-(((17R)-3β-((*tert*-Butyldimethylsilyl)oxy)-androst-5-en-17-yl)-3-(pyridin-3-yl)-4,5-dihydroisoxazol-5-ol (**22f**)

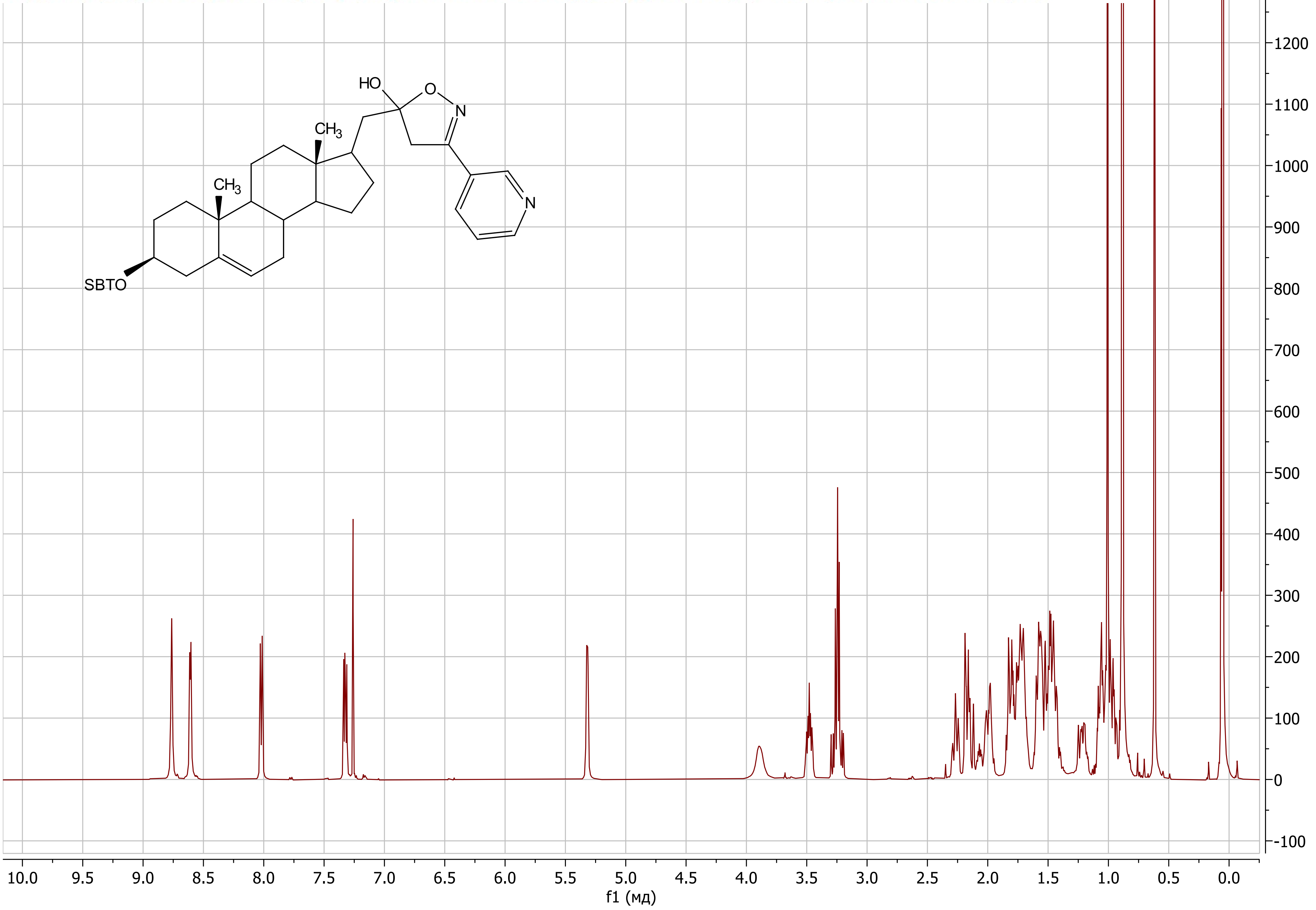

5-(((17R)-3β-((*tert*-Butyldimethylsilyl)oxy)-androst-5-en-17-yl)-3-(pyridin-3-yl)-4,5-dihydroisoxazol-5-ol (**22f**)

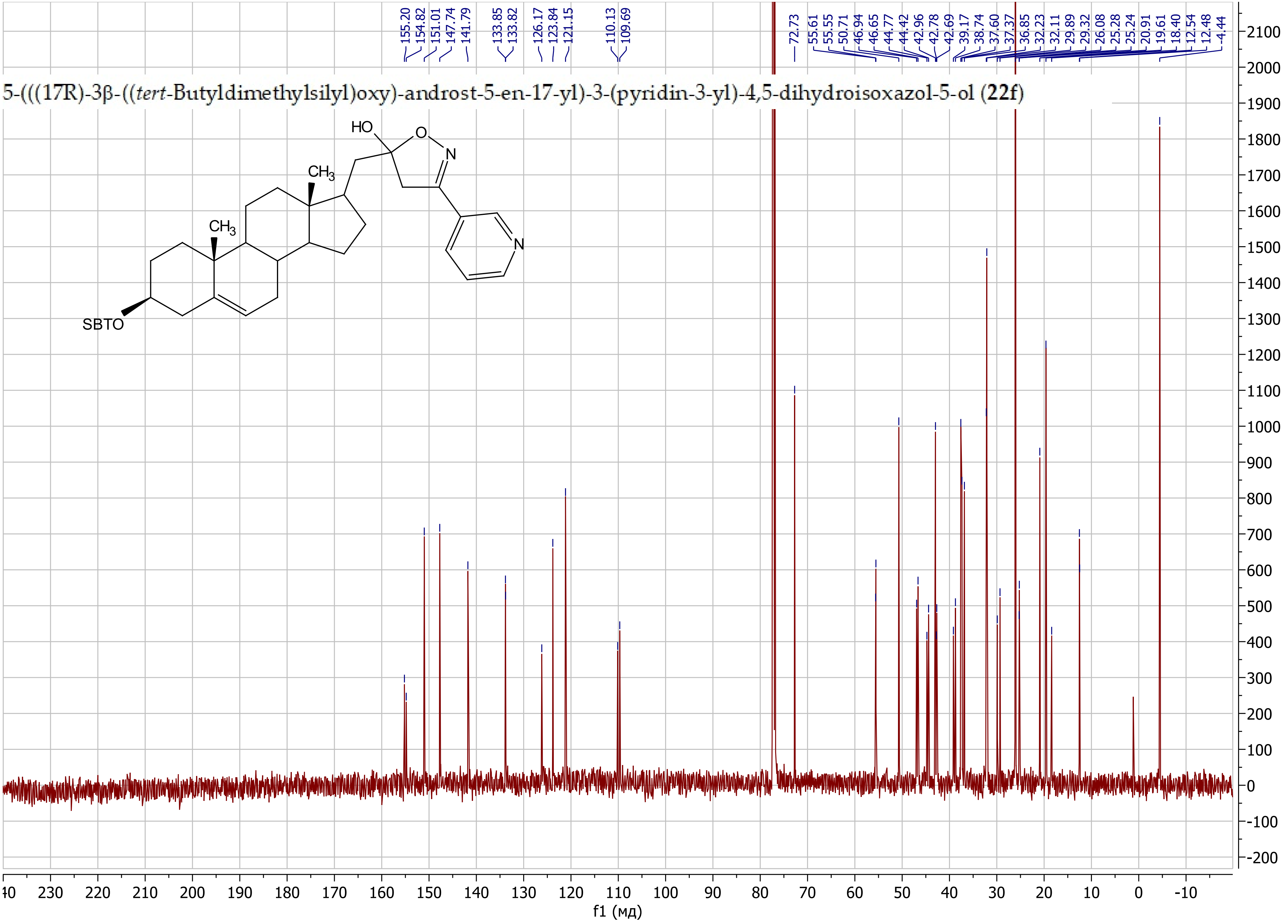

5-(((17R)-3β-((*tert*-Butyldimethylsilyl)oxy)-androst-5-en-17-yl)-3-(2-fluorophenyl)-4,5-dihydroisoxazol-5-ol (**22g**)

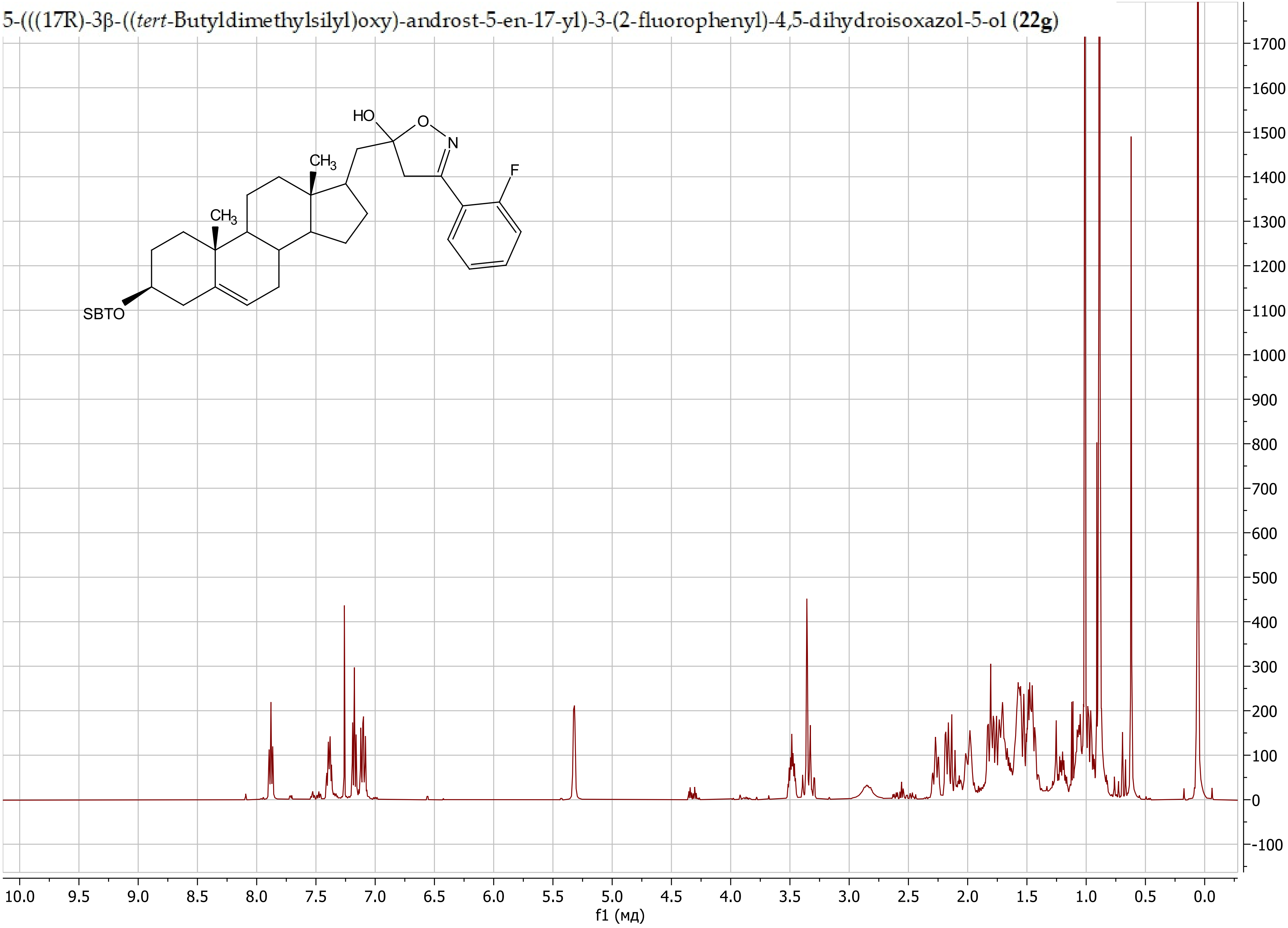

5-(((17R)-3β-((*tert*-Butyldimethylsilyl)oxy)-androst-5-en-17-yl)-3-(2-fluorophenyl)-4,5-dihydroisoxazol-5-ol (**22g**)

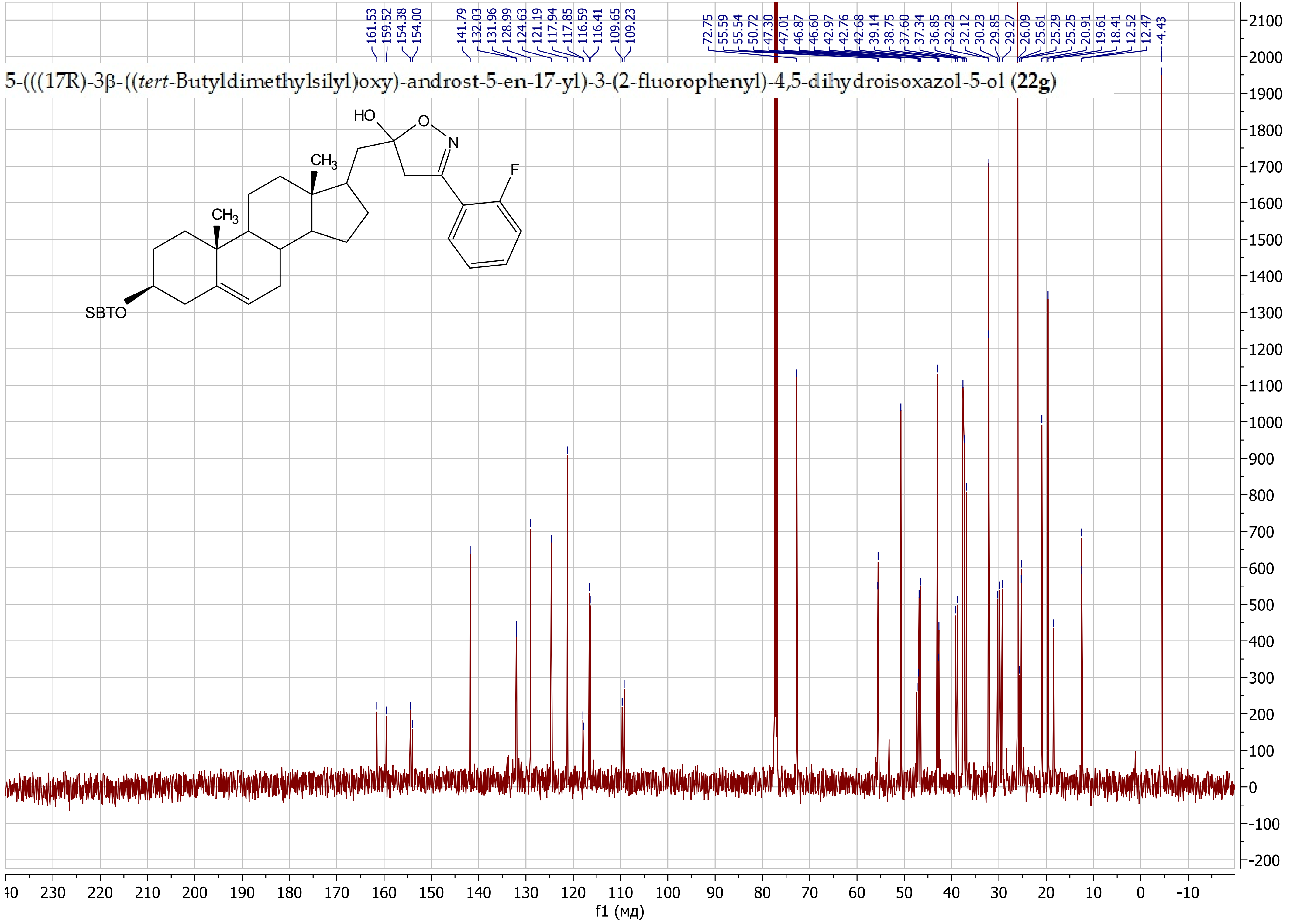

5-(((17R)-3β-((*tert*-Butyldimethylsilyl)oxy)-androst-5-en-17-yl)-3-(2-((tetrahydro-2*H*-pyran-2-yl)oxy)propan-2-yl)-4,5-dihydroisoxazol-5-ol (**22h**)

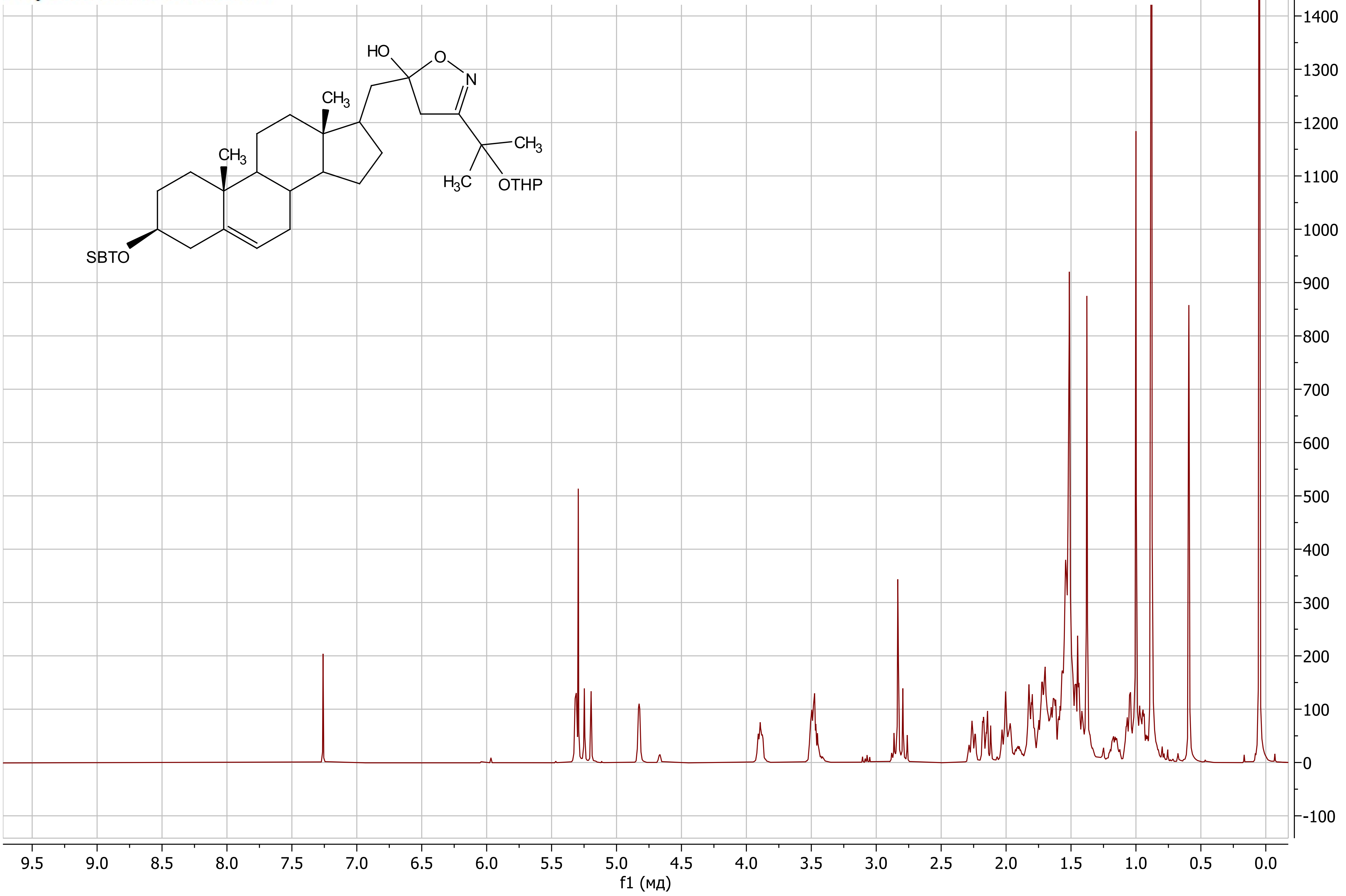

5-(((17R)-3β-((*tert*-Butyldimethylsilyl)oxy)-androst-5-en-17-yl)-3-(2-((tetrahydro-2*H*-pyran-2-yl)oxy)propan-2-yl)-4,5-dihydroisoxazol-5-ol (**22h**)

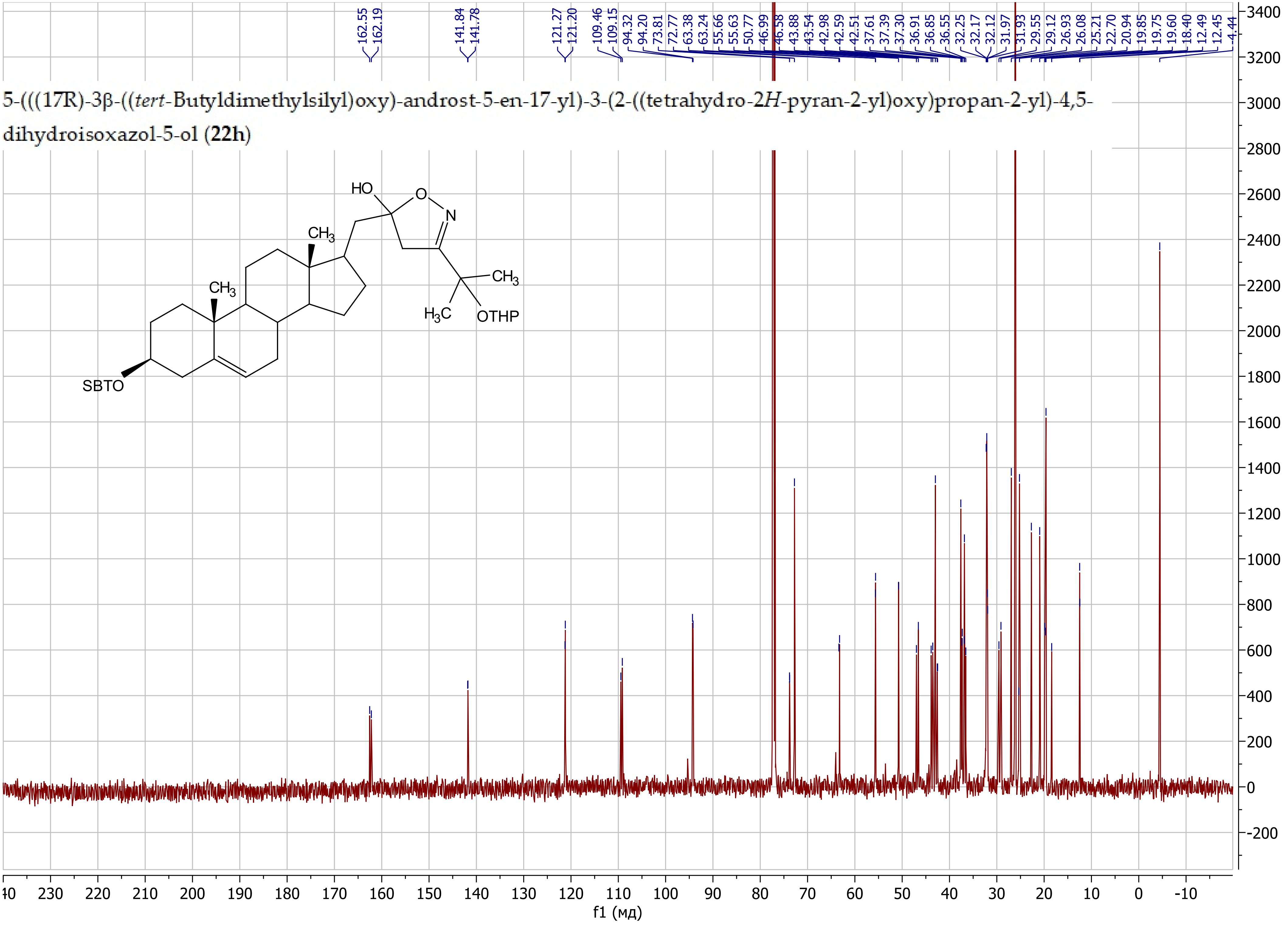

5-(((17R)-3β-((*tert*-Butyldimethylsilyl)oxy)-androst-5-en-17-yl)methyl)-3-(((tetrahydro-2*H*-pyran-2-yl)oxy)methyl)-4,5-dihydroisoxazol-5-ol (**22i**)

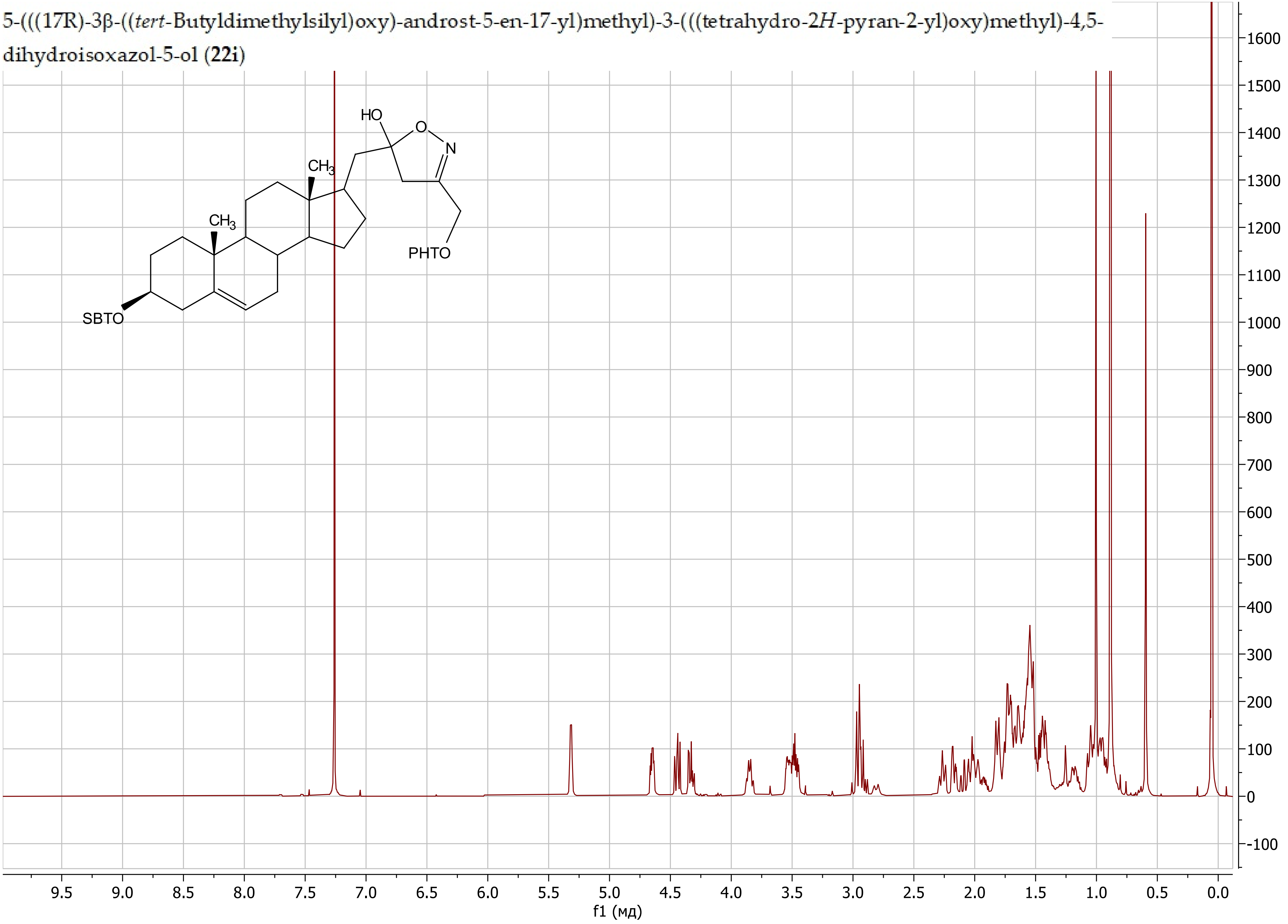

5-(((17R)-3β-((*tert*-Butyldimethylsilyl)oxy)-androst-5-en-17-yl)methyl)-3-(((tetrahydro-2*H*-pyran-2-yl)oxy)methyl)-4,5-dihydroisoxazol-5-ol (**22i**)

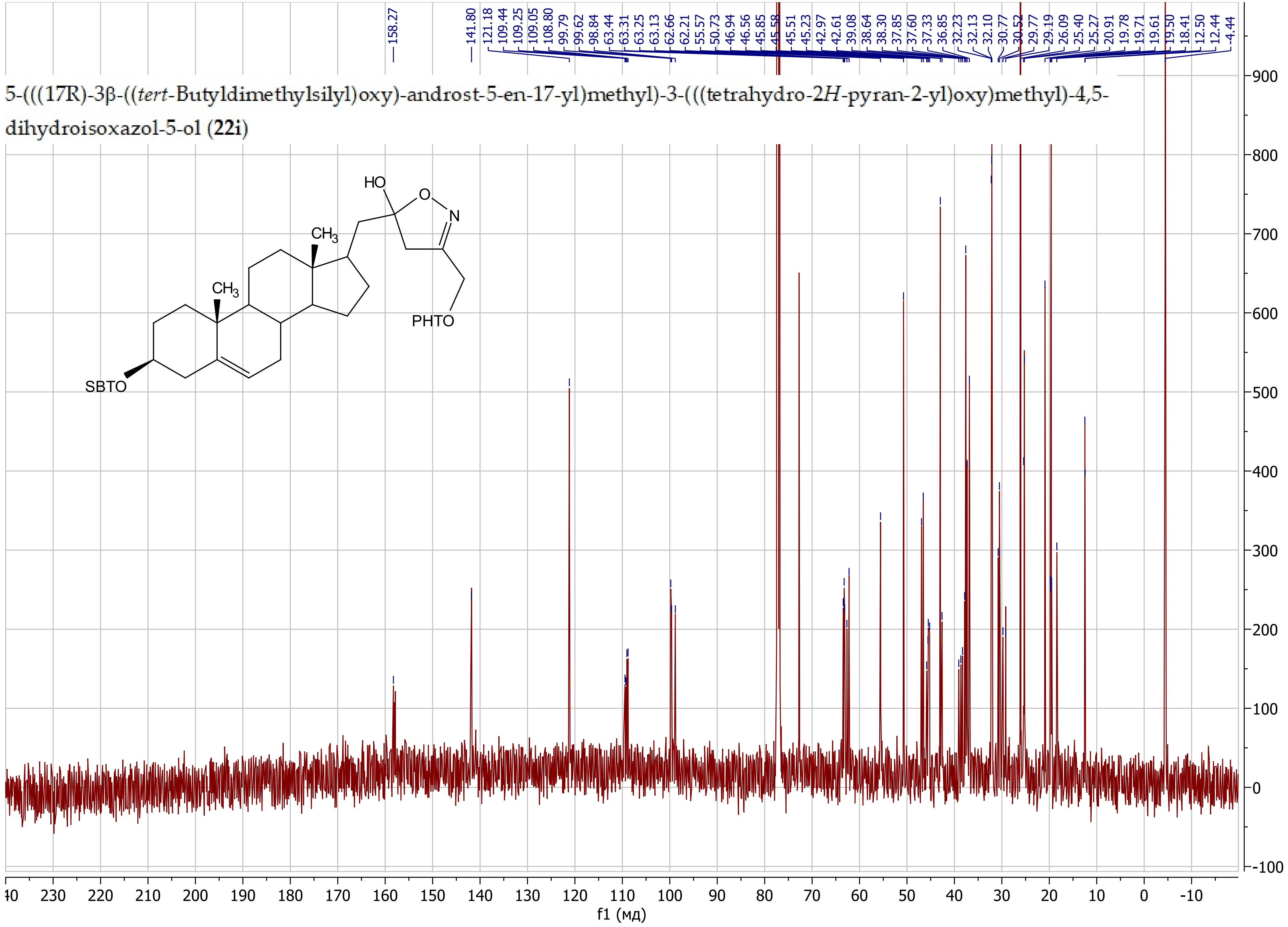

5-(((17*R*)-3β-((*tert*-butyldimethylsilyl)oxy)-androst-5-en-17-yl)methyl)isoxazole (**23a**)

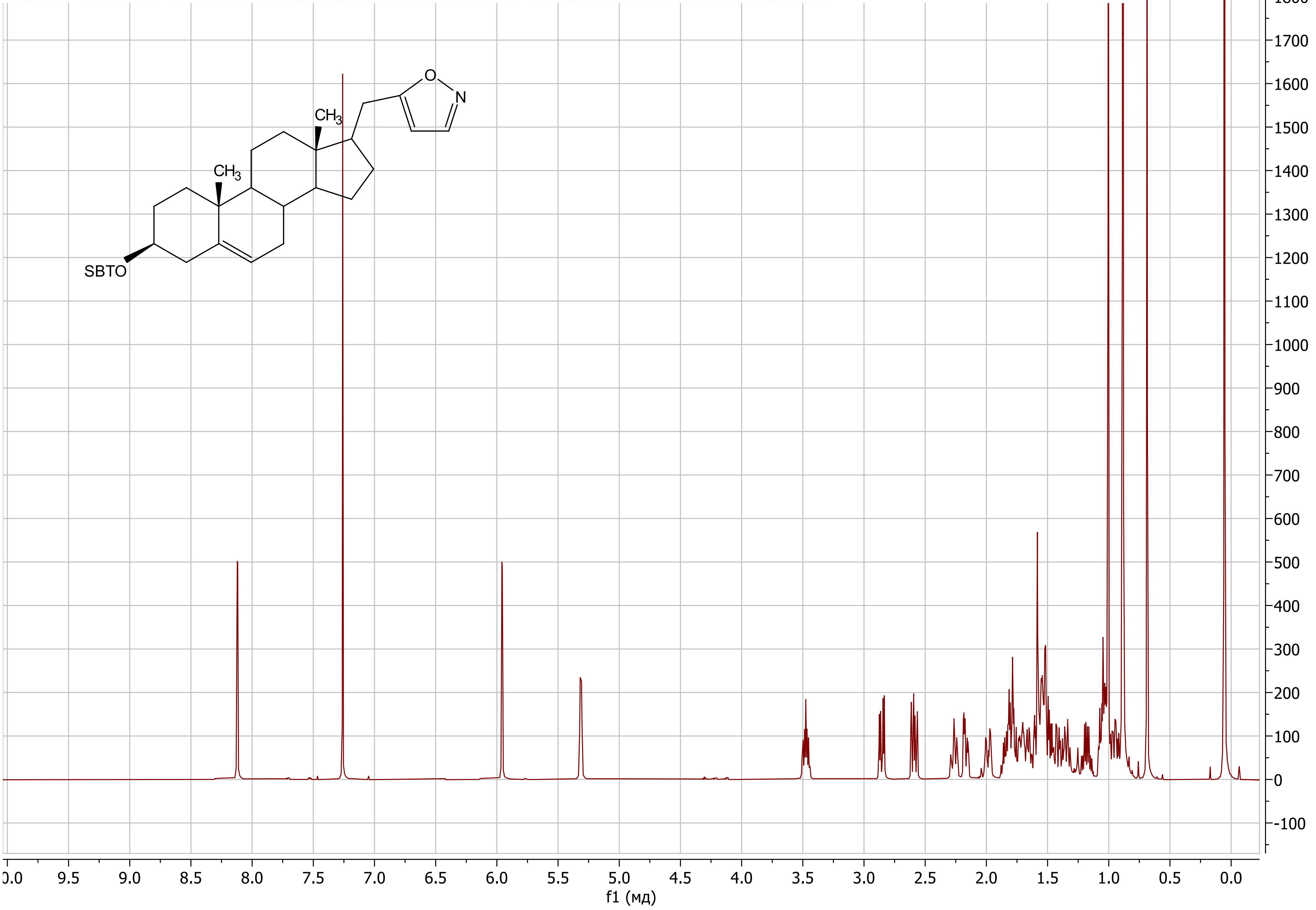

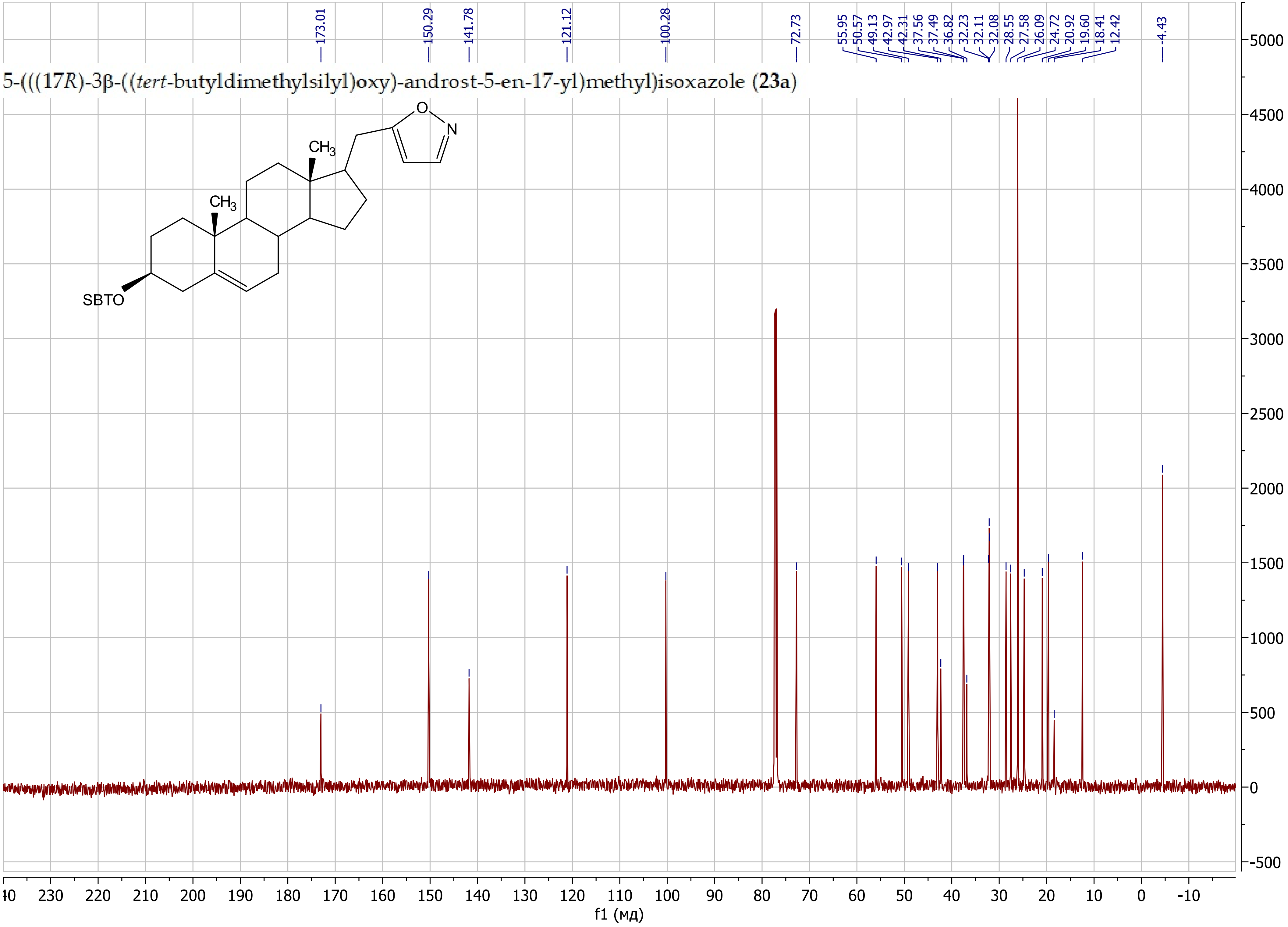

5-(((17R)-3β-((tert-butyldimethylsilyl)oxy)-androst-5-en-17-yl)methyl)isoxazole (23a)
173.01
150.29
141.78
121.12
100.28
72.73
55.95
50.57
49.13
42.97
42.31
37.56
37.49
36.82
32.23
32.11
32.08
28.55
27.58
26.09
24.72
20.92
19.60
18.41
12.42
-4.43
CH3
CH3
O
N
SBTO
f1 (мд)

5-(((17*R*)-3β-((*tert*-Butyldimethylsilyl)oxy)-androst-5-en-17-yl)methyl)-3-isopropylisoxazole (**23b**)

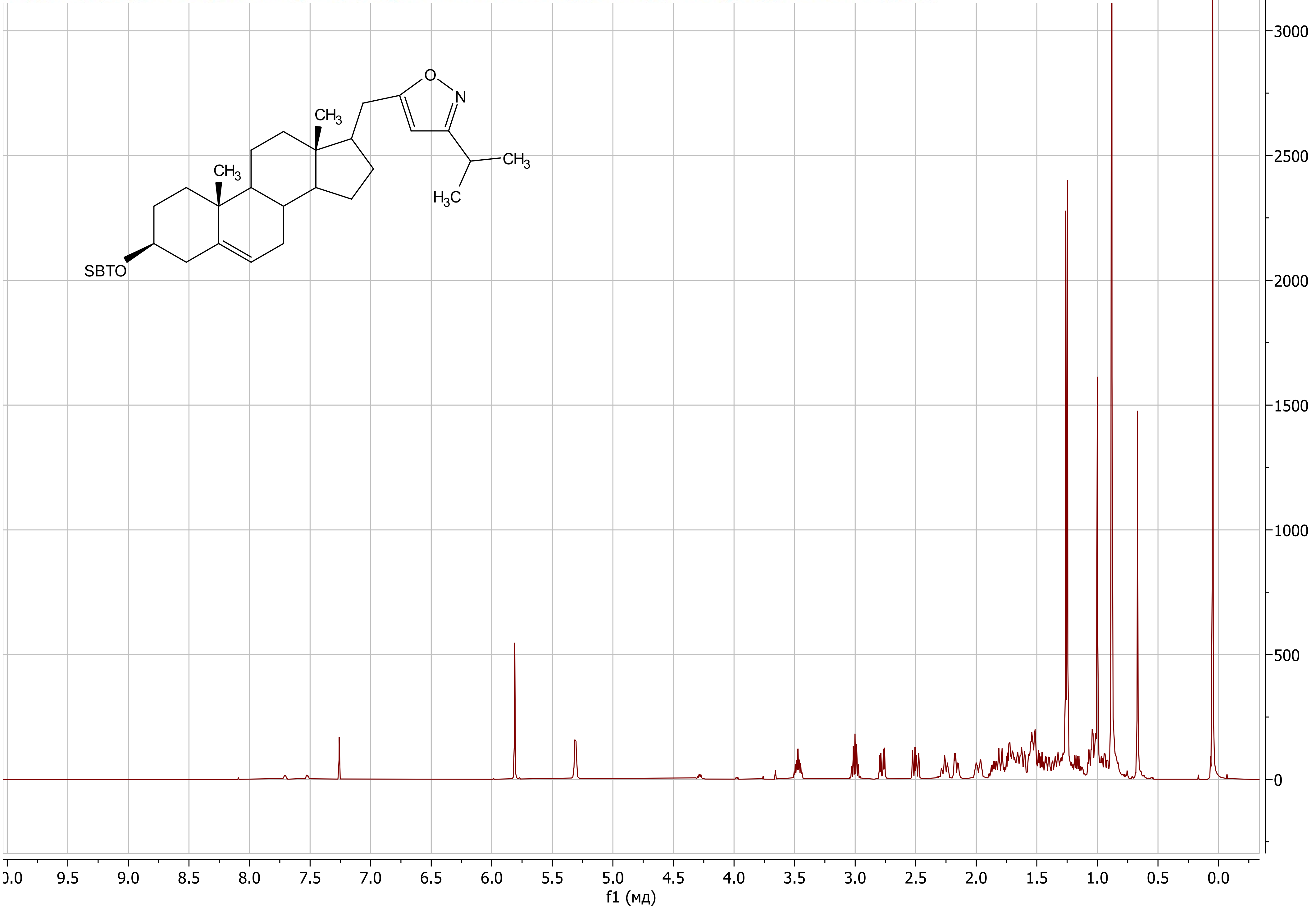

5-(((17*R*)-3β-((*tert*-Butyldimethylsilyl)oxy)-androst-5-en-17-yl)methyl)-3-isopropylisoxazole (**23b**)

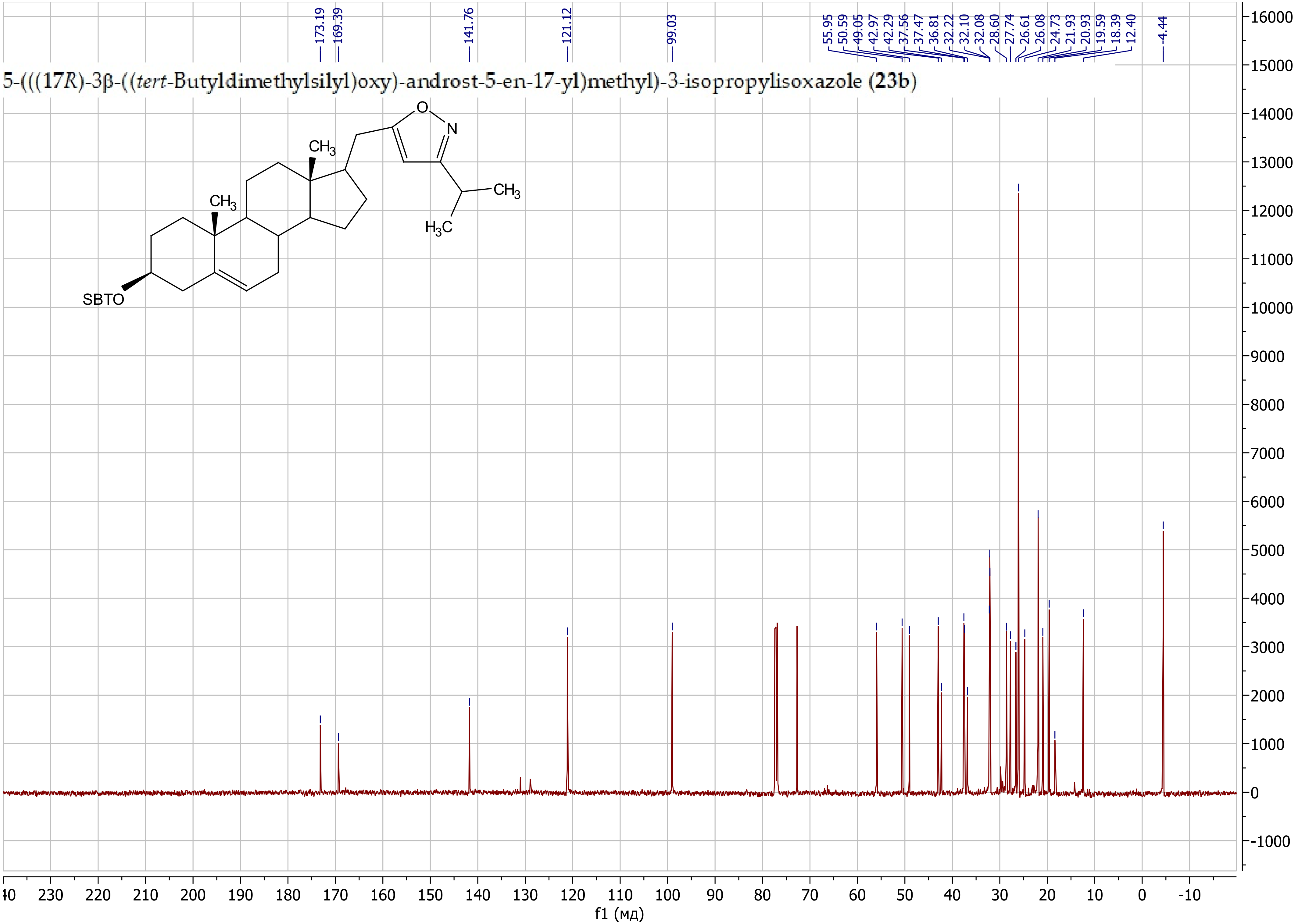

5-(((17*R*)-3β-((*tert*-Butyldimethylsilyl)oxy)-androst-5-en-17-yl)methyl)-3-cyclopropylisoxazole (**23c**)

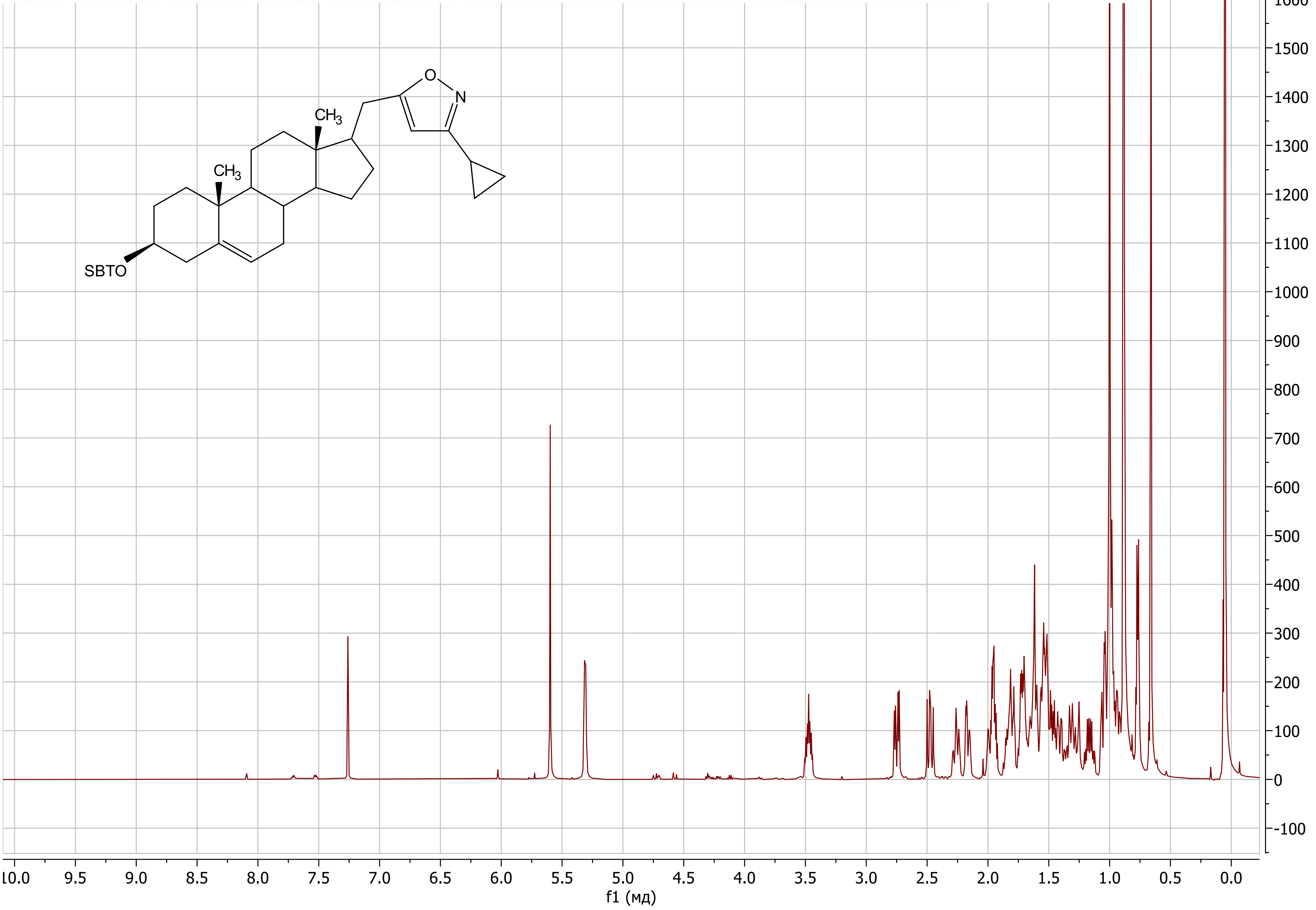

#### 5-(((17*R*)-3β-((*tert*-Butyldimethylsilyl)oxy)-androst-5-en-17-yl)methyl)-3-cyclopropylisoxazole (**23c**)

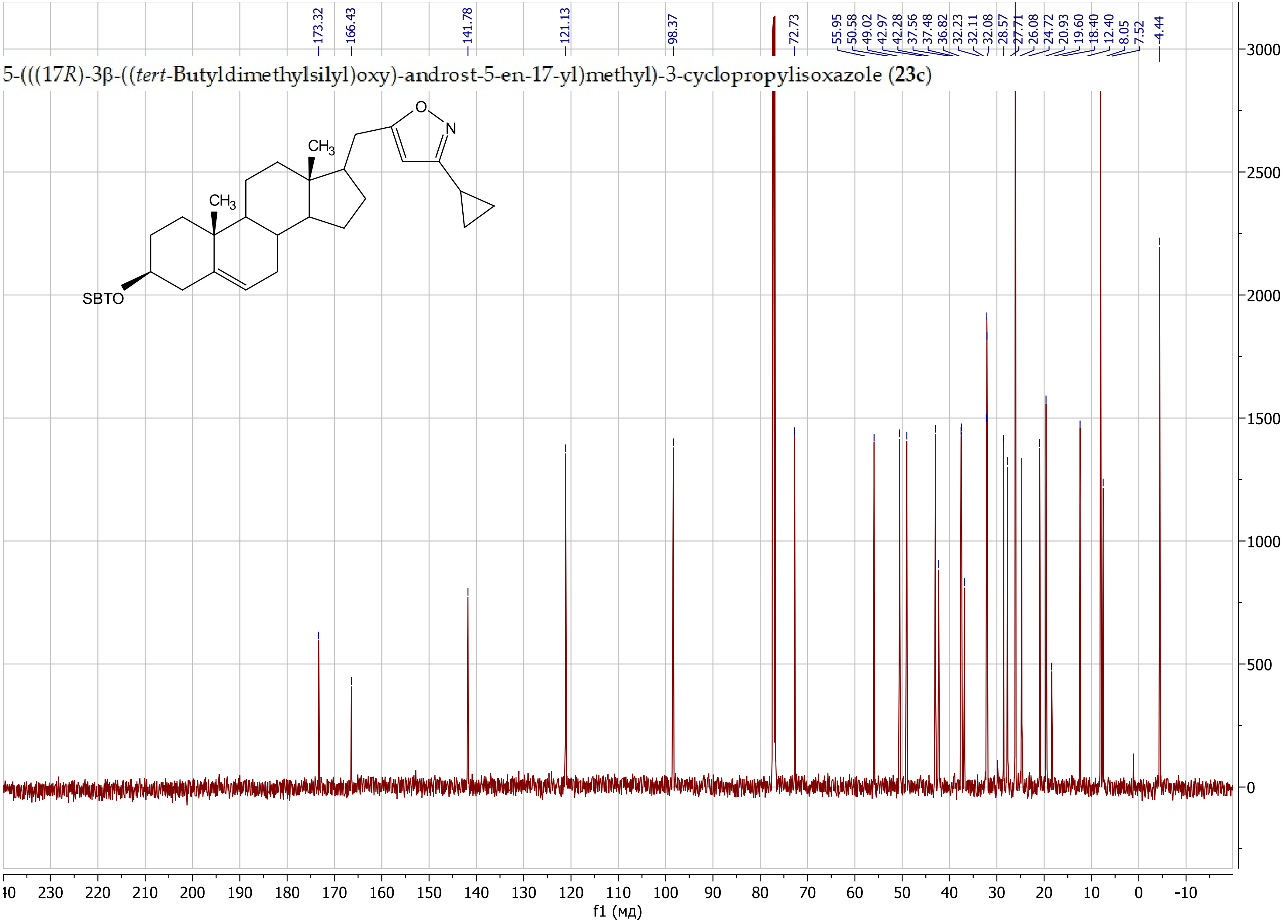

3-Butyl-5-(((17*R*)-3β-((*tert*-butyldimethylsilyl)oxy)-androst-5-en-17-yl)methyl)isoxazole (**23d**)

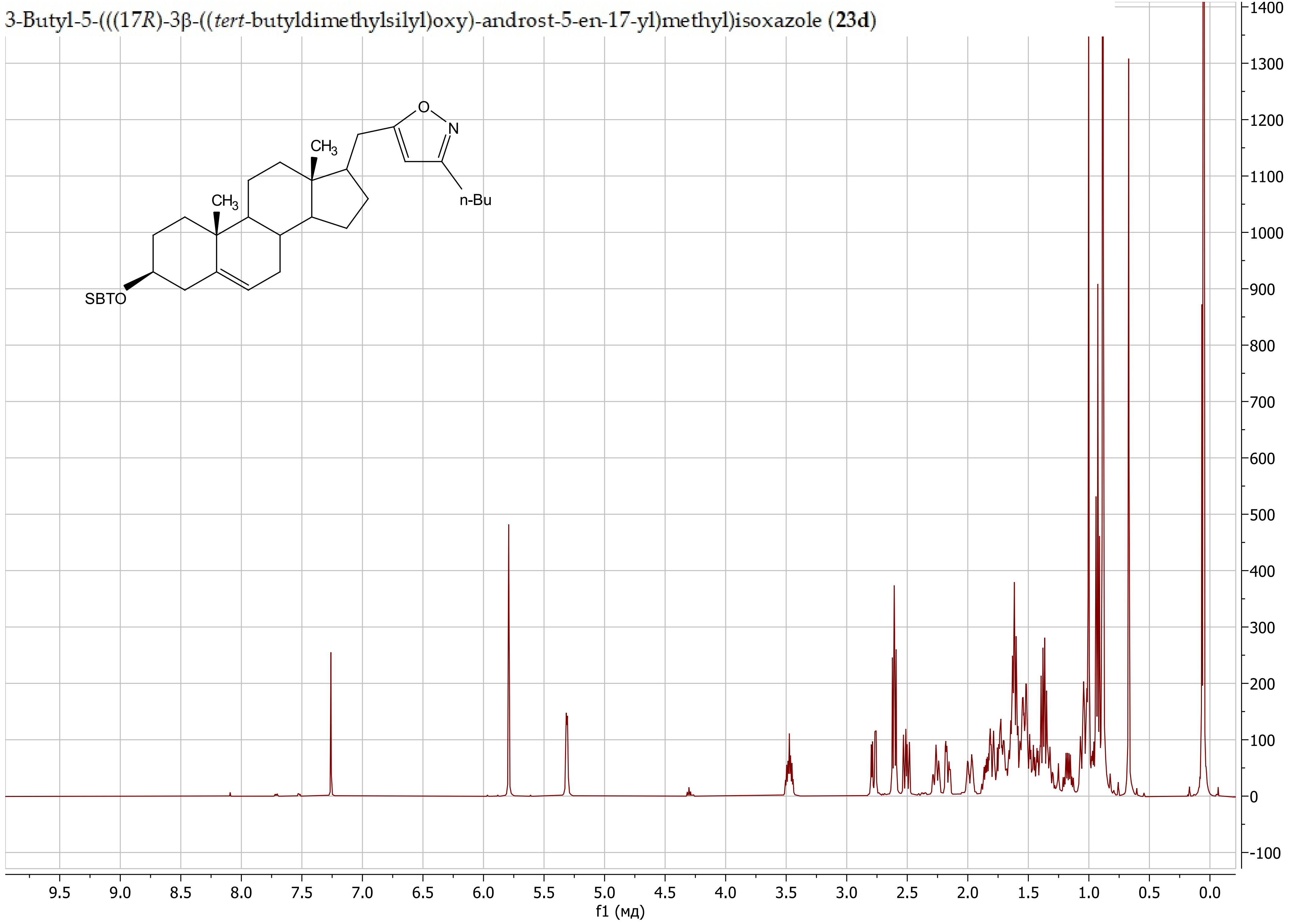

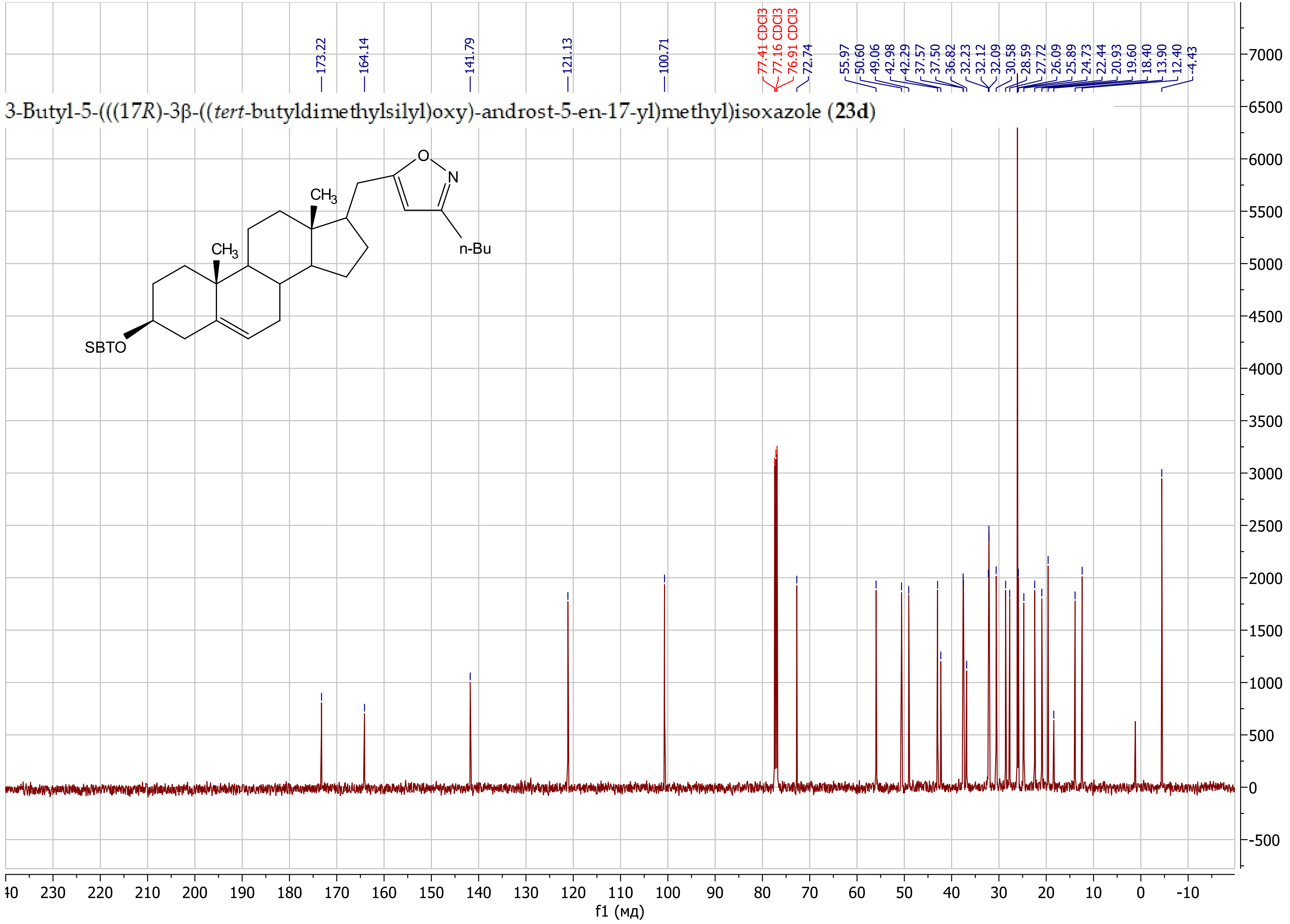
3-Butyl-5-(((17*R*)-3β-((*tert*-butyldimethylsilyl)oxy)-androst-5-en-17-yl)methyl)isoxazole (**23d**)
173.22
164.14
141.79
121.13
100.71
77.41 CDCl3
77.16 CDCl3
76.91 CDCl3
72.74
55.97
50.60
49.06
42.98
42.29
37.57
37.50
36.82
32.23
32.12
32.09
30.58
28.59
27.72
26.09
25.89
24.73
22.44
20.93
19.60
18.40
13.90
12.40
-4.43
O
N
n-Bu
$CH_3$
$CH_3$
SBTO
f1 (мд)

5-(((17*R*)-3β-((*tert*-Butyldimethylsilyl)oxy)-androst-5-en-17-yl)methyl)-3-phenylisoxazole (**23e**)

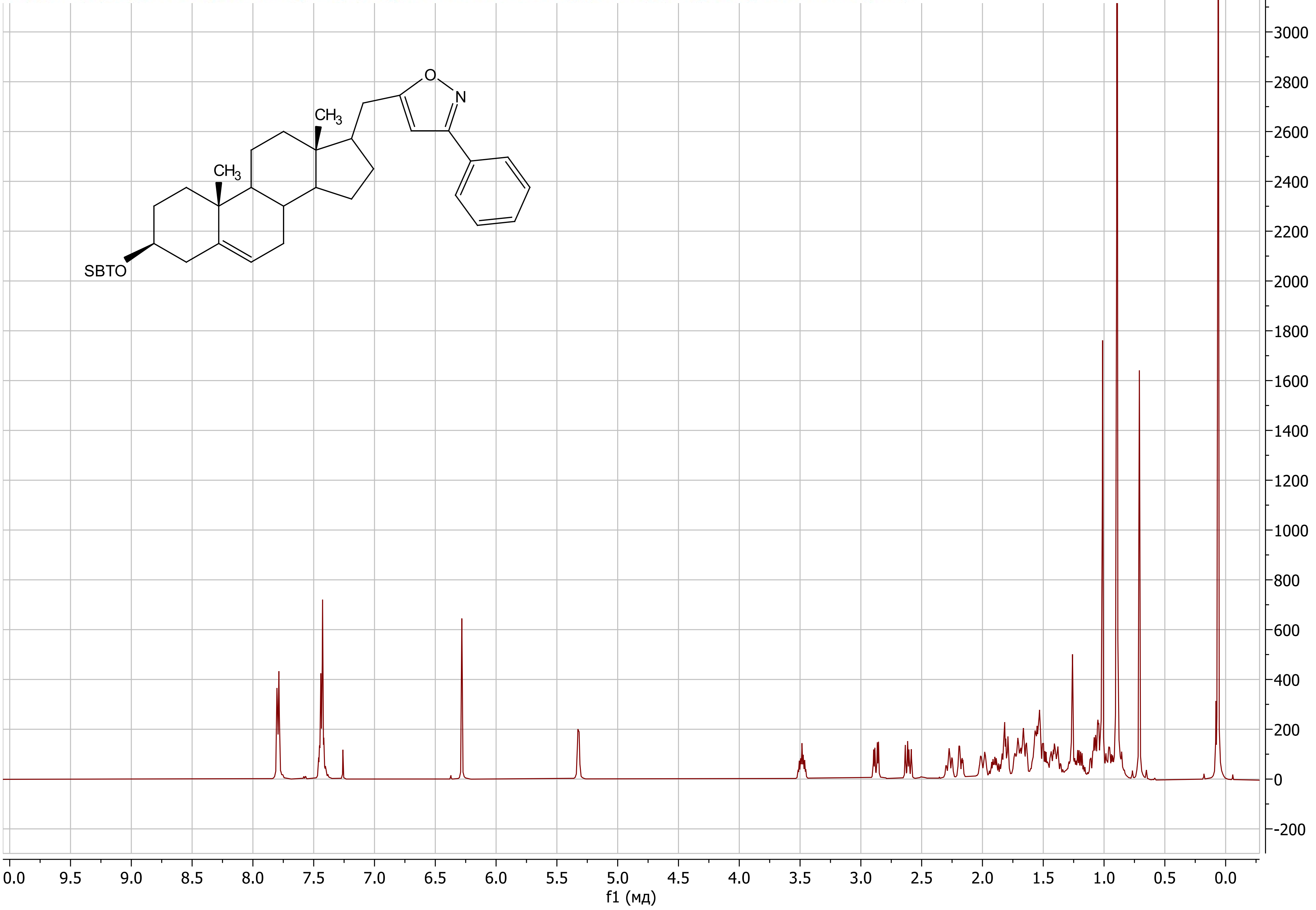

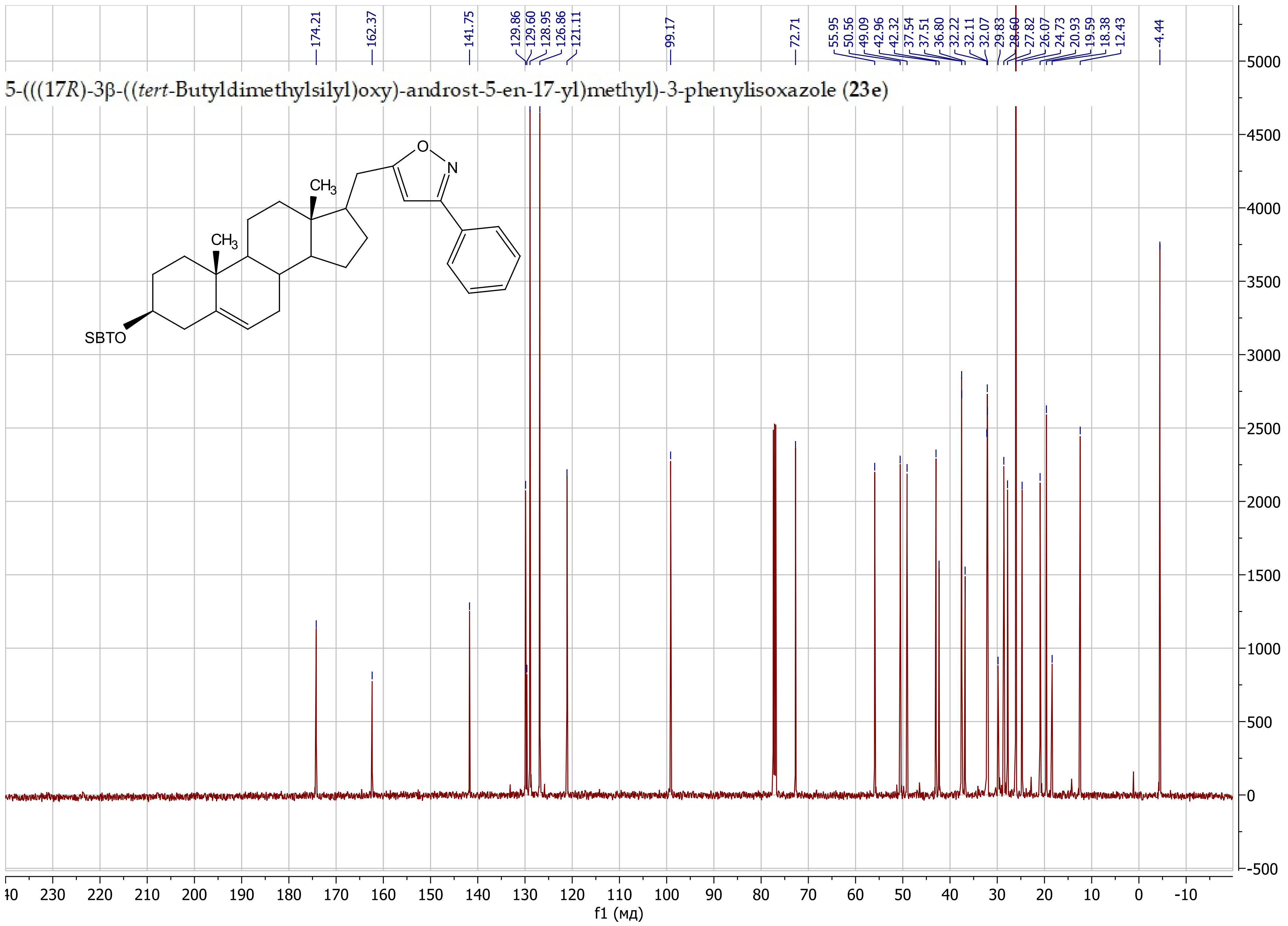
5-(((17*R*)-3β-((*tert*-Butyldimethylsilyl)oxy)-androst-5-en-17-yl)methyl)-3-phenylisoxazole (**23e**)
174.21
162.37
141.75
129.86
129.60
128.95
126.86
121.11
99.17
72.71
55.95
50.56
49.09
42.96
42.32
37.54
37.51
36.80
32.22
32.11
32.07
29.83
28.60
27.82
26.07
24.73
20.93
19.59
18.38
12.43
-4.44
O
N
CH3
CH3
SBTO
f1 (мд)

5-(((17*R*)-3β-((*tert*-Butyldimethylsilyl)oxy)-androst-5-en-17-yl)methyl)-3-(pyridin-3-yl)isoxazole (**23f**)

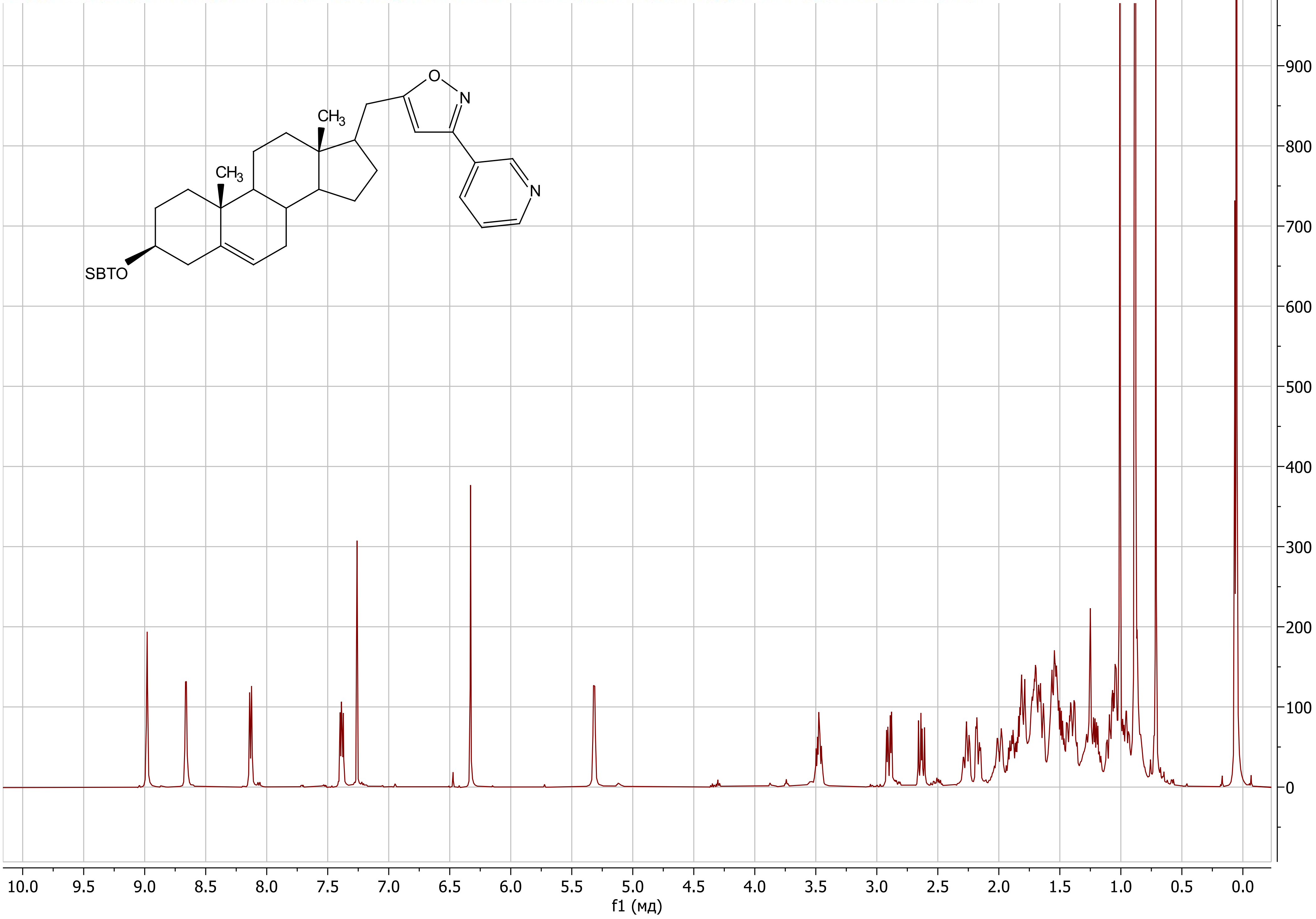

#### 5-(((17*R*)-3β-((*tert*-Butyldimethylsilyl)oxy)-androst-5-en-17-yl)methyl)-3-(pyridin-3-yl)isoxazole (**23f**)

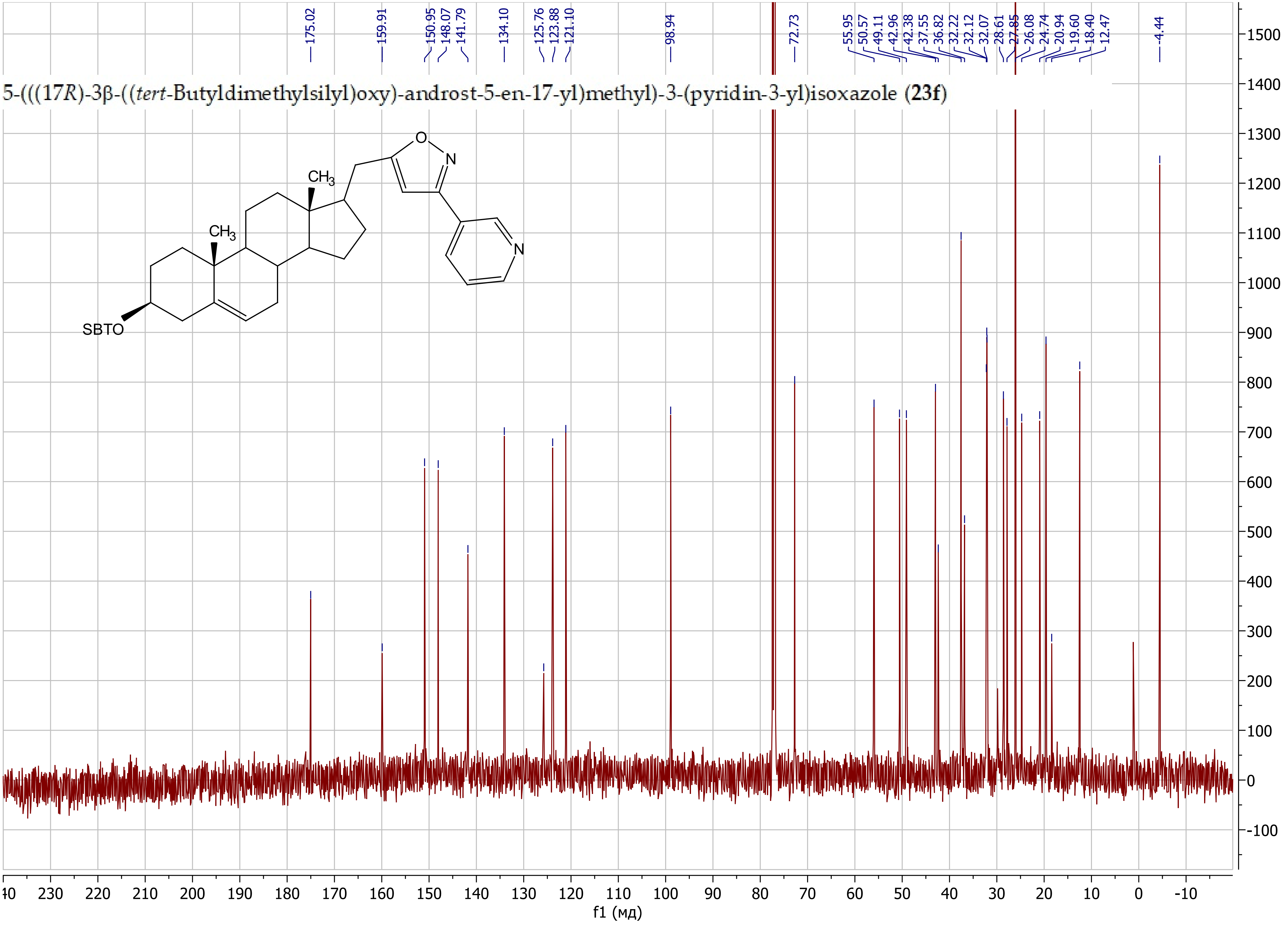

5-(((17*R*)-3β-((*tert*-Butyldimethylsilyl)oxy)-androst-5-en-17-yl)methyl)-3-(2-fluorophenyl)isoxazole (**23g**)

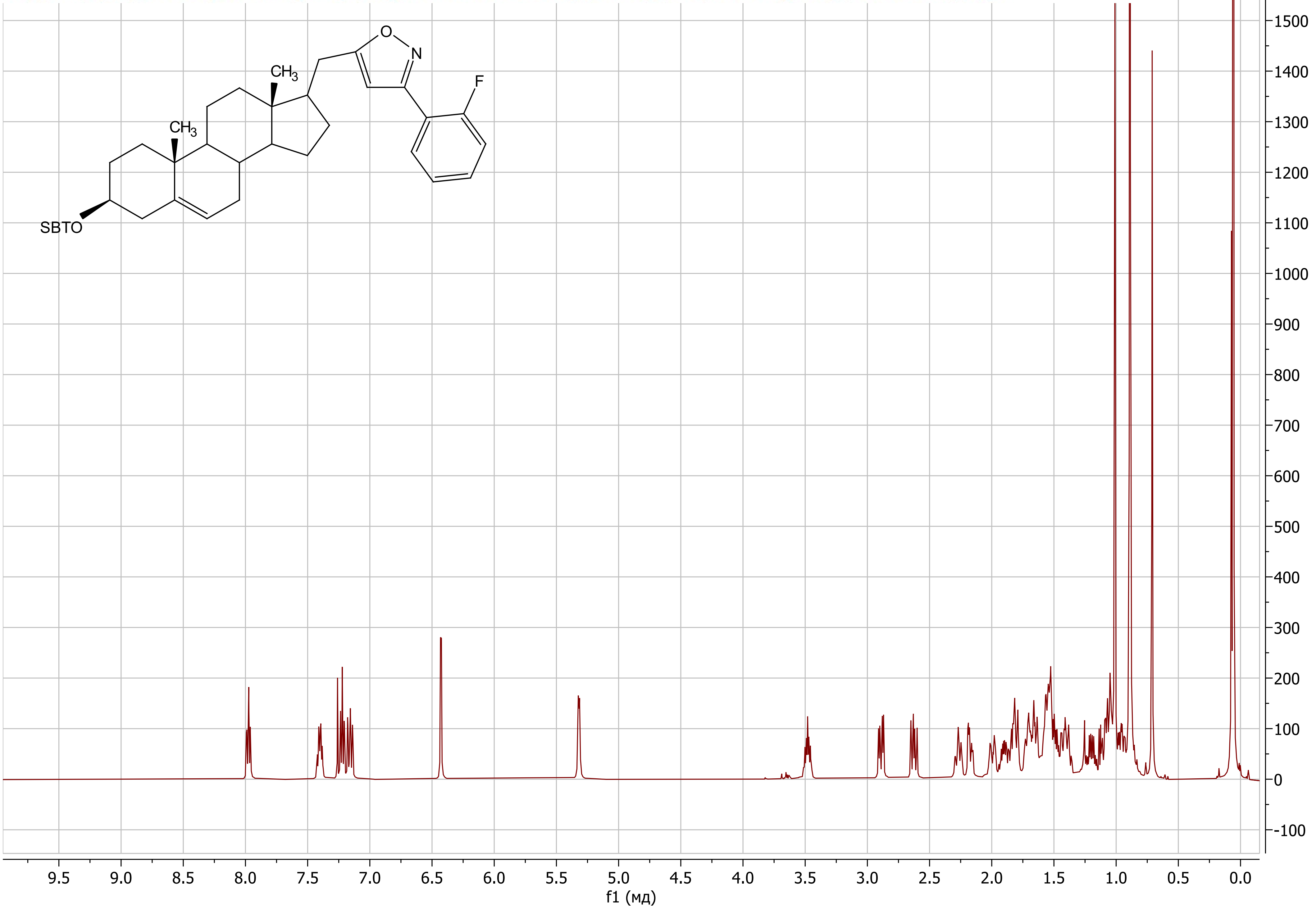

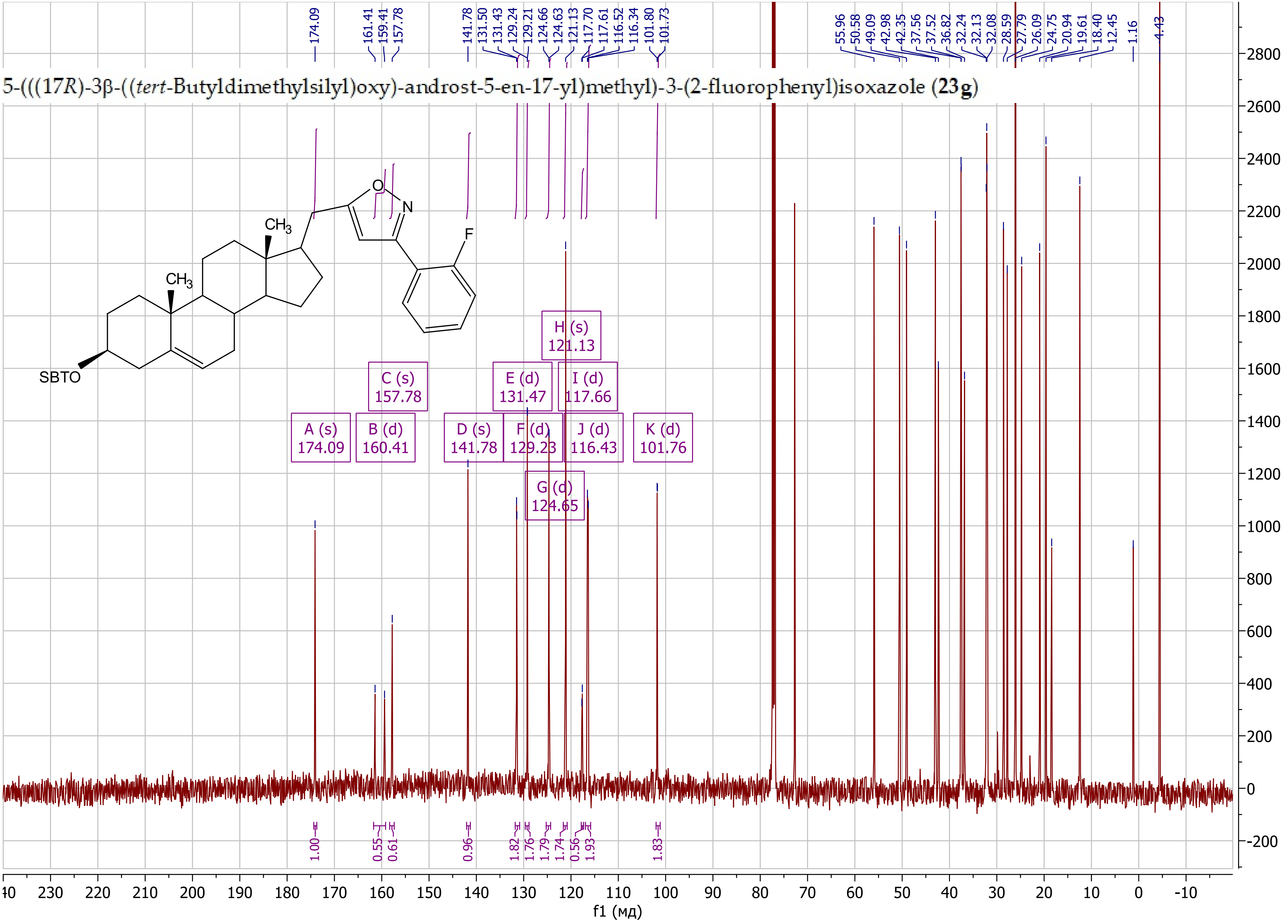
5-(((17R)-3β-((tert-Butyldimethylsilyl)oxy)-androst-5-en-17-yl)methyl)-3-(2-fluorophenyl)isoxazole (23g)
174.09
161.41
159.41
157.78
141.78
131.50
131.43
129.24
129.21
124.66
124.63
121.13
117.70
117.61
116.52
116.34
101.80
101.73
55.96
50.58
49.09
42.98
42.35
37.56
37.52
36.82
32.24
32.13
32.08
28.59
27.79
26.09
24.75
20.94
19.61
18.40
12.45
1.16
-4.43
SBTO
CH3
CH3
O
N
F
A (s) 174.09
B (d) 160.41
C (s) 157.78
D (s) 141.78
E (d) 131.47
F (d) 129.23
G (d) 124.65
H (s) 121.13
I (d) 117.66
J (d) 116.43
K (d) 101.76
1.00
0.55
0.61
0.96
1.82
1.76
1.79
1.74
0.56
1.93
1.83
f1 (мд)

5-(((17*R*)-3β-((*tert*-Butyldimethylsilyl)oxy)-androst-5-en-17-yl)-3-(2-((tetrahydro-2*H*-pyran-2-yl)oxy)propan-2-yl)isoxazole (**23h**)

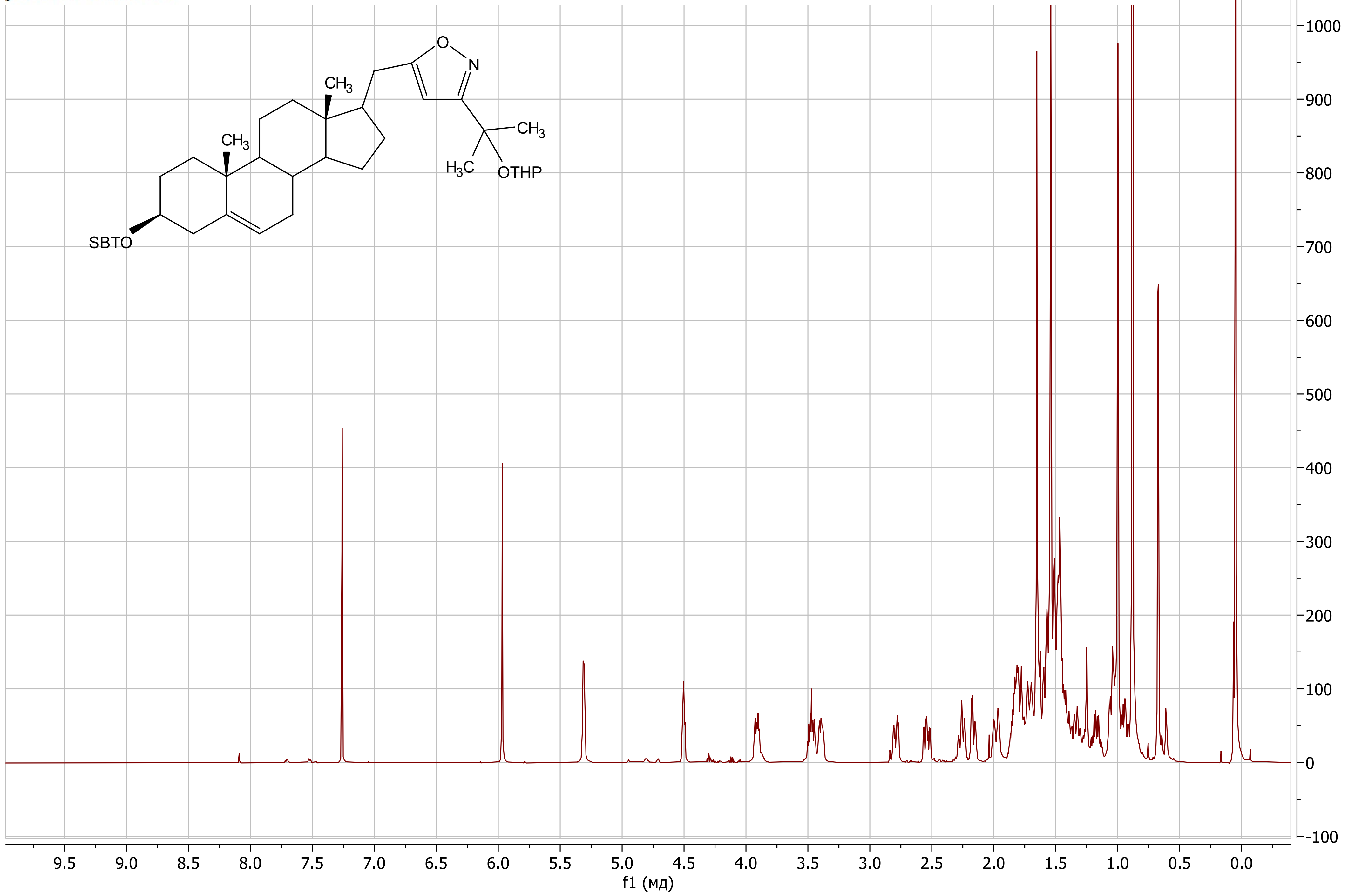

5-(((17*R*)-3β-((*tert*-Butyldimethylsilyl)oxy)-androst-5-en-17-yl)-3-(2-((tetrahydro-2*H*-pyran-2-yl)oxy)propan-2-yl)isoxazole (**23h**)

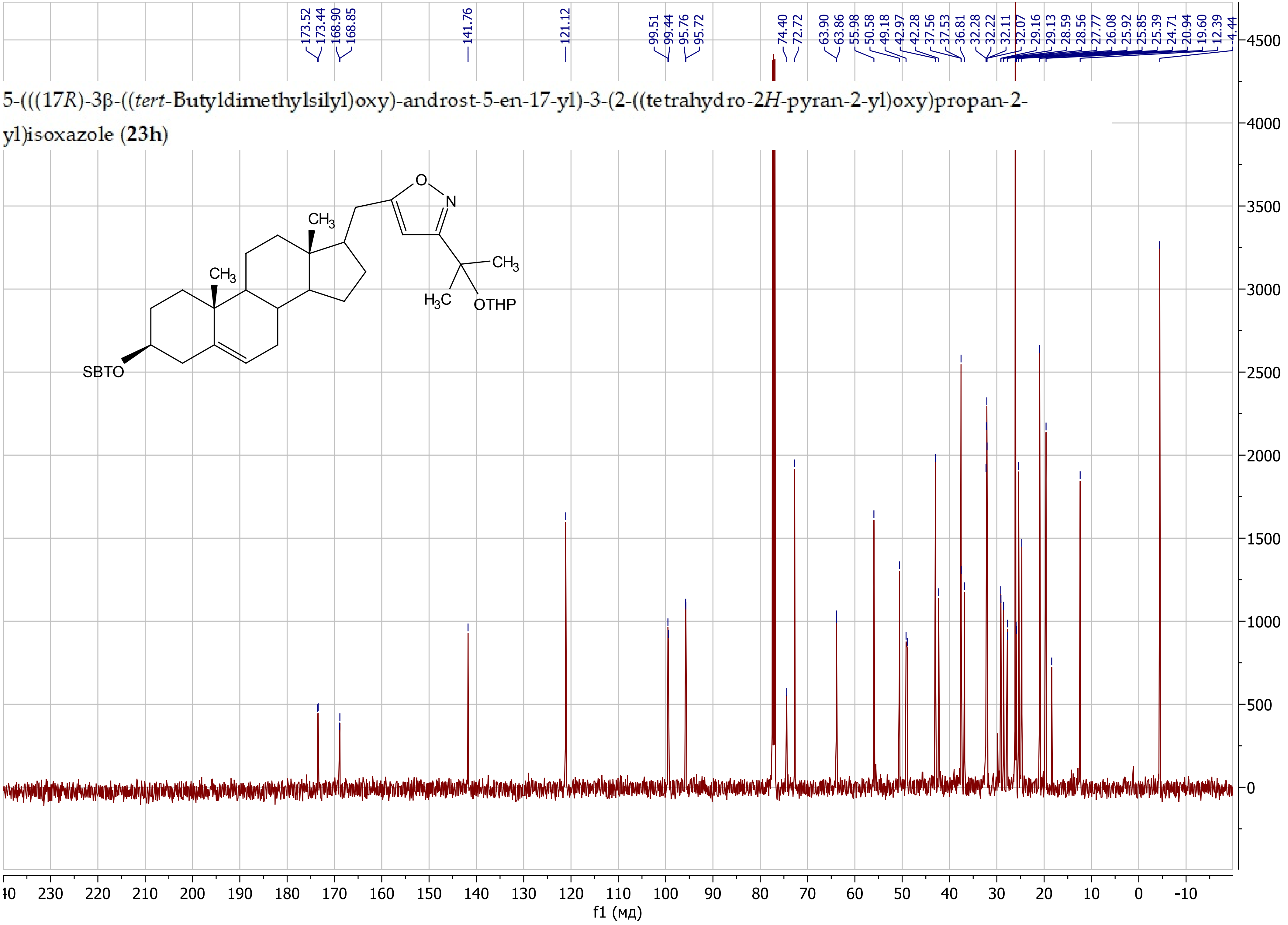

5-(((17*R*)-3β-((*tert*-Butyldimethylsilyl)oxy)-androst-5-en-17-yl)-3-(((tetrahydro-2*H*-pyran-2-yl)oxy)methyl)isoxazole (**23i**)

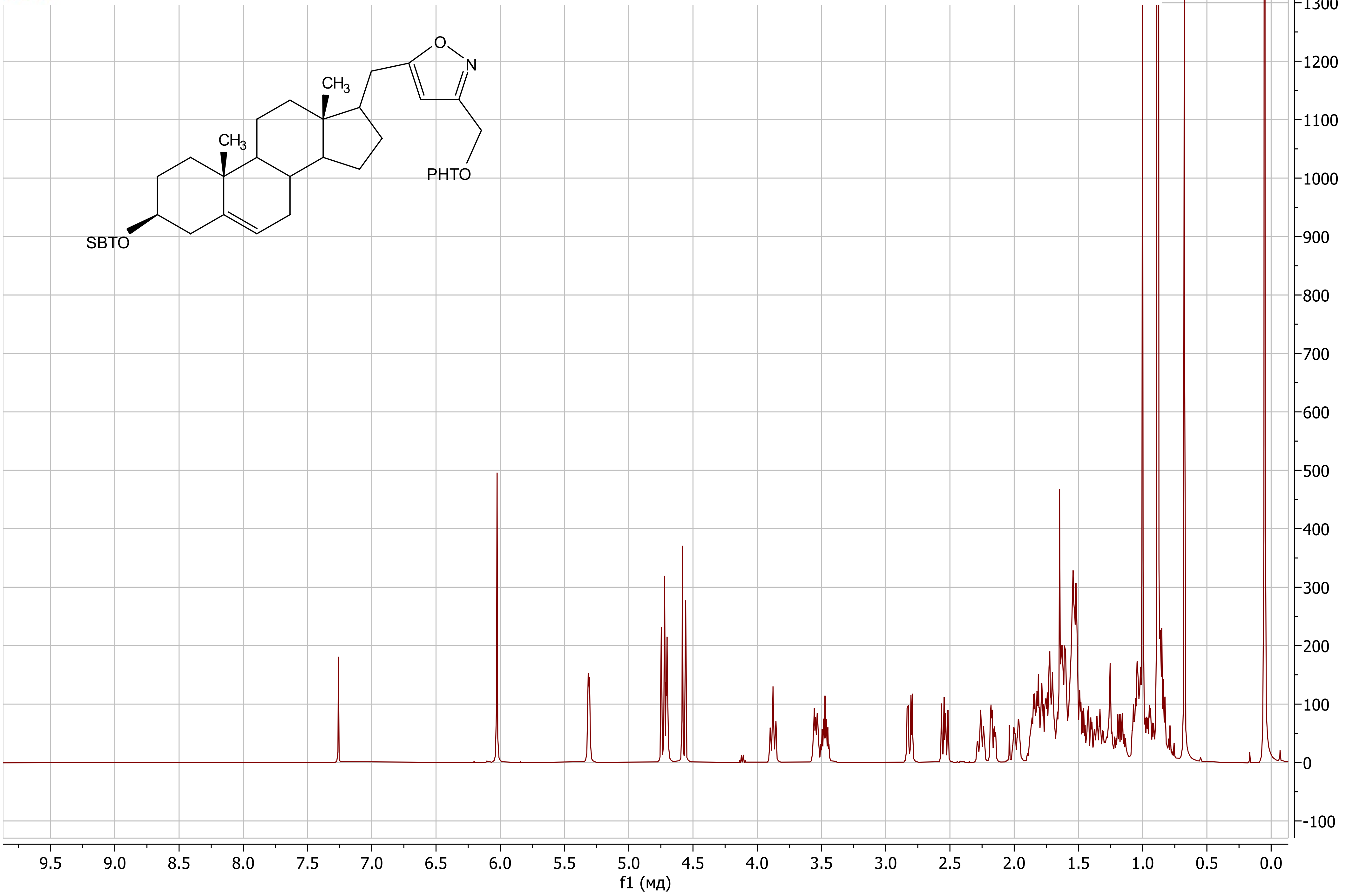

5-(((17*R*)-3β-((*tert*-Butyldimethylsilyl)oxy)-androst-5-en-17-yl)-3-(((tetrahydro-2*H*-pyran-2-yl)oxy)methyl)isoxazole (**23i**)

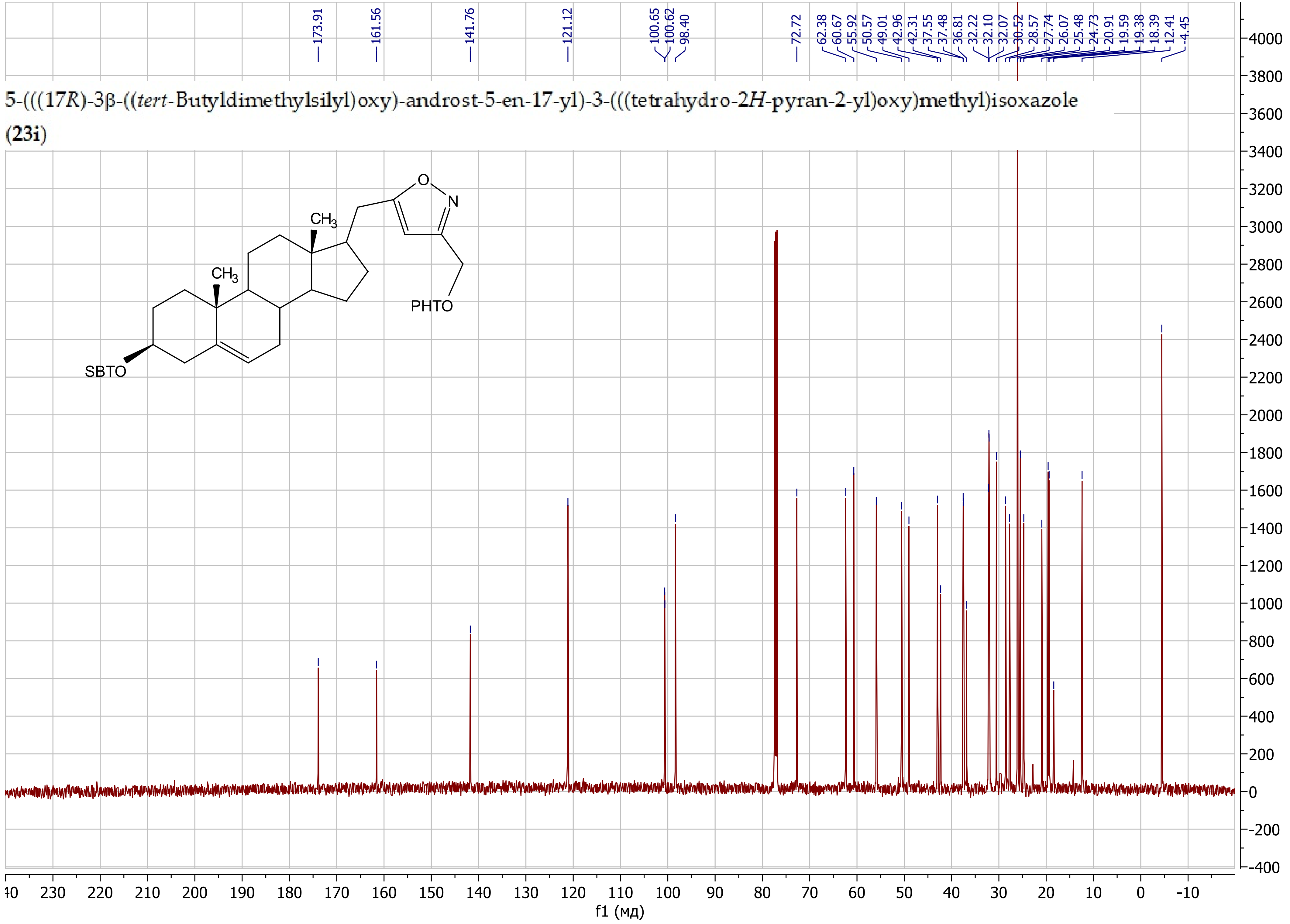

(17*R*)-(Isoxazol-5-ylmethyl)-androst-5-en-3β-ol (**24a**)

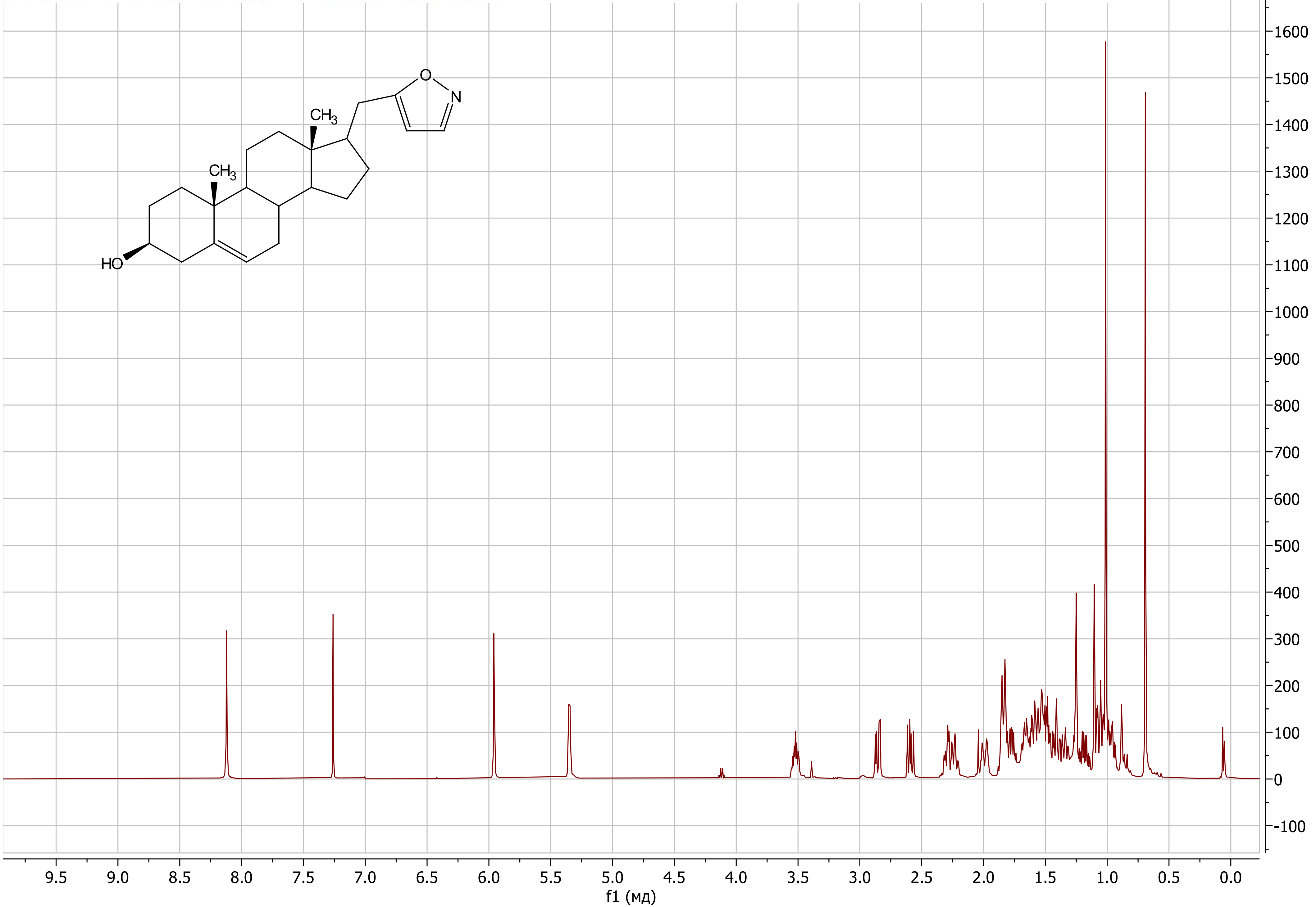

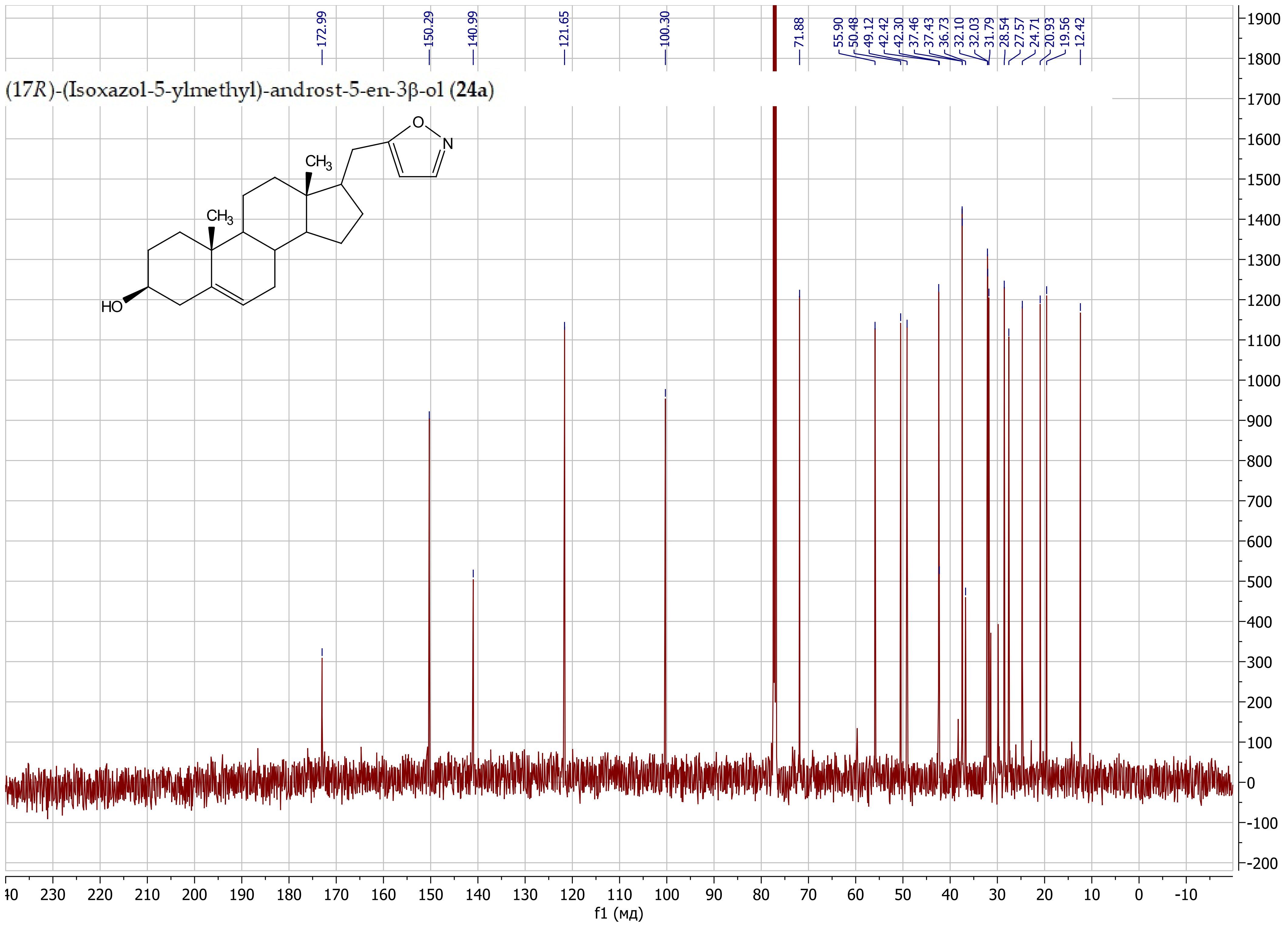
(17R)-(Isoxazol-5-ylmethyl)-androst-5-en-3β-ol (24a)
172.99
150.29
140.99
121.65
100.30
71.88
55.90
50.48
49.12
42.42
42.30
37.46
37.43
36.73
32.10
32.03
31.79
28.54
27.57
24.71
20.93
19.56
12.42
O
N
CH3
CH3
HO
f1 (мд)

(17*R*)-17-((3-isopropylisoxazol-5-yl)methyl)-androst-5-en-3β-ol (**24b**)

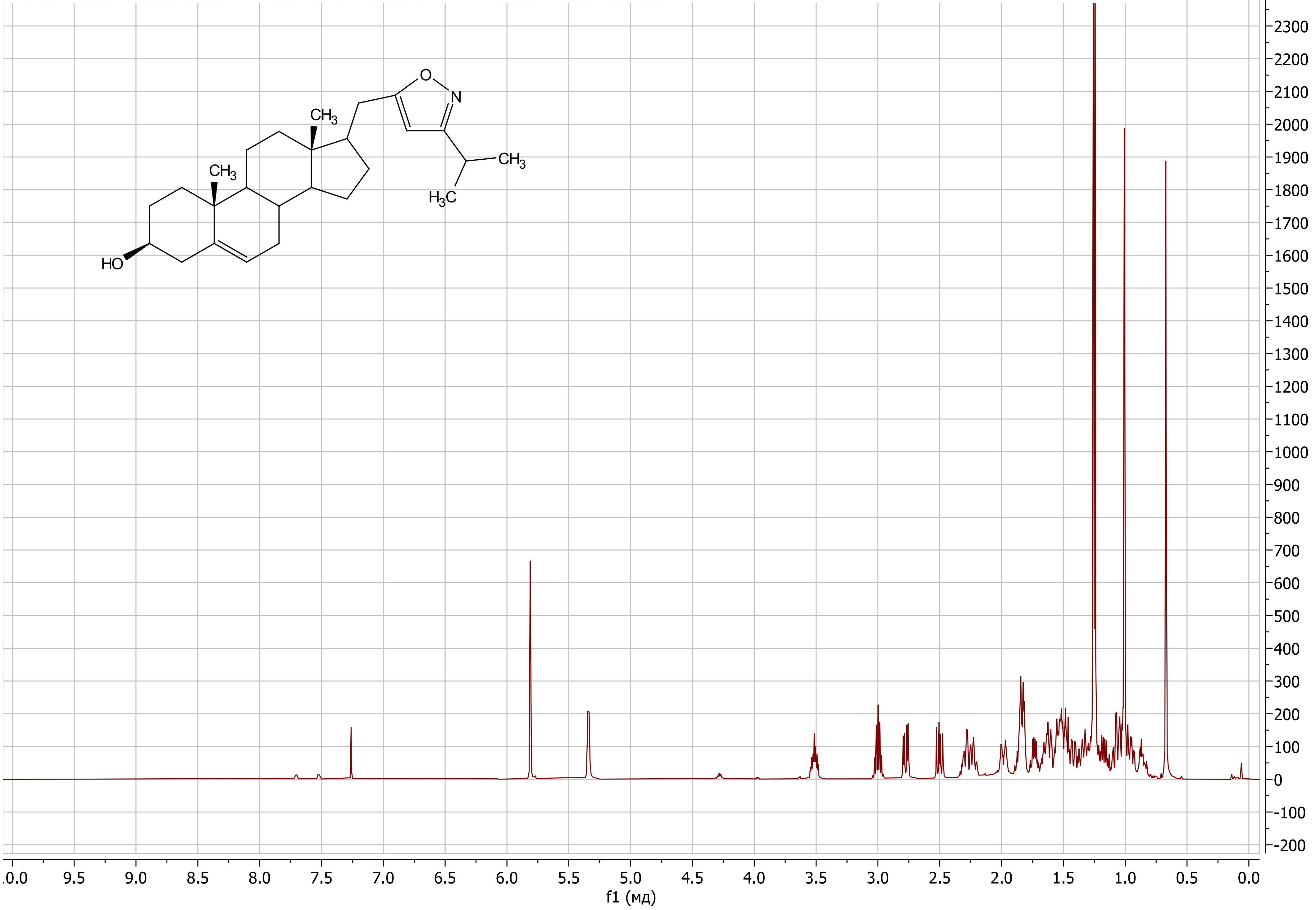

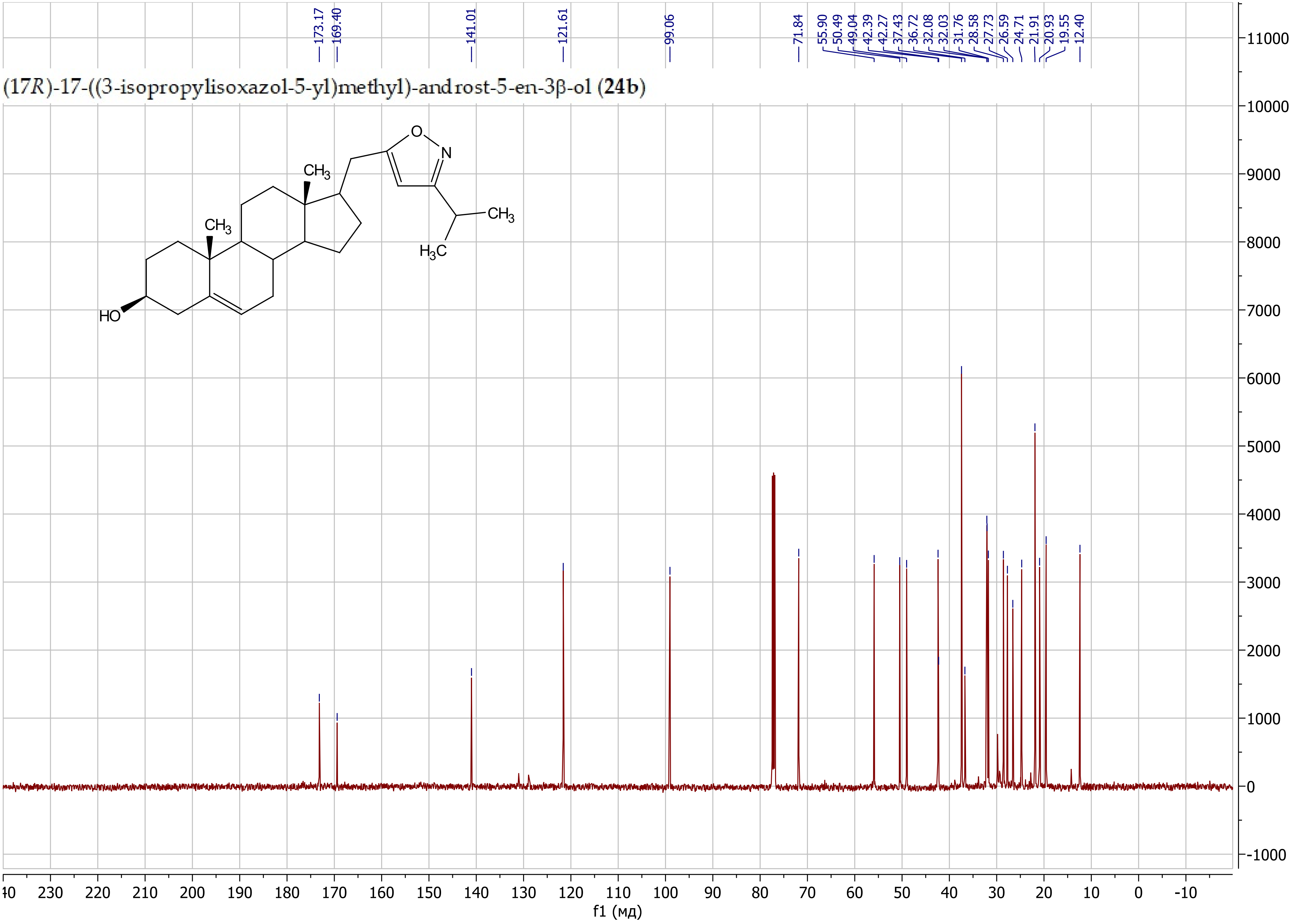
173.17
169.40
141.01
121.61
99.06
71.84
55.90
50.49
49.04
42.39
42.27
37.43
36.72
32.08
32.03
31.76
28.58
27.73
26.59
24.71
21.91
20.93
19.55
12.40
(17*R*)-17-((3-isopropylisoxazol-5-yl)methyl)-androst-5-en-3β-ol (**24b**)
O
N
CH3
CH3
CH3
H3C
HO
f1 (мд)

### 17β-((3-Butylisoxazol-5-yl)methyl)-androst-5-en-3β-ol (**24d**)

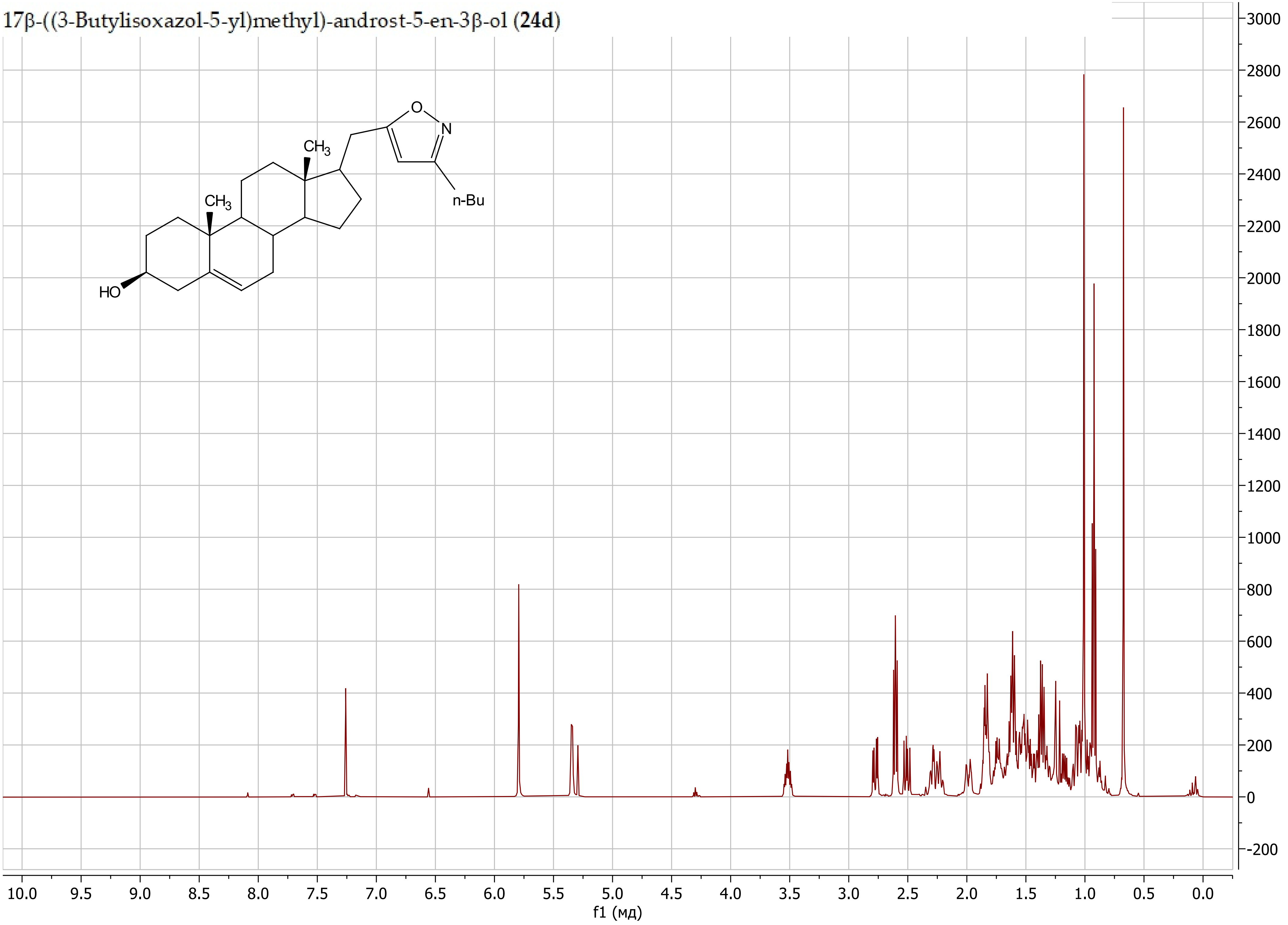

17β-((3-Butylisoxazol-5-yl)methyl)-androst-5-en-3β-ol (**24d**)

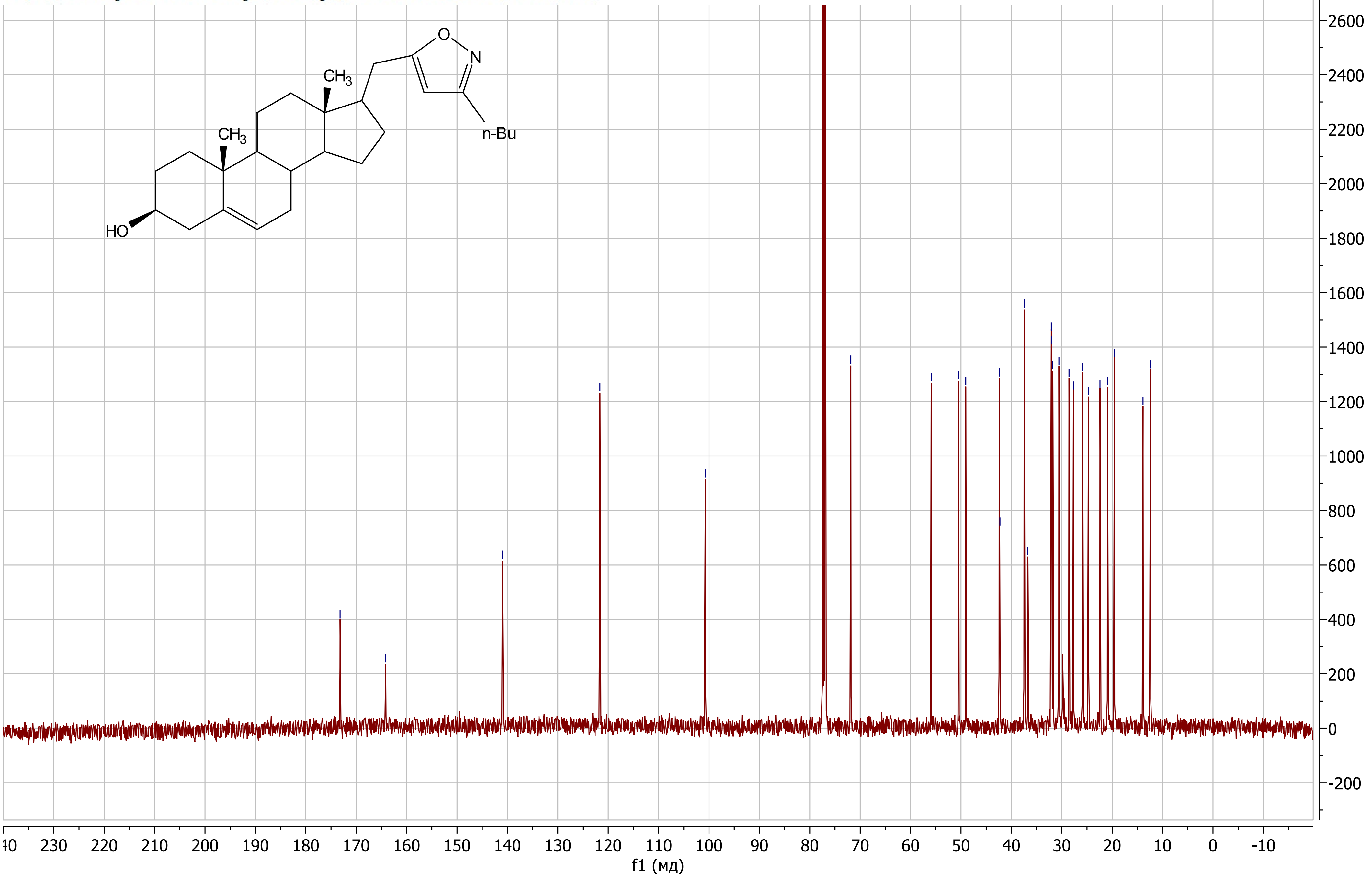

(17*R*)-17-((3-Phenylisoxazol-5-yl)methyl)-androst-5-en-3β-ol (**24e**)

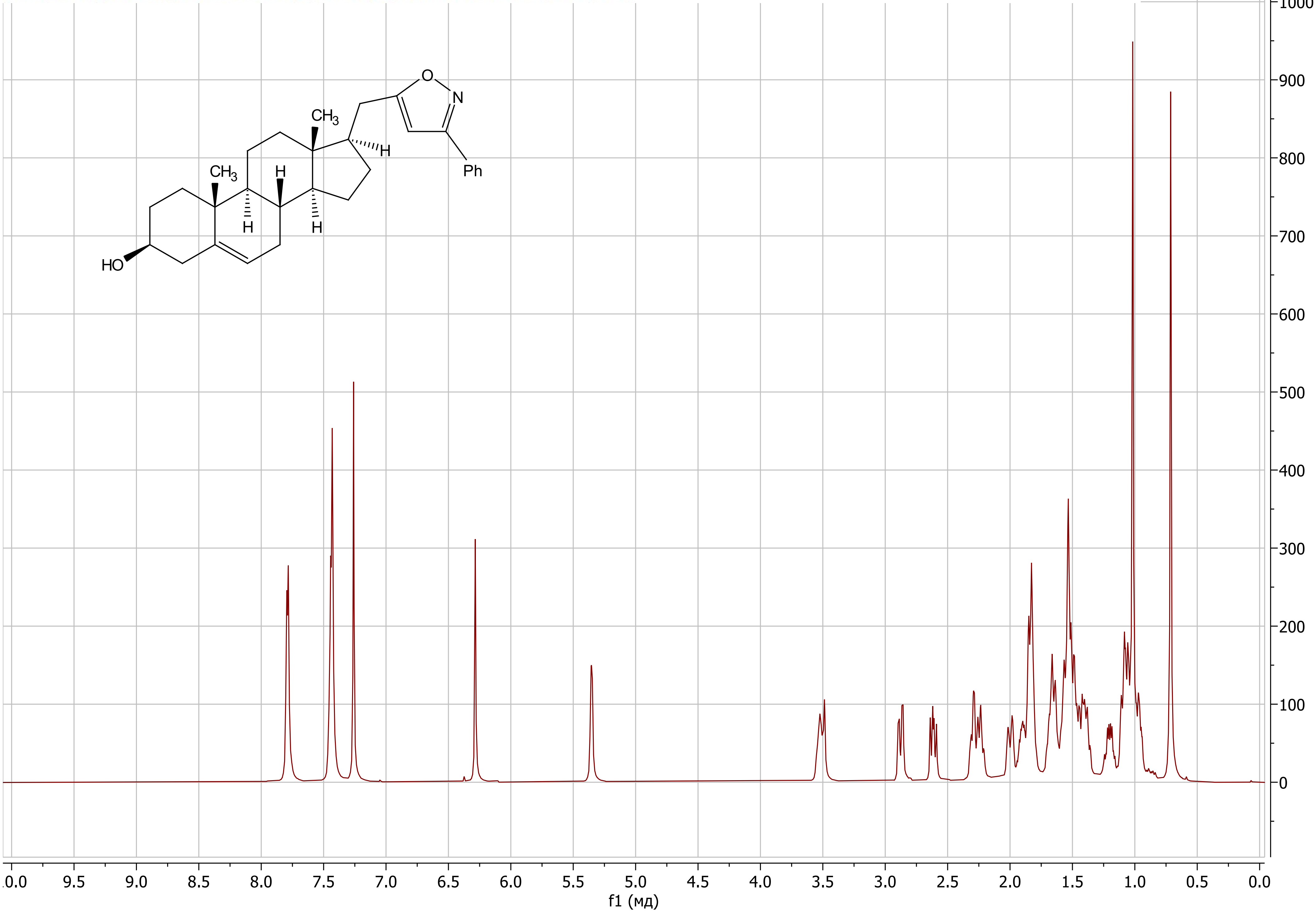

(17*R*)-17-((3-Phenylisoxazol-5-yl)methyl)-androst-5-en-3β-ol (**24e**)

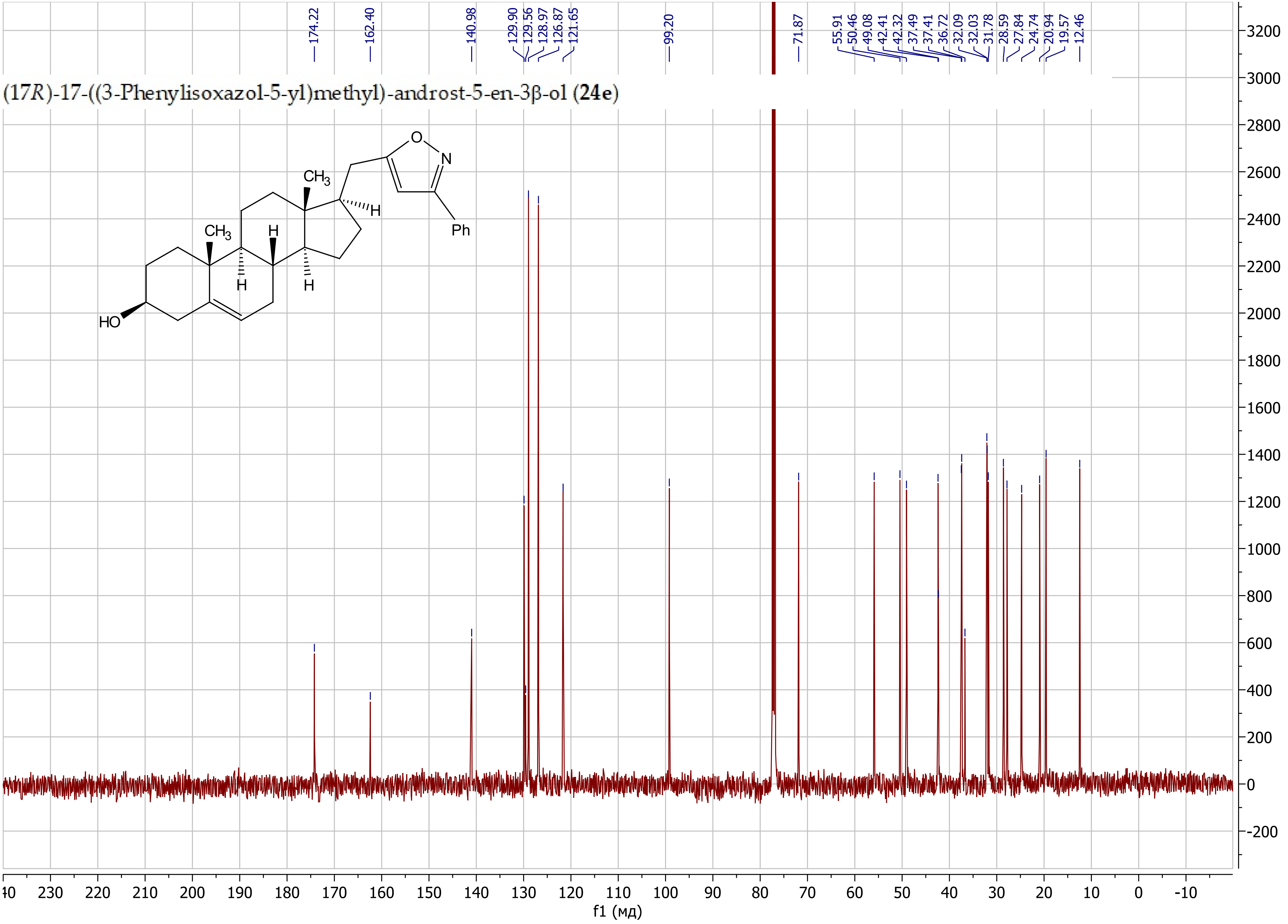

(17*R*)-17-((3-(Pyridin-3-yl)isoxazol-5-yl)methyl)-androst-5-en-3β-ol (**24f**)

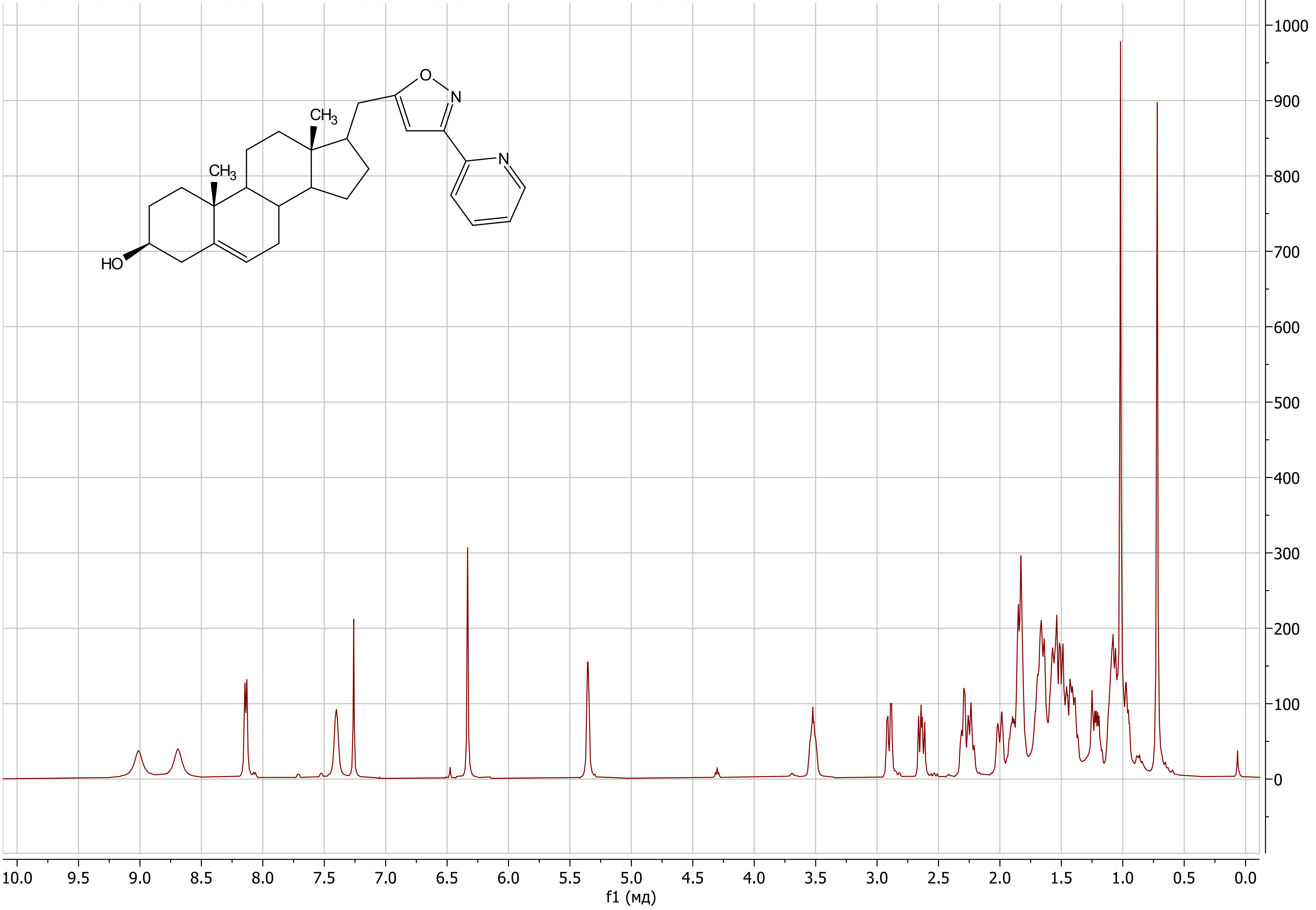

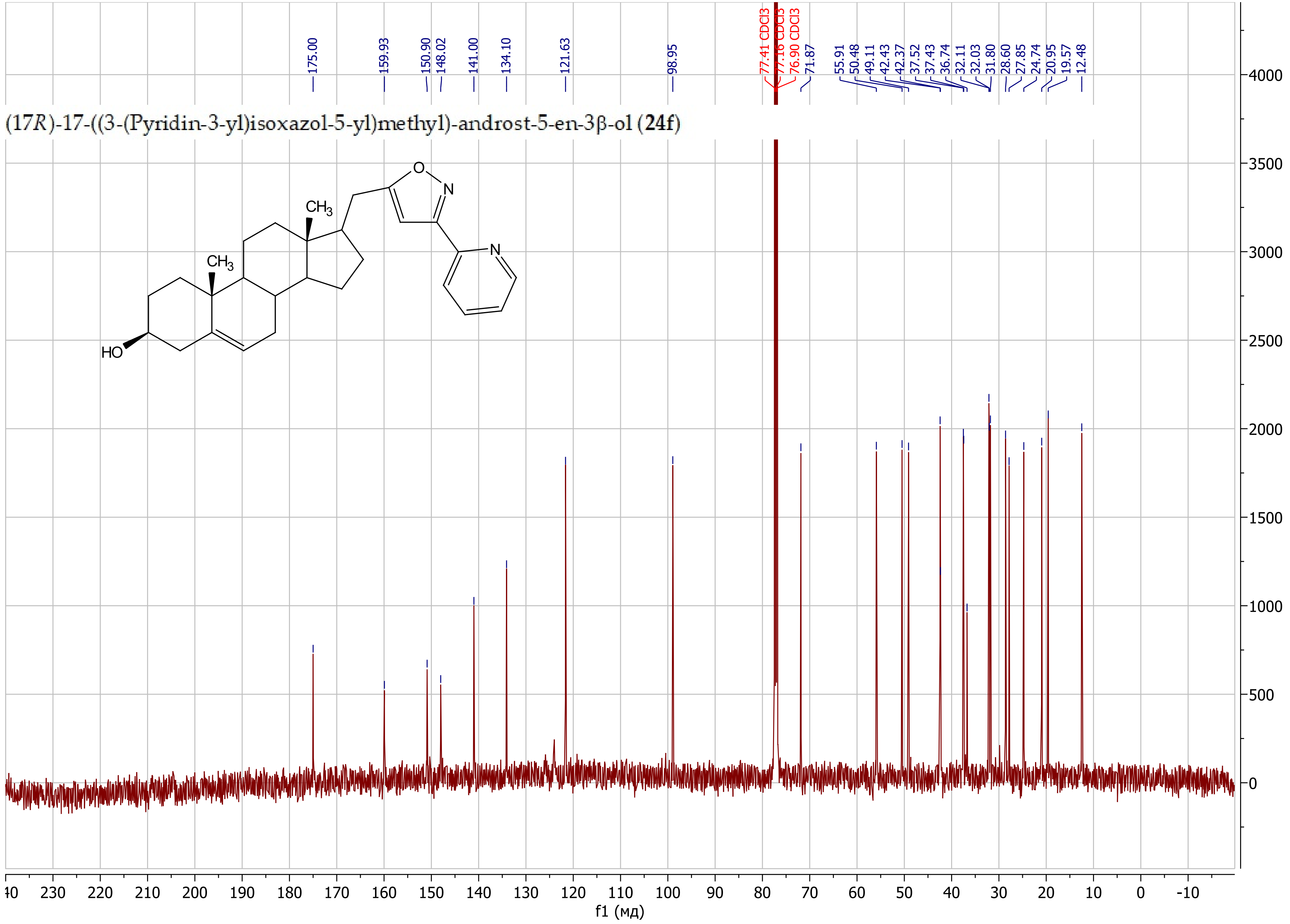


(17*R*)-17-((3-(Pyridin-3-yl)isoxazol-5-yl)methyl)-androst-5-en-3β-ol (**24f**)

17β-((3-(2-Fluorophenyl)isoxazol-5-yl)methyl)-androst-5-en-3β-ol (**24g**)

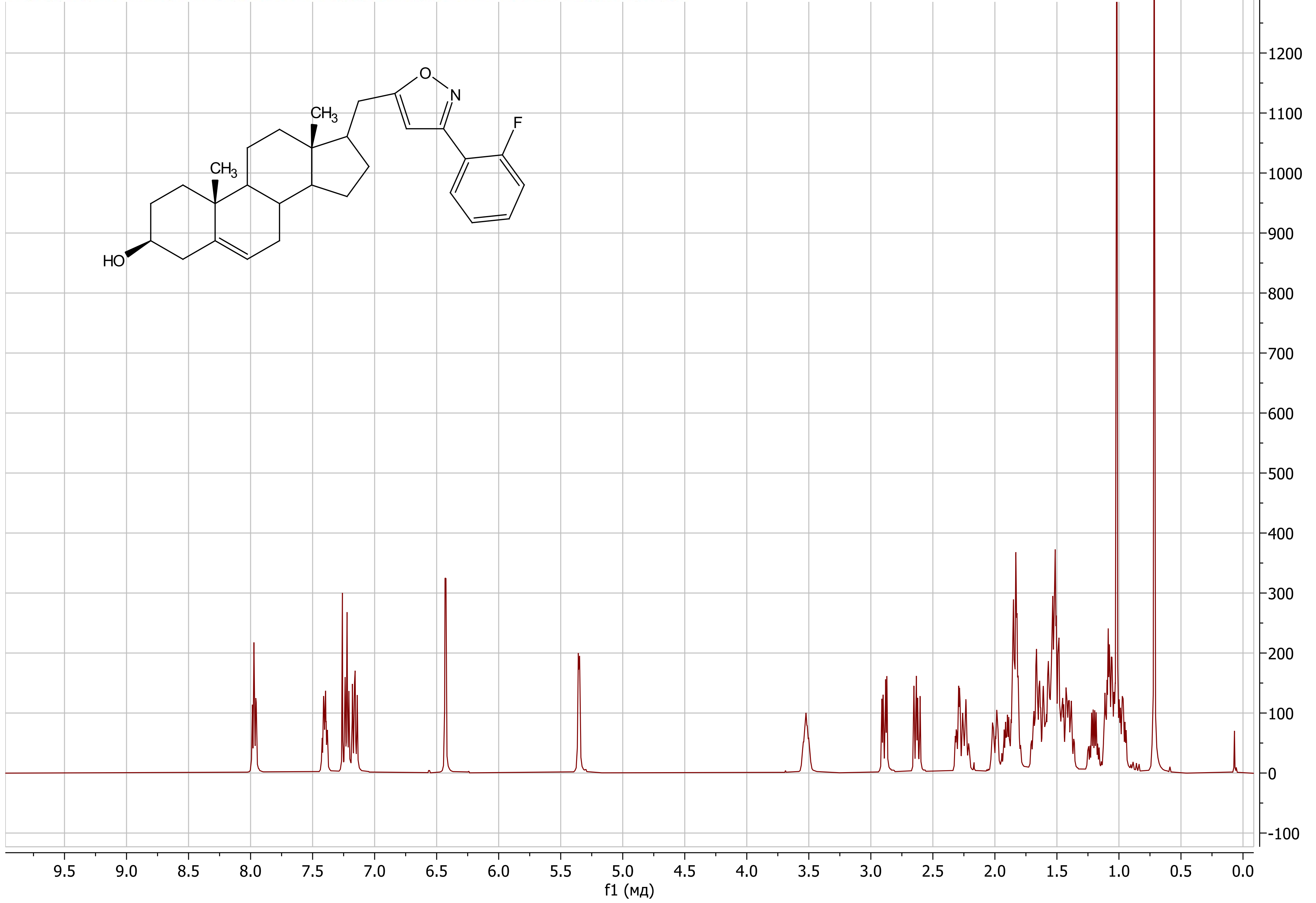

17β-((3-(2-Fluorophenyl)isoxazol-5-yl)methyl)-androst-5-en-3β-ol (**24g**)

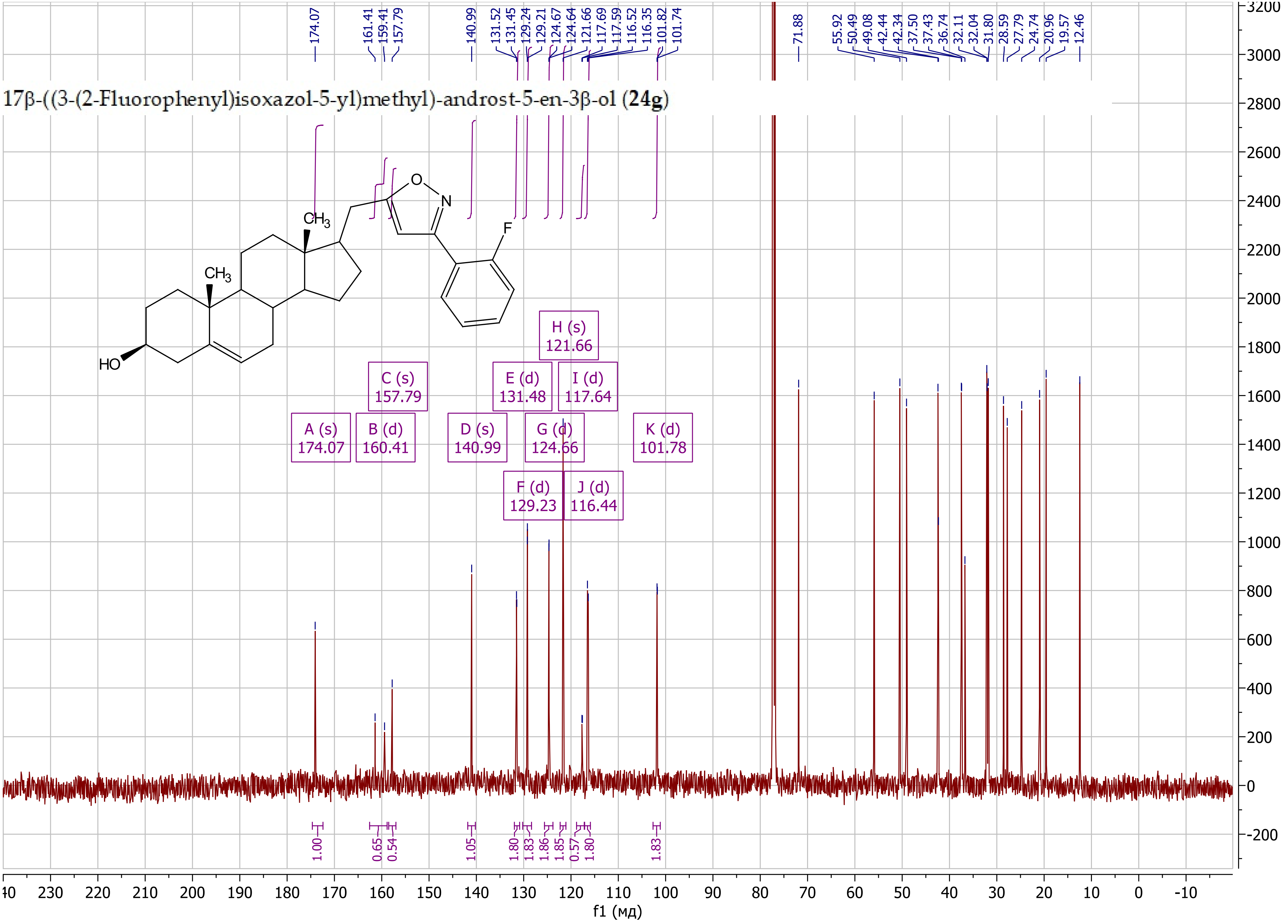

(17*R*)-17-((3-(2-Hydroxypropan-2-yl)isoxazol-5-yl)methyl)-androst-5-en-3β-ol (**24j**)

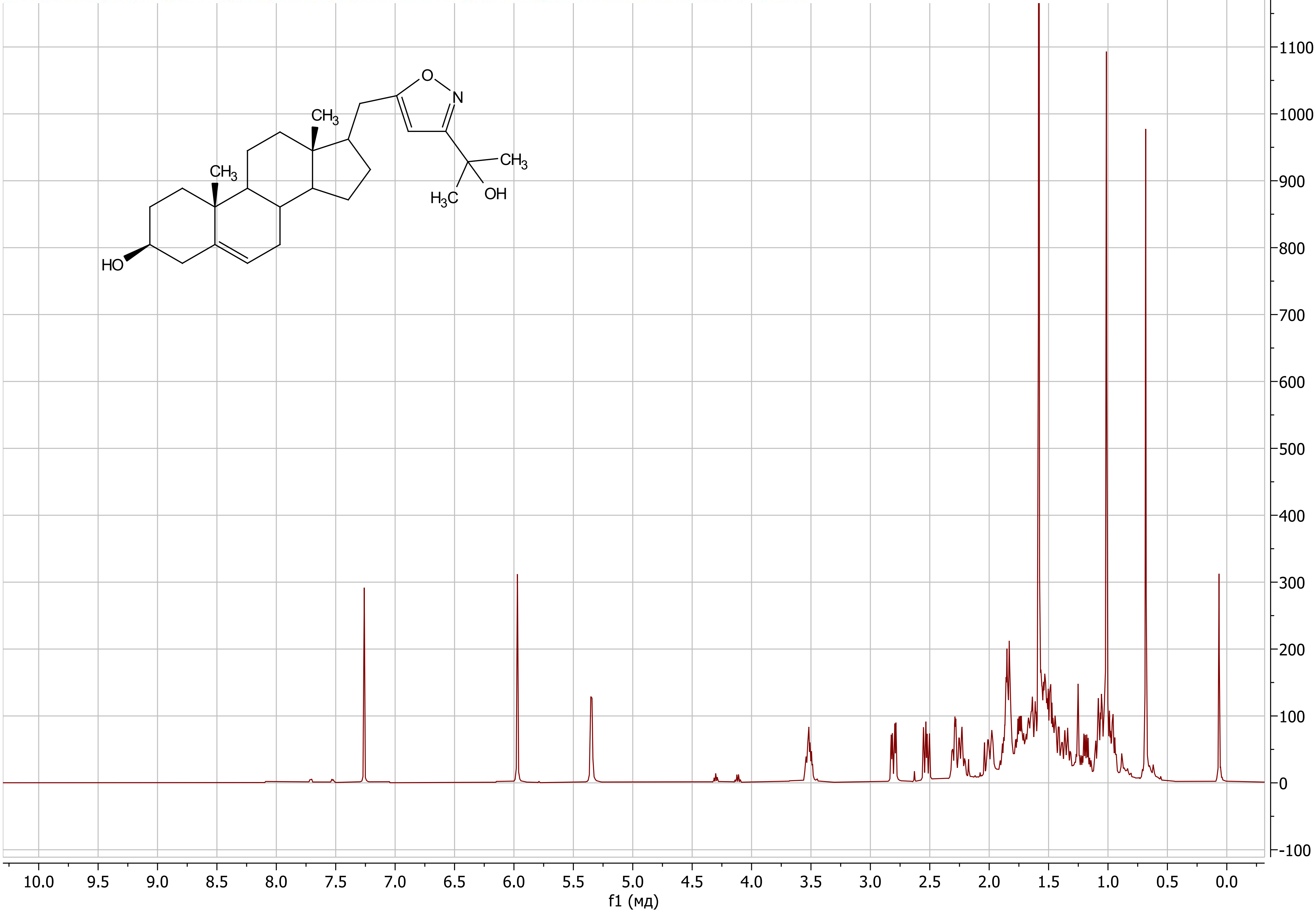

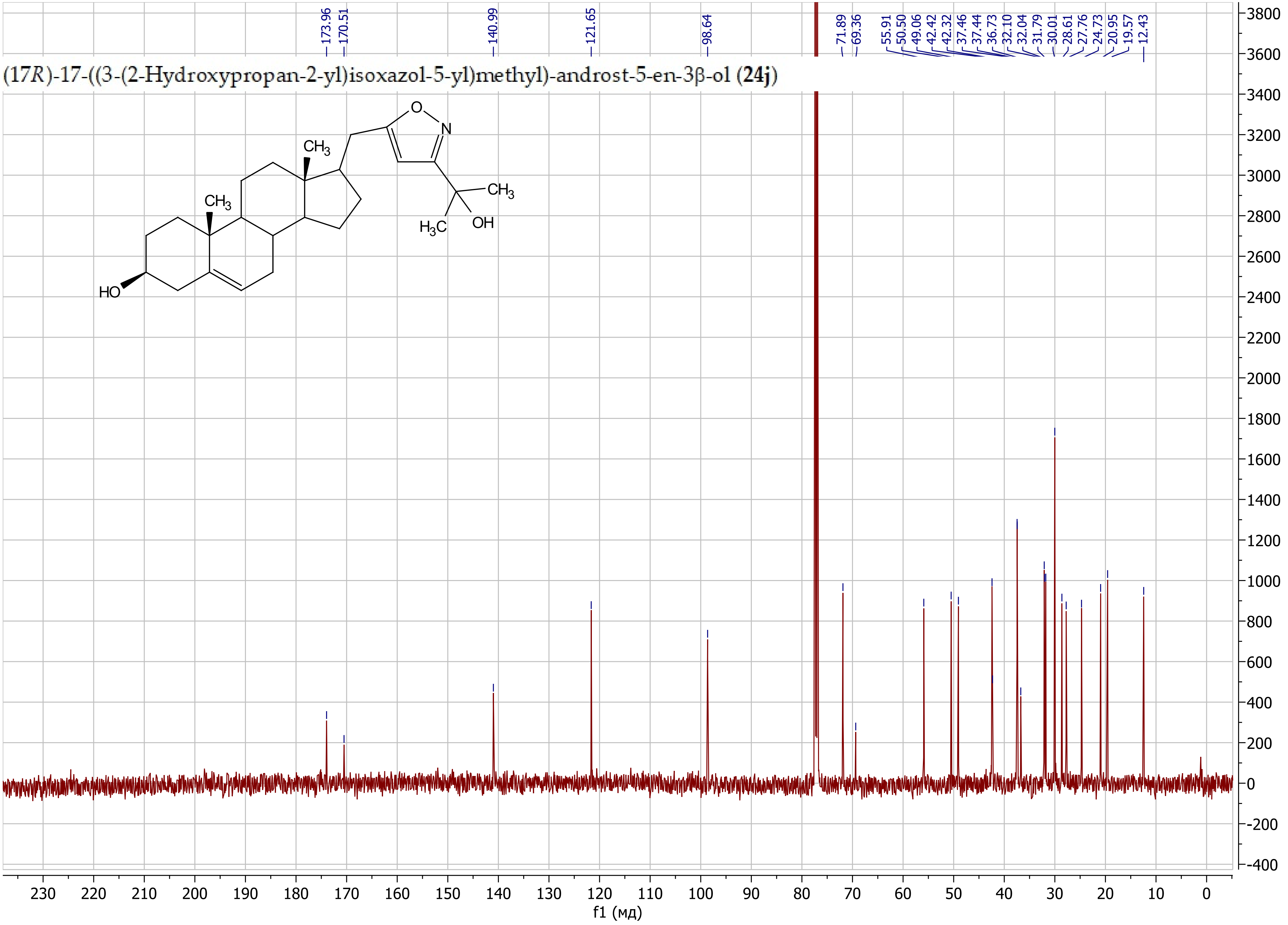

(17R)-17-((3-(2-Hydroxypropan-2-yl)isoxazol-5-yl)methyl)-androst-5-en-3β-ol (24j)
173.96
170.51
140.99
121.65
98.64
71.89
69.36
55.91
50.50
49.06
42.42
42.32
37.46
37.44
36.73
32.10
32.04
31.79
30.01
28.61
27.76
24.73
20.95
19.57
12.43
HO
CH3
CH3
O
N
CH3
H3C
OH
f1 (мд)

(17*R*)-17-((3-Cyclopropylisoxazol-5-yl)methyl)-androst-5-en-3β-ol (**24c**)

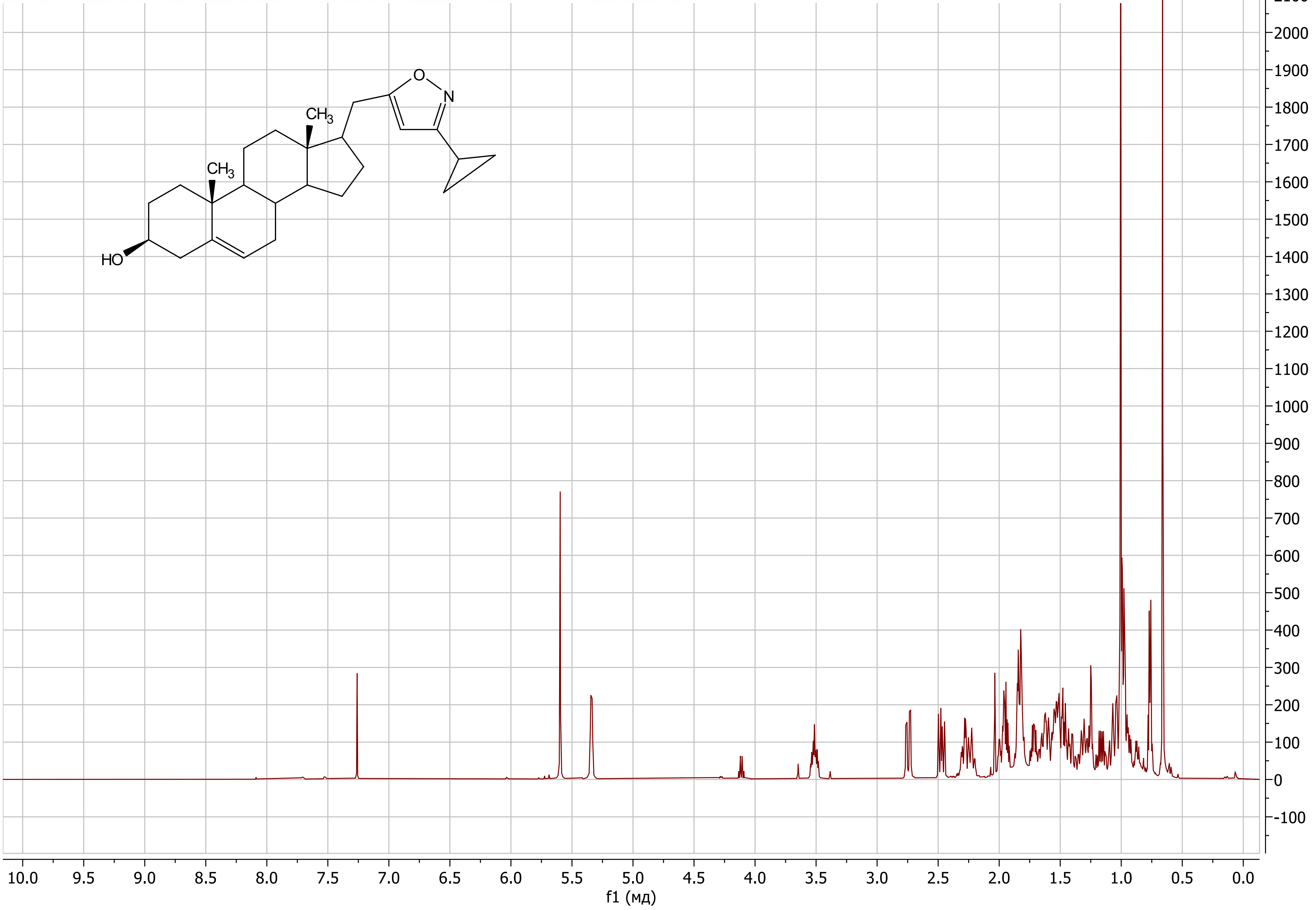

(17*R*)-17-((3-Cyclopropylisoxazol-5-yl)methyl)-androst-5-en-3β-ol (**24c**)

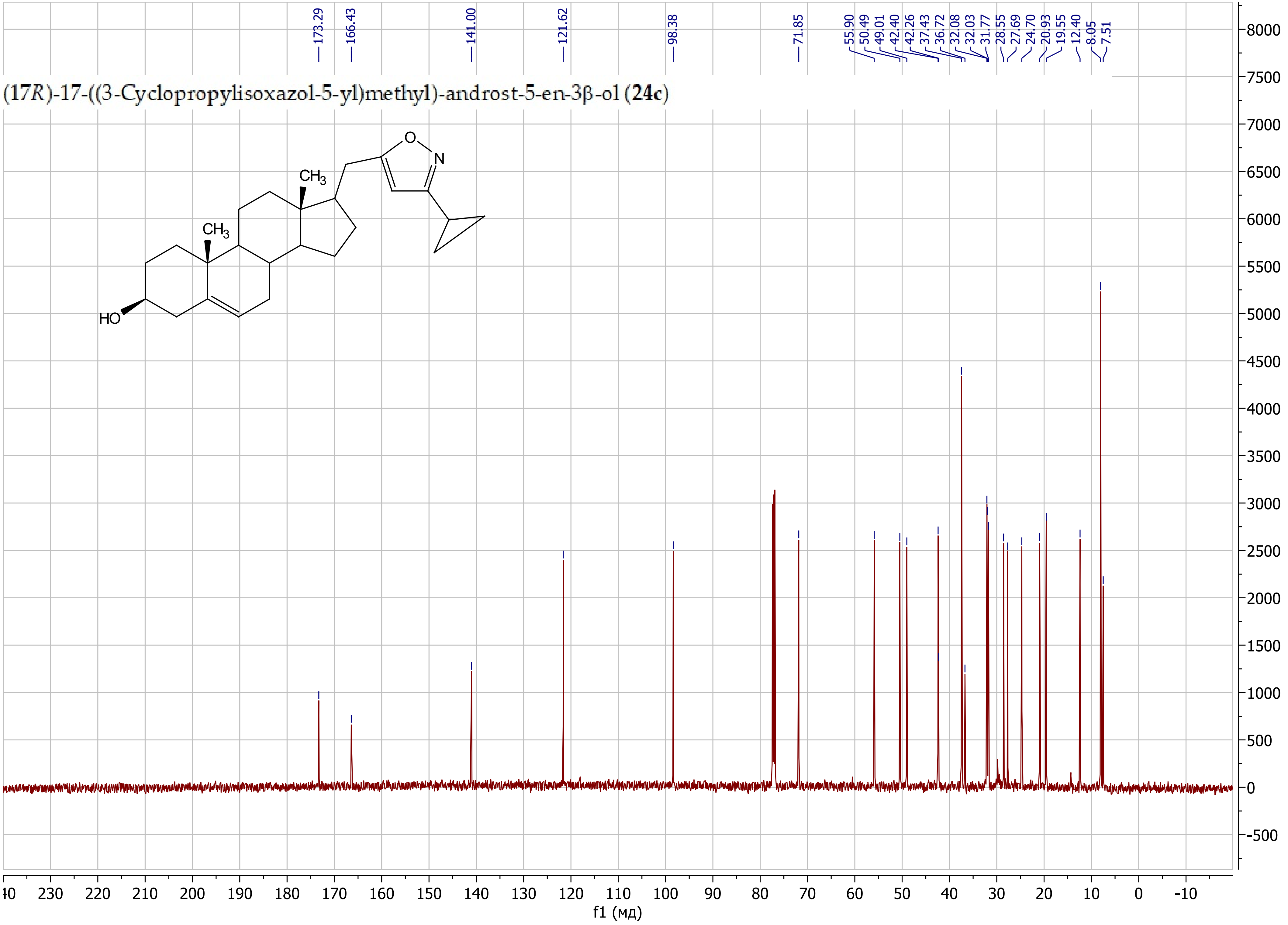

(*E*)-4-Amino-1-((17*R*)-3β-((*tert*-butyldimethylsilyl)oxy)-androst-5-en-17-yl)-5-methyl-2-oxohex-3-en-3-yl 1*H*-imidazole-1-carboxylate (**26b**)

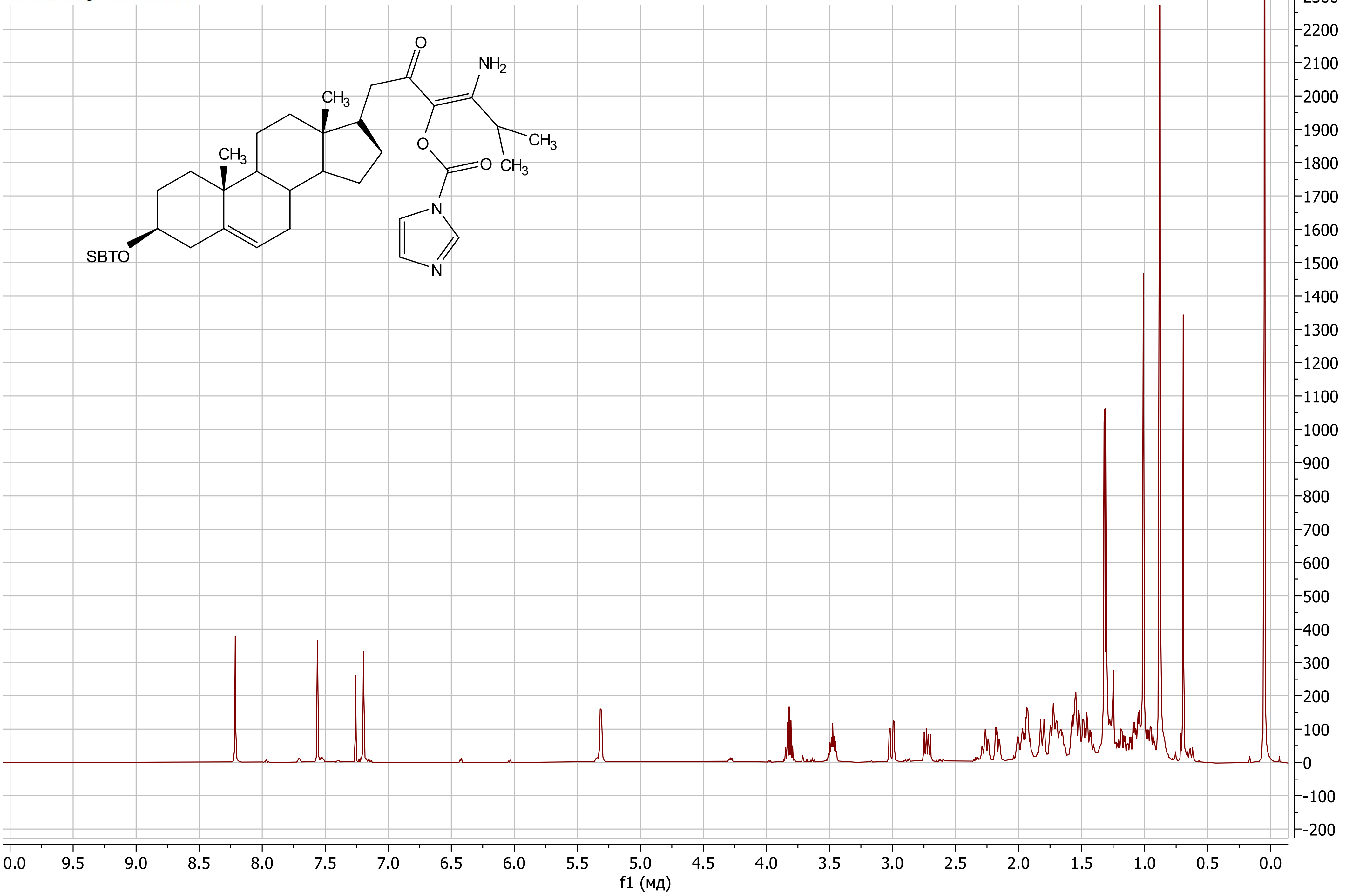

(*E*)-4-Amino-1-((17*R*)-3β-((*tert*-butyldimethylsilyl)oxy)-androst-5-en-17-yl)-5-methyl-2-oxohex-3-en-3-yl 1*H*-imidazole-1-carboxylate (**26b**)

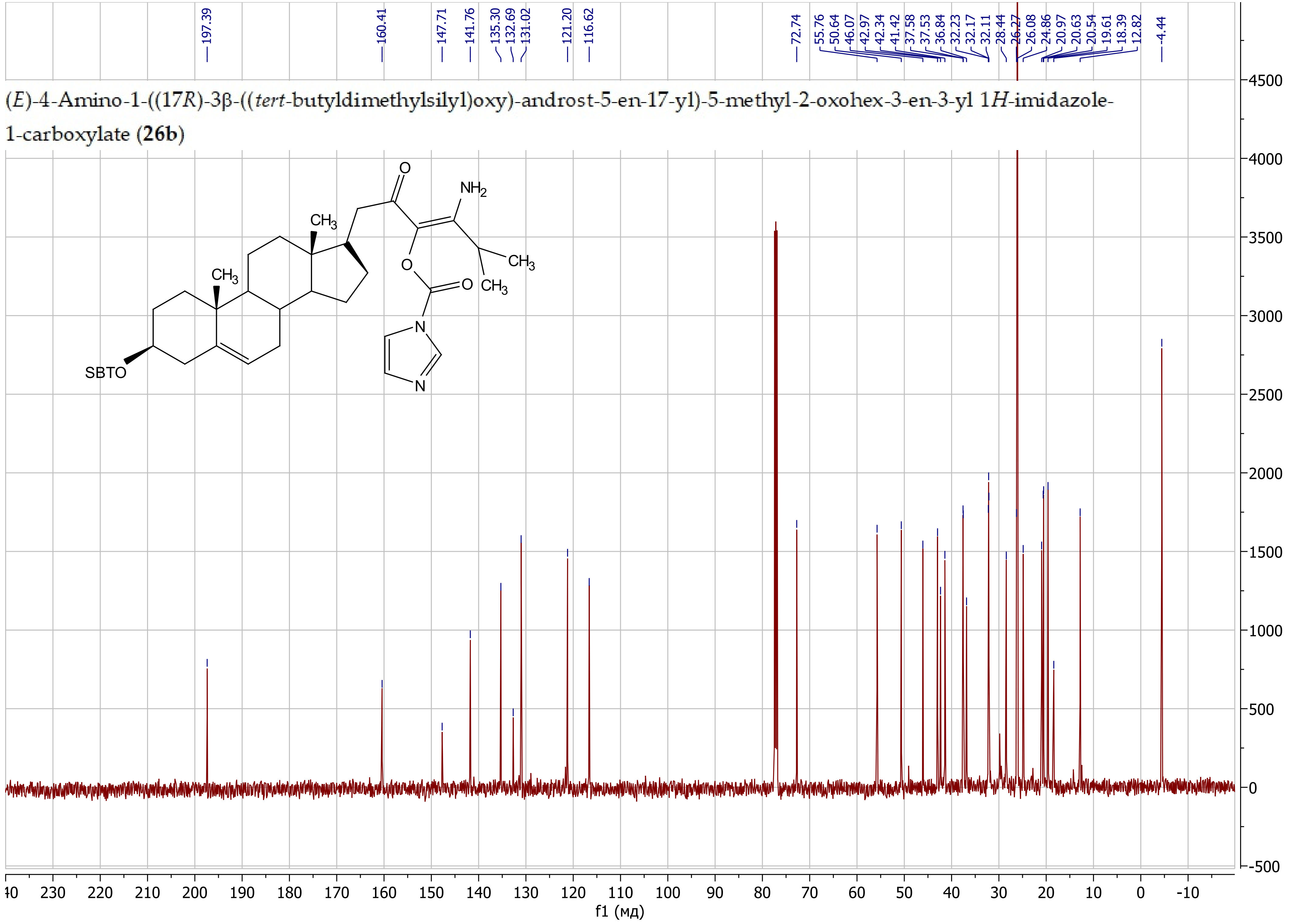

(*E*)-1-Amino-4-((17*R*)-3β-((*tert*-butyldimethylsilyl)oxy)-androst-5-en-17-yl)-1-cyclopropyl-3-oxobut-1-en-2-yl 1*H*-imidazole-1-carboxylate (**26c**)

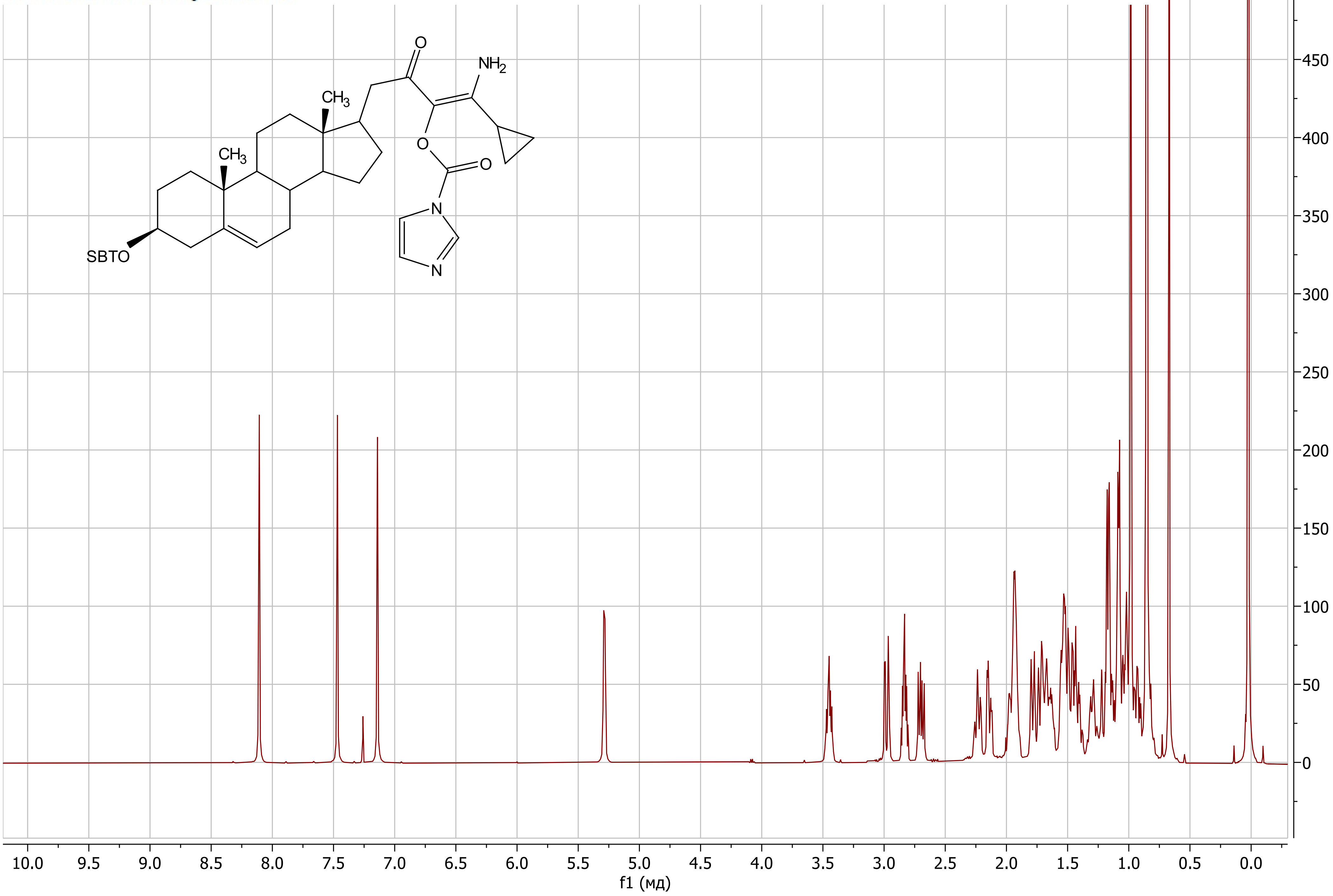

(*E*)-1-Amino-4-((17*R*)-3β-((*tert*-butyldimethylsilyl)oxy)-androst-5-en-17-yl)-1-cyclopropyl-3-oxobut-1-en-2-yl 1*H*-imidazole-1-carboxylate (**26c**)

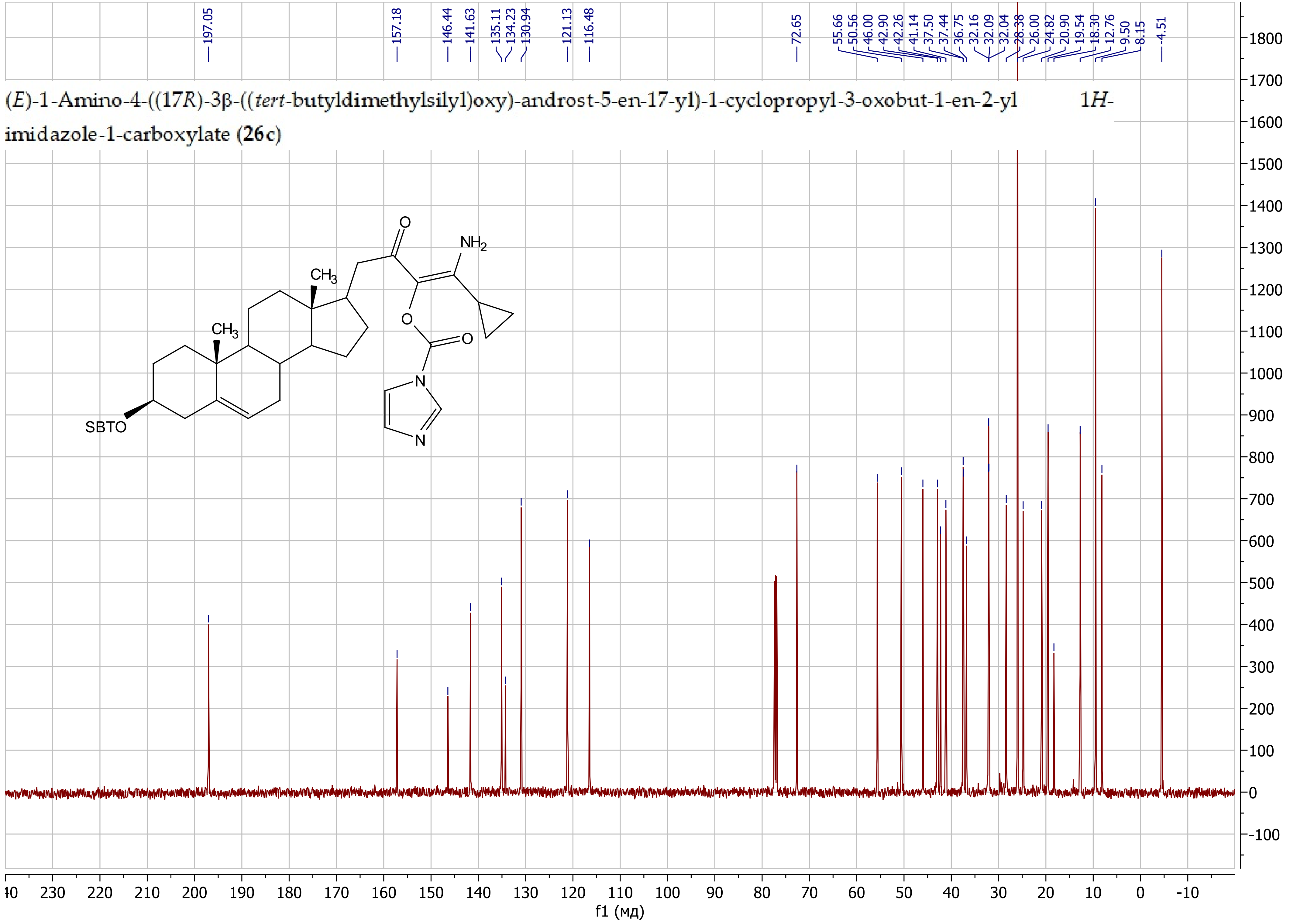

(*E*)-1-Amino-4-((17*R*)-3β-hydroxy-androst-5-en-17-yl)-1-cyclopropyl-3-oxobut-1-en-2-yl 1H-imidazole-1-carboxylate

(**27**)

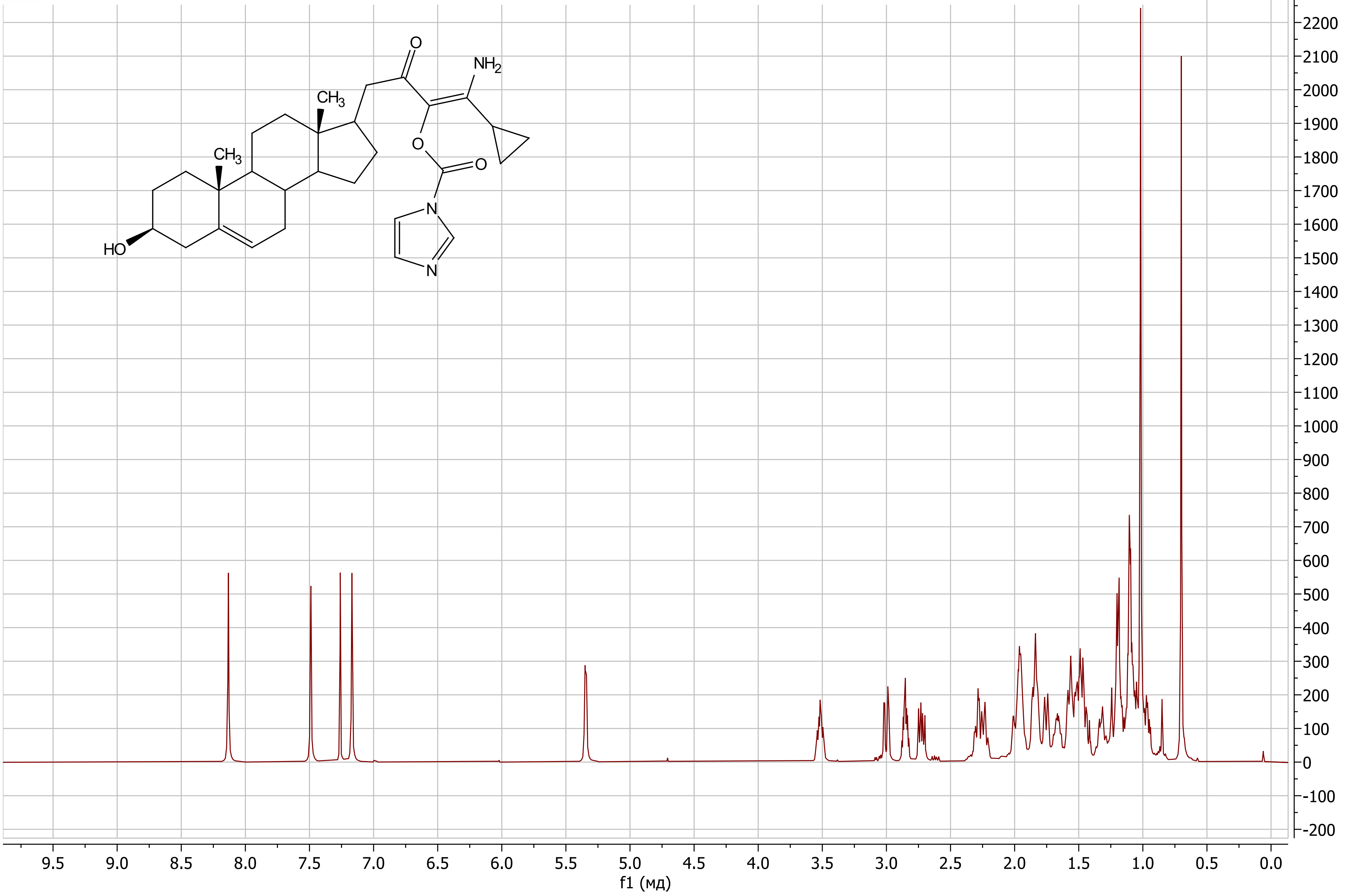

(*E*)-1-Amino-4-((17*R*)-3β-hydroxy-androst-5-en-17-yl)-1-cyclopropyl-3-oxobut-1-en-2-yl 1H-imidazole-1-carboxylate (**27**)

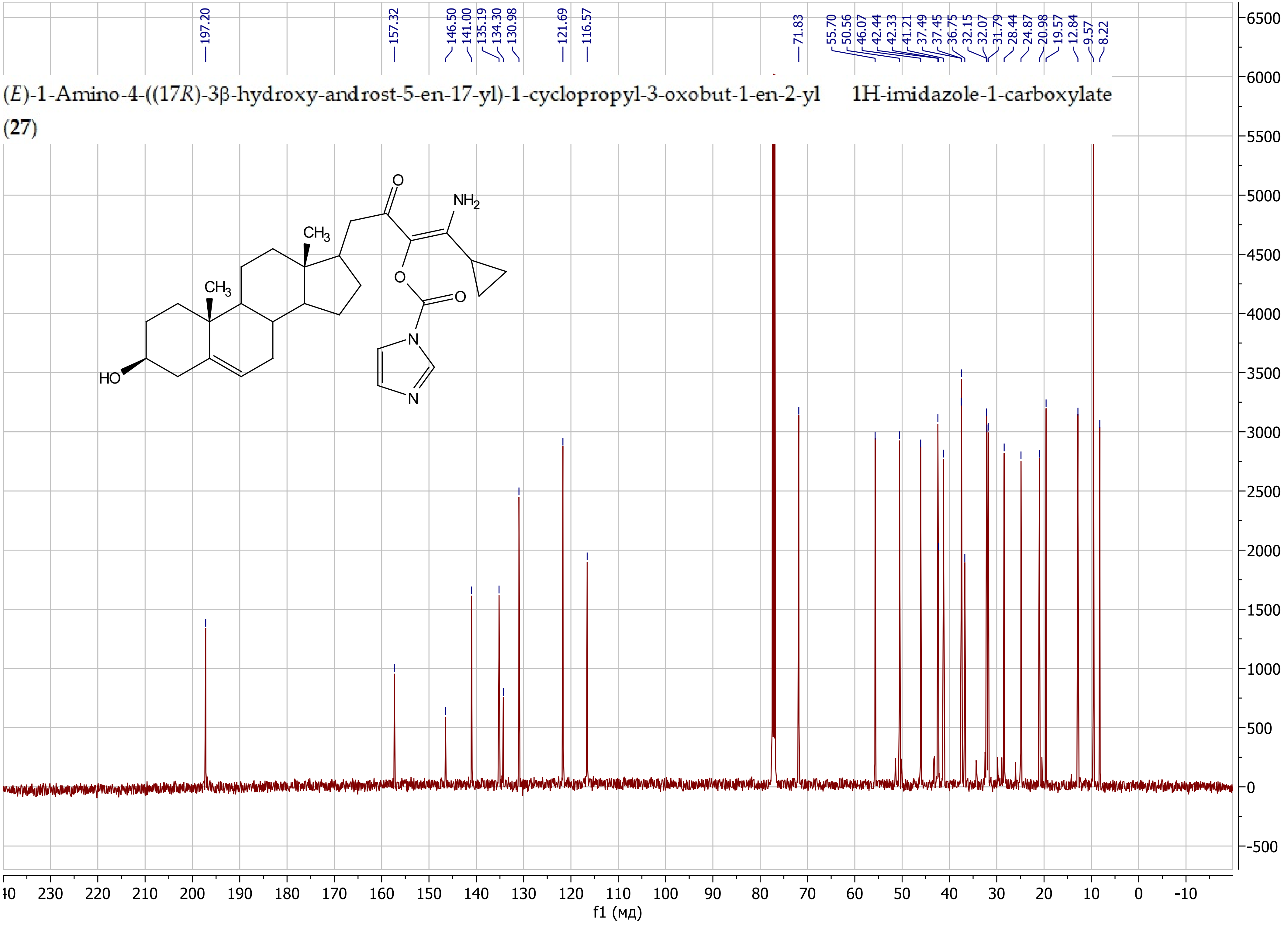

5-(2-((17*R*)-3β-((*tert*-Butyldimethylsilyl)oxy)-androst-5-en-17-yl)acetyl)-4-cyclopropyloxazol-2(3*H*)-one (**28**)

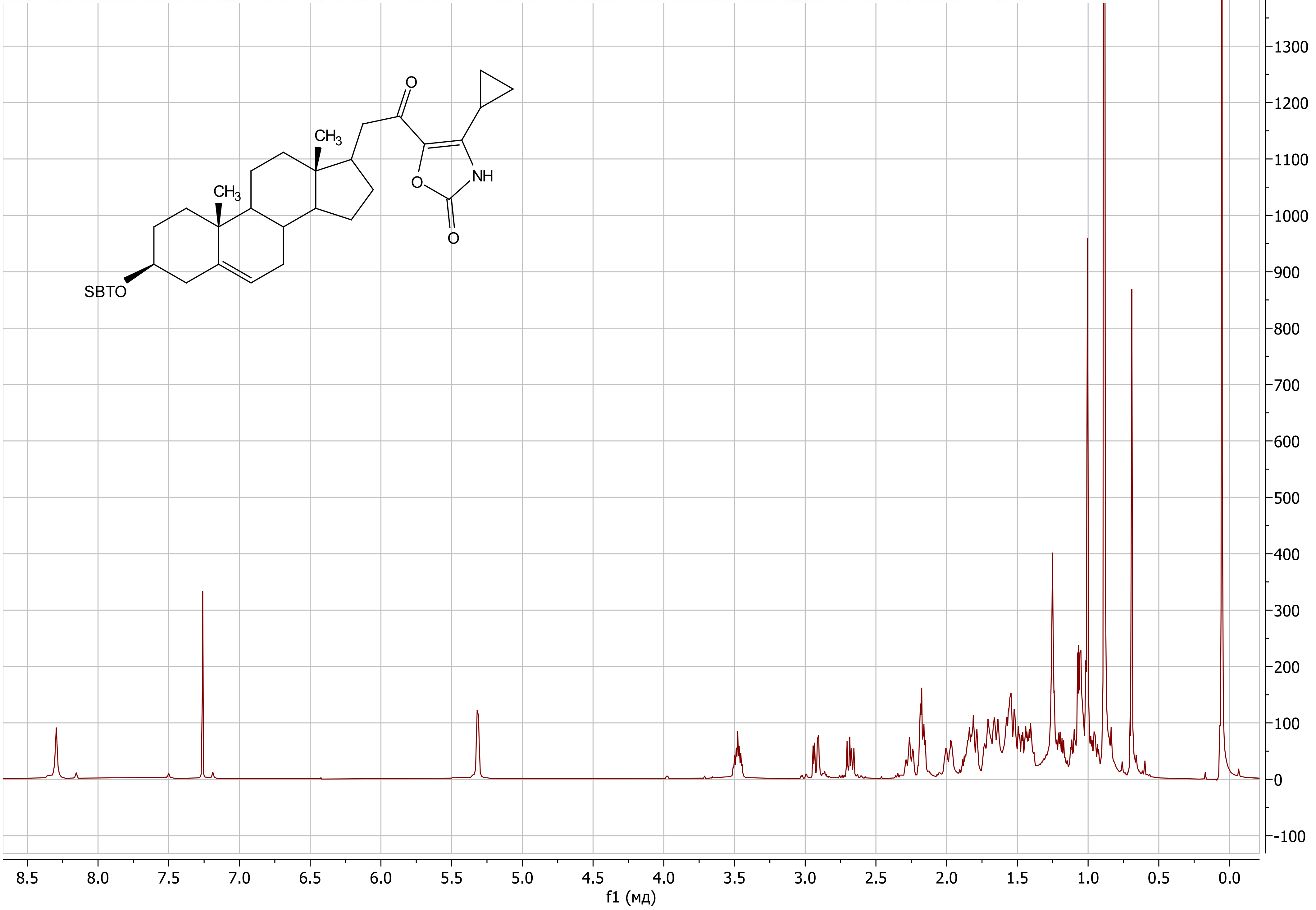

5-(2-((17*R*)-3β-((*tert*-Butyldimethylsilyl)oxy)-androst-5-en-17-yl)acetyl)-4-cyclopropyloxazol-2(3*H*)-one (**28**)

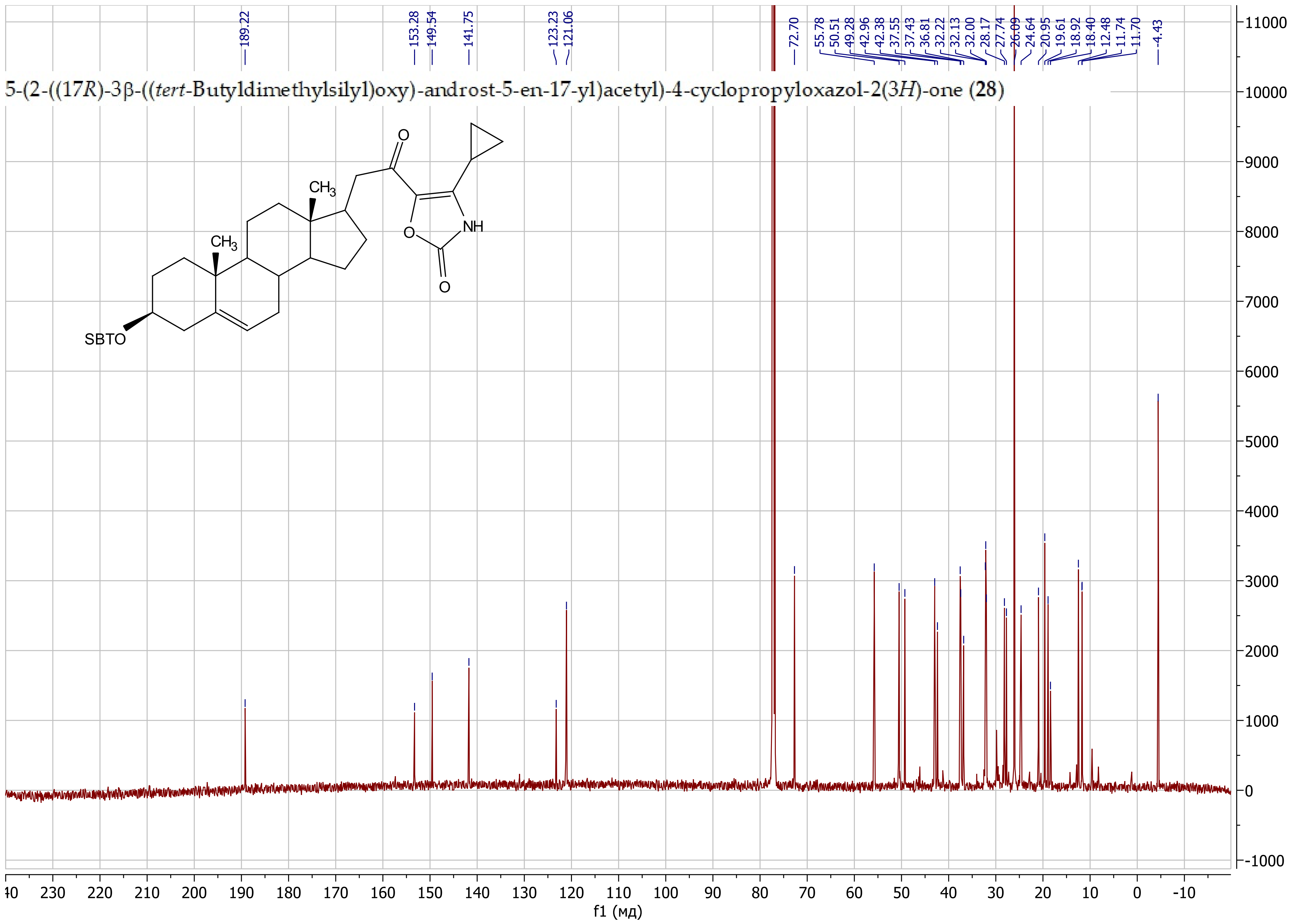

4-(3β-((*tert*-Butyldimethylsilyl)oxy)-androst-5-en-17-yl)-3-oxobutanenitrile (**31**)

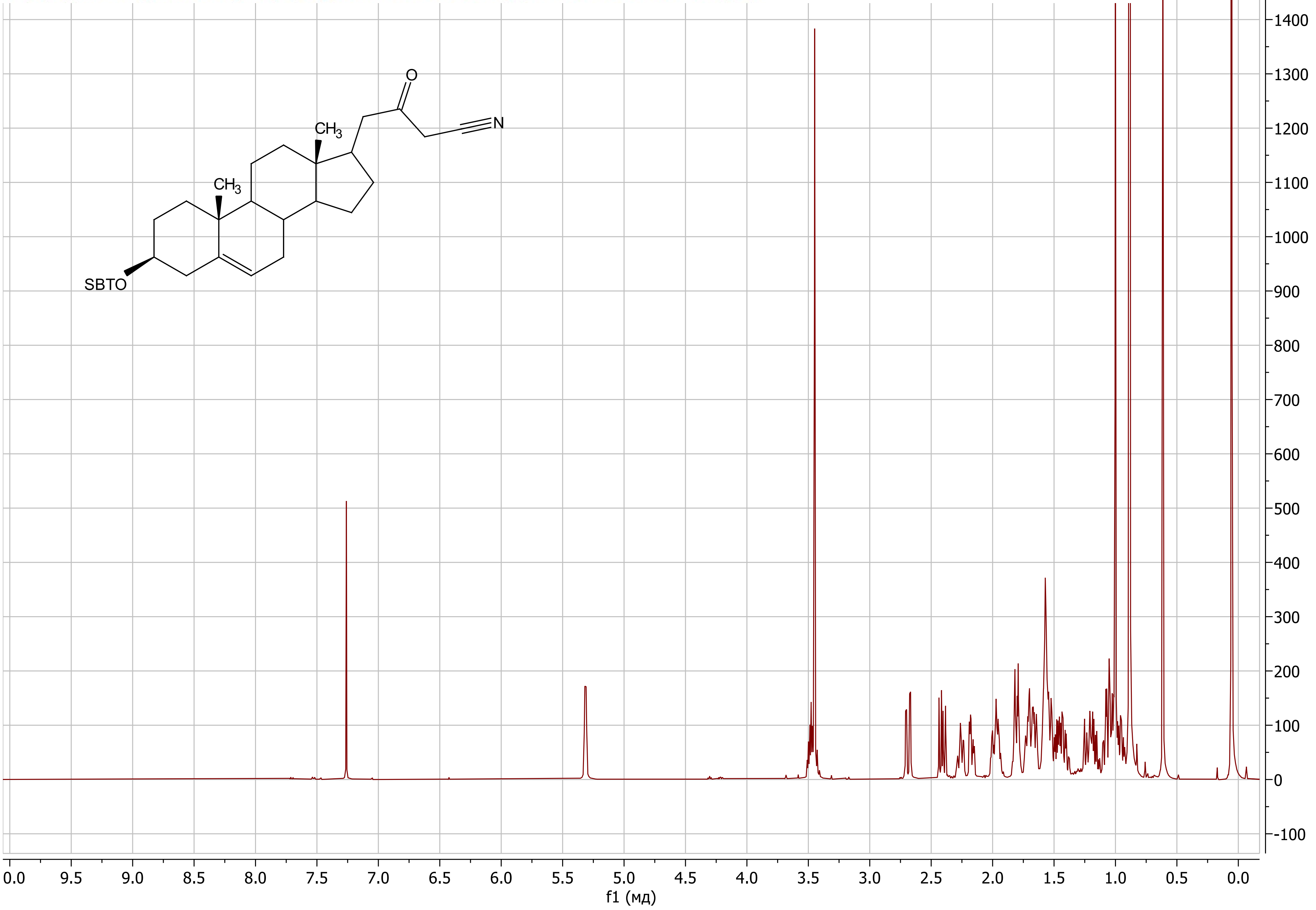

4-(3β-((*tert*-Butyldimethylsilyl)oxy)-androst-5-en-17-yl)-3-oxobutanenitrile (**31**)

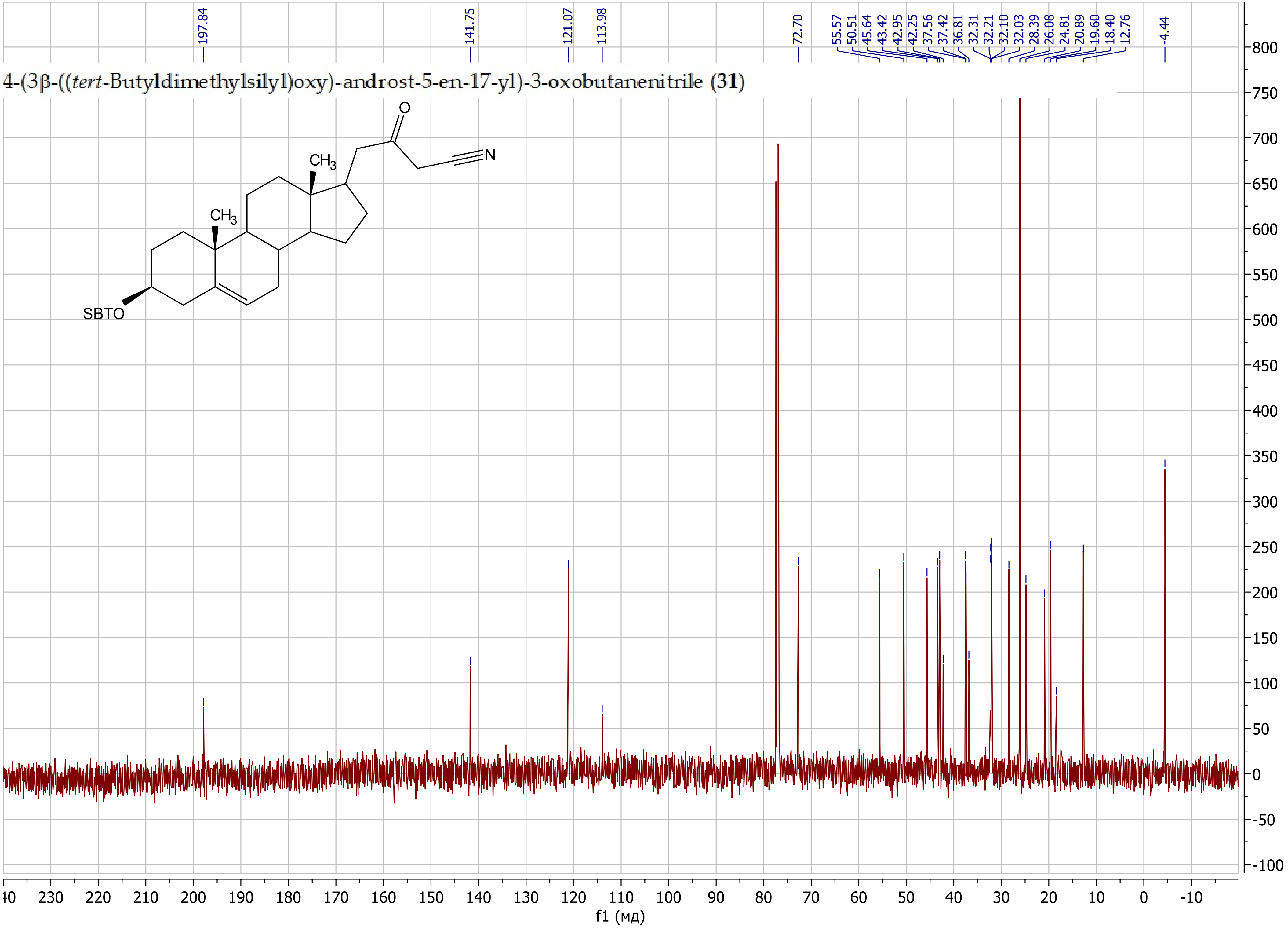

(17*R*)-17-((3-(Hydroxymethyl)isoxazol-5-yl)methyl)-androst-5-en-3β-ol (**32**)

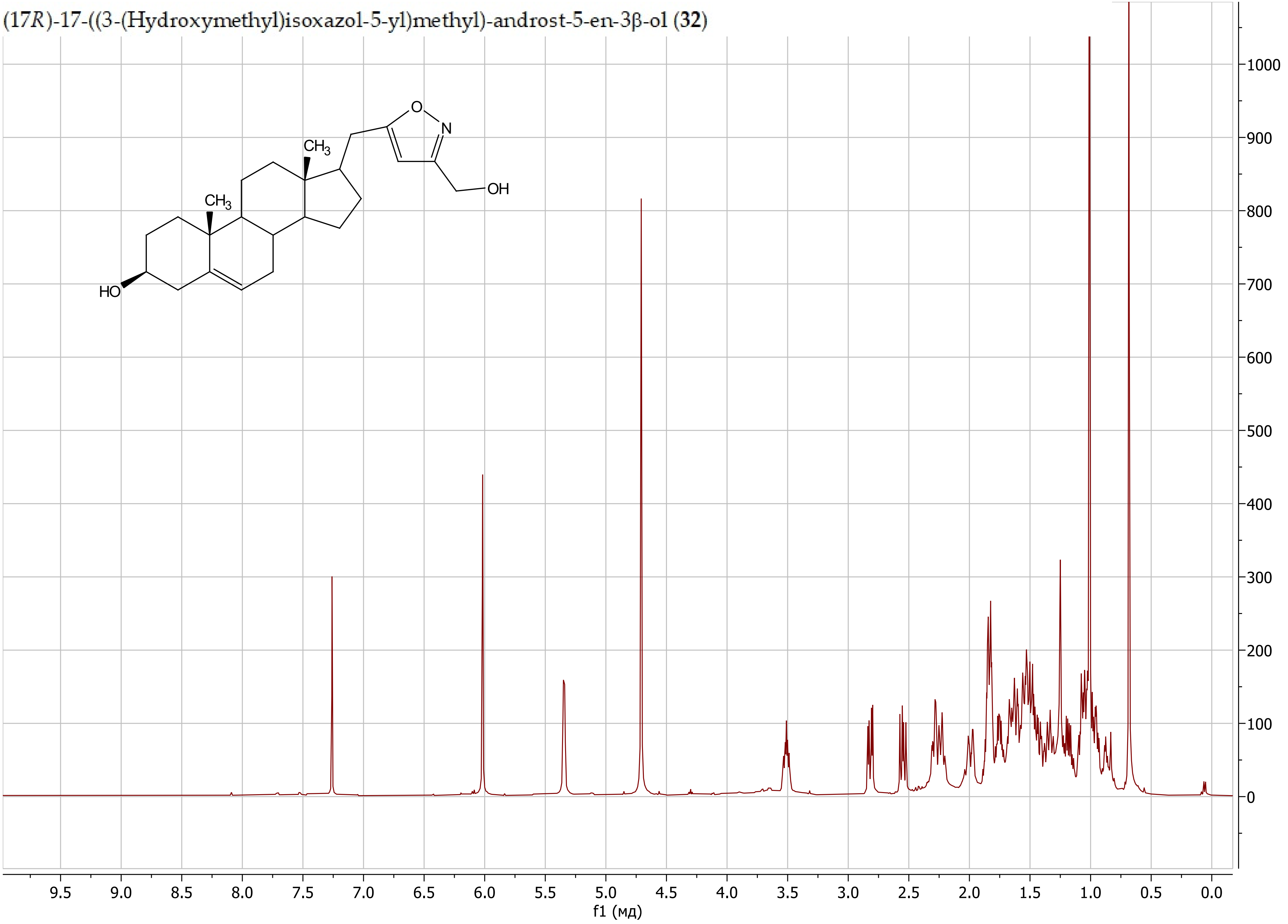

(17*R*)-17-((3-(Hydroxymethyl)isoxazol-5-yl)methyl)-androst-5-en-3β-ol (**32**)

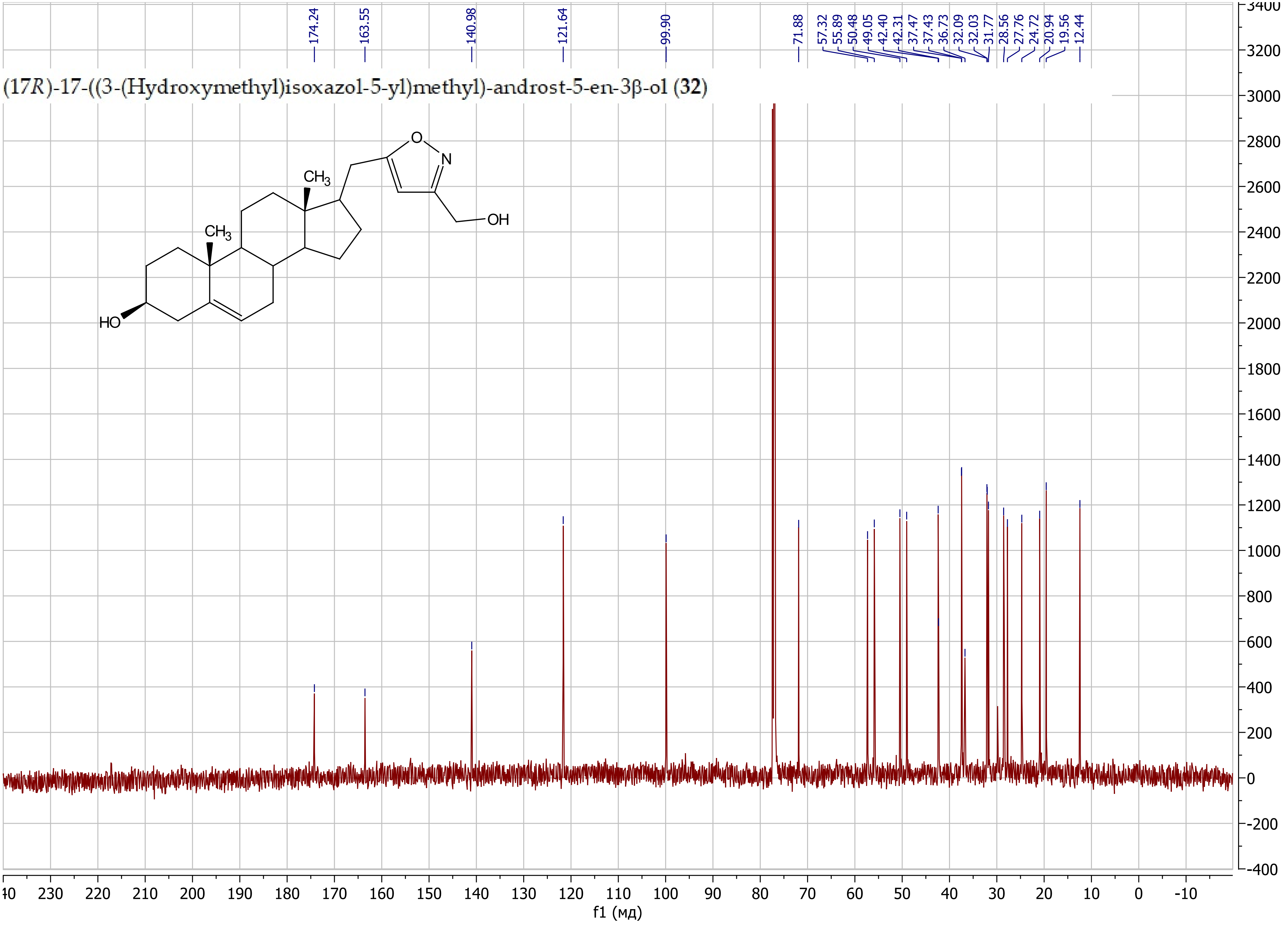

5-(((17*R*)-3β-((*tert*-Butyldimethylsilyl)oxy)-androst-5-en-17-yl)methyl)isoxazol-3-yl)methanol (**33**)

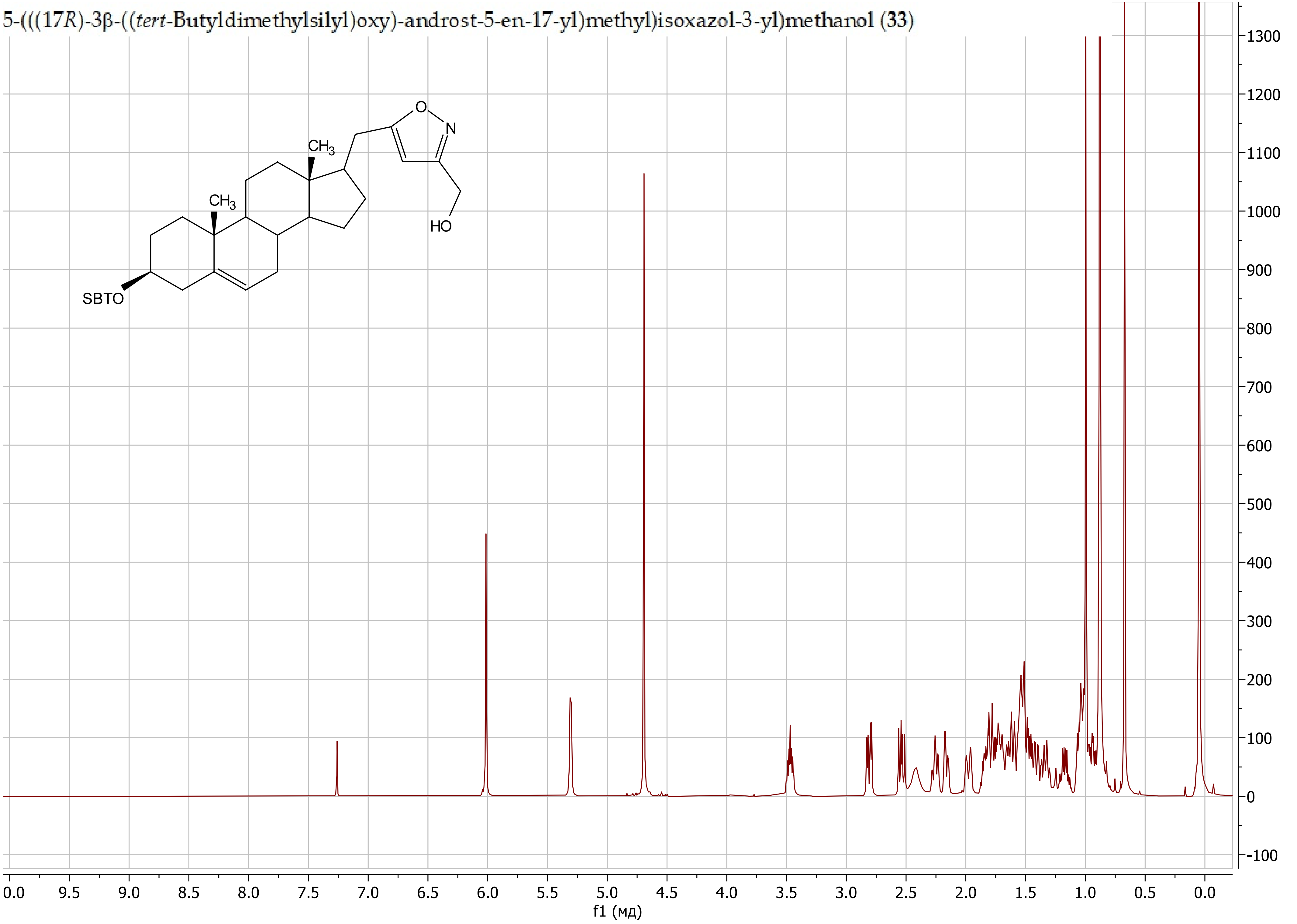

5-(((17*R*)-3β-((*tert*-Butyldimethylsilyl)oxy)-androst-5-en-17-yl)methyl)isoxazol-3-yl)methanol (**33**)

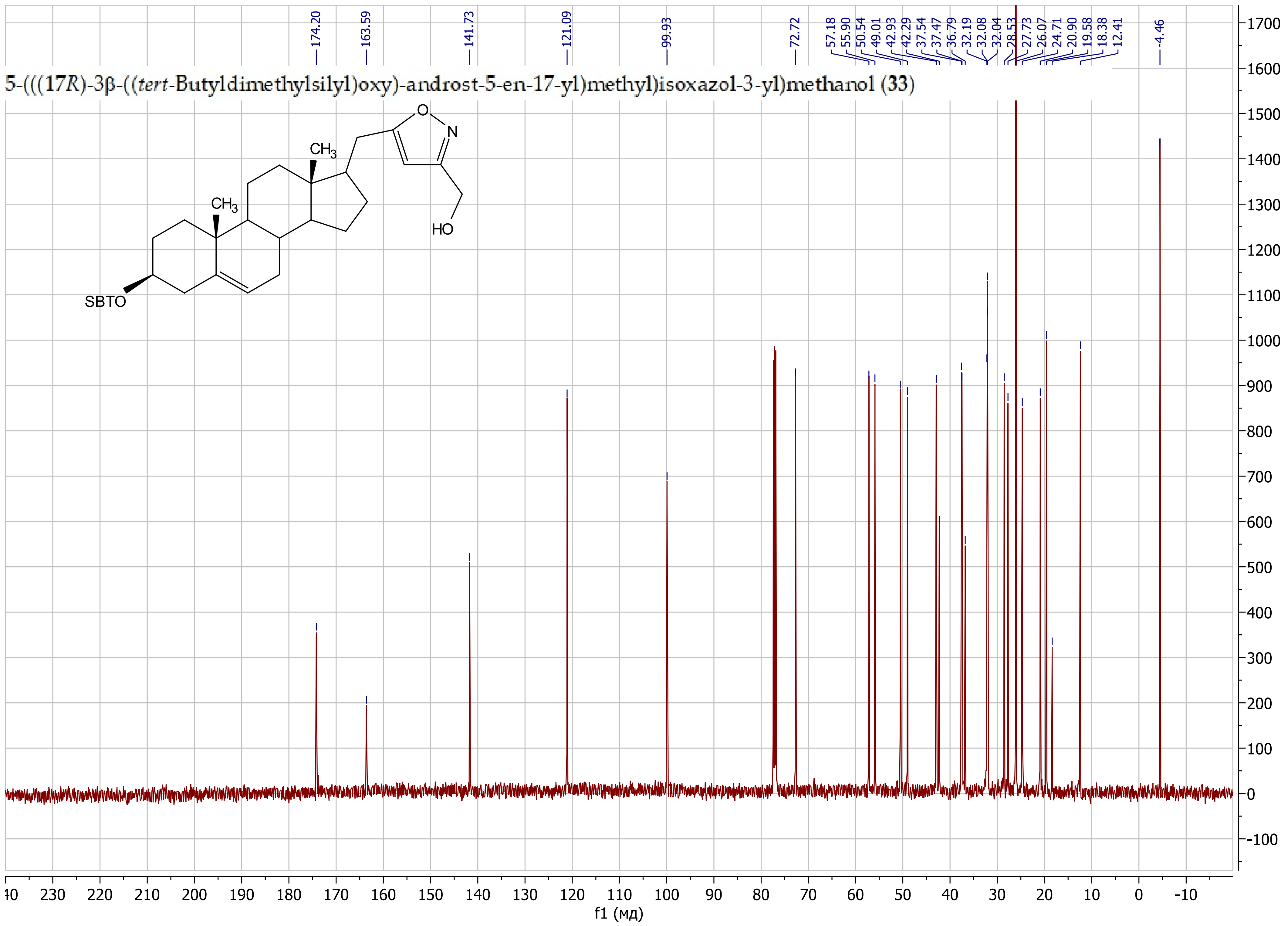

5-((3β-((*tert*-Butyldimethylsilyl)oxy)-androst-5-en-17-yl)methyl)isoxazol-3-yl)methyl methanesulfonate (**34**)

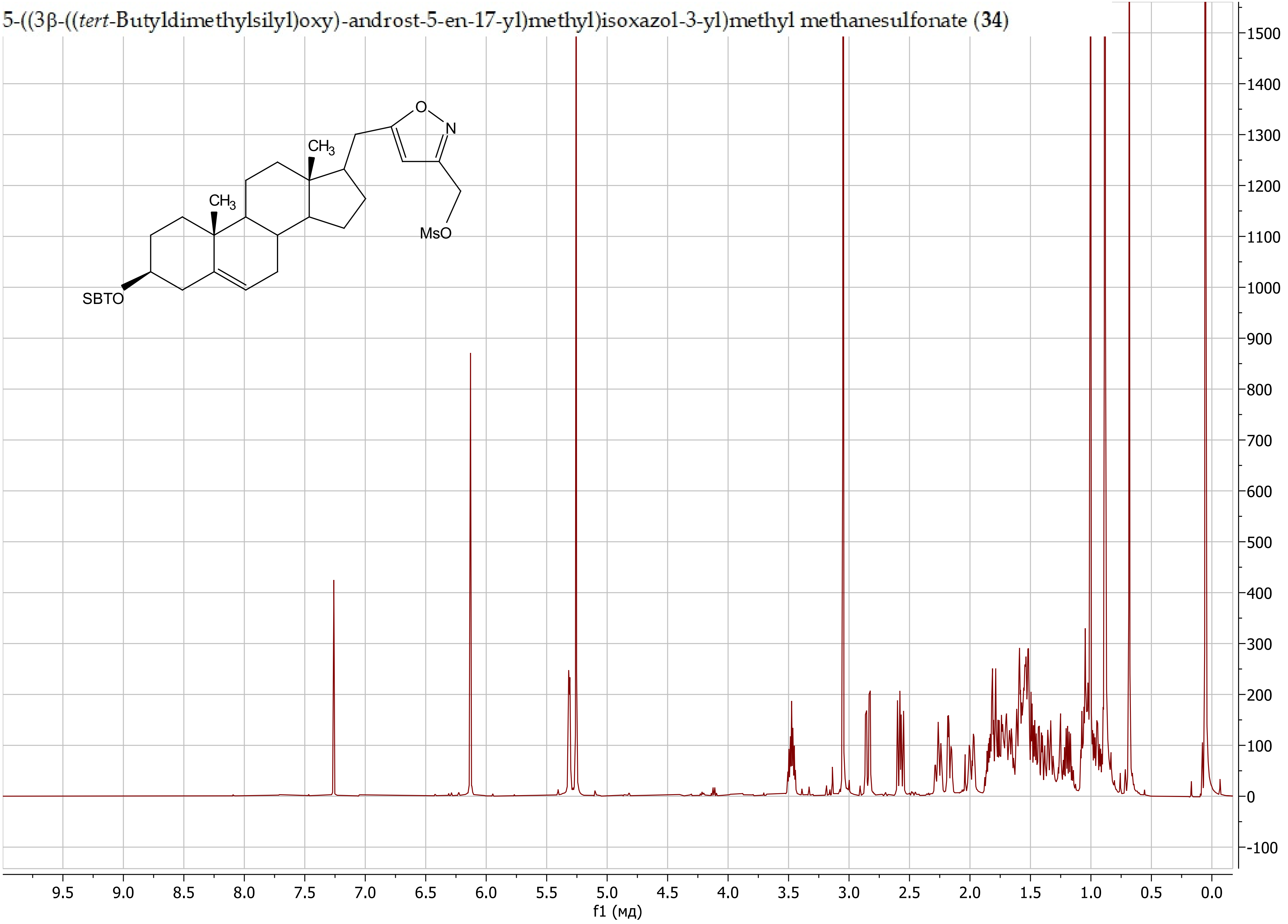

5-((3β-((*tert*-Butyldimethylsilyl)oxy)-androst-5-en-17-yl)methyl)isoxazol-3-yl)methyl methanesulfonate (**34**)

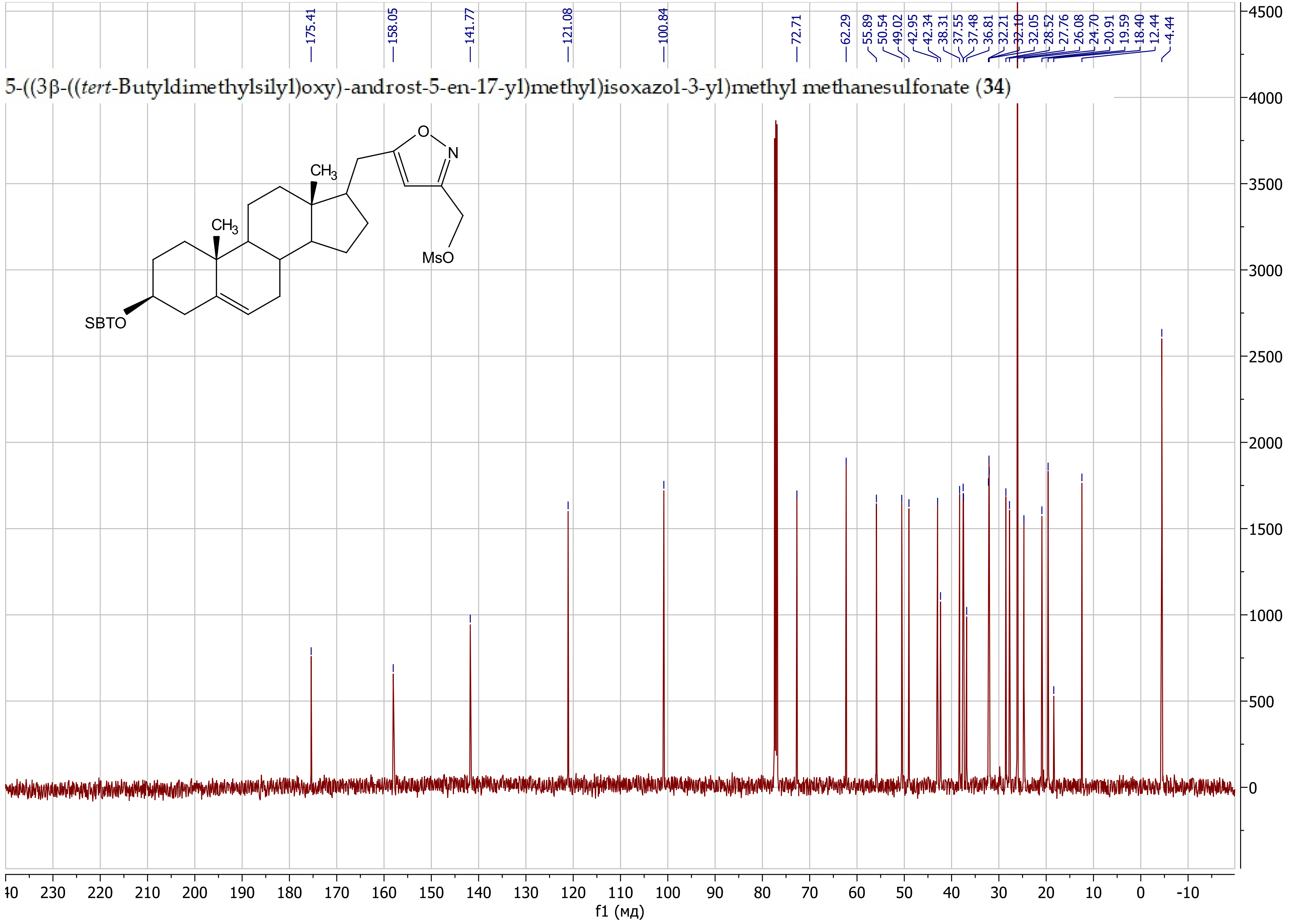

3-(Azidomethyl)-5-(((17*R*)-3β-((*tert*-butyldimethylsilyl)oxy)-androst-5-en-17-yl)methyl)isoxazole (**35**)

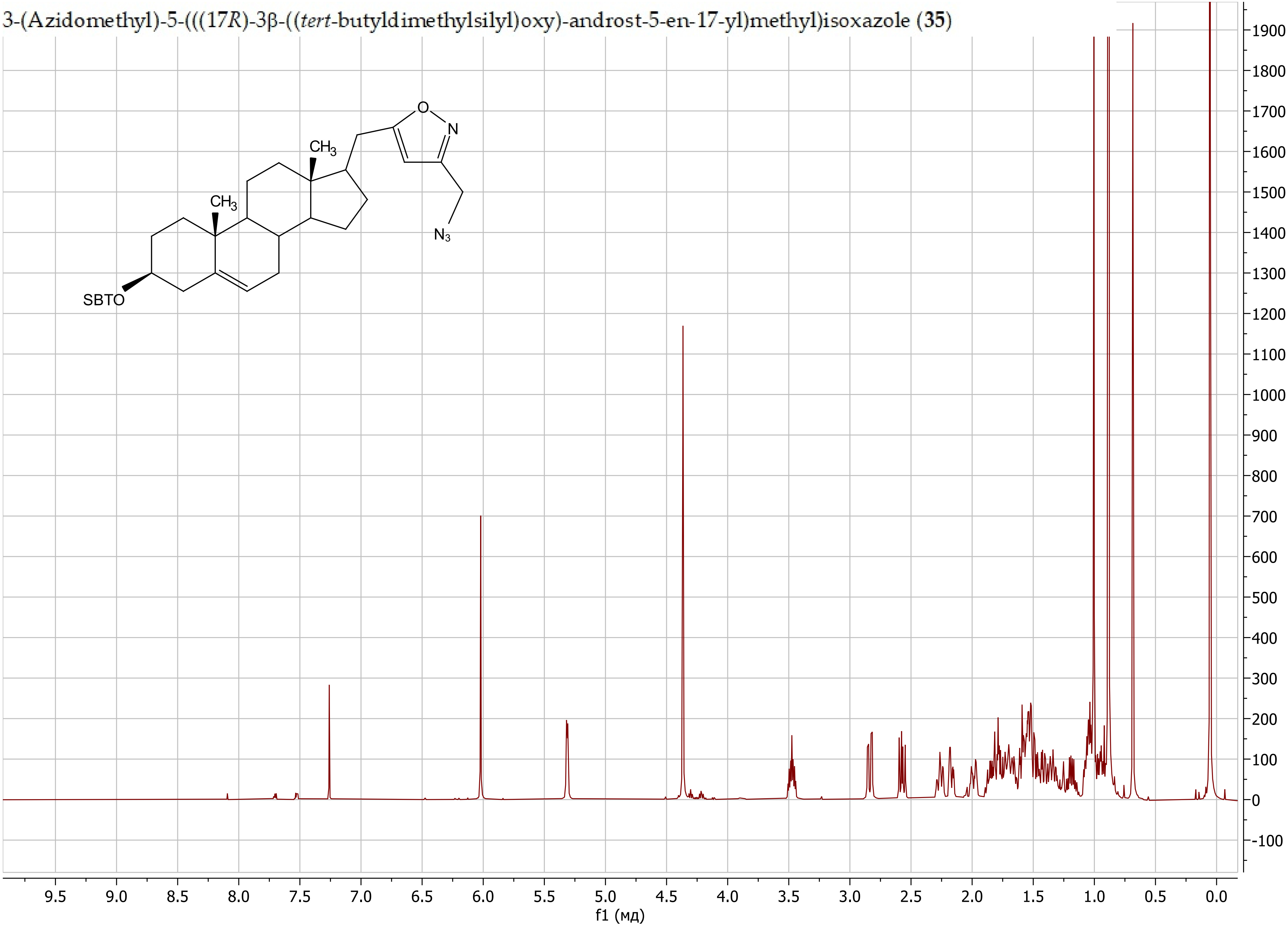

3-(Azidomethyl)-5-(((17*R*)-3β-((*tert*-butyldimethylsilyl)oxy)-androst-5-en-17-yl)methyl)isoxazole (**35**)

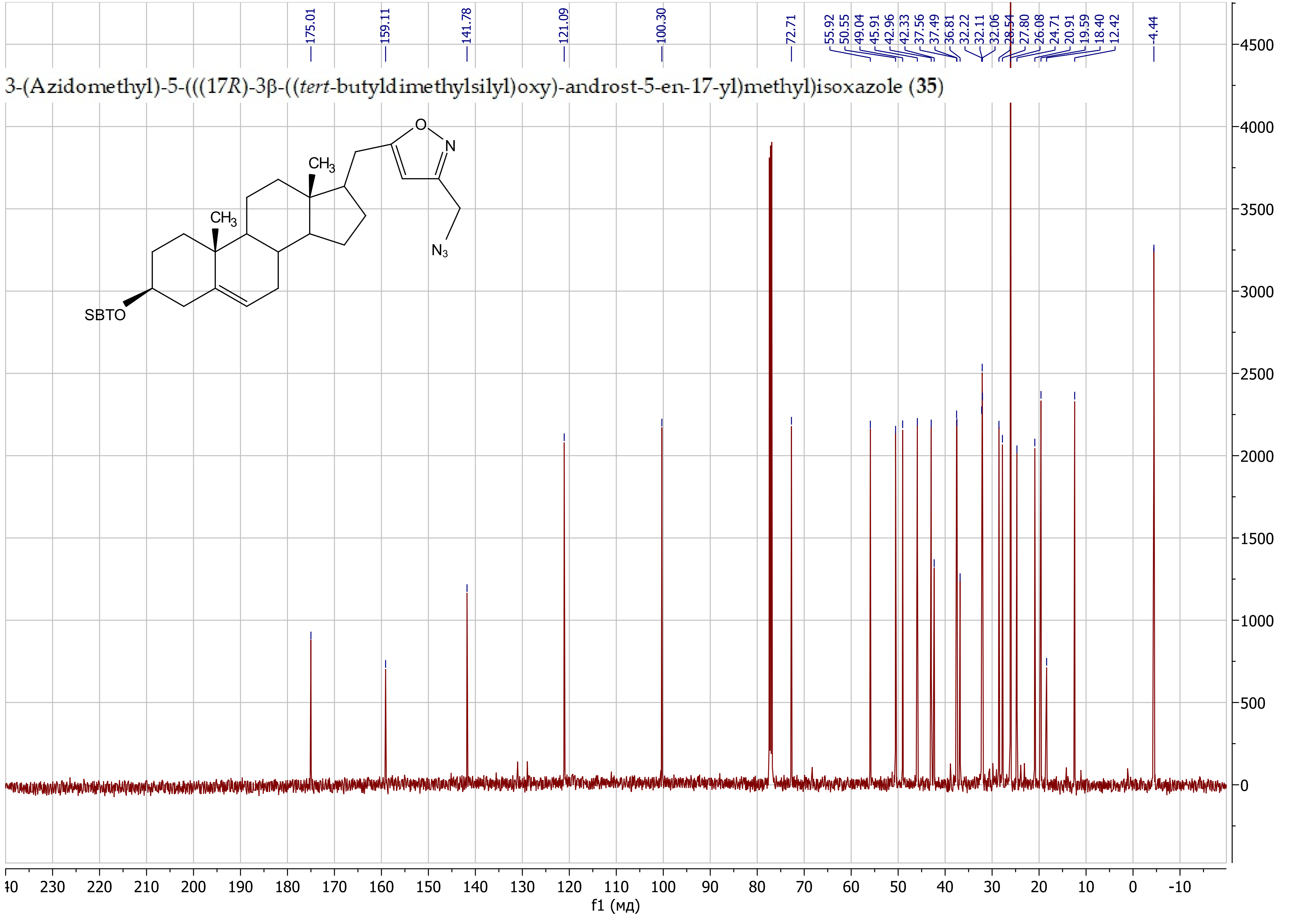

(17*R*)-17-((3-(Azidomethyl)isoxazol-5-yl)methyl)-androst-5-en-3β-ol (**36**)

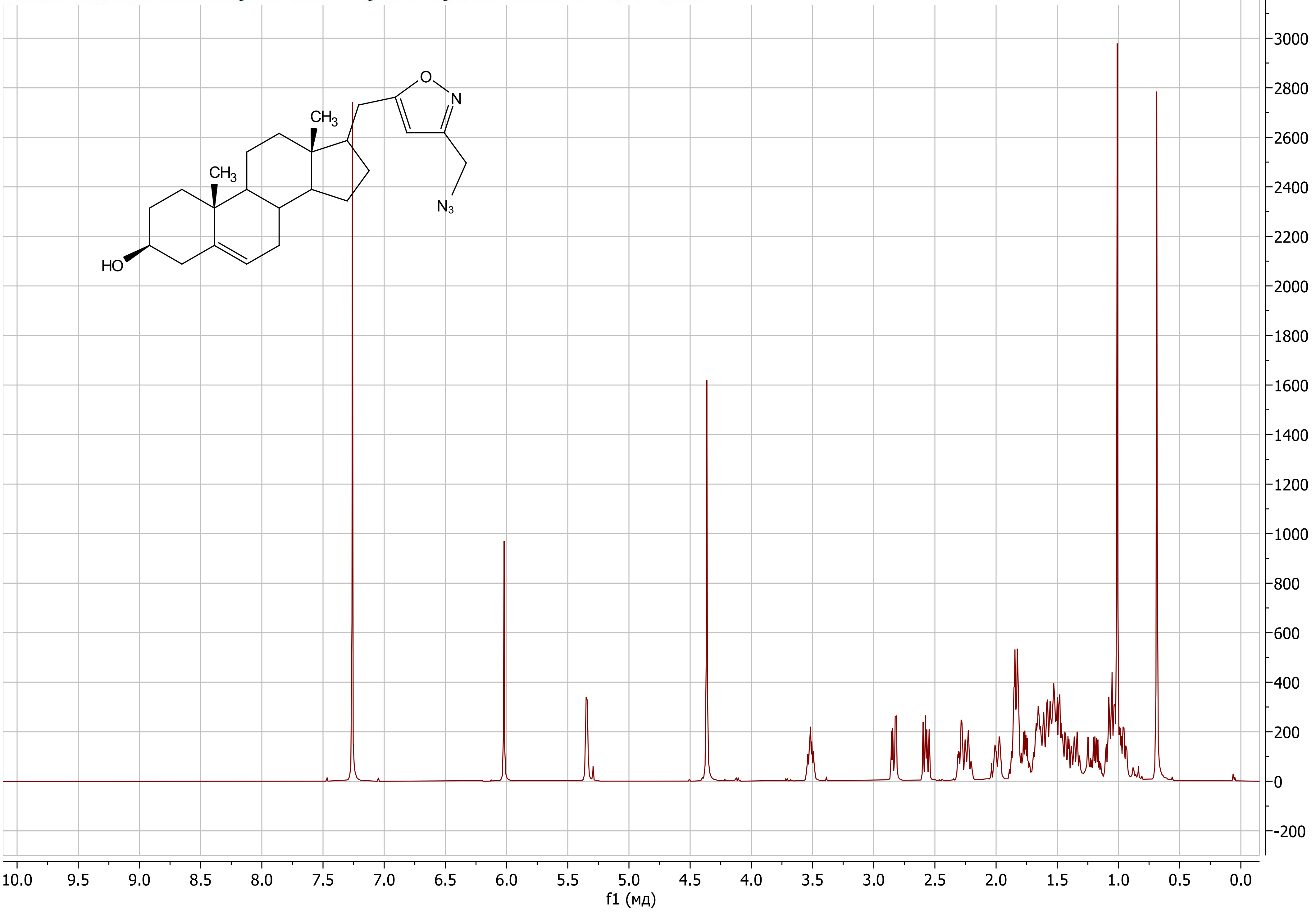

(17*R*)-17-((3-(Azidomethyl)isoxazol-5-yl)methyl)-androst-5-en-3β-ol (**36**)

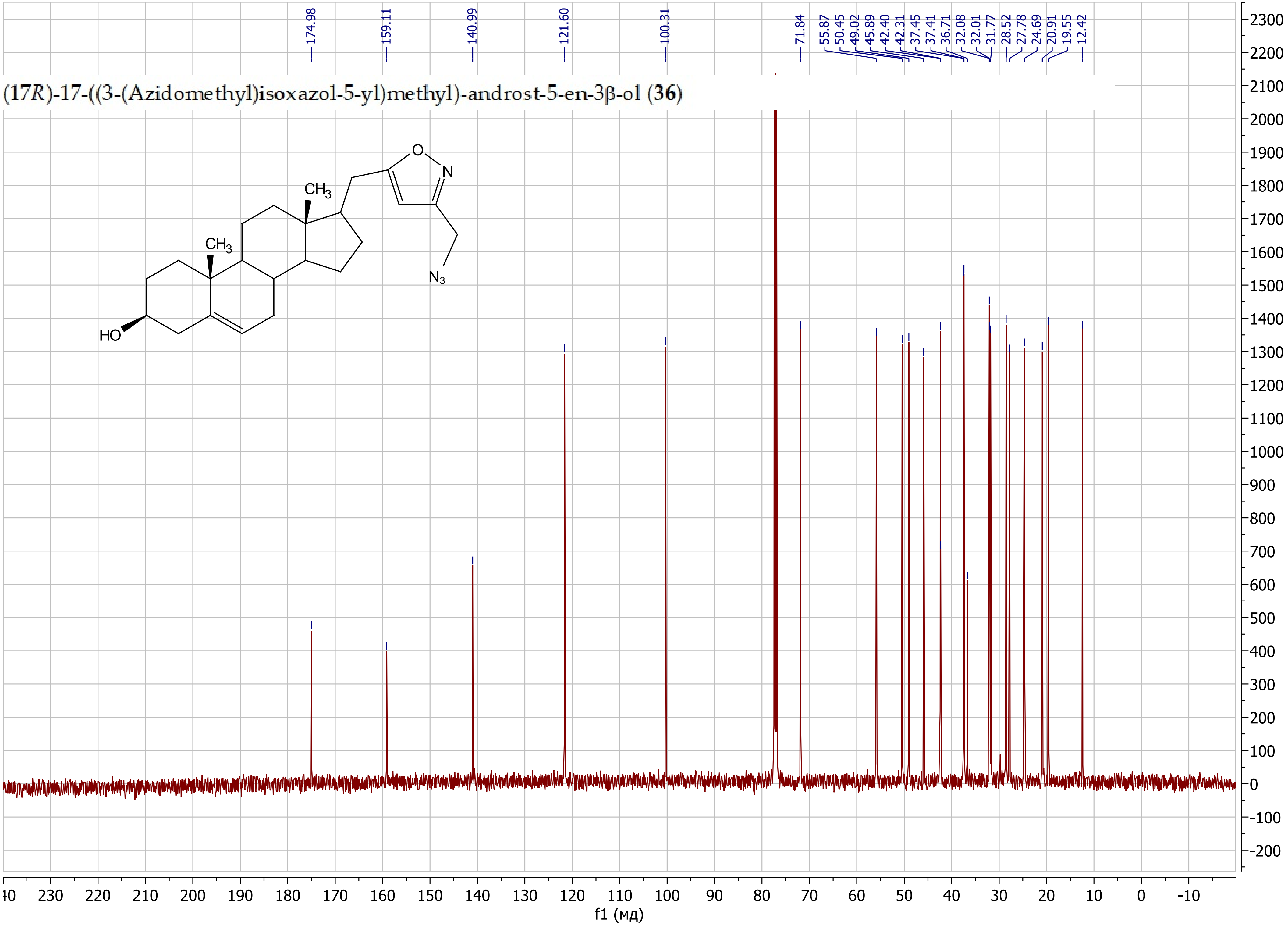

5-(((17*R*)-3β-((*tert*-Butyldimethylsilyl)oxy)-androst-5-en-17-yl)methyl)-3-(chloromethyl)isoxazole (**37**)

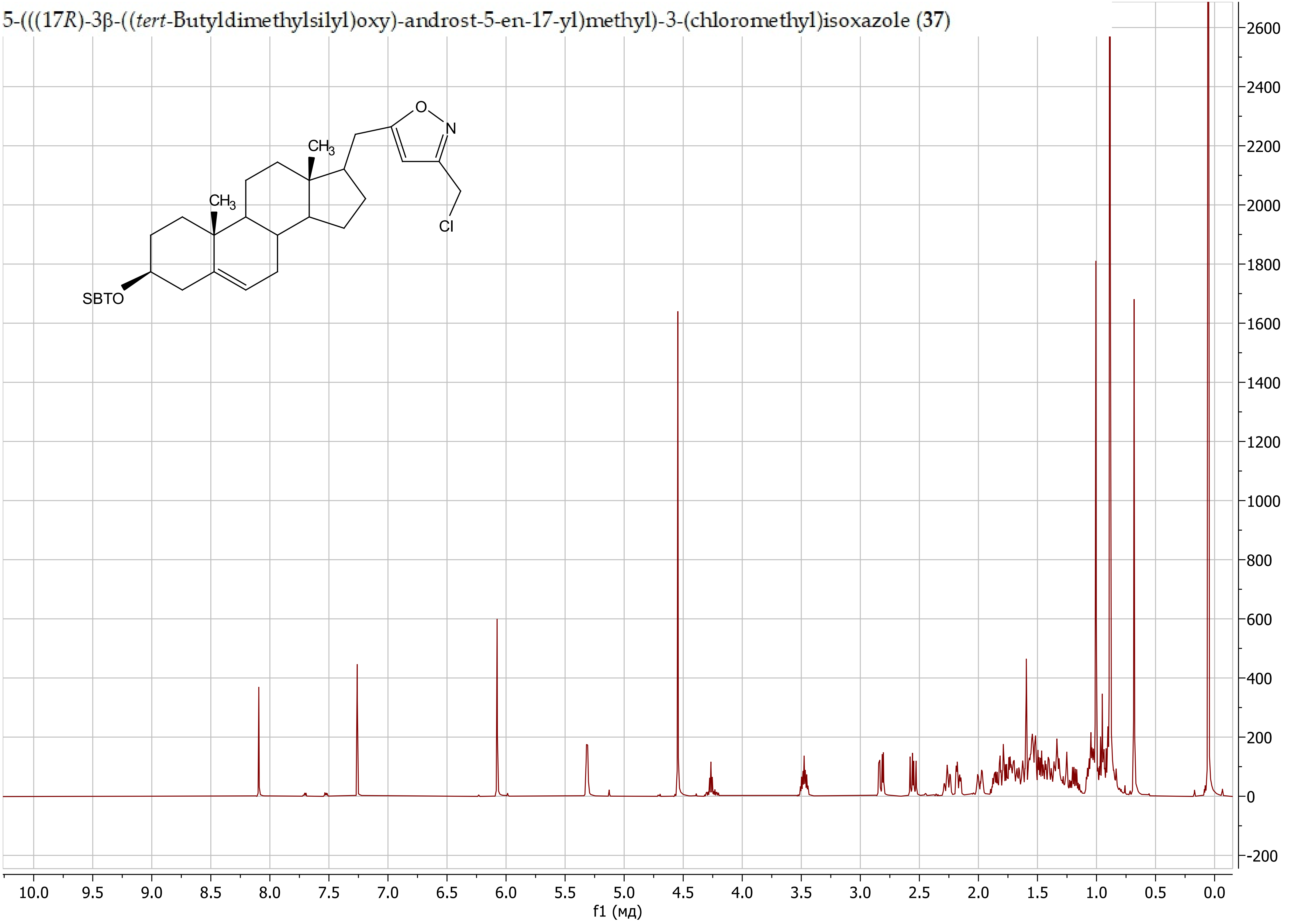

5-(((17*R*)-3β-((*tert*-Butyldimethylsilyl)oxy)-androst-5-en-17-yl)methyl)-3-(chloromethyl)isoxazole (**37**)

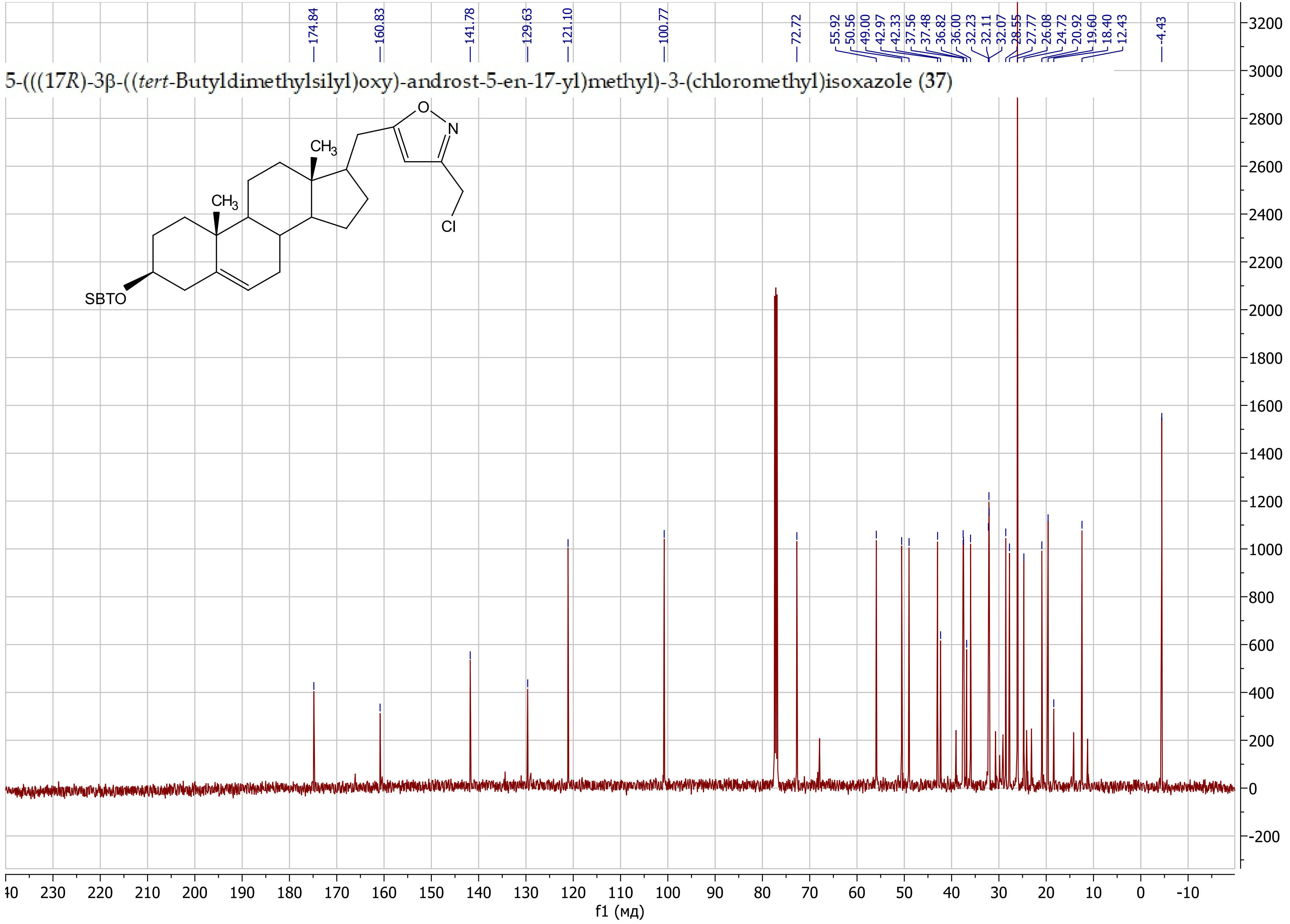

(17*R*)-17-((3-(Chloromethyl)isoxazol-5-yl)methyl)-androst-5-en-3β-ol (38)

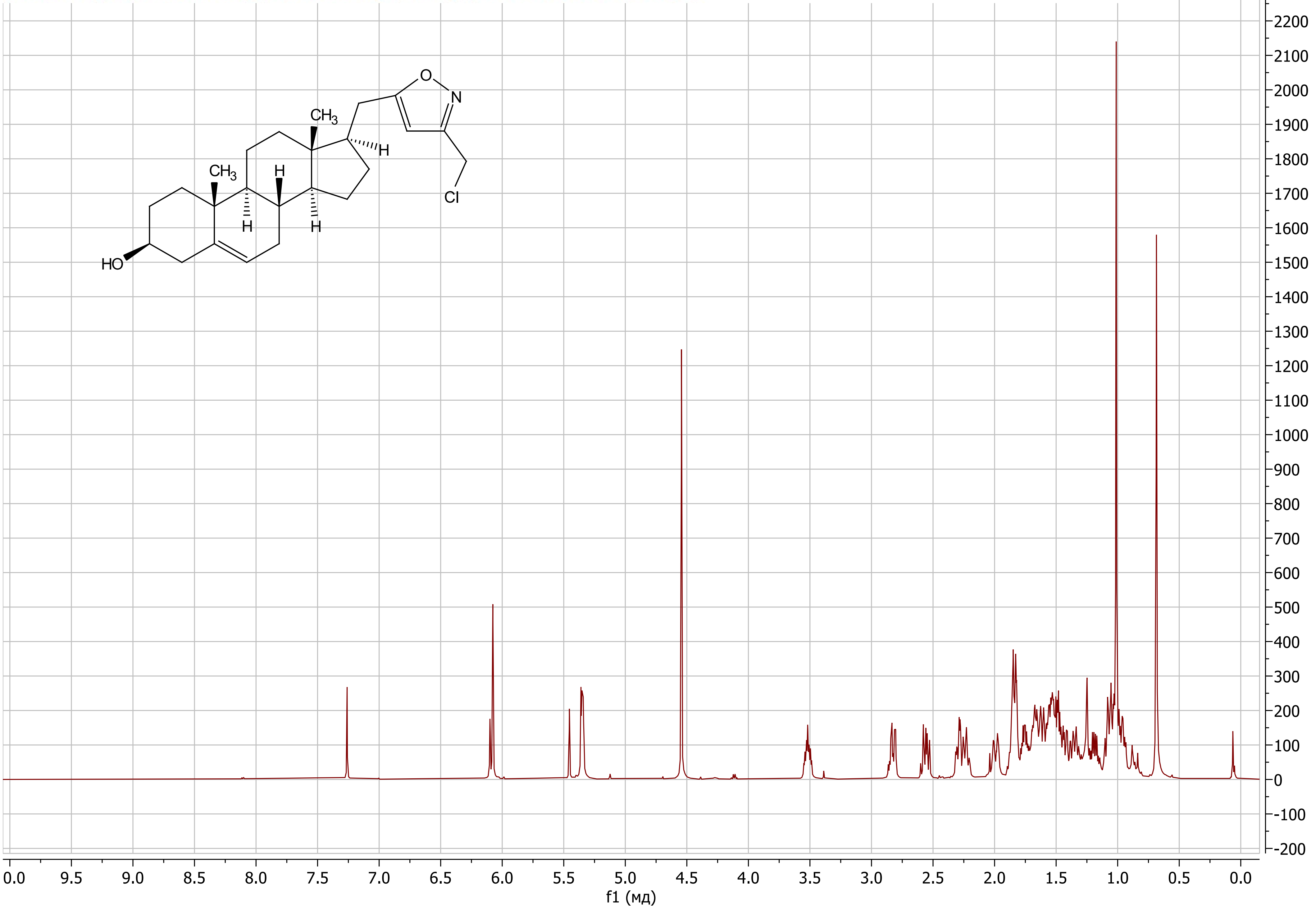

(17*R*)-17-((3-(Chloromethyl)isoxazol-5-yl)methyl)-androst-5-en-3β-ol (38)

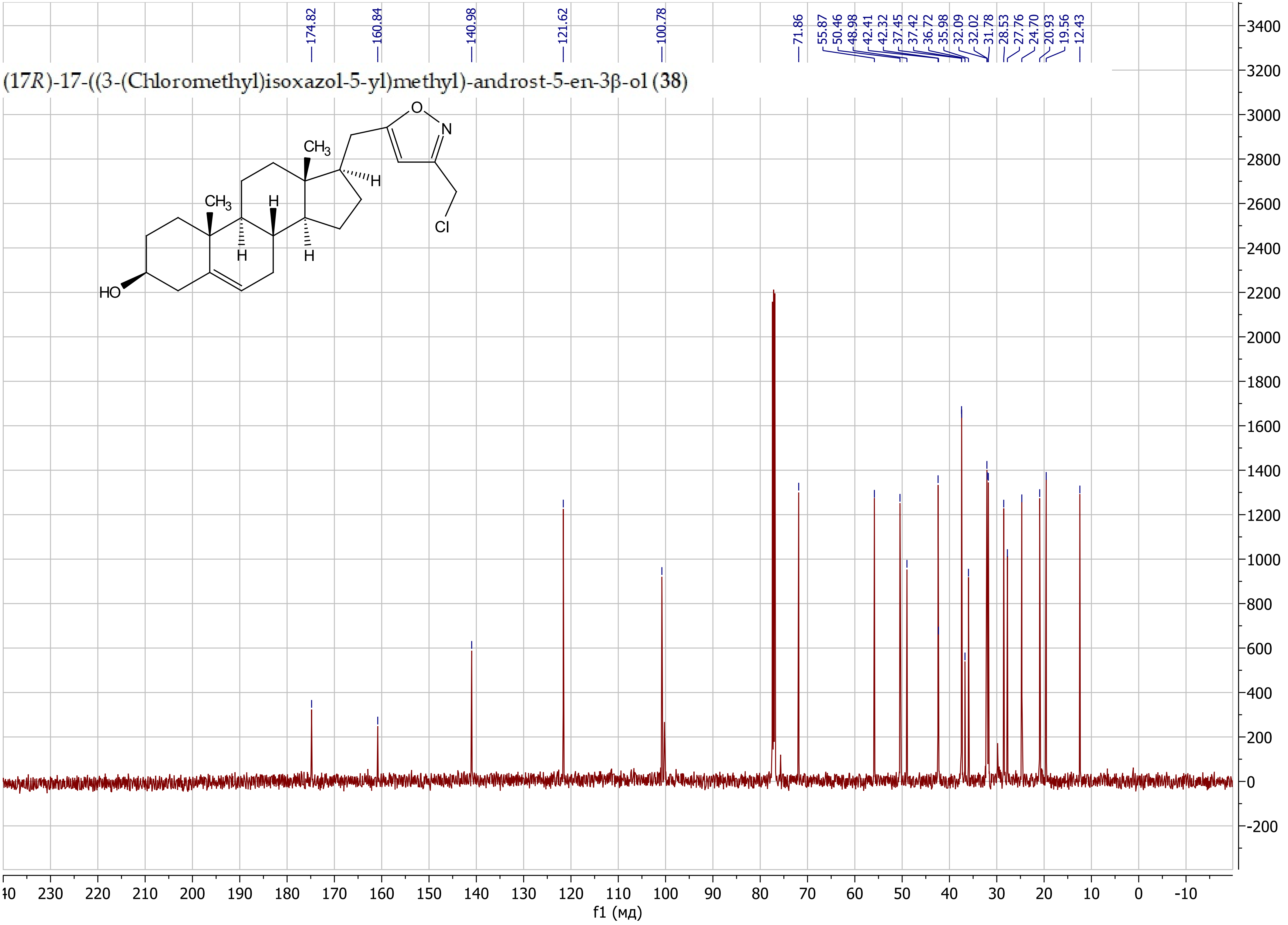

3-(((17*R*)-3β-((*tert*-Butyldimethylsilyl)oxy)-androst-5-en-17-yl)methyl)isoxazole (**40a**)

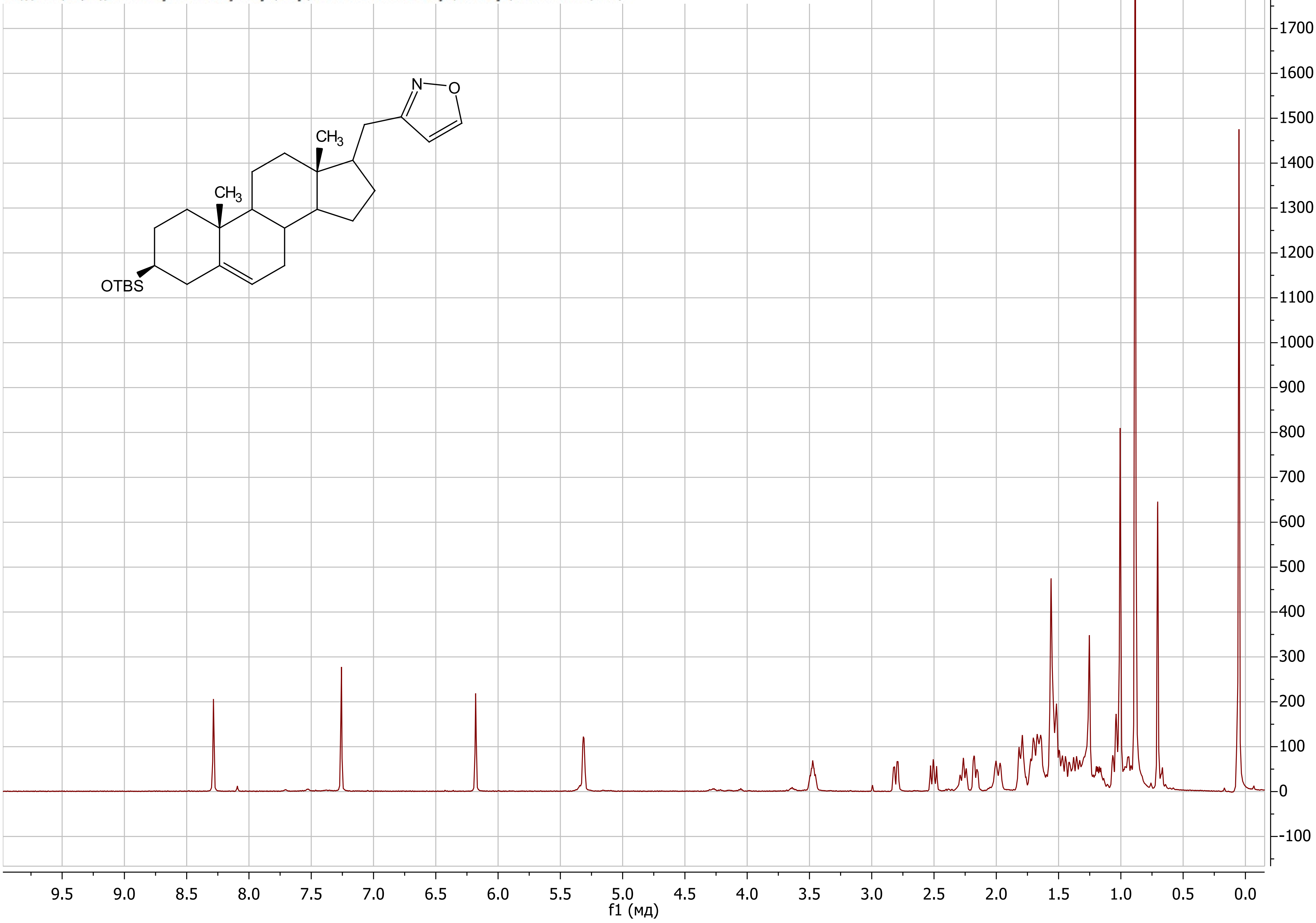

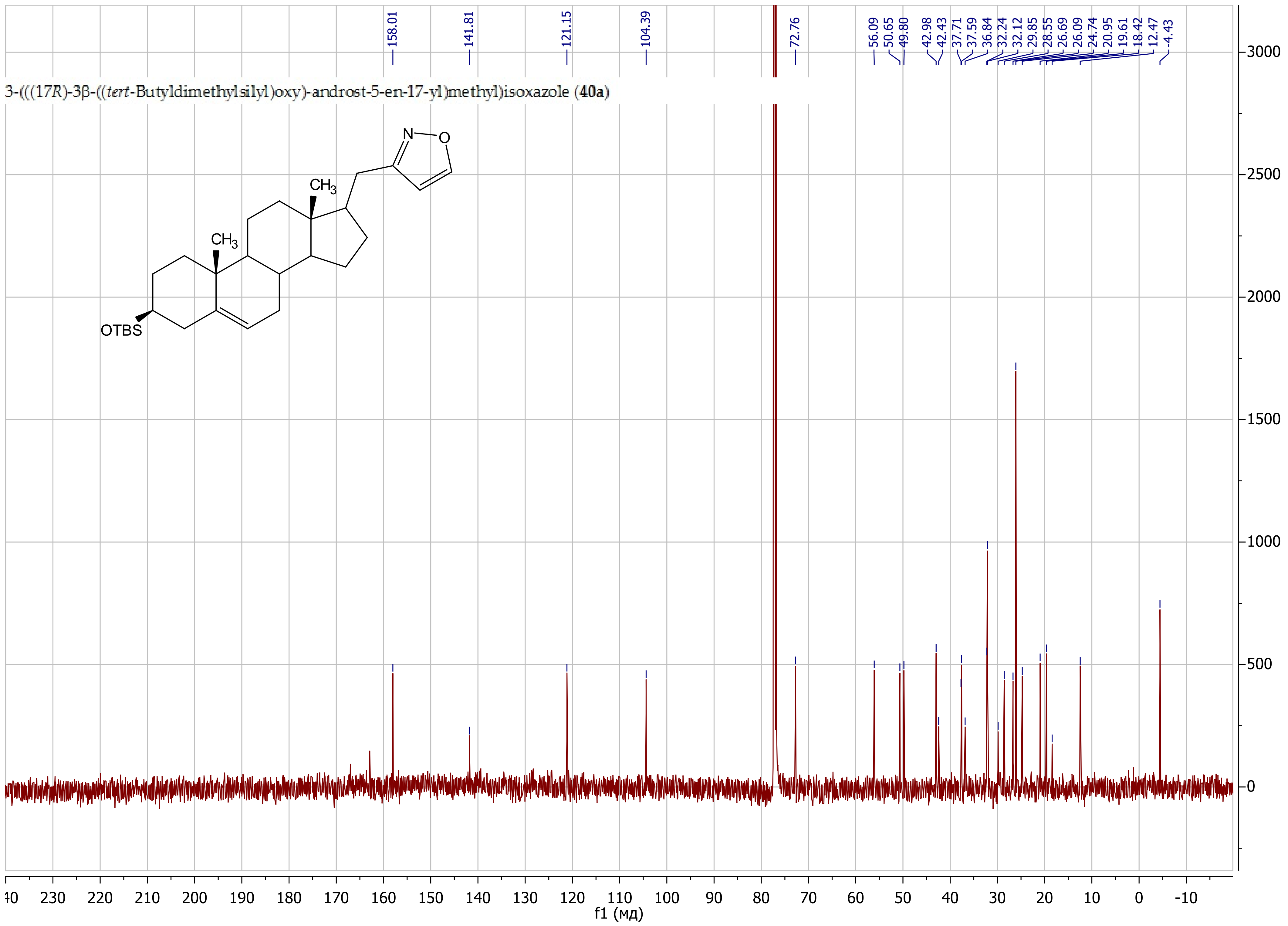

158.01
141.81
121.15
104.39
72.76
56.09
50.65
49.80
42.98
42.43
37.71
37.59
36.84
32.24
32.12
29.85
28.55
26.69
26.09
24.74
20.95
19.61
18.42
12.47
-4.43
3-(((17R)-3β-((tert-Butyldimethylsilyl)oxy)-androst-5-en-17-yl)methyl)isoxazole (40a)
N
O
CH3
CH3
OTBS
f1 (мд)

5-Butyl-3-(((17*R*)-3β-((*tert*-butyldimethylsilyl)oxy)-androst-5-en-17-yl)methyl)isoxazole (**40d**)

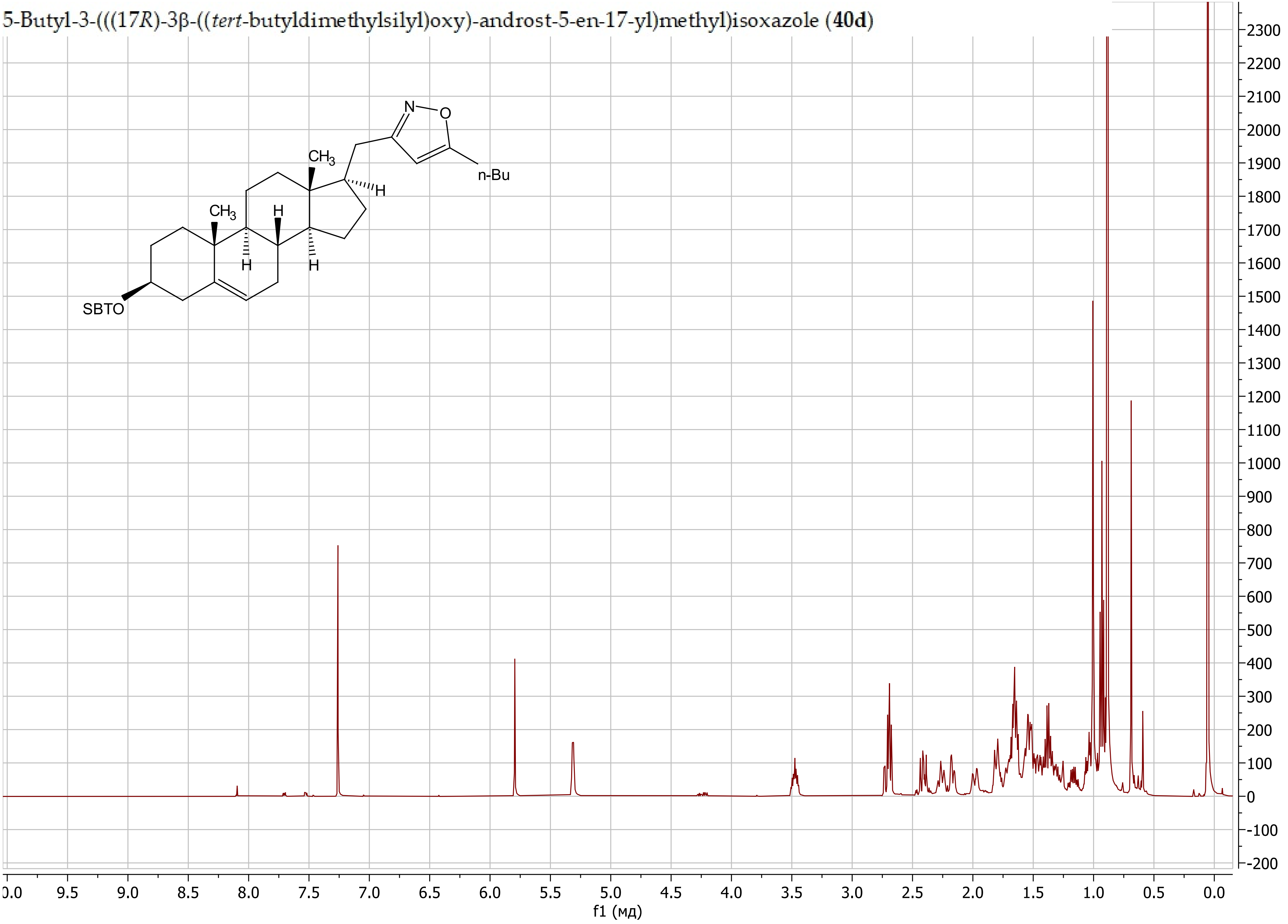

5-Butyl-3-(((17*R*)-3β-((*tert*-butyldimethylsilyl)oxy)-androst-5-en-17-yl)methyl)isoxazole (**40d**)

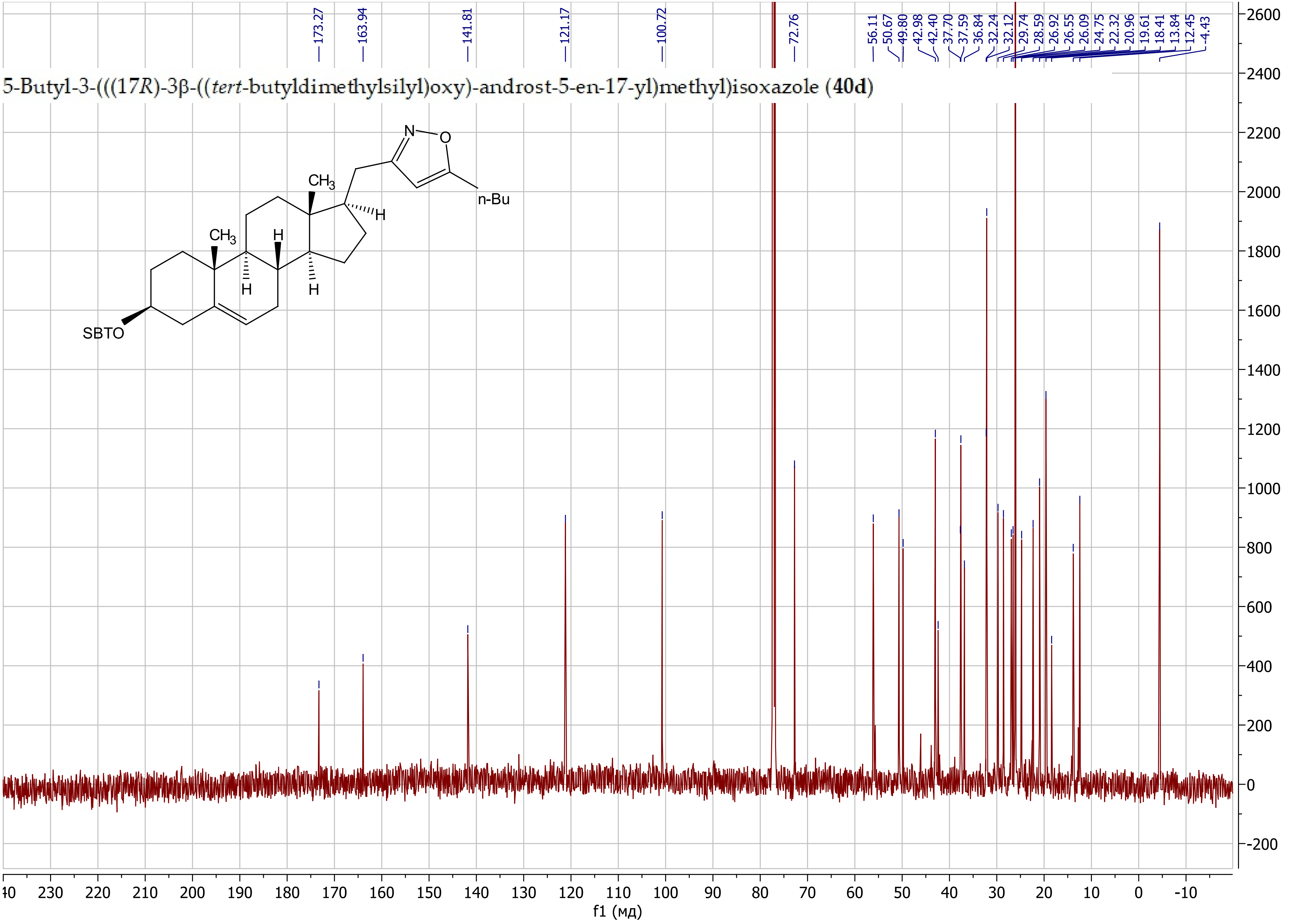

3-(((17*R*)-3β-((*tert*-Butyldimethylsilyl)oxy)-androst-5-en-17-yl)methyl)-5-phenylisoxazole (**40e**)

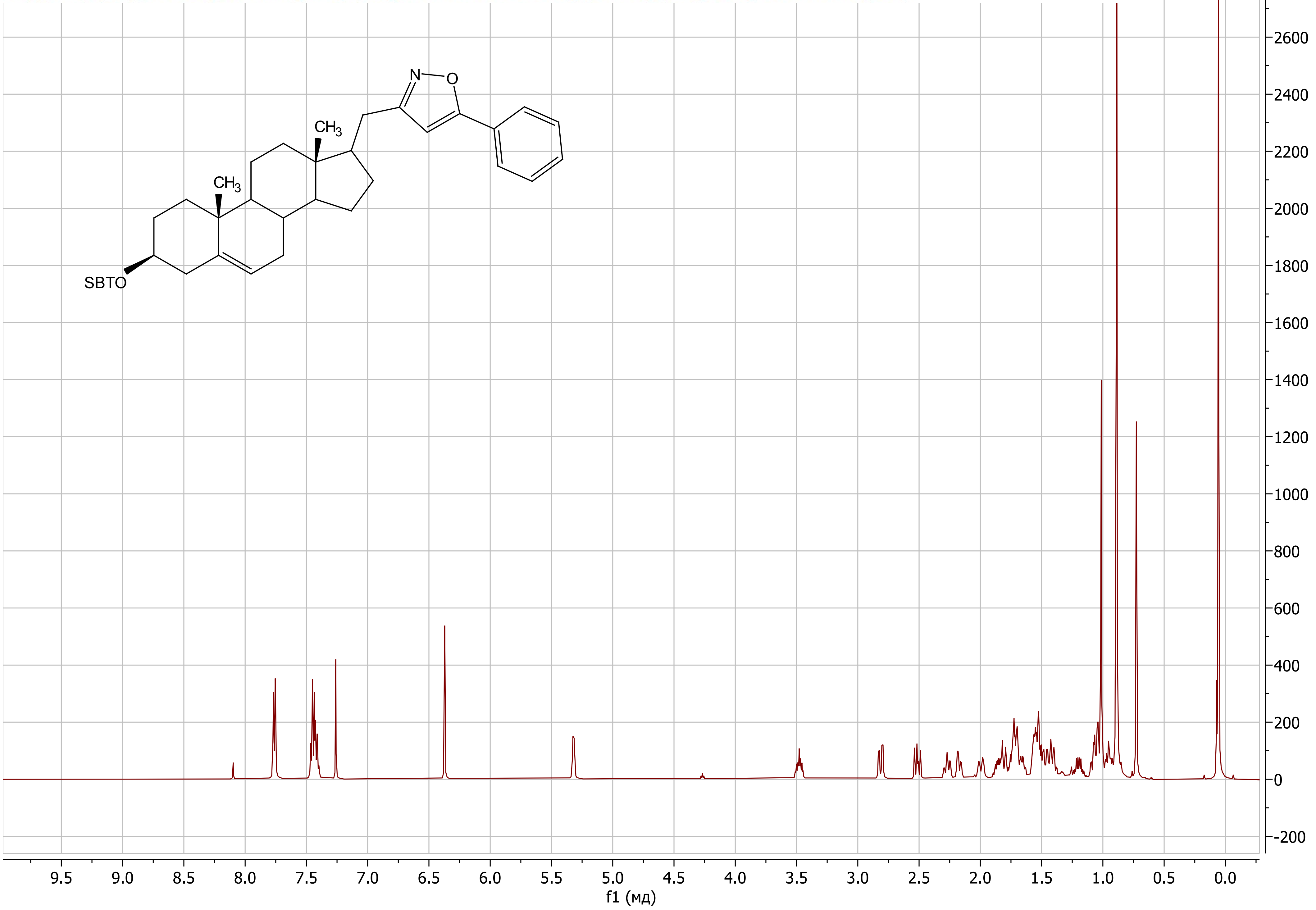

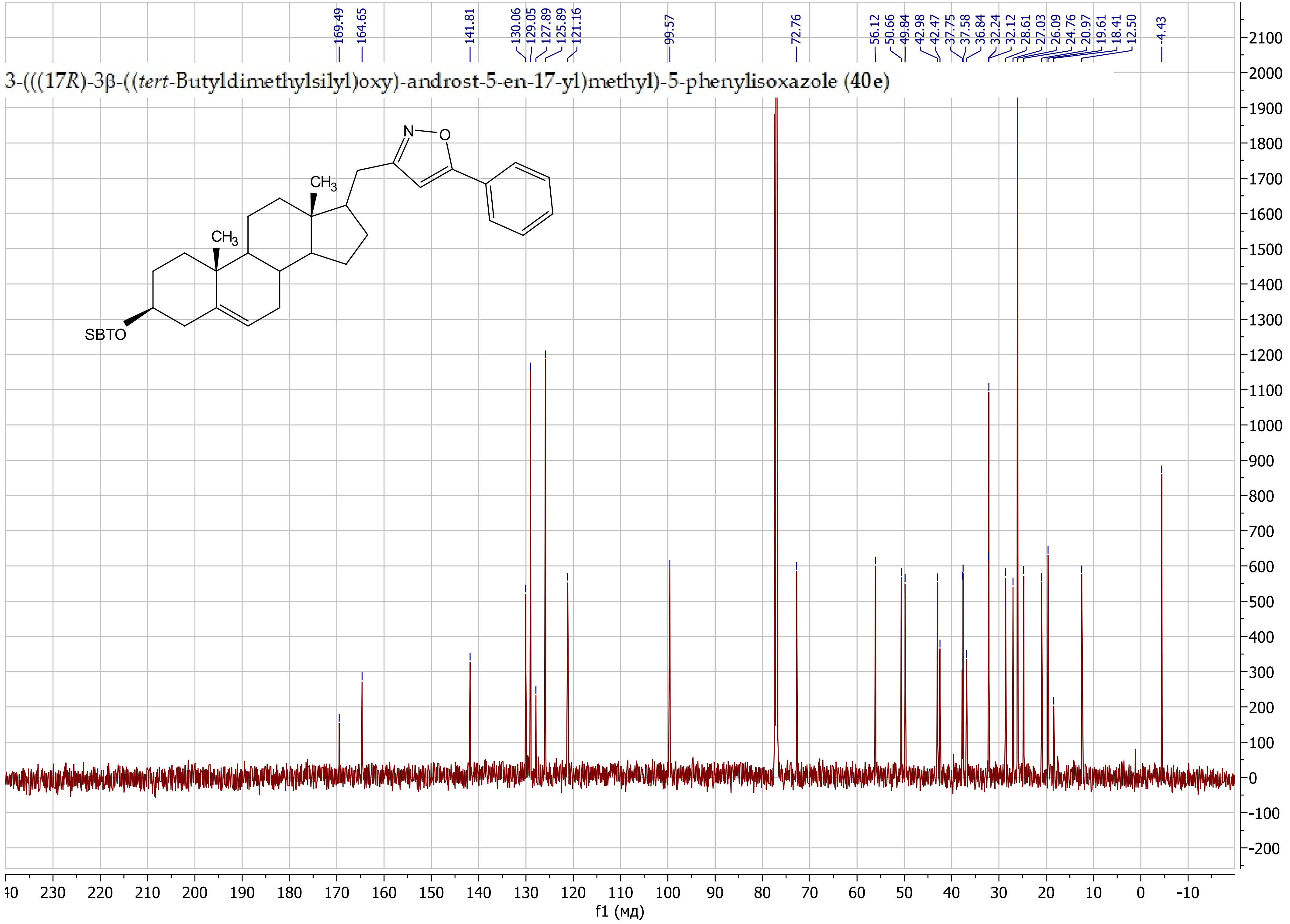
3-(((17R)-3β-((tert-Butyldimethylsilyl)oxy)-androst-5-en-17-yl)methyl)-5-phenylisoxazole (40e)
169.49
164.65
141.81
130.06
129.05
127.89
125.89
121.16
99.57
72.76
56.12
50.66
49.84
42.98
42.47
37.75
37.58
36.84
32.24
32.12
28.61
27.03
26.09
24.76
20.97
19.61
18.41
12.50
-4.43
N
O
CH3
CH3
SBTO
f1 (мд)

3-(((17*R*)-3β-((*tert*-Butyldimethylsilyl)oxy)-androst-5-en-17-yl)methyl)-5-(pyridin-3-yl)isoxazole (**40f**)

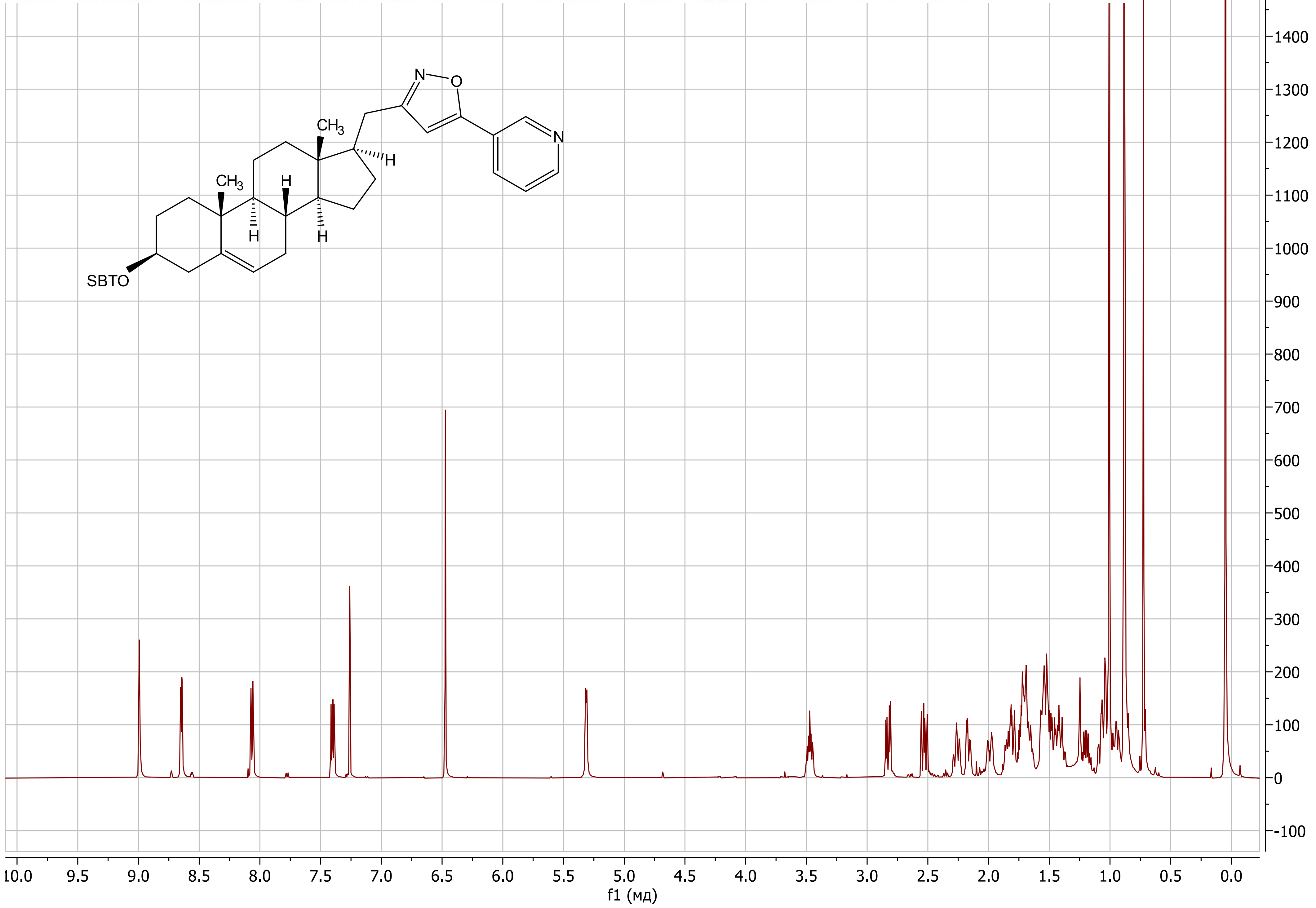

3-(((17*R*)-3β-((*tert*-Butyldimethylsilyl)oxy)-androst-5-en-17-yl)methyl)-5-(pyridin-3-yl)isoxazole (**40f**)

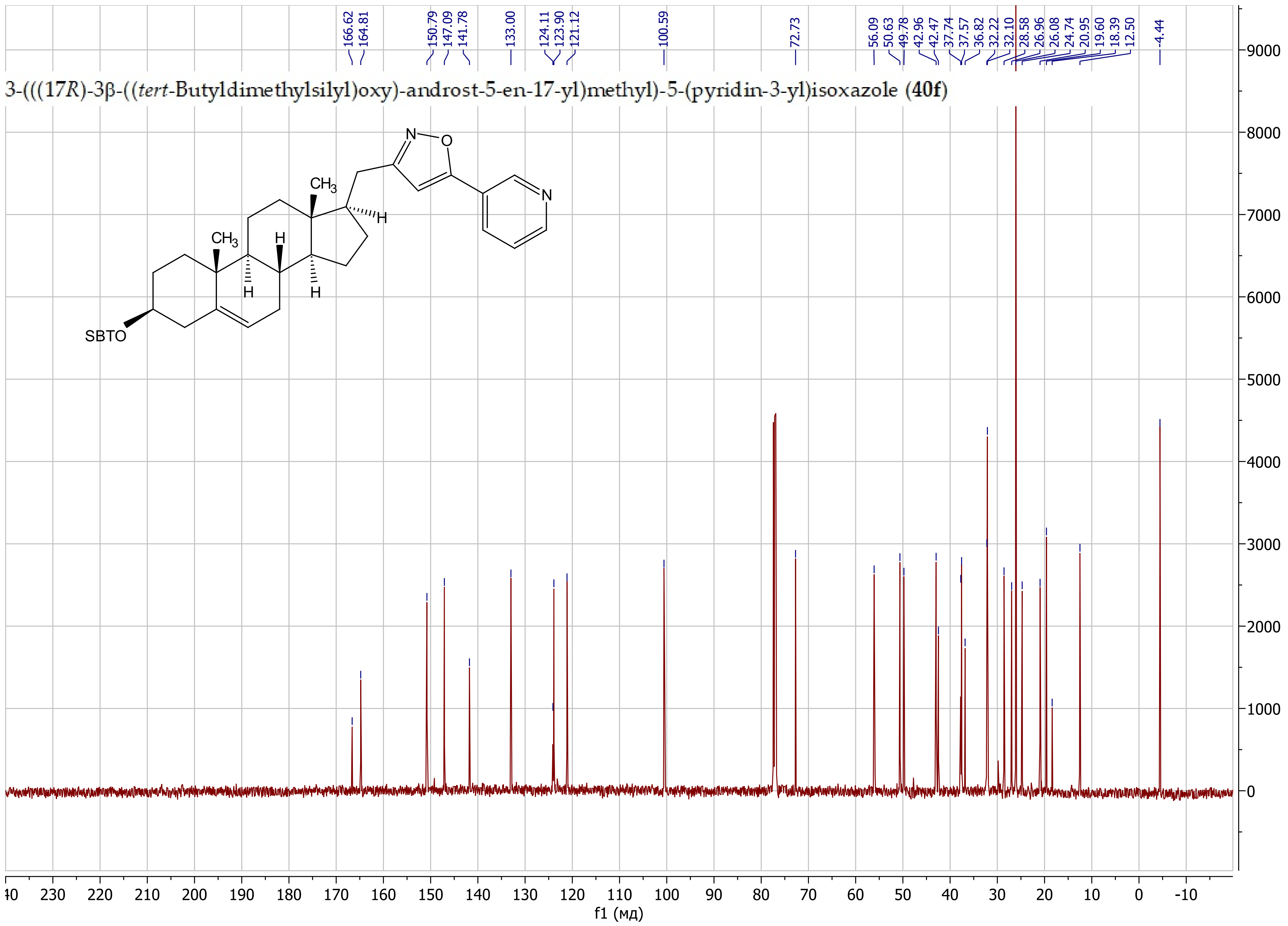

3-(((17R)-3β-((*tert*-Butyldimethylsilyl)oxy)-androst-5-en-17-yl)methyl)-5-(2-fluorophenyl)isoxazole (**40g**)

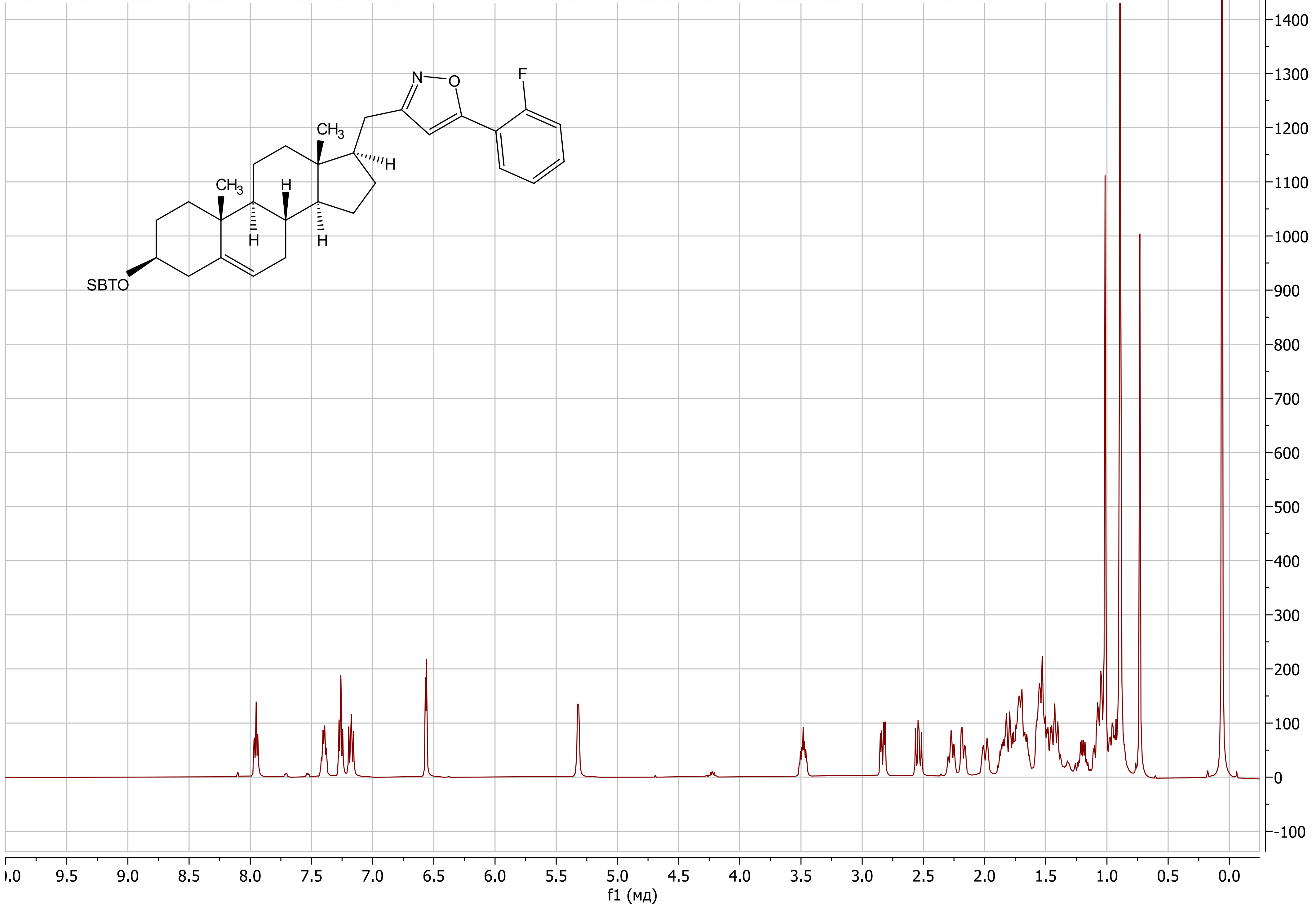

3-(((17R)-3β-((*tert*-Butyldimethylsilyl)oxy)-androst-5-en-17-yl)methyl)-5-(2-fluorophenyl)isoxazole (**40g**)

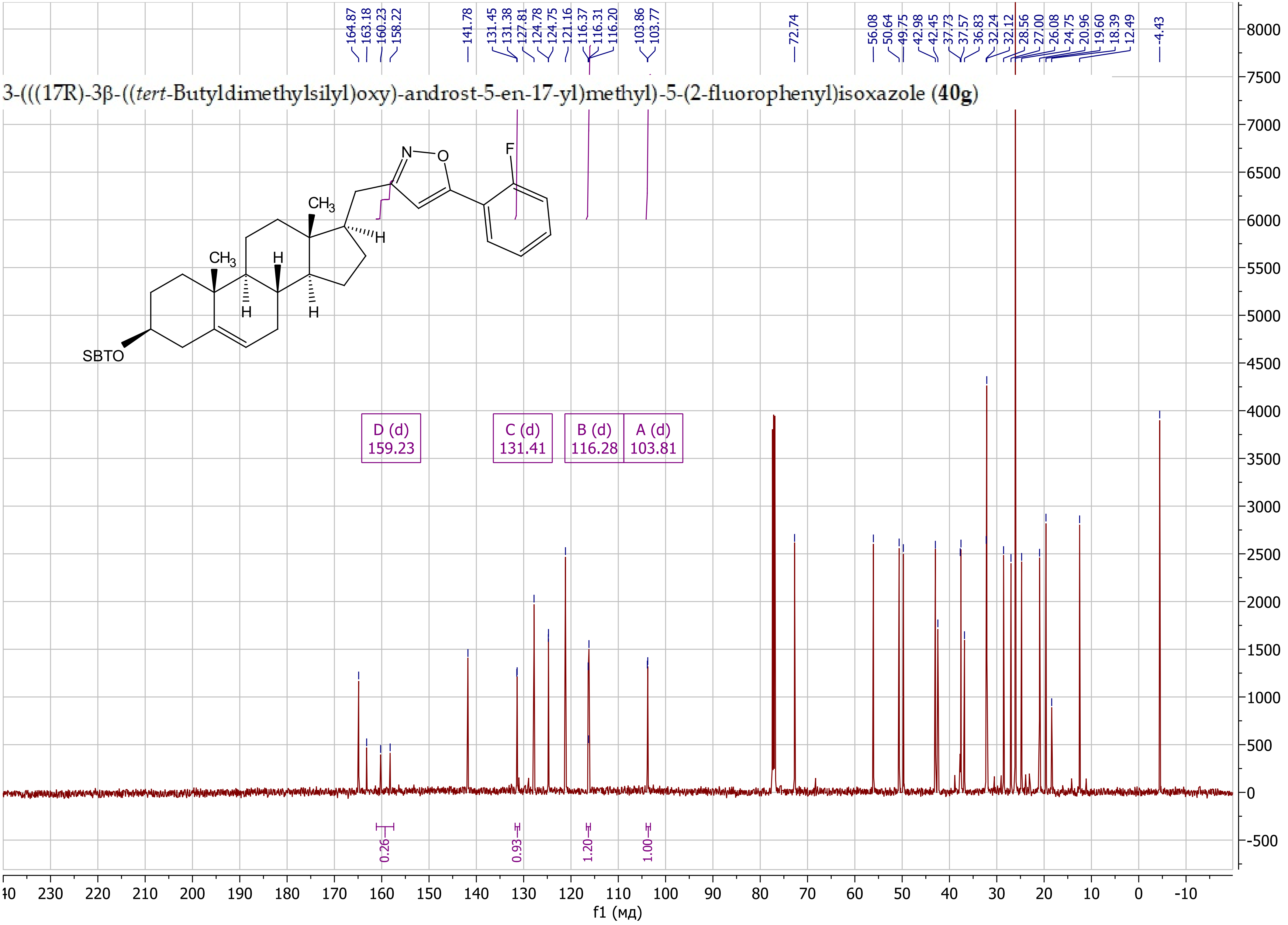

3-(((17*R*)-3β-((*tert*-Butyldimethylsilyl)oxy)-androst-5-en-17-yl)methyl)-5-(2-((tetrahydro-2*H*-pyran-2-yl)oxy)propan-2-yl)isoxazole (**40h**)

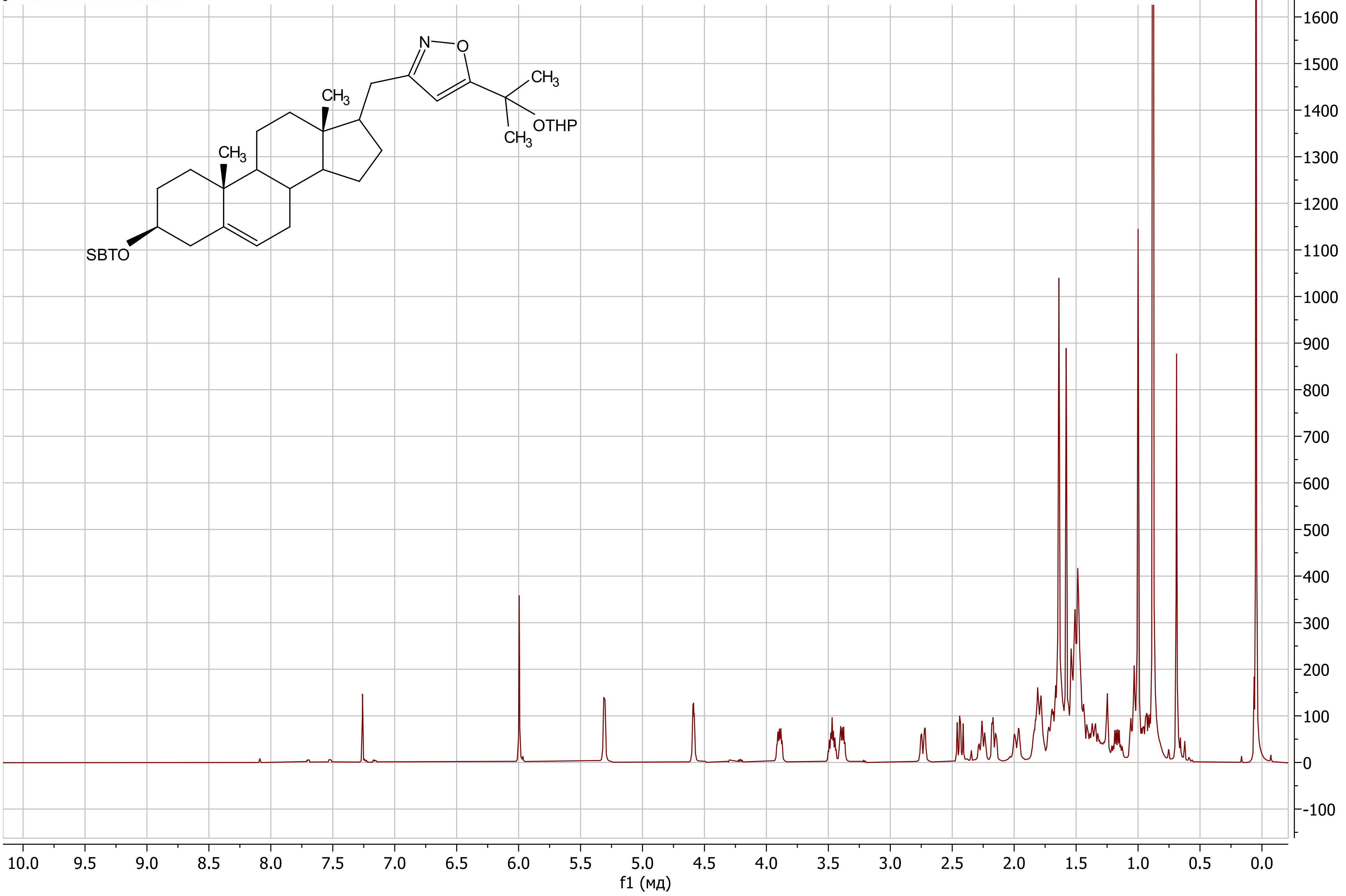

##### 3-(((17*R*)-3β-((*tert*-Butyldimethylsilyl)oxy)-androst-5-en-17-yl)methyl)-5-(2-((tetrahydro-2*H*-pyran-2-yl)oxy)propan-2-yl)isoxazole (**40h**)

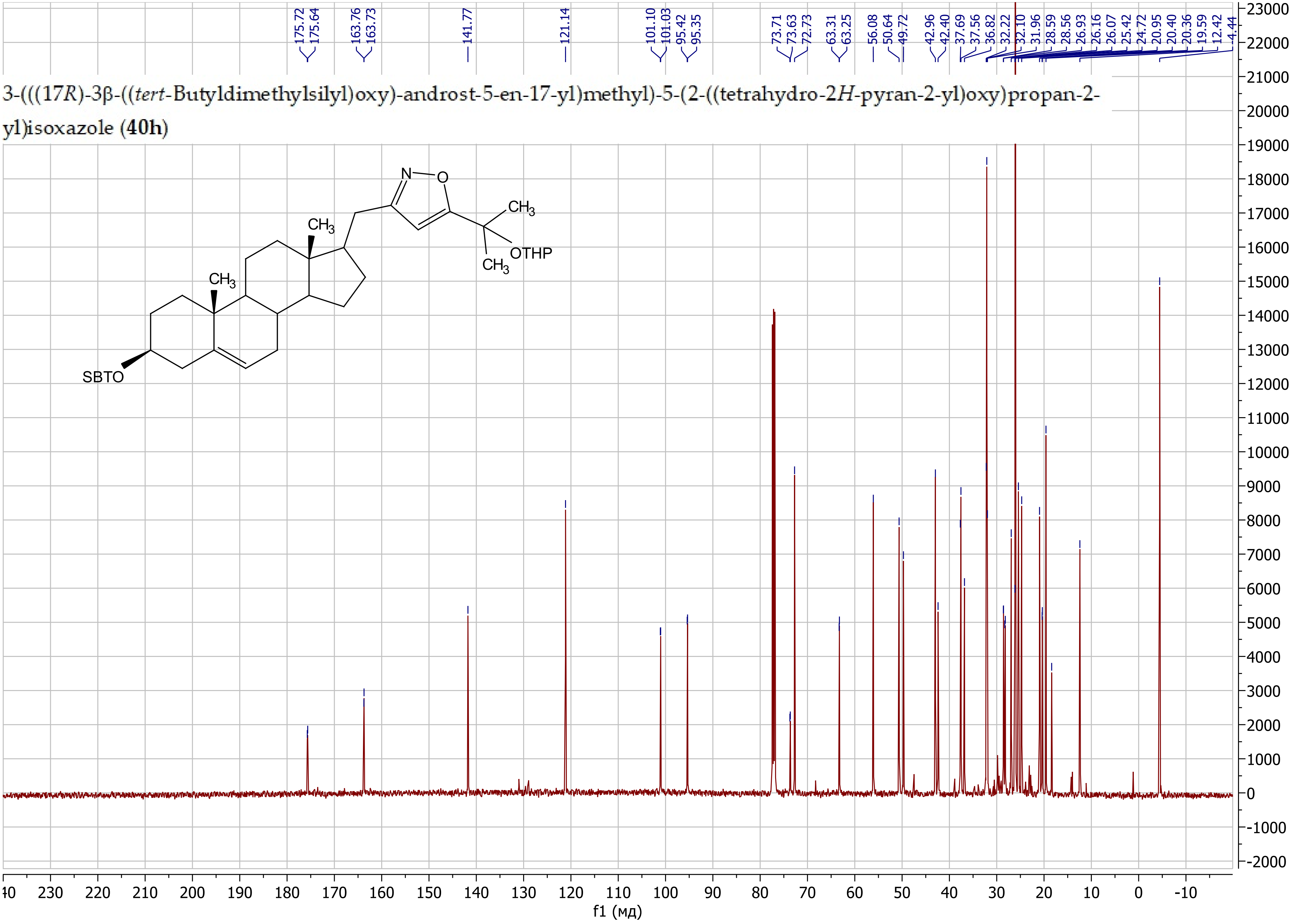

3-(((17*R*)-3β-((*tert*-Butyldimethylsilyl)oxy)-androst-5-en-17-yl)methyl)-5-(((tetrahydro-2*H*-pyran-2-yl)oxy)methyl)isoxazole (**40i**)

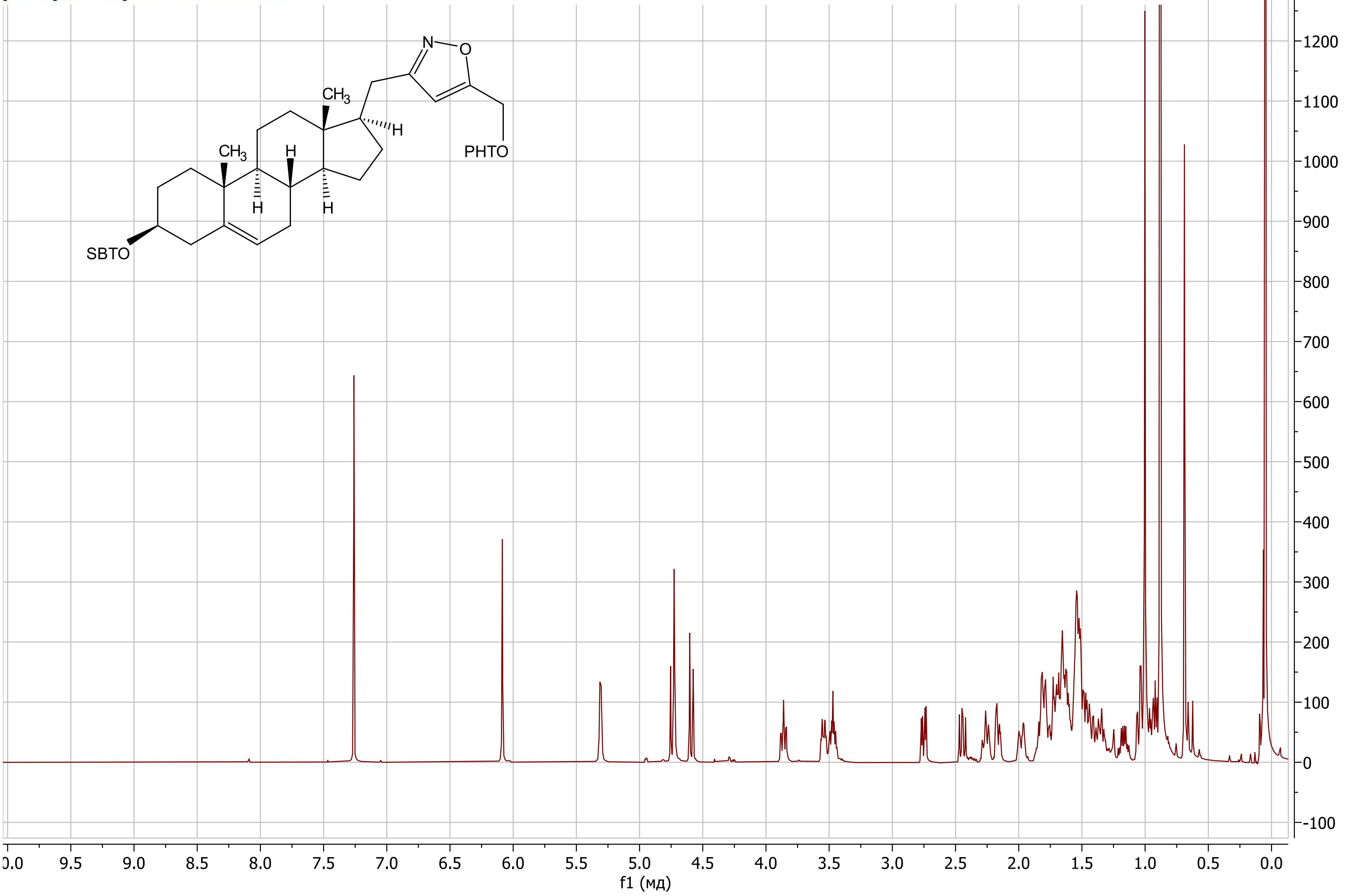

3-(((17*R*)-3β-((*tert*-Butyldimethylsilyl)oxy)-androst-5-en-17-yl)methyl)-5-(((tetrahydro-2*H*-pyran-2-yl)oxy)methyl)isoxazole (**40i**)

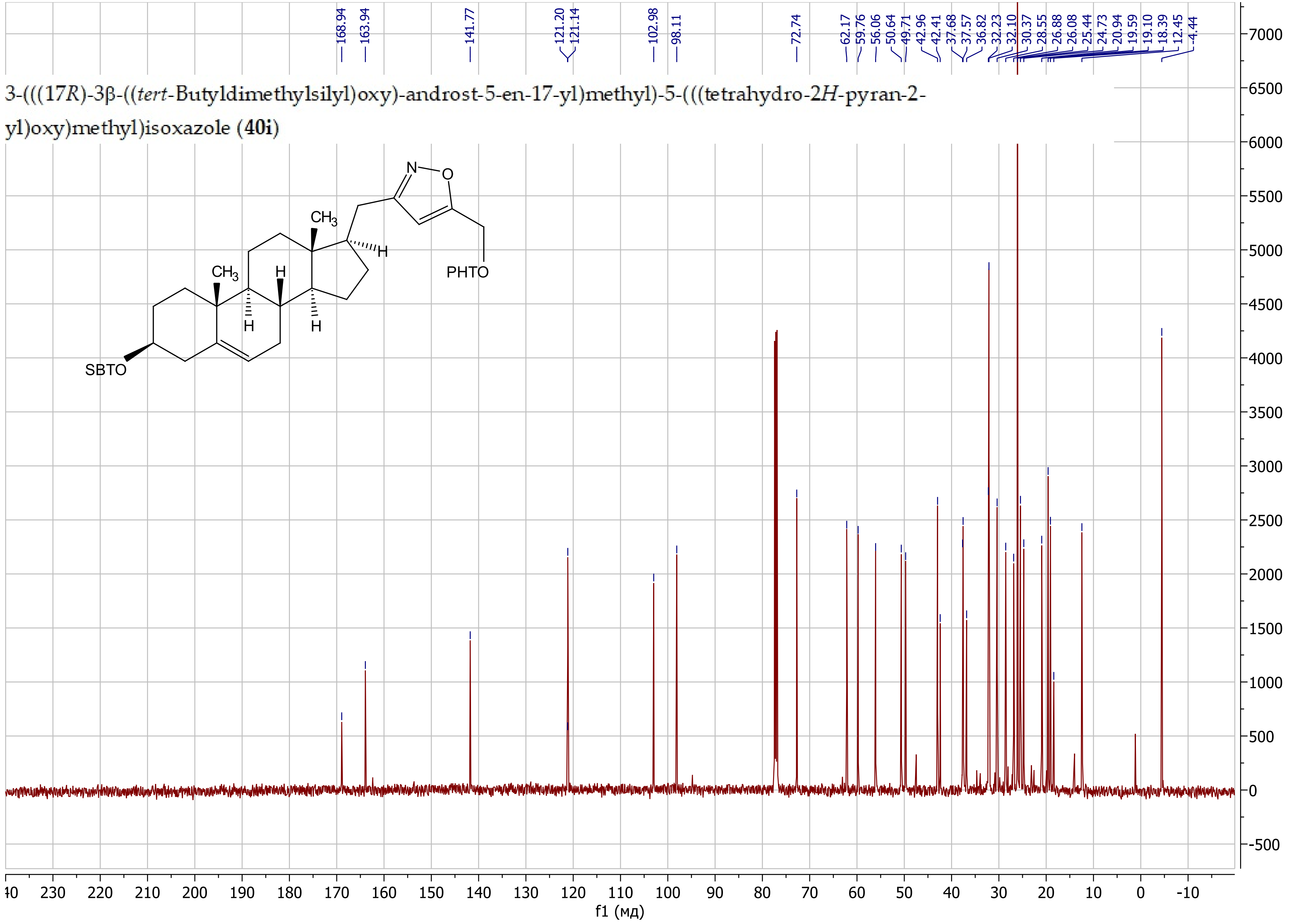

(17*R*)-17-(Isoxazol-3-ylmethyl)-androst-5-en-3β-ol (**41a**)

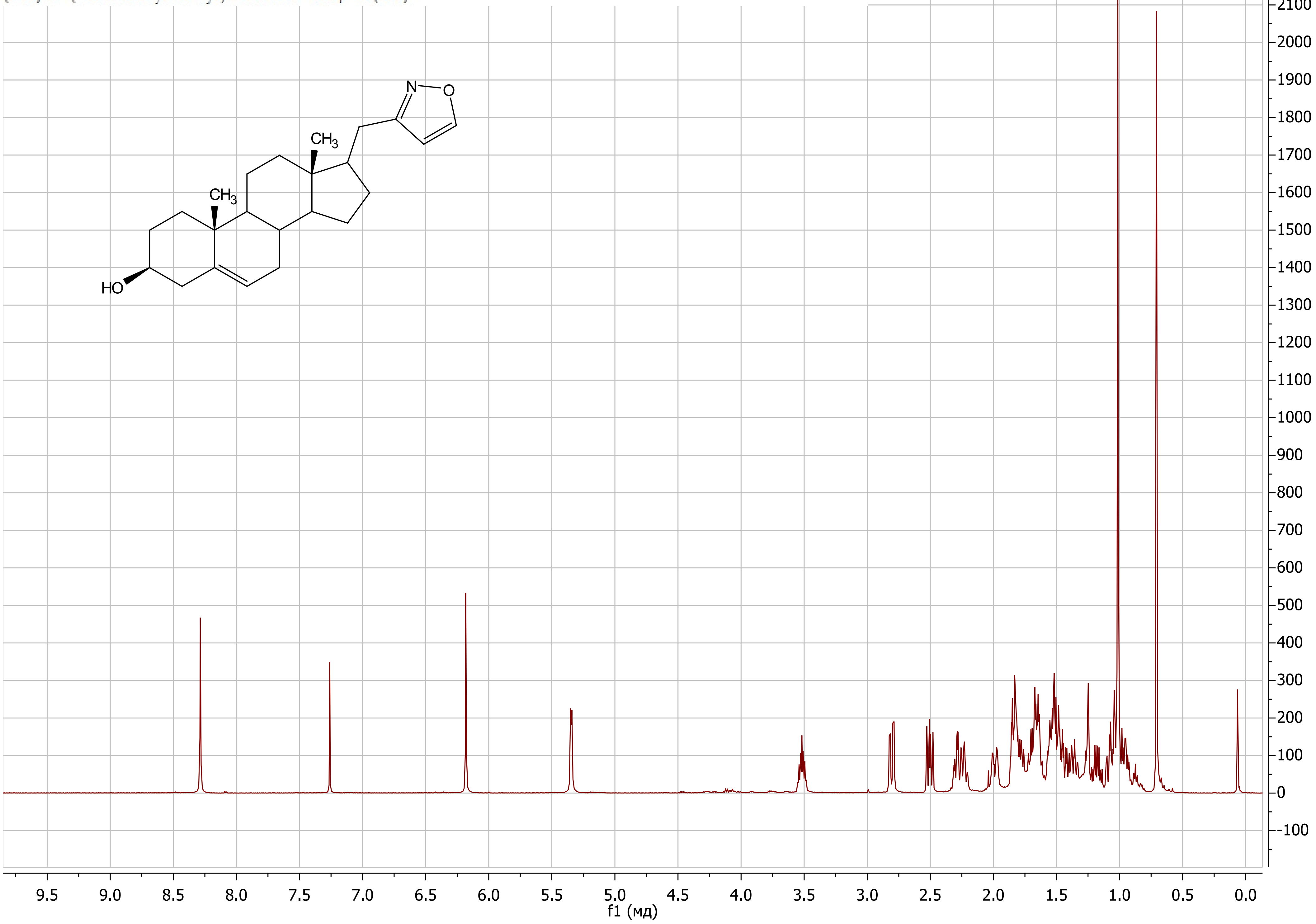

(17*R*)-17-(Isoxazol-3-ylmethyl)-androst-5-en-3β-ol (**41a**)

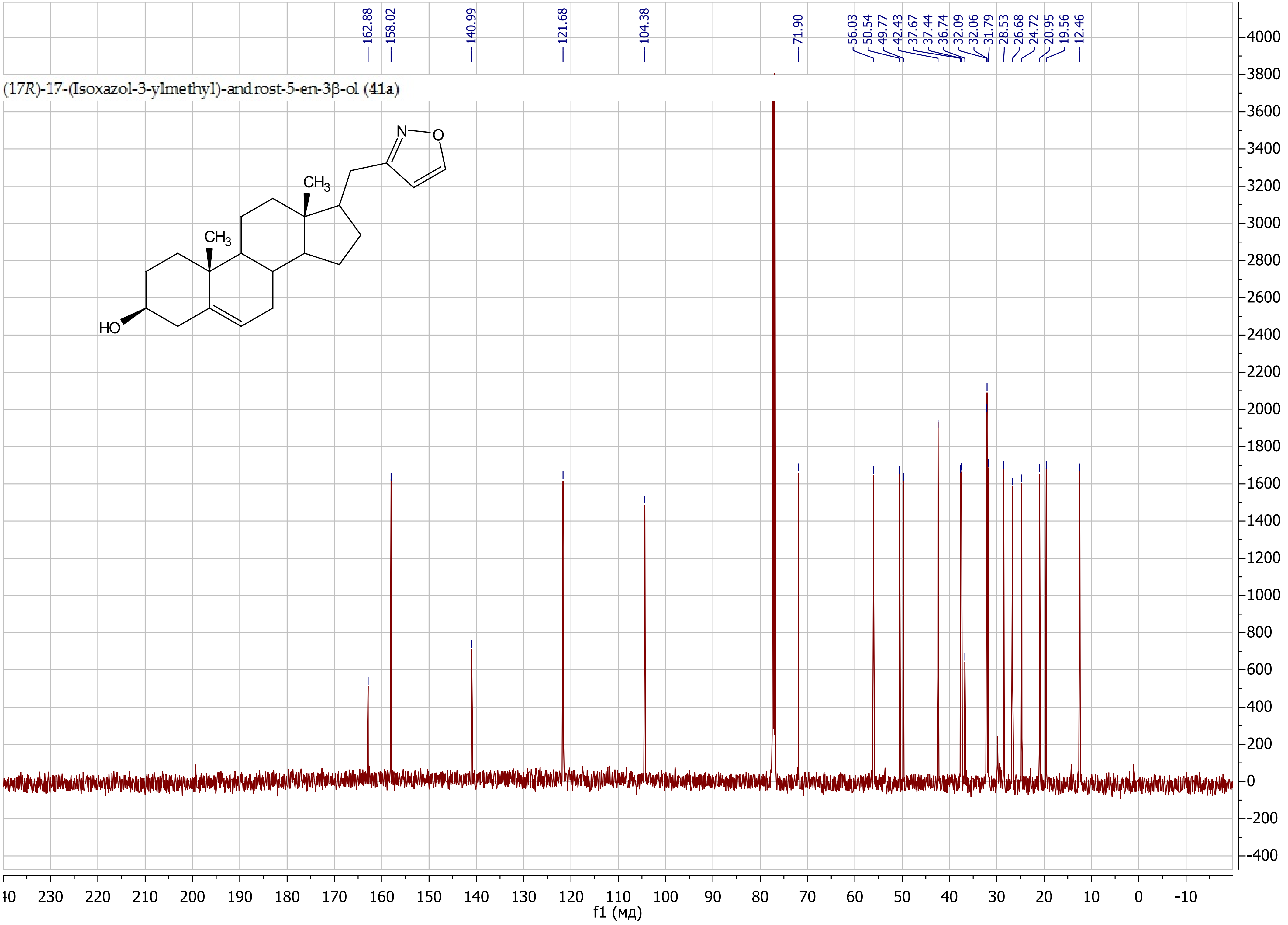

(17*R*)-17-((5-Butylisoxazol-3-yl)methyl)-androst-5-en-3β-ol (**41d**)

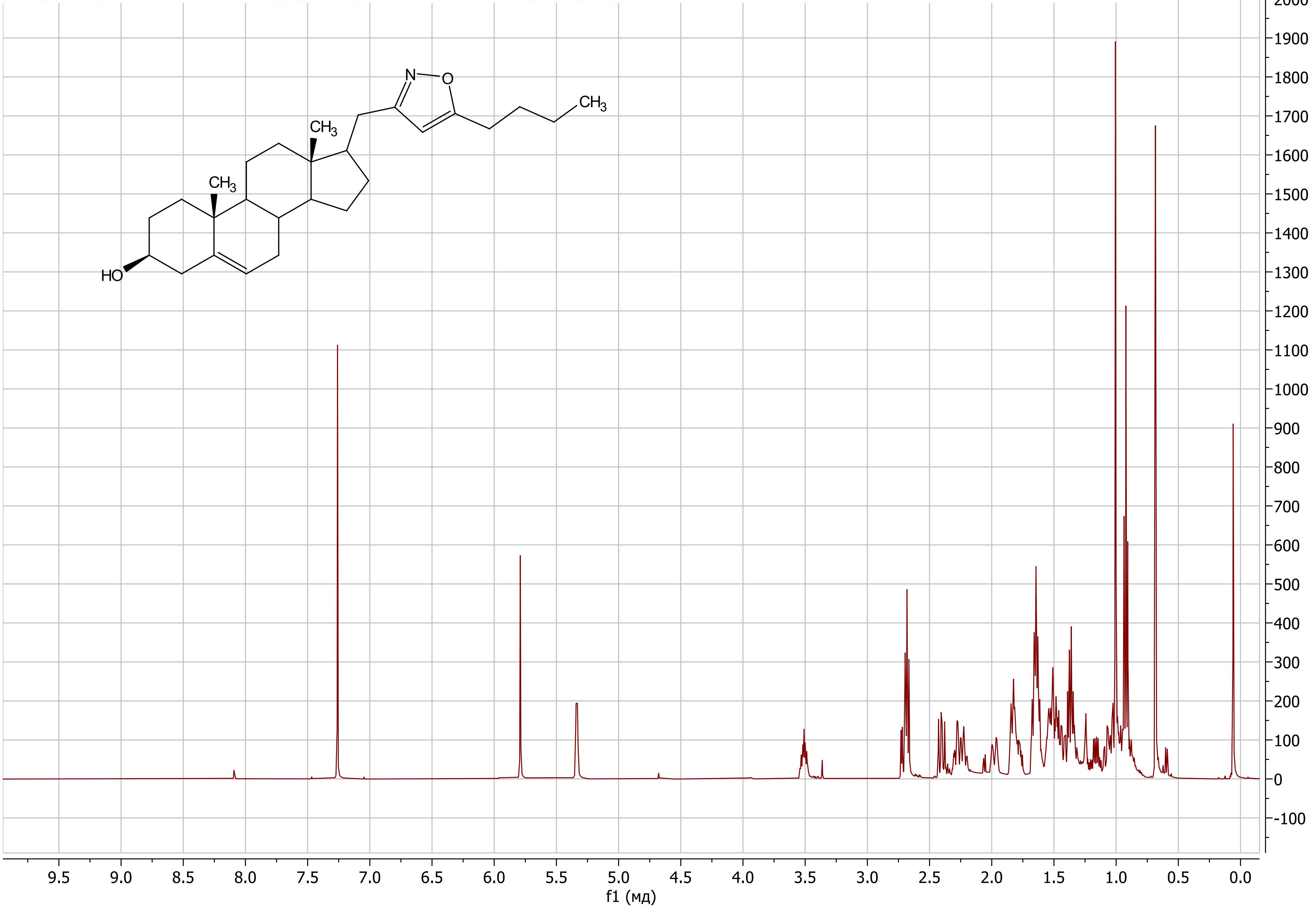

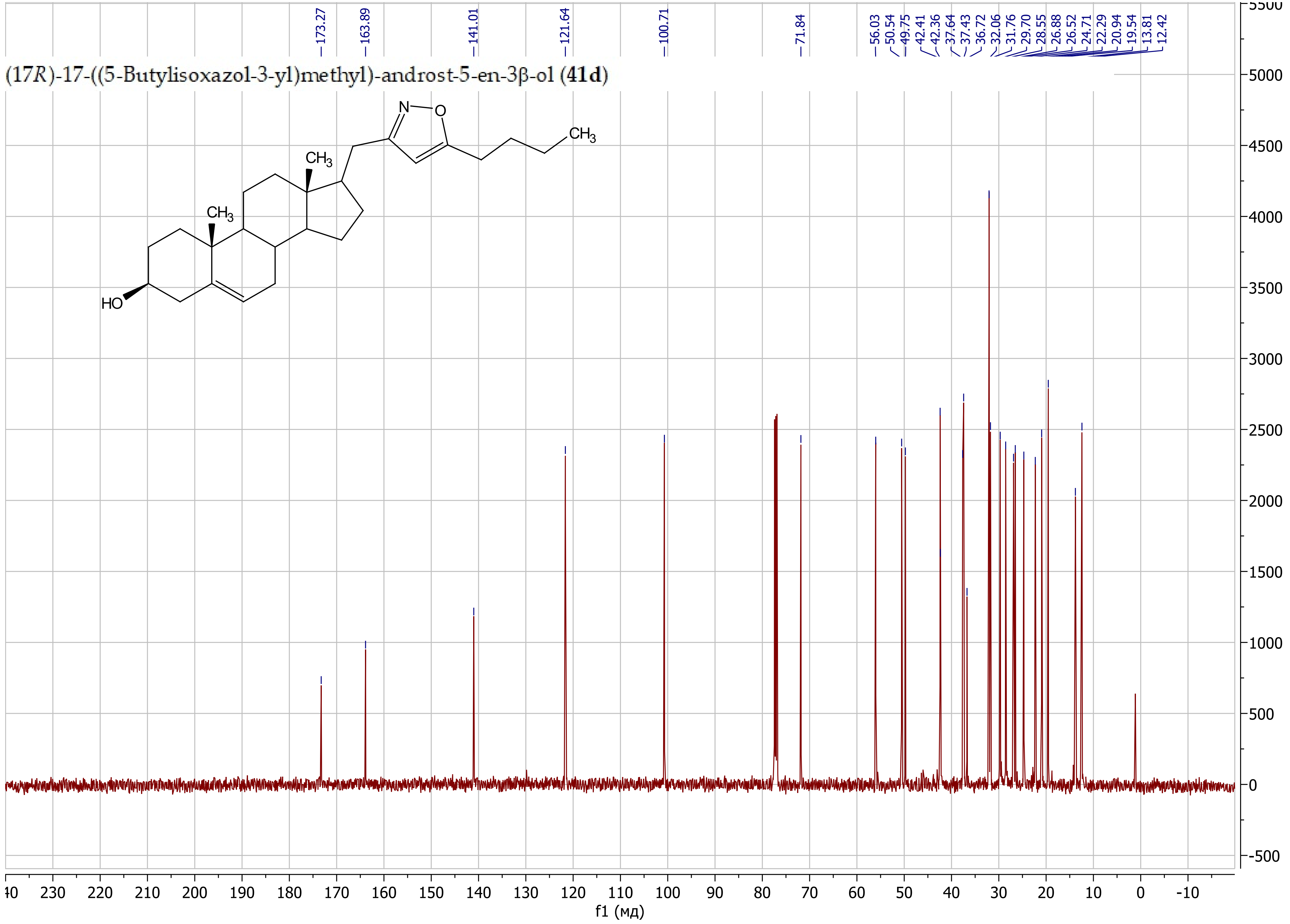
(17*R*)-17-((5-Butylisoxazol-3-yl)methyl)-androst-5-en-3β-ol (**41d**)
173.27
163.89
141.01
121.64
100.71
71.84
56.03
50.54
49.75
42.41
42.36
37.64
37.43
36.72
32.06
31.76
29.70
28.55
26.88
26.52
24.71
22.29
20.94
19.54
13.81
12.42
N
O
CH3
CH3
CH3
HO
f1 (мд)

(17*R*)-17-((5-Phenylisoxazol-3-yl)methyl)-androst-5-en-3β-ol (**41e**)

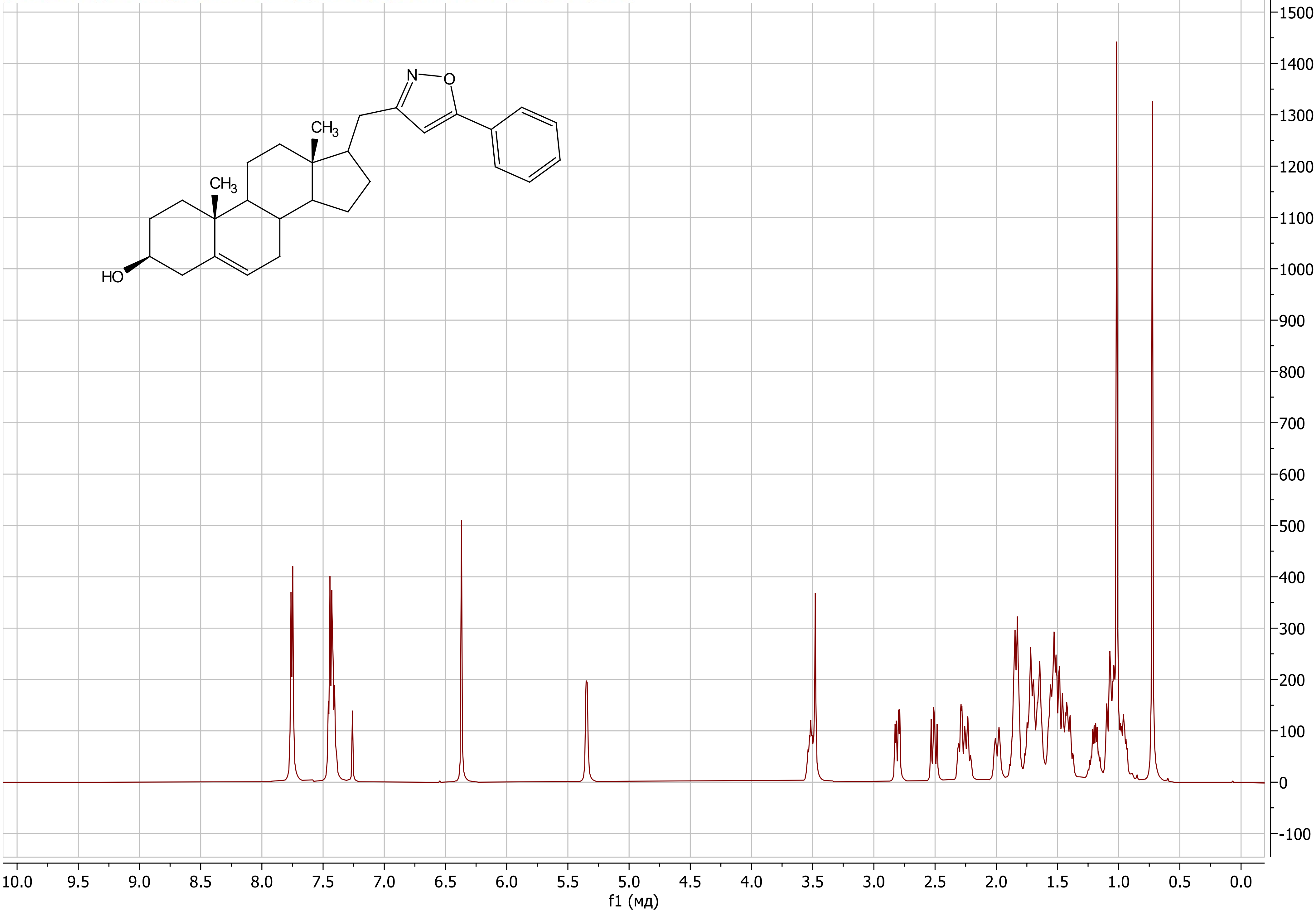

(17*R*)-17-((5-Phenylisoxazol-3-yl)methyl)-androst-5-en-3β-ol (**41e**)

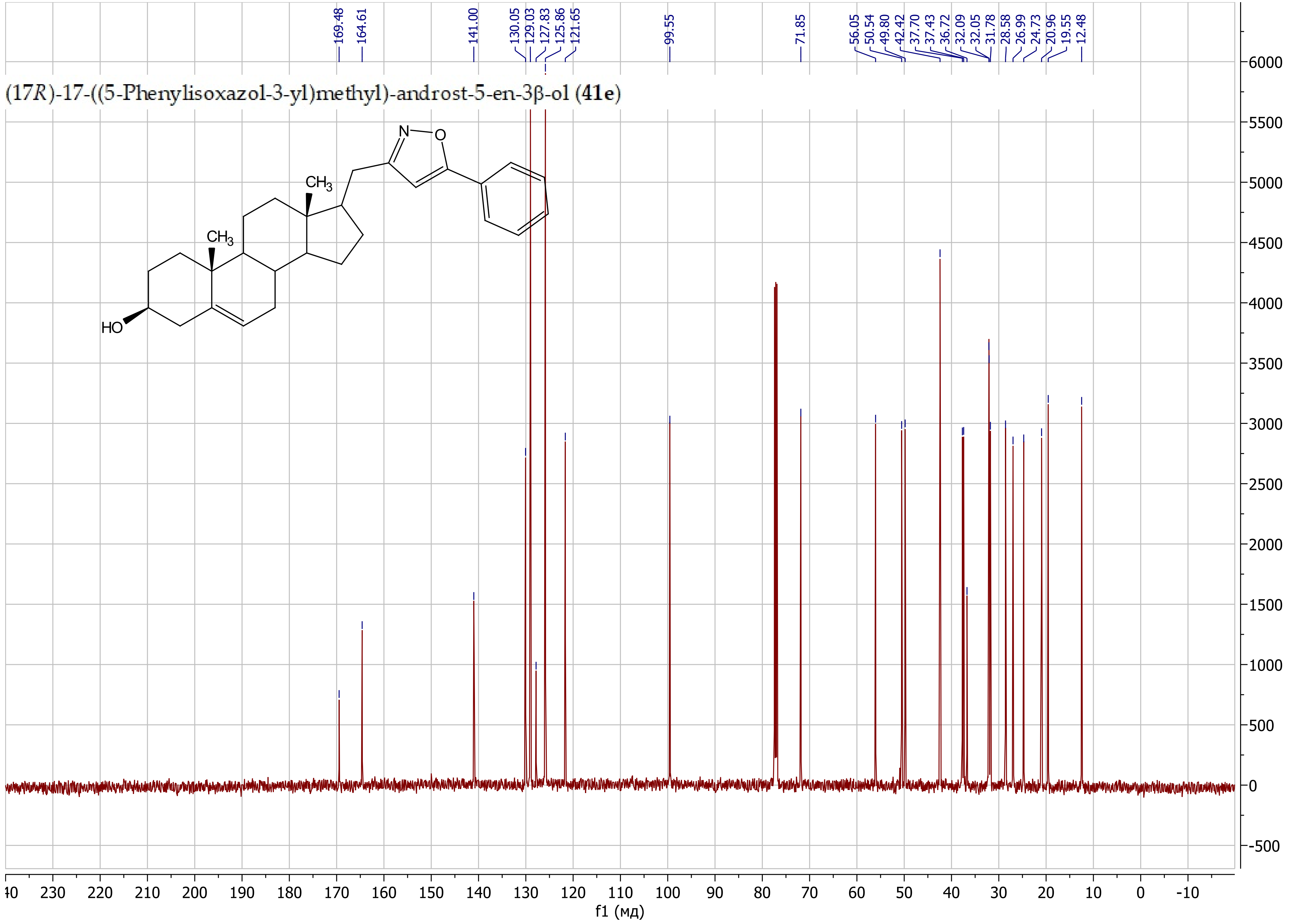

(17*R*)-17-((5-(Pyridin-3-yl)isoxazol-3-yl)methyl)-androst-5-en-3β-ol (**41f**)

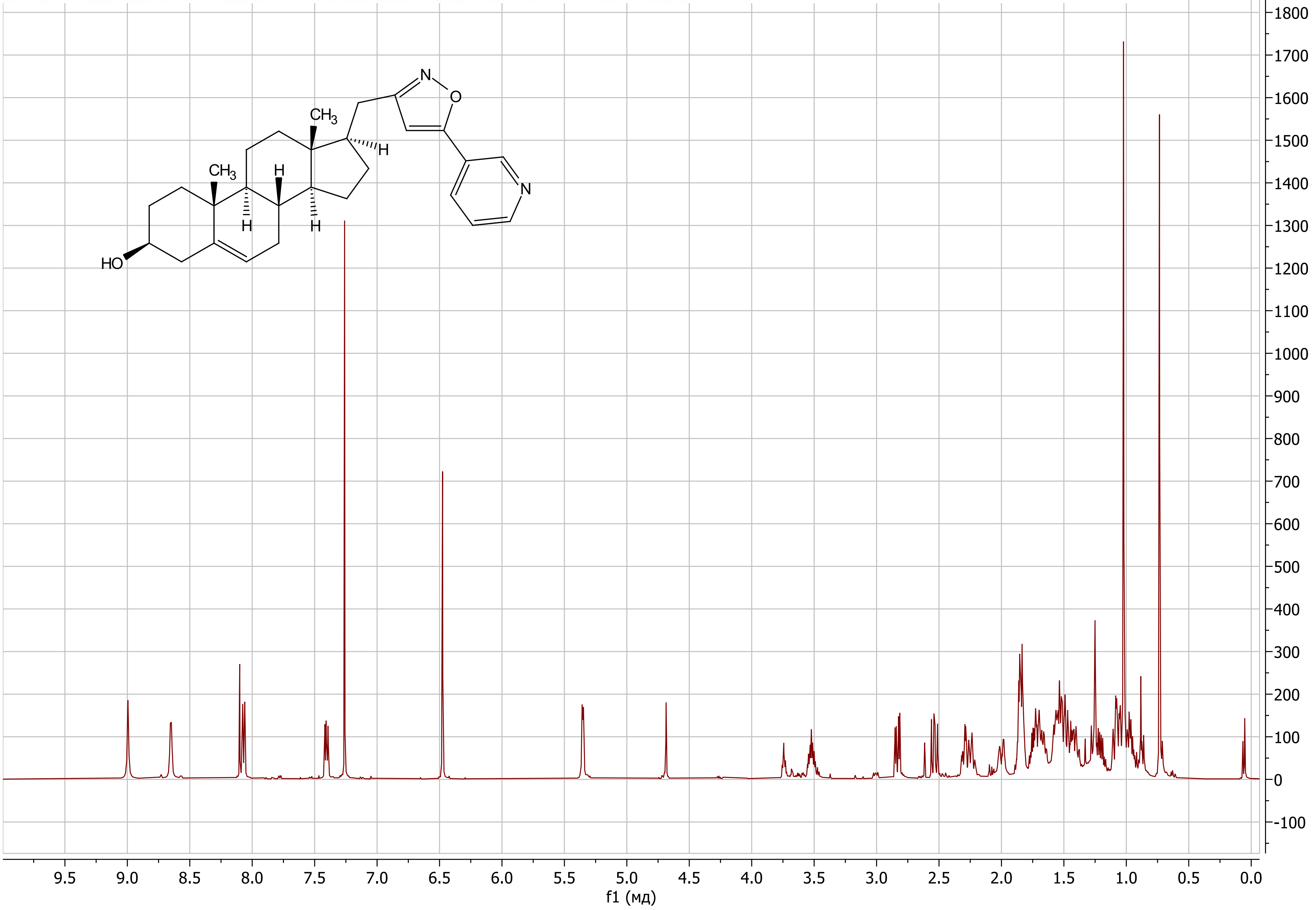

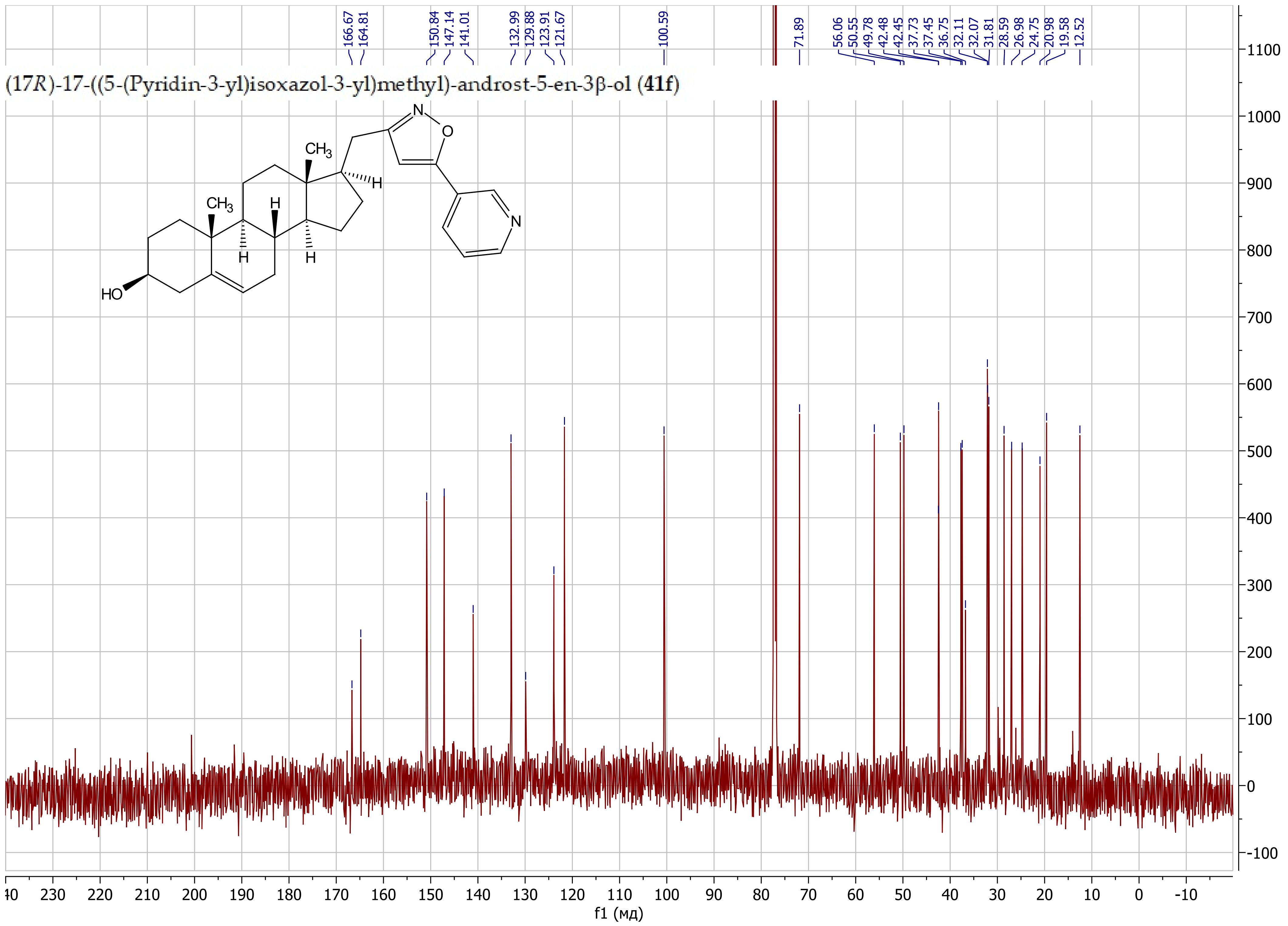

(17R)-17-((5-(Pyridin-3-yl)isoxazol-3-yl)methyl)-androst-5-en-3β-ol (41f)
166.67
164.81
150.84
147.14
141.01
132.99
129.88
123.91
121.67
100.59
71.89
56.06
50.55
49.78
42.48
42.45
37.73
37.45
36.75
32.11
32.07
31.81
28.59
26.98
24.75
20.98
19.58
12.52
CH3
H
HO
N
O
f1 (мд)

(17*R*)-17-((5-(2-Fluorophenyl)isoxazol-3-yl)methyl)-androst-5-en-3β-ol (**41g**)

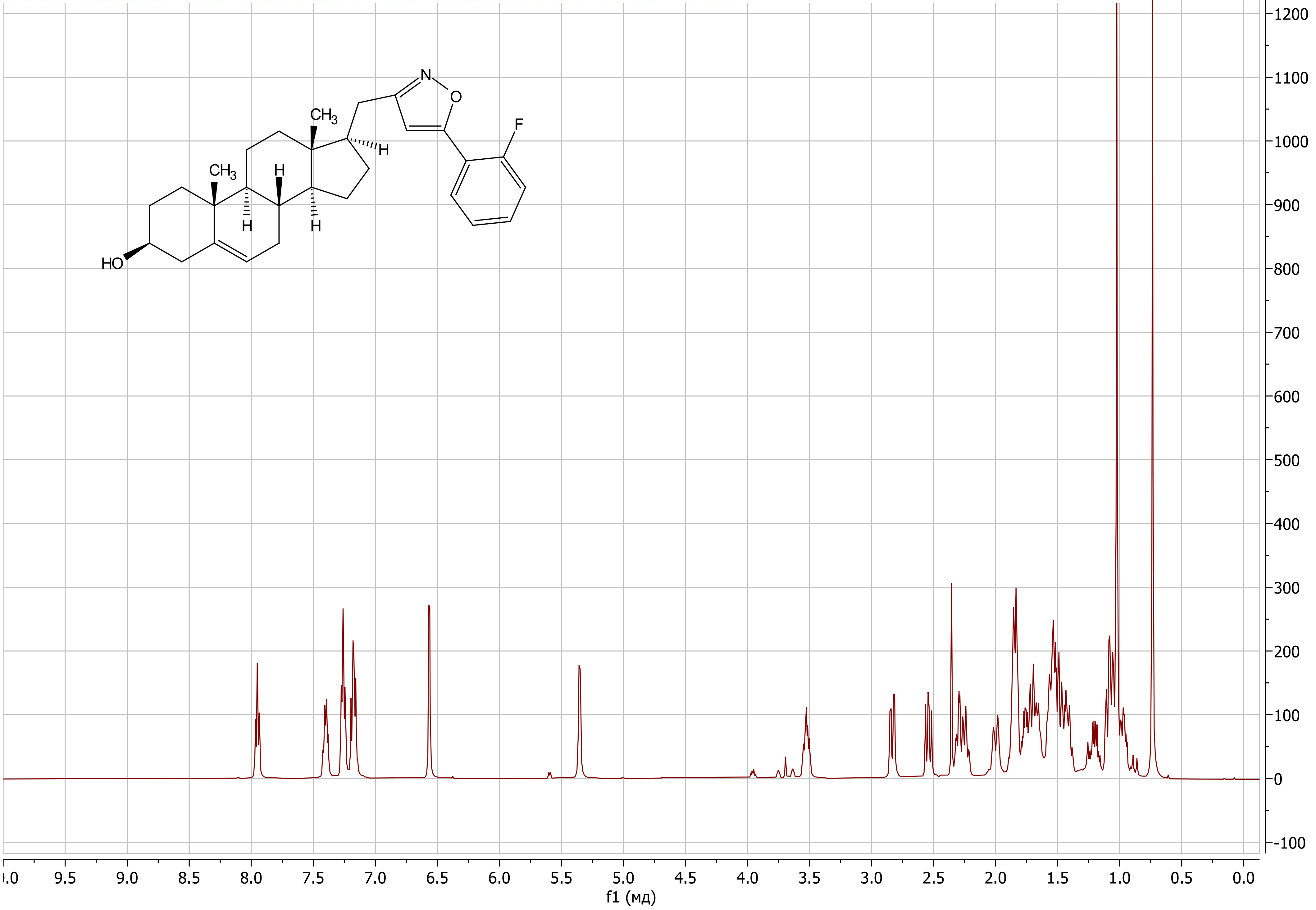

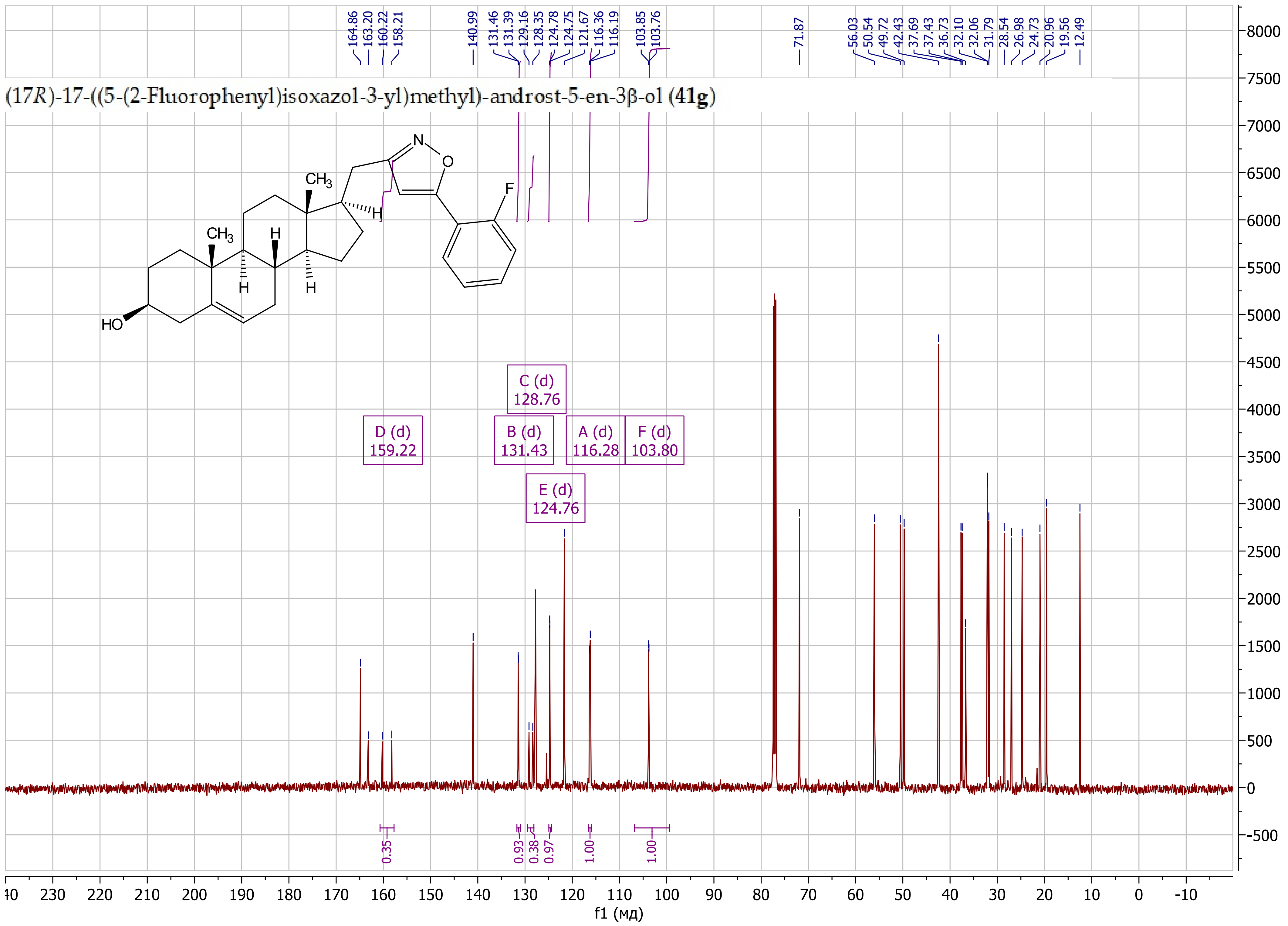

(17R)-17-((5-(2-Fluorophenyl)isoxazol-3-yl)methyl)-androst-5-en-3β-ol (41g)
164.86
163.20
160.22
158.21
140.99
131.46
131.39
129.16
128.35
124.78
124.75
121.67
116.36
116.19
103.85
103.76
71.87
56.03
50.54
49.72
42.43
37.69
37.43
36.73
32.10
32.06
31.79
28.54
26.98
24.73
20.96
19.56
12.49
C (d) 128.76
D (d) 159.22
B (d) 131.43
A (d) 116.28
F (d) 103.80
E (d) 124.76
0.35
0.93
0.38
0.97
1.00
1.00
f1 (мд)

(17*R*)-17-((5-(2-Hydroxypropan-2-yl)isoxazol-3-yl)methyl)-androst-5-en-3β-ol (**41j**)

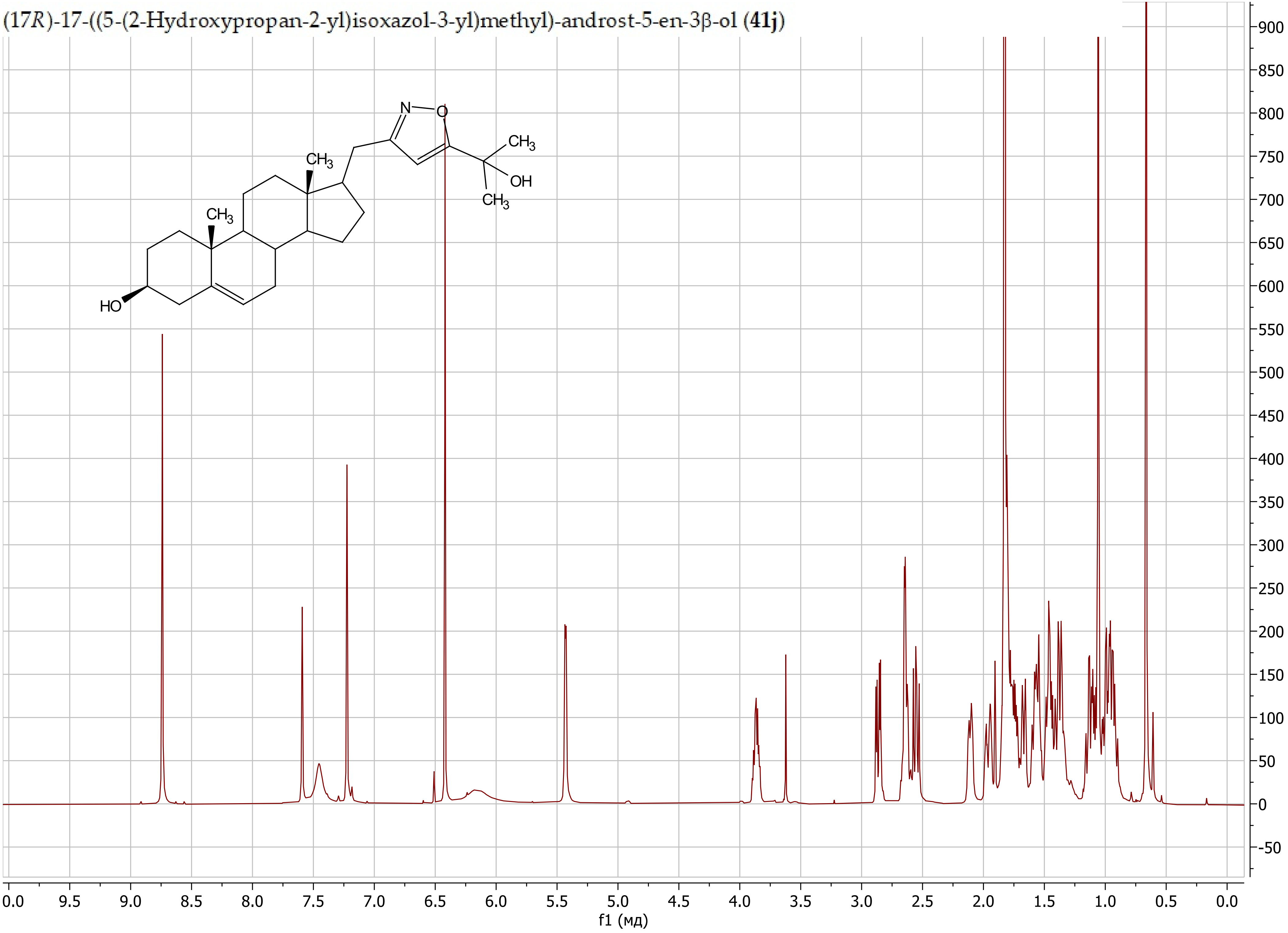

(17*R*)-17-((5-(2-Hydroxypropan-2-yl)isoxazol-3-yl)methyl)-androst-5-en-3β-ol (**41j**)

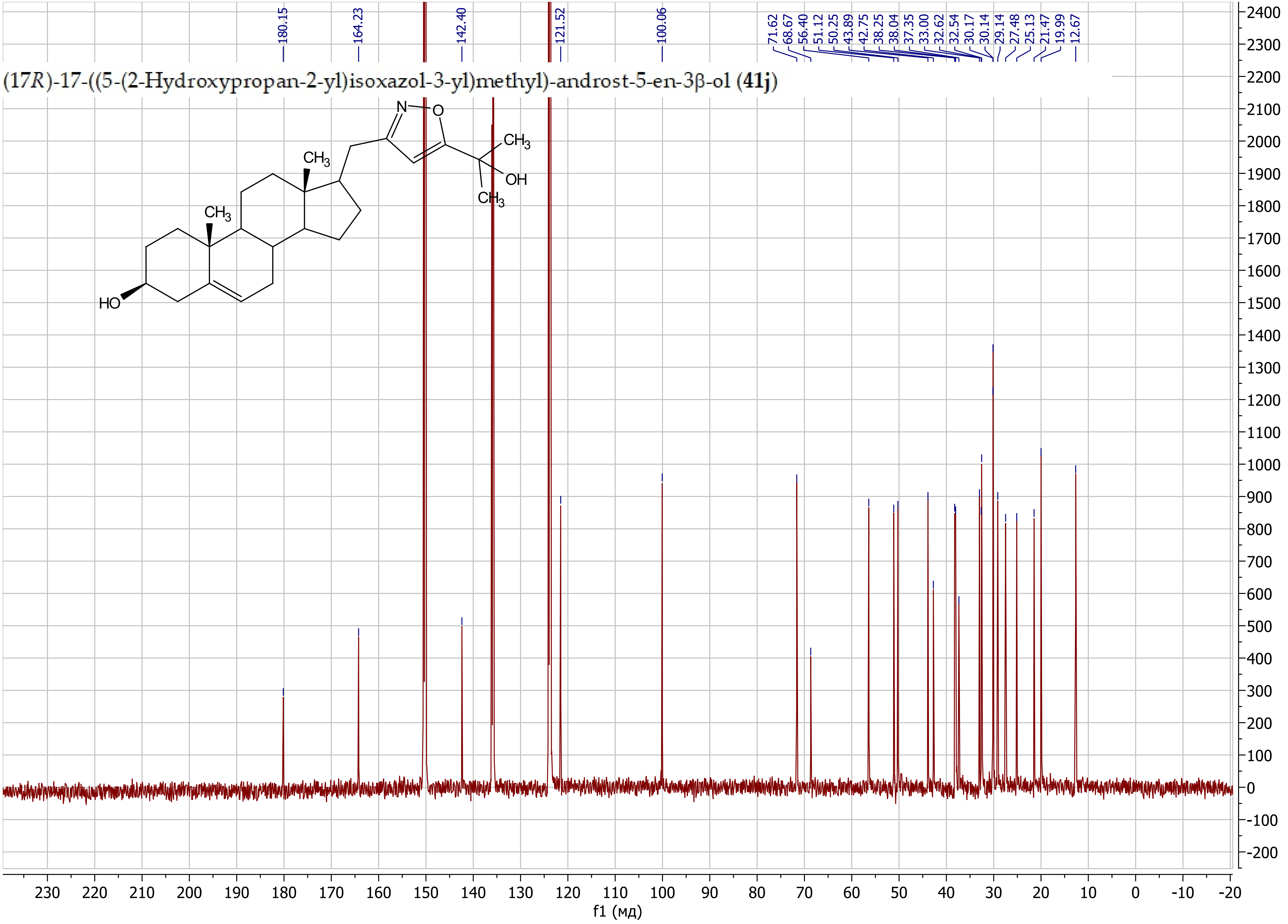

(17*R*)-17-((5-(Hydroxymethyl)isoxazol-3-yl)methyl)-androst-5-en-3β-ol (**41k**)

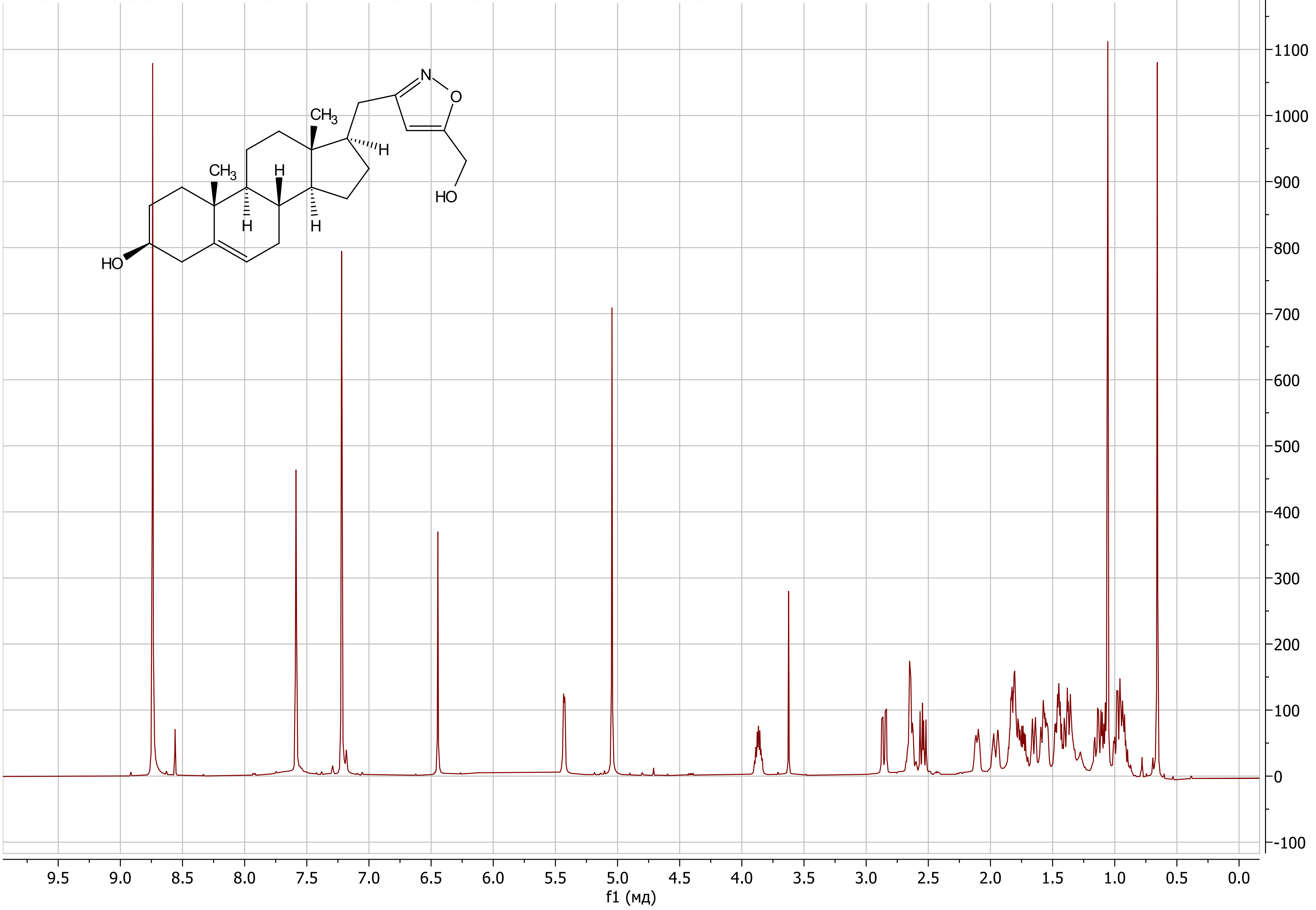

(17*R*)-17-((5-(Hydroxymethyl)isoxazol-3-yl)methyl)-androst-5-en-3β-ol (**41k**)

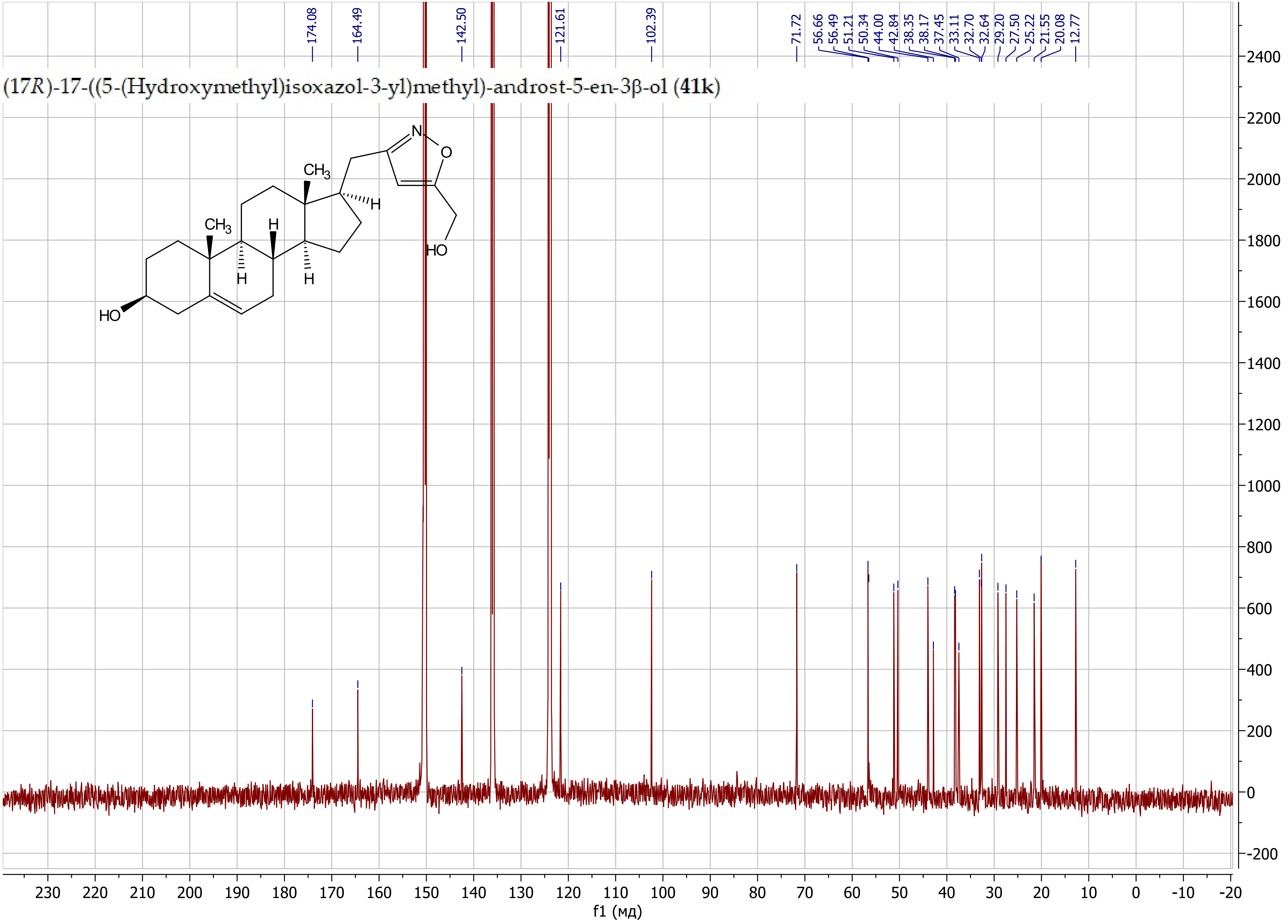
